# Decoding the lethal effect of *SARS-CoV-2* (novel coronavirus) strains from global perspective: molecular pathogenesis and evolutionary divergence

**DOI:** 10.1101/2020.04.06.027854

**Authors:** Shuvam Banerjee, Shrinjana Dhar, Sandip Bhattacharjee, Pritha Bhattacharjee

## Abstract

**Background:** COVID-19 is a disease with global public health emergency that have shook the world since its’ first detection in China in December, 2019. Severe acute respiratory syndrome Coronavirus 2 (*SARS-CoV-2*) is the pathogen responsible behind this pandemic. The lethality of different viral strains is found to vary in different geographical locations but the molecular mechanism is yet to be known.

**Methods:** Available data of whole genome sequencing of different viral strains published by different countries were retrieved and then analysed using Multiple Sequence Alignment and Pair-wise Sequence Alignment leading to Phylogenetic tree construction. Each location and the corresponding genetic variations were screened in depth. Then the variations are analysed at protein level giving special emphasis on Non Synonymous amino acid substitutions. The fatality rates in different countries were matched against the mutation number, rarity of the nucleotide alterations and functional impact of the Non Synonymous changes at protein level, separately and in combination.

**Findings:** All the viral strains have been found to evolve from the viral strain of Taiwan (MT192759) which is 100% identical with the ancestor *SARS-CoV-2* sequences of Wuhan (NC 045512.2; submitted on 5^th^ Jan, 2020). Transition from C to T (C>T) is the most frequent mutation in this viral genome and mutations A>T, G>A, T>A are the rarest ones, found in countries with maximum fatality rate i.e Italy, Spain and Sweden. 20 Non Synonymous mutations are located in viral genome spanning Orf1ab polyprotein, Surface glycoprotein, Nucleocapsid protein etc. The functional effect on the structure and function of the protein can favourably or unfavourably interact with the host body.

**Interpretation:** The fatality outcome depends on three important factors (a) number of mutation (b) rarity of the allelic variation and (c) functional consequence of the mutation at protein level. The molecular divergence, evolved from the ancestral strain (S) lead to extremely lethal (E), lethal(L) and non lethal (N) strains with the involvement of an Intermediate strain(I).

## Introduction

In the new decade of 21^st^ century, surfaced the first public health emergency of global concern in Wuhan, China from the menace of COVID-19 (*nCoV/*β*-coronavirus*), also known as *SARS-CoV-2*. COVID-19 is one of the seven pathogenic members of family *coronaviridae*, other notable ones are severe acute respiratory syndrome (SARS) coronavirus (*SARS-CoV*), identified first in southern China (November, 2002), and Middle East respiratory syndrome (MERS) coronavirus (*MERS-CoV*), from Saudi Arabia (2012). Less severe ones are *HKU1, NL63, OC-43* and *229E*.^1^ The enveloped virus possesses crown-like spikes on their surface (Latin word *corona* i.e crown) and a RNA genome, single stranded positive-sense strand with 26 to 32 kb length.^2^ The microbe can co-infect different vertebrates including humans and affect organs of respiratory, gastrointestinal and central nervous system.

According to World Health Organization (WHO), 80% COVID infection are mild or asymptomatic, 15% are severe infection, while 5% are critical. Crude mortality ratio (no. of reported deaths divided by reported cases) is between 3-4%. The infection mortality rate (the number of reported deaths divided by the number of infection) will be lower than seasonal influenza (0·1%). Globally there are 6,68,690 confirmed cases; 31,065 deaths and so far 1,43,107 people recovered in 199 countries and territories.^3,4^

The most vulnerable group in corona epidemic is elderly, malnourished, hypertensive, diabetic, immune compromised, cancer and cardiovascular patients, as well as pregnant women. Despite wide spread COVID-19 infection, the numbers of fatally infected children cases are less reported so far, possible mechanism might be an unknown protective interaction between immune system and respiratory pathway.^5^ COVID-19 spreads through droplet infection and fomites, from infected person during coughing and sneezing. Majority of symptoms are sore throat, breathing difficulty and fever, although GI and musculoskeletal system are also involved.^6,7^ Clinical course is somehow showing a predictable pattern. On around day five, flu like symptom starts, common are fever, headache, dry cough, myalgia (and back pain), nausea (without vomiting), abdominal discomfort with some diarrhea, loss of smell, anorexia and fatigue. Around same time symptoms can worsen leading to shortness of breath due to bilateral viral pneumonia from direct viral damage to lung parenchyma. Around day10, cytokine storm kicks in, subsequently acute respiratory distress syndrome (ARDS) and multi organ failure ensues. This degradation usually happens in matter of hours. Hospitalized patients in moderate to severe cases usually come in hypoxic stage without dyspnea. China reported 15% cardiac involvement and serious final outcome. In most of the chest X-rays, bilateral interstitial pneumonia or ground glass opacities are seen. Hypoxia mostly does not correlate well with the chest x-ray findings. Chest auscultations are not of great help. Blood reports show, in the most cases, WBC is low, i.e. mostly lymphocytopenia, low platelet count whereas pro-calcitonin are mostly normal. Most consistently, CRP and ferritin levels are elevated and similarly CPK, D-dimer, LDH, ALK-phos/AST-ALT levels are also high. It has been seen a ratio of absolute neutrophil count to absolute lymphocyte count, greater than 3.5, may be the highest predictor of poor outcome. An elevated level of IL-6 is of grave concern of cytokine storm.^8,9^

Among the four classified subfamilies, α- and β-coronaviruses both were found in bats, e.g Leaf-nose bats like *Hipposideros armige* harbor *α-coronavirus* and *β-coronavirus* in *Hipposideros larvatus*;^10^ while *γ-* and *δ-coronavirus* found in pigs and birds respectively.^11^ The genesis of *β-coronavirus* cluster in afore mentioned bat species, consisting of primeval and genetically independent types might took favour of host/pathogen interaction despite of wide geographical distribution.^10^ Genomic characterization of *SARS-CoV-2* reveals 96% identity to that of bat coronavirus whole genome;^1^ while 79% and 50% to that of *SARS-CoV* and MERS-CoV respectively.^12^ Like other betacoronaviruses, the genome of *SARS-CoV-2* has a long ORF1ab polyprotein at the 5’ end, followed by four major structural proteins, including the spike like surface glycoprotein, small envelope protein, matrix protein, and nucleocapsid protein. ^11,13^ The spike (S) protein has two distinct functional domains, termed S1 and S2, both of which are necessary for a coronavirus to successfully enter a cell. It has been found that S protein of *SARS-CoV-2* is 10-20 times more likely to bind to human ACE2 than the S protein of the early 2000s *SARS-CoV* strain. ACE2 is essentially a carboxypeptidase, which can remove carboxy-terminal of hydrophobic or basic amino acids. The expression of ACE2 is comparatively higher in mouth and tongue and is also normally expressed in human lower lungs on type-I and type-II alveolar epithelial cells. The heightened affinity for a prevalent cellular receptor may be a factor which increases a quick transmission of *SARS-CoV-2* in the upper respiratory tract. As an RNA virus, 2019-nCoV has the inherent feature of a high mutation rate. Due to mutations and recombination effects, different viral strains are originating with new characteristics; however because of its genome encoded exonuclease, the mutation rate might be somewhat lower than other RNA viruses.^12,14^

In this study, we comprehensively analyzed the whole genome sequence homology from the available patient data uploaded by affected countries in NCBI Virus database, identified the mutations developed by different strains from the ancestor strain and studied the impact of those mutations at functional level. Our endeavour was to categorize all the strains into major groups depending upon their lethal effect and mutations observed.

## Methodology

### Whole genomic data retrieval from the database

Retrieved the whole genome sequences from “NCBI Virus” database, specific input was “*SARS-CoV-2*”, during a period from Jan 5 through March 24, 2020. Later submissions or the sequences with many undetermined nucleotides (denoted as N) were not considered.^15^ Countries those had multiple entries, highest number of entries with whole genome sequence were matched for identifying representative sequence to consider (e.g. for USA, out of 44 submitted complete genomic sequences, the sequence with 29 submissions were of complete genomic sequence with 100% identity and thus one representative sequence was considered for the present study). Thus, 13 complete whole genome sequences submitted by 13 different affected countries were taken into consideration (figure 1).

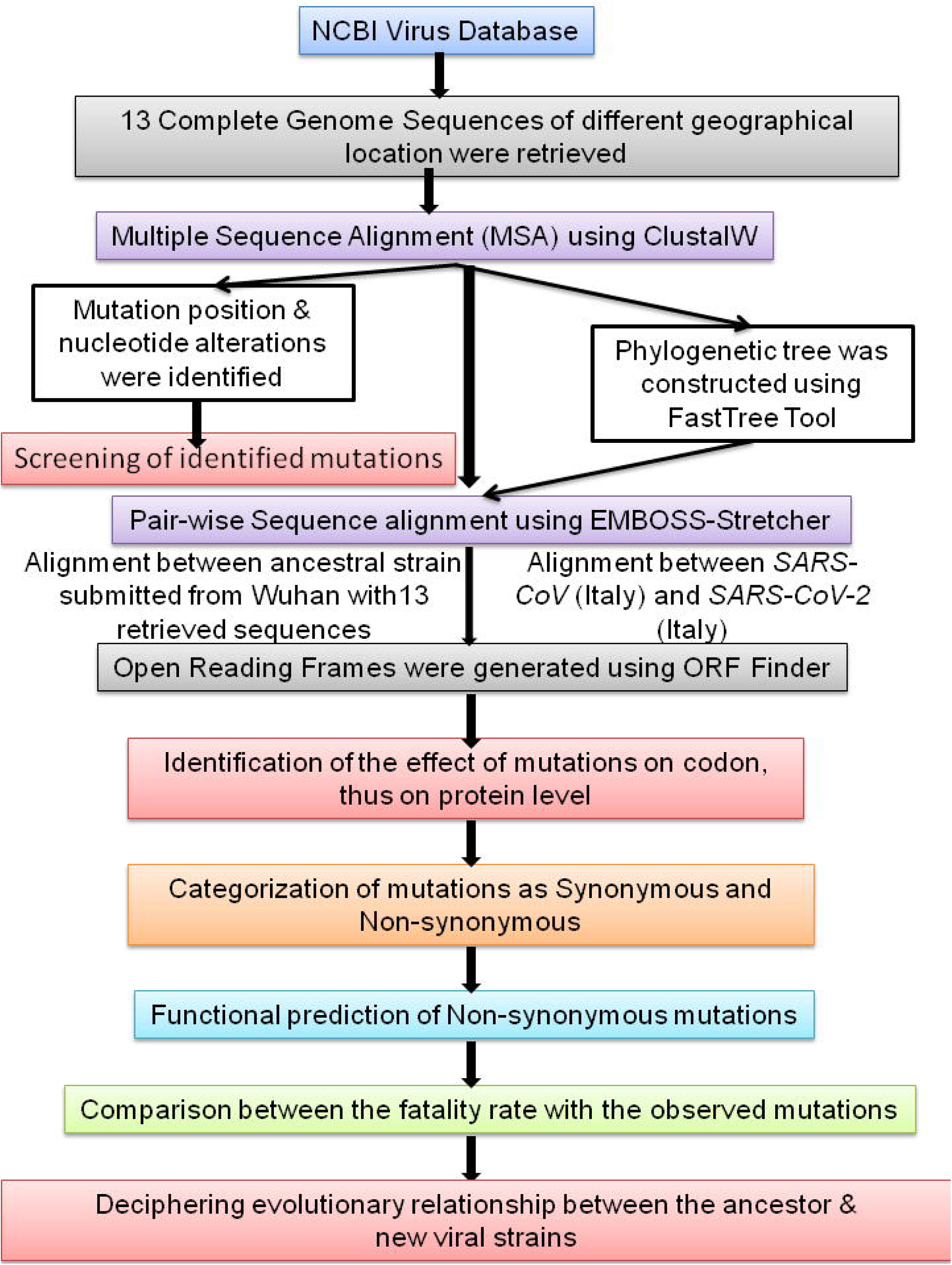

### Multiple Sequence Alignment and Evolutionary Dynamics

ClustalW (version 2.1) was employed to align multiple sequences, representative of 13 countries, with the purpose (a) to understand the similarity and variation in all genomic position, (b) to establish the evolutionary relationship among the viral strains affecting the whole world and (c) finally, to distinguish the ancestry of different viral strains using FastTree tool, which is based on Neighbour joining method.

### Pair-wise Sequence Alignment

EMBOSS-Stretcher was used for whole genomic pair-wise alignment between the ancestor *SARS-CoV-2* sequences of Wuhan (NC045512.2; submitted on 5^th^ Jan, 2020) with all 13 sequences. Similar strategy was used for pair wise alignment between the genome sequences of *SARS-CoV* submitted by Italy, 2003 with the *SARS-CoV-2* strain of Italy, 2020.

### Screening of Mutations

All alignment results were thoroughly screened to find out the exact location of the mutations and the genetic variation affecting that position. Functional implications of each type of nucleotide variations were understood from the LODS ratio and it is calculated by the formulae log_e_ (Obs/Exp), where ‘Obs’ is the observed frequency of a specific alteration in the genome and ‘Exp’ is the expected frequency of that specific alteration due to individual proportion of the nucleotides in the genome sequences.

### Variation analysis at Protein level

To obtain the different protein sequences of the translated genome, open reading frame was generated using ORF Finder. Six different ORF were generated which covered the entire genome sequence. Our protein sequences were aligned with existing protein sequences of *SARS-CoV-2* present in the ‘NCBI Virus’ database to check which ORF corresponds to the which known protein. The amino acid alterations were identified from the nucleotide information and were classified into Synonymous (S) and Non Synonymous (NS). The functional impacts of all NS mutations at protein were analysed using different SNP annotation tools (i.e. SIFT, SNAP, Polyphen2 and MetaSNP).

### Fatality rate and underlying molecular predisposition

The fatality rate of a country was calculated considering the summation of total number of death and critical case and then dividing the same with total number of detected case. [Fatality rate = (Total no. of deaths + total no of Severe cases)/ Total no. of detected cases]). The fatality rates were matched against the rarity of the nucleotide alterations and functional impact of the Non-synonymous changes at protein level.

### Origin and evolutionary explanations of *SARS-CoV-2* strains

All the 13 Strains of 13 different geographical locations were categorized depending upon the no of mutations and type of mutations they had.

## Results

### MSA of whole genome sequences and identified mutations

Multiple sequence alignment of 13 whole genome sequences, representative entries from 13 countries, clearly showed extremely conserved sequence with only 35 mutations (Supplementary Material 1). Only viral genome sequence identified from Taiwan entry does not show any mutation, suggesting this is the ancestral genome sequence. All the mutations of *SARS-CoV-2* identified were reported in Table 1. Among all mutations, a three nucleotide deletion (from 21964-21966 position) was observed only from MT012098 strain of India.

**Table 1:**
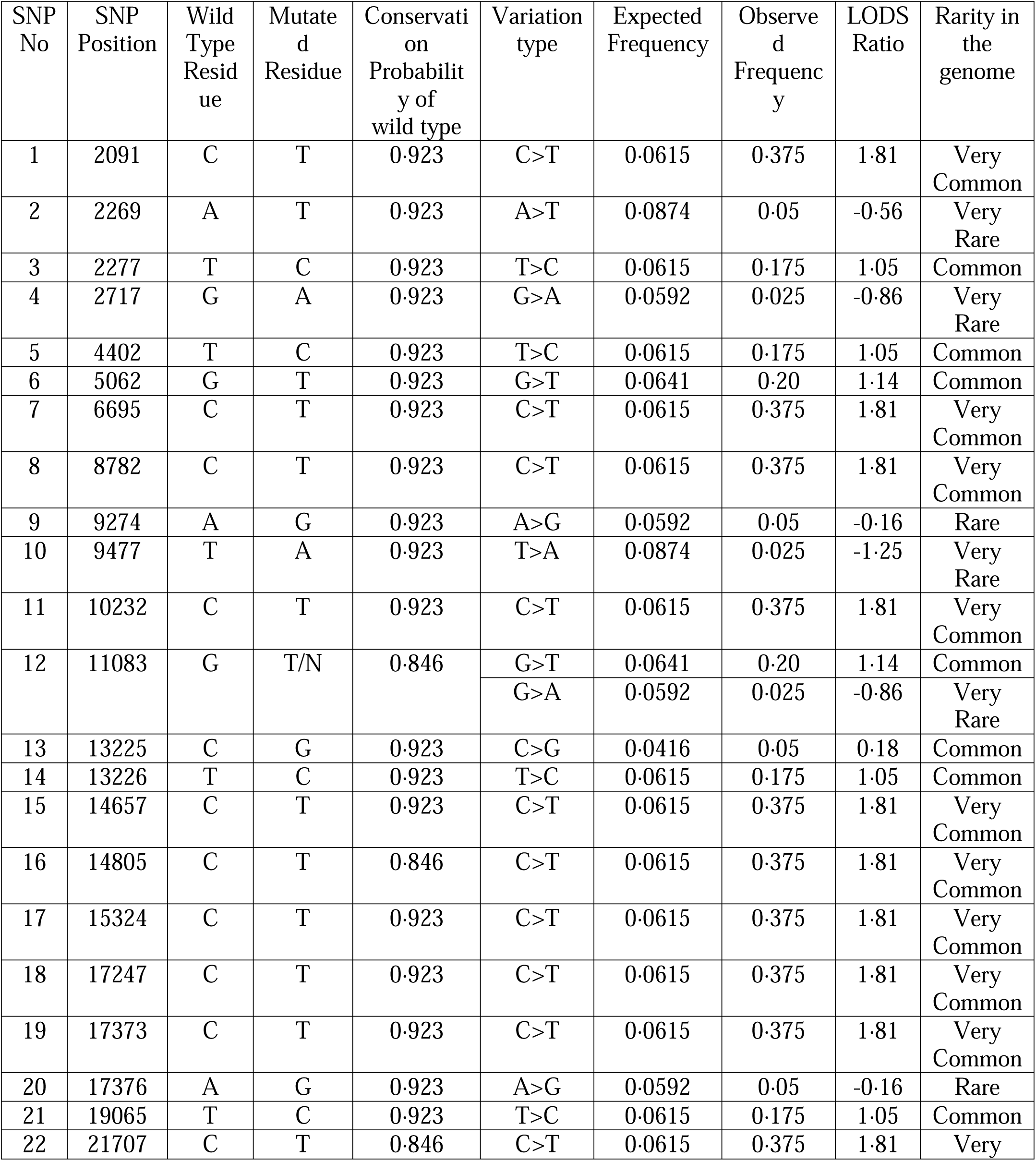
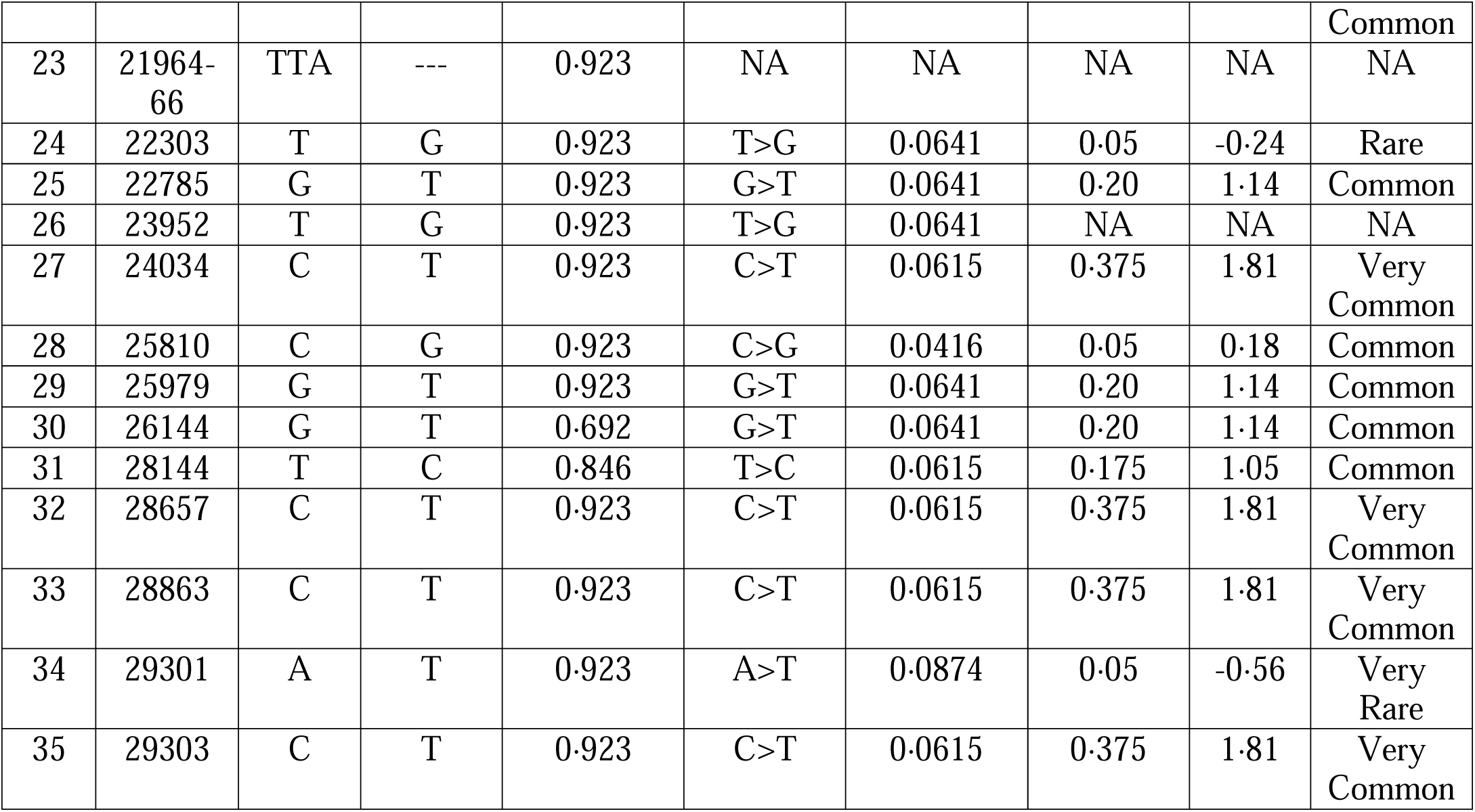
Database of Single Nucleotide Polymorphisms found by whole genome analysis of different strains of *SARS-CoV-2* retrieved from ‘NCBI virus’ database and their rarity in *SARS-CoV-2* genome

### Phylogenetic tree and evolutionary dynamics

The phylogenetic tree analysis showed the proximity of the viral strains of different COVID-19 affecting countries. Three distinct clusters have been found (figure 2) - one comprises of Sweden (MT093571), Australia (MT007544), Italy (MT066156) and Brazil (MT126808); second one comprises of Finland (MT020781), USA (MT027064) and the 3rd cluster comprises of China (MT135043) and Spain (MT233523). MT192759 strain of Taiwan is at middle position of the phylogram depicting that all the strains have diverged from it through mutations. MSA results also depicted it to be the ancestor strain of *SARS-CoV-2* with zero mutation.

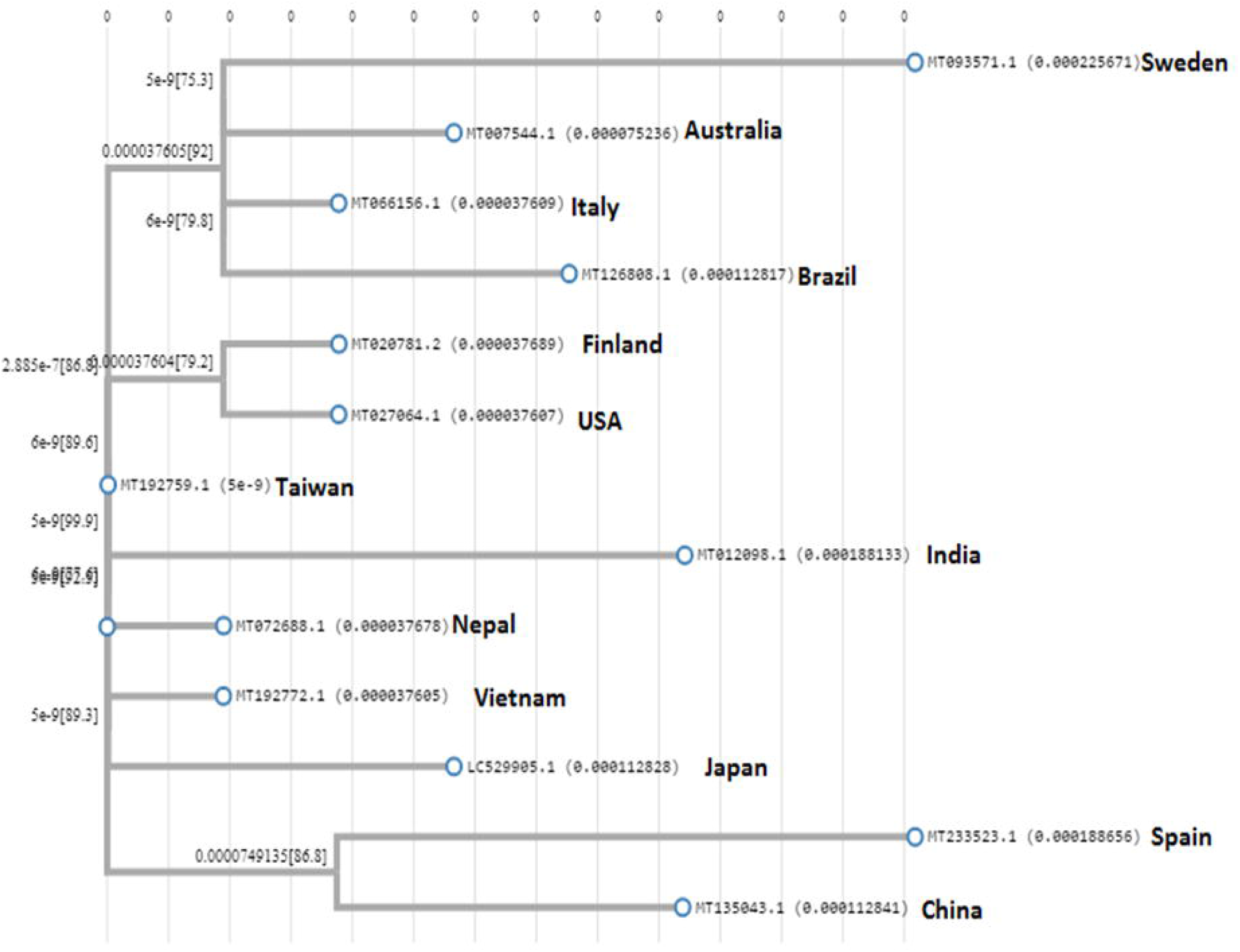

### Pair-wise Alignment between the ancestor viral strains of Wuhan with all 13 sequences

The first submitted Ancestor viral Strain of Wuhan (Submitted 5^th^ Jan, 2020; NC_045512·2) showed 100% identity with the viral Strain of Taiwan (MT192759·1) only and confirmed our previous observation (Supplementary Material 2). The comparison between *SARS-CoV* and *SARS-CoV-2* Strain showed approximately 79.2% identity; and that of *SARS-CoV* (AY323977, 2003 strain) and *SARS-CoV-2* (MT066156, 2020 strain) of Italy found 11 out of 35 mutations to be common, suggesting viral strain of Wuhan and Taiwan are identical and *SARS-CoV* shares common features with different strains of *SARS-CoV-2* (Supplementary Material 3).

### Screening of Mutations

All the mutations are screened depending upon their rarity to occur in this viral genome and A>T, G>A and T>A are found to be very rare alterations with LOD score value less than −0·5 where as C>T is the most common alteration with LOD score 1·81.

### Variation analysis at Protein level

Different protein sequences are obtained using ORF finder and those sequences were matched with existing Orf1ab polyprotein, spike like surface glycoprotein, nucleocapsid phosphoprotein etc. to ensure prediction of *in silico* ORF with corresponding *SARS-CoV-2* protein. Next, observed mutations were confirmed at amino acid level and identified a total of 20 Non-synonymous (NS) and 14 Synonymous (S) mutations (Table 2). The analysis with different SNP annotation tools like SIFT,^16^ SNAP,^17^ Polyphen2,^18^ MetaSNP^19^ showed the functional impact of NS mutations at different level, predicting either tolerance of the mutation, or disease causing ability, or executing risk or damage, which altogether can infer the fatality rate.

**Table 2:** Effect of mutation in protein level and Fatality rate in different geographical location (Shown in ascending order of achieving mutation by the strain)

| Viral Strain (Accession No) | Country Name | Total no of Mutation | Mutation position | Variation type | Codon Alteration | Effect on Protein Sequence | Mutation Catagory | Fatality Rate |
| --- | --- | --- | --- | --- | --- | --- | --- | --- |
| MT192759 | Taiwan | 0 | NA | NA | NA | NA |  | 0.80 |
| MT072688 | Nepal | 1 | 24034 | C>T | AAC>AAT | N833N | S | 0.00 |
| MT020781 | Finland | 1 | 21707 | C>T | CAT>TAT | H58Y | NS | 2.63 |
| MT192772 | Vietnam | 1 | 10232 | C>T | CGT>TGT | R3323C | NS | 2.01 |
| MT027064 | USA | 2 | 2091<br>21707 | C>T<br>C>T | ACT>ATT<br>CAT>TAT | T609I<br>H58Y | NS<br>NS | 3.80 |
| MT007544 | Australia | 3 | 19065<br>22303<br>26144 | T>C<br>T>G<br>G>T | CCT>CCC<br>AGT>AGG<br>GGT>GTT | P1766P<br>S256R<br>G251V | S<br>NS<br>NS | 0.91 |
| LC529905 | Japan | 3 | 15324<br>25810<br>29303 | C>T<br>C>G<br>C>T | AAC>AAT<br>CTT>GTT<br>CCA>TCA | N519N<br>L140V<br>P344S | S<br>NS<br>NS | 7.57 |
| MT066156 | Italy | 3 | 2269<br>11083<br>26144 | A>T<br>G>N<br>G>T | GCA>GCT<br>TTG>TTT/<br>A<br>GGT>GTT | A668A<br>L3606F<br>G251V | S<br>NS<br>NS | 14.82 |
| MT126808 | Brazil | 4 | 11083<br>14805<br>17247<br>26144 | G>T<br>C>T<br>T>C<br>G>T | TTG>TTT<br>TAC>TAT<br>CGT>CGC<br>GGT>GTT | L3606F<br>Y346Y<br>R1160R<br>G251V | NS<br>S<br>S<br>NS | 7.33 |
| MT135043 | China | 5 | 4402<br>5062<br>8782<br>28144<br>29301 | T>C<br>G>T<br>C>T<br>T>C<br>A>T | CTT>CTC<br>TTG>TTT<br>AGC>AGT<br>TTT>TTC<br>GAT>GTT | L1379L<br>L1599F<br>S2839S<br>F87F<br>D343V | S<br>NS<br>S<br>S<br>NS | 5.45 |
| MT012098 | India | 6<br>(includes a deletion) | 2277<br>6695<br>14657<br>17373<br>22785<br>21964-66 | T>C<br>C>T<br>C>T<br>C>T<br>G>T<br>TTA Deletion | ATT>ACT<br>CCT>TCT<br>GCT>GTT<br>GCC>GCT<br>AGA>ATA<br>TATTAC>TAC | I671T<br>P2144S<br>A306V<br>A1202A<br>R417I<br>Y154 Deletion | NS<br>NS<br>NS<br>S<br>NS<br>Deletion | 2.06 |
| MT093571 | Sweden | 7 | 2717<br>9274<br>13225<br>13226<br>17376<br>23952<br>26144 | G>A<br>A>G<br>C>G<br>T>C<br>A>G<br>T>G<br>G>T | GGT>AGT<br>AGA>AGG<br>TCC>TCG<br>TTT>CTT<br>ACA>ACG<br>TTT>TGT<br>GGT>GTT | G818S<br>R3003R<br>S4320S<br>F4321L<br>T1203T<br>F806C<br>G251V | NS<br>S<br>S<br>NS<br>S<br>NS<br>NS | 9.20 |
| MT233523 | Spain | 7 | 8782 | C>T | AGC<u>→</u>AGT | S2839S | S | 13.77 |
|  |  |  | 9477 | T>A | TTT>TAT | F3071Y | NS |  |
|  |  |  | 14805 | C>T | TAC<u>→</u>TAT | Y346Y | S |  |
|  |  |  | 25979 | G>T | GGA>GTA | G196V | NS |  |
|  |  |  | 28144 | T>C | TTT>TTC | F87F | S |  |
|  |  |  | 28657 | C>T | GAC<u>→</u>GAT | D128D | S |  |
|  |  |  | 28863 | C>T | TCA>TTA | S197L | NS |  |

S = Synonymous Mutation, NS = Non Synonymous Mutation

### Fatality Rate Calculation and underlying molecular predisposition

The fatality rate of Italy (14·82%), Spain (13·77%) and Sweden (9·20%) were significantly higher, where it was less than 4% in countries like Nepal (Zero fatality), Finland (2·63%), Vietnam (2·01%), USA (3·80), Australia (0·91%) and India (2·06). In case of Japan, Brazil and China, fatality rate was moderate i.e approximately 4-8% (figure 3). The number of mutations and presence of either of the very rare mutations or functionally important NS mutations or both are found to be strongly linked with the fatality rate (Table 3A-Table 3C).

**Table 3A:**
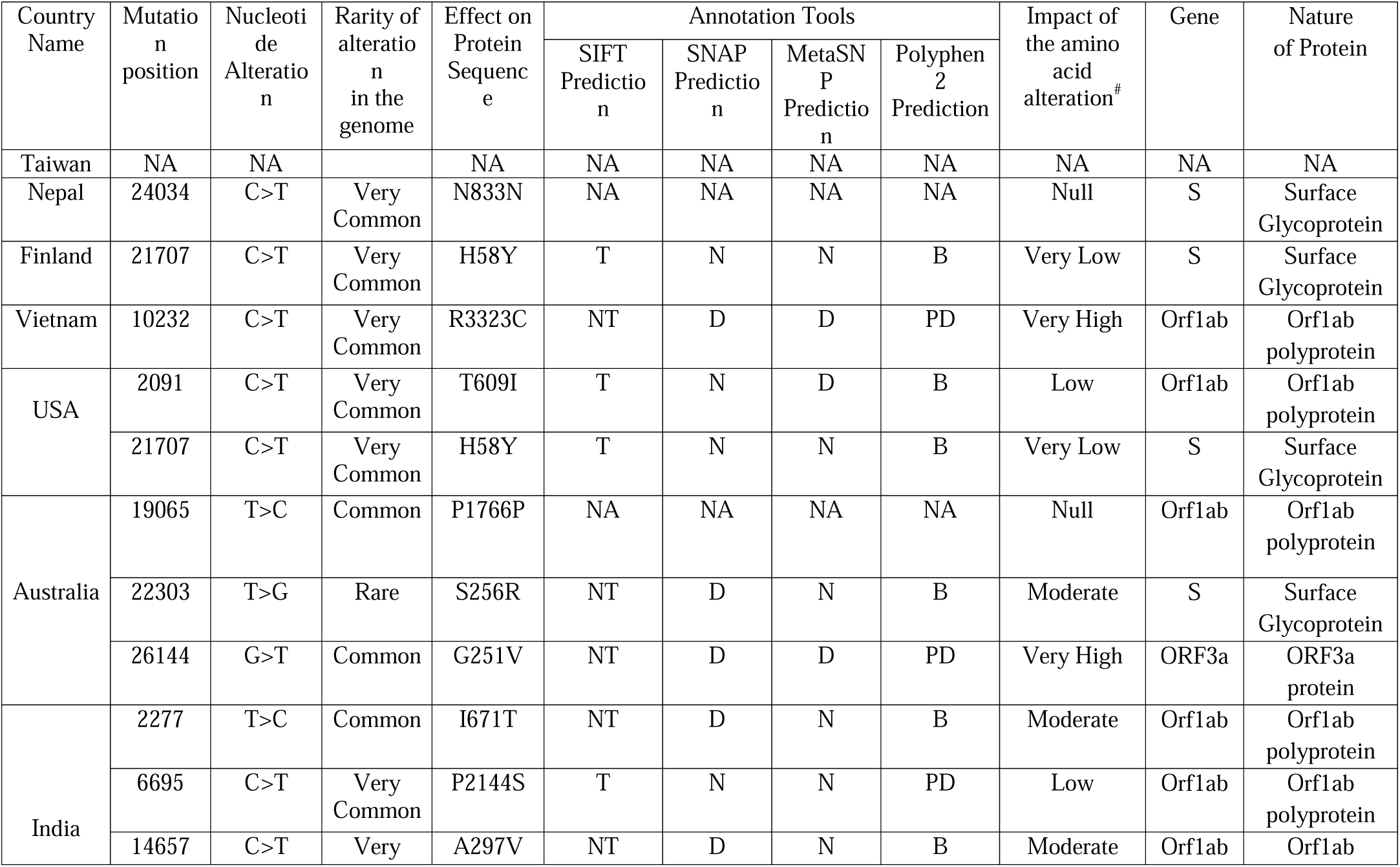
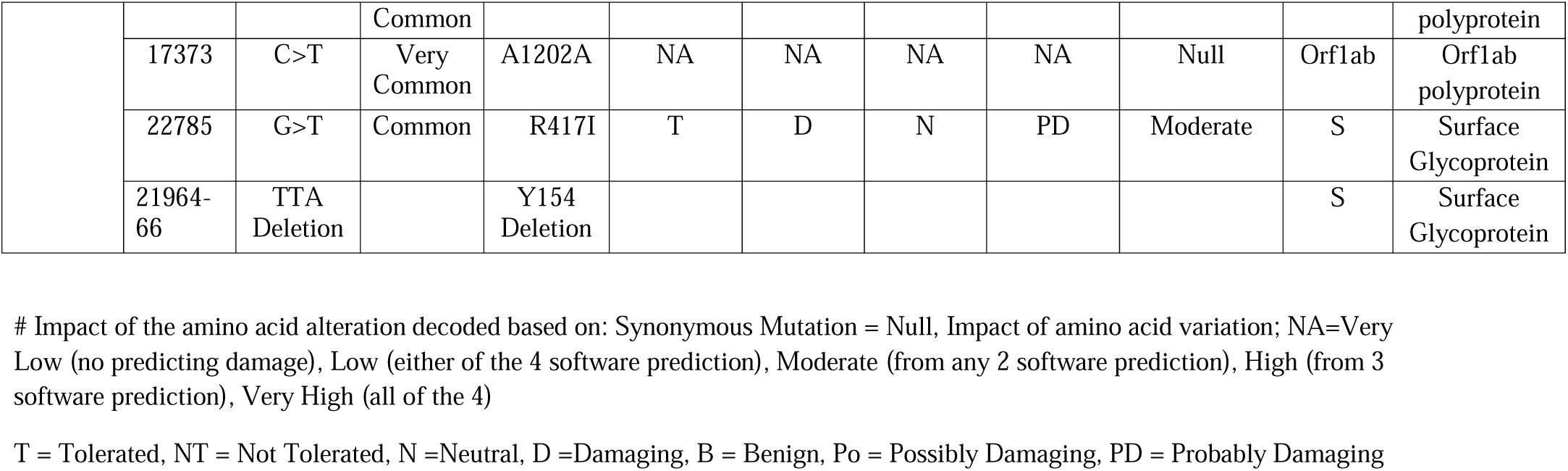
Countries with Fatality Rate <4% and underlying molecular predisposition

**Table 3B:**
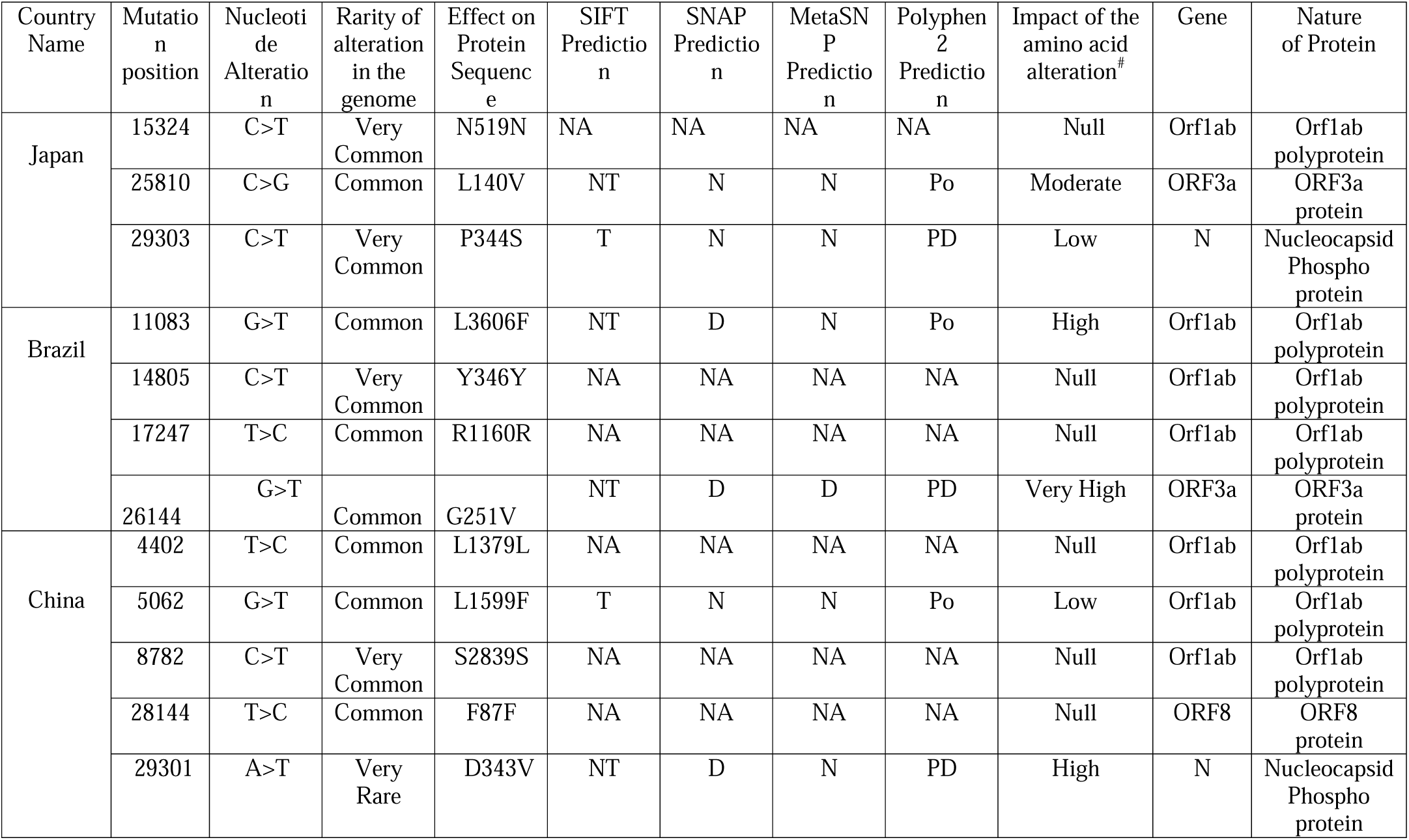
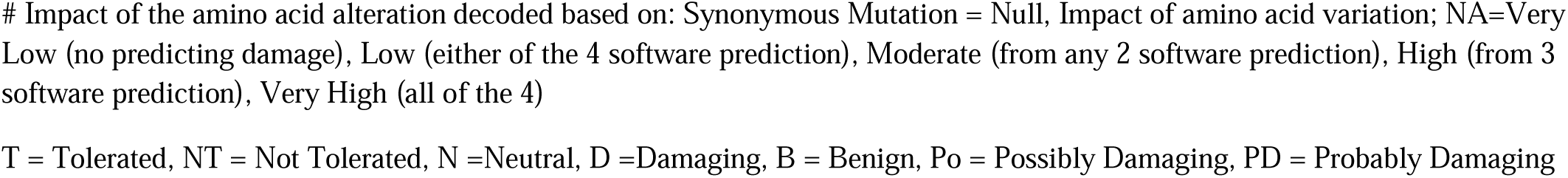
Countries with Fatality Rate 4-8% and underlying molecular predisposition.

**Table 3C:**
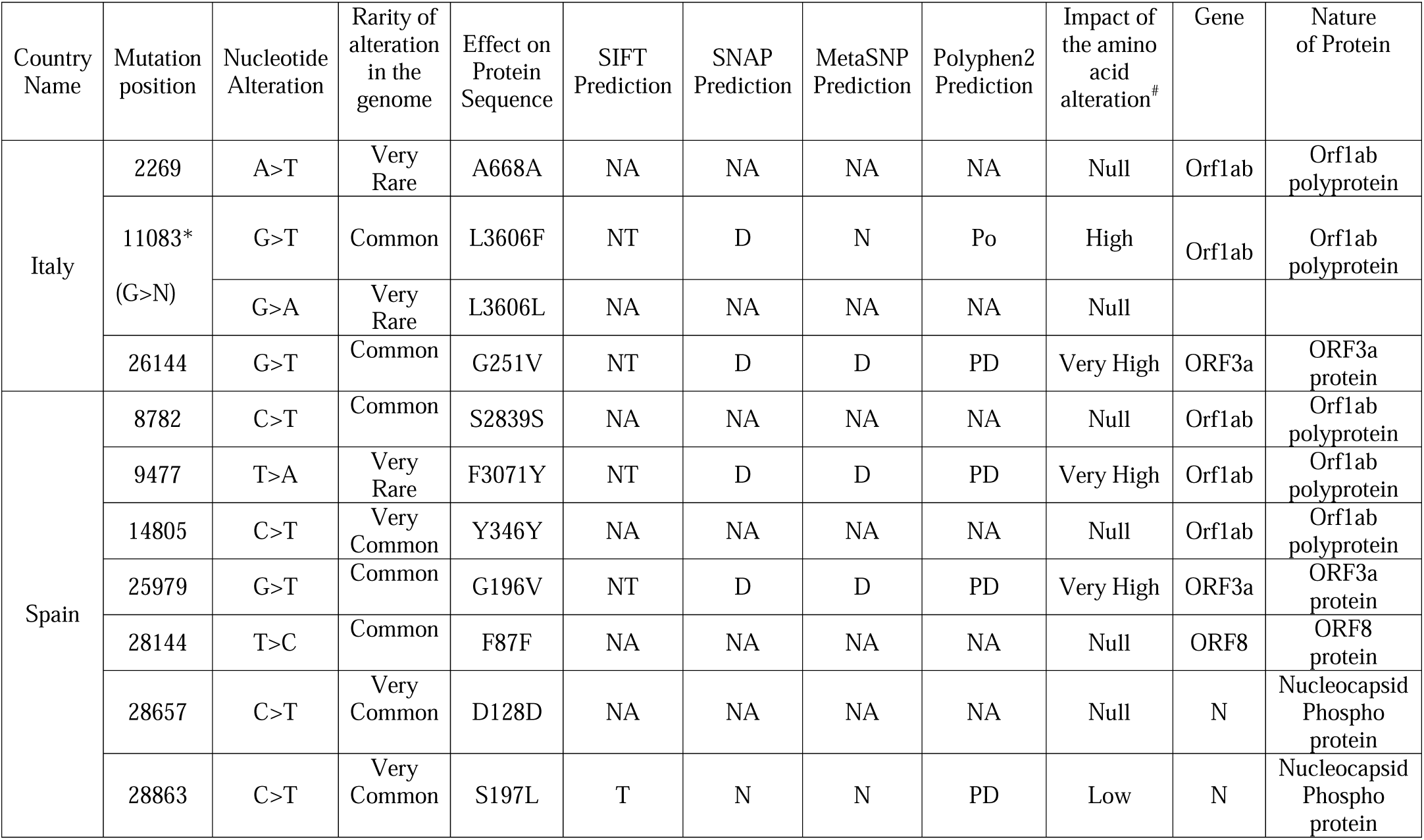
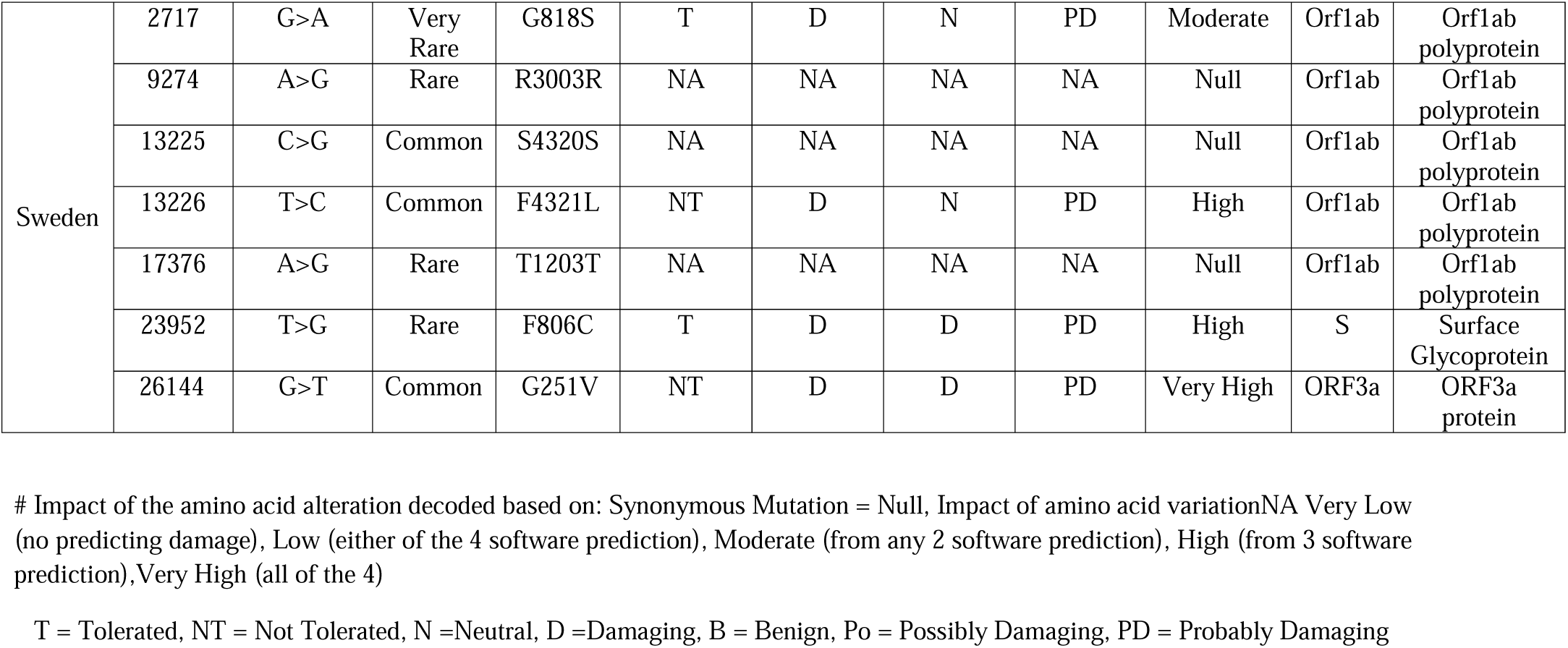
Countries with Fatality Rate >8 % and underlying molecular predisposition.

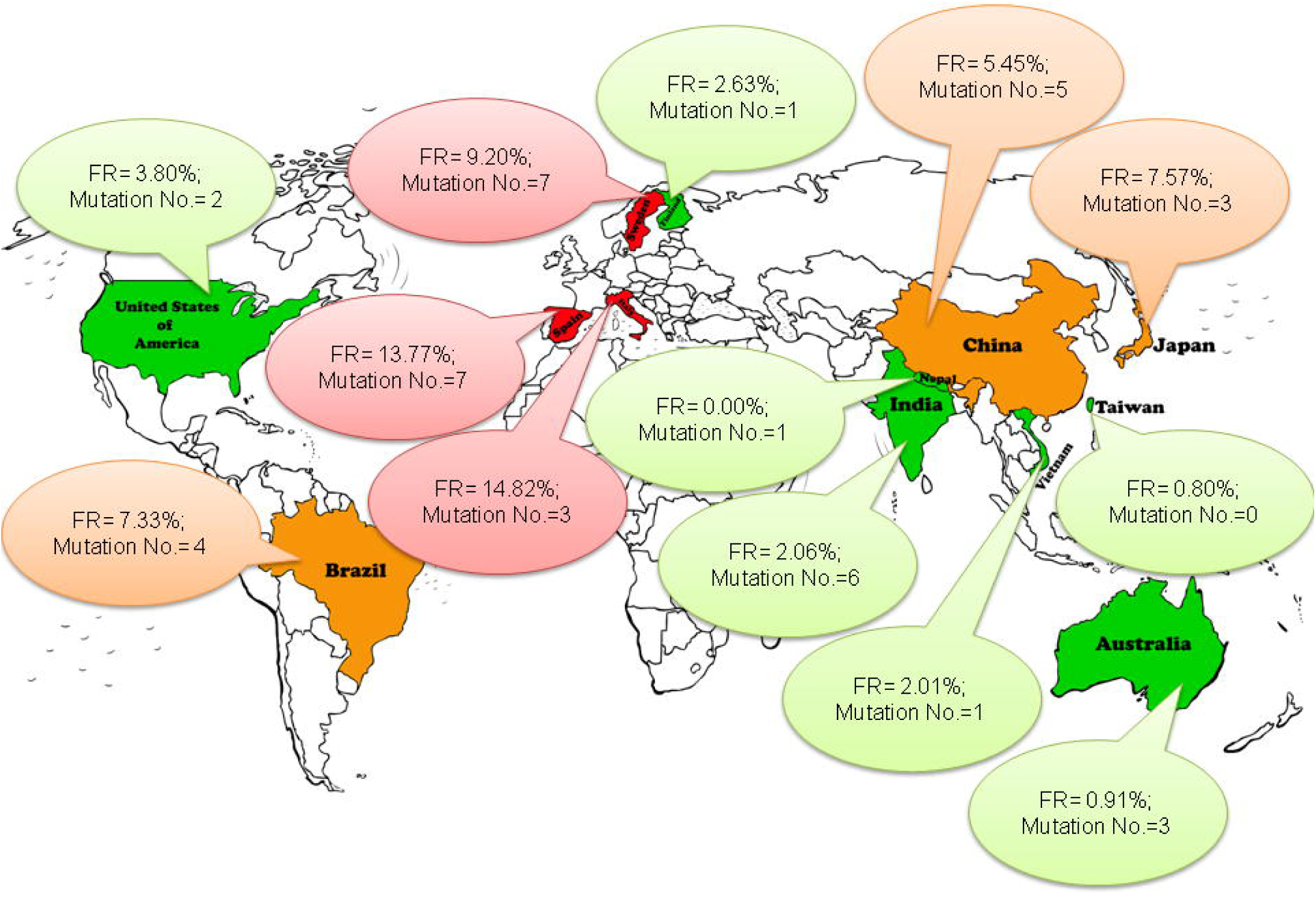

### Origin and evolution of *SARS-CoV-2* strains

It is extremely important to understand how the ancestral strain (S) leads to lethal strain (L) and other clusters observed according to phylogeny. Our analysis showed Ancestral strain (S) give rise to an Intermediate strain (I) with single very common mutation, a transition from C to T. From I, three different strains originate, i.e (a) Lethal strain (L), with additional mutations over I, (b) Extremely lethal strain (E) that contains very rare mutations and a (c) Non Lethal strain (N) that contains favourable mutations at surface glycoprotein, which possibly inhibit the interaction of ACE2 and favouring non-lethal outcome (figure 4).

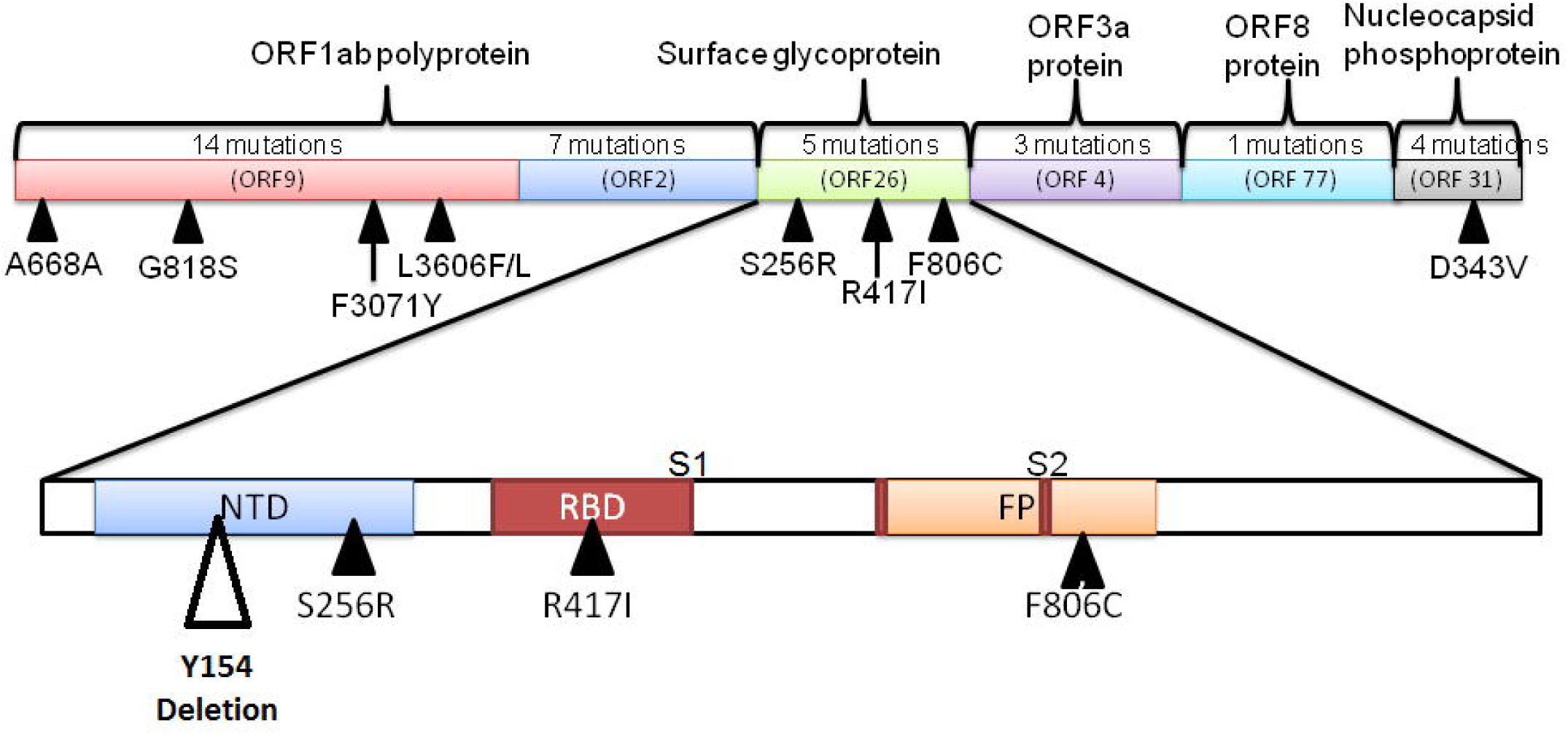

## Discussion

The current global pandemic of the novel coronavirus Covid-19 created two epicenters first at Hubei Province in People’s Republic of China and presently in Europe. *SARS-CoV-2*, like its close relatives *SARS-CoV* and *MERS-CoV*, is also pathogenic but with a higher infectivity rate. The increasing number of cases and wide spread disease raise grave concerns about the future trajectory of the pandemic and thus, a better understanding regarding the molecular divergence of viral strains and pathogenesis, is utmost important.

The present study attempted to categorize COVID affected countries based on molecular pathogenesis. Three important factors were considered, i.e, number of mutations during evolution, rarity of the allelic substitution and functional alteration of the non-synonymous mutations. We screened and compared extent of mutations observed in genome sequence of *SARS-CoV-2*. All reported genome sequences of 13 affected countries have been analyzed. So far, studies indicated that there are only two strains, S, possibly the ancestral one and L, might be the lethal and more aggressive one. The distinct clusters that we observed from our phylogenetic tree construction raise the assumption of existence of many more unknown strains (figure 5). One interesting report substantiating our finding is from Shenzhen which showed possible generation of new strain which neither belongs to ‘S’ nor to ‘L’ subtype.^12^ We evaluated the extent of molecular divergence by MSA and Pair-wise alignment and subsequently tried to decipher the protein alteration parallel to the mutations.

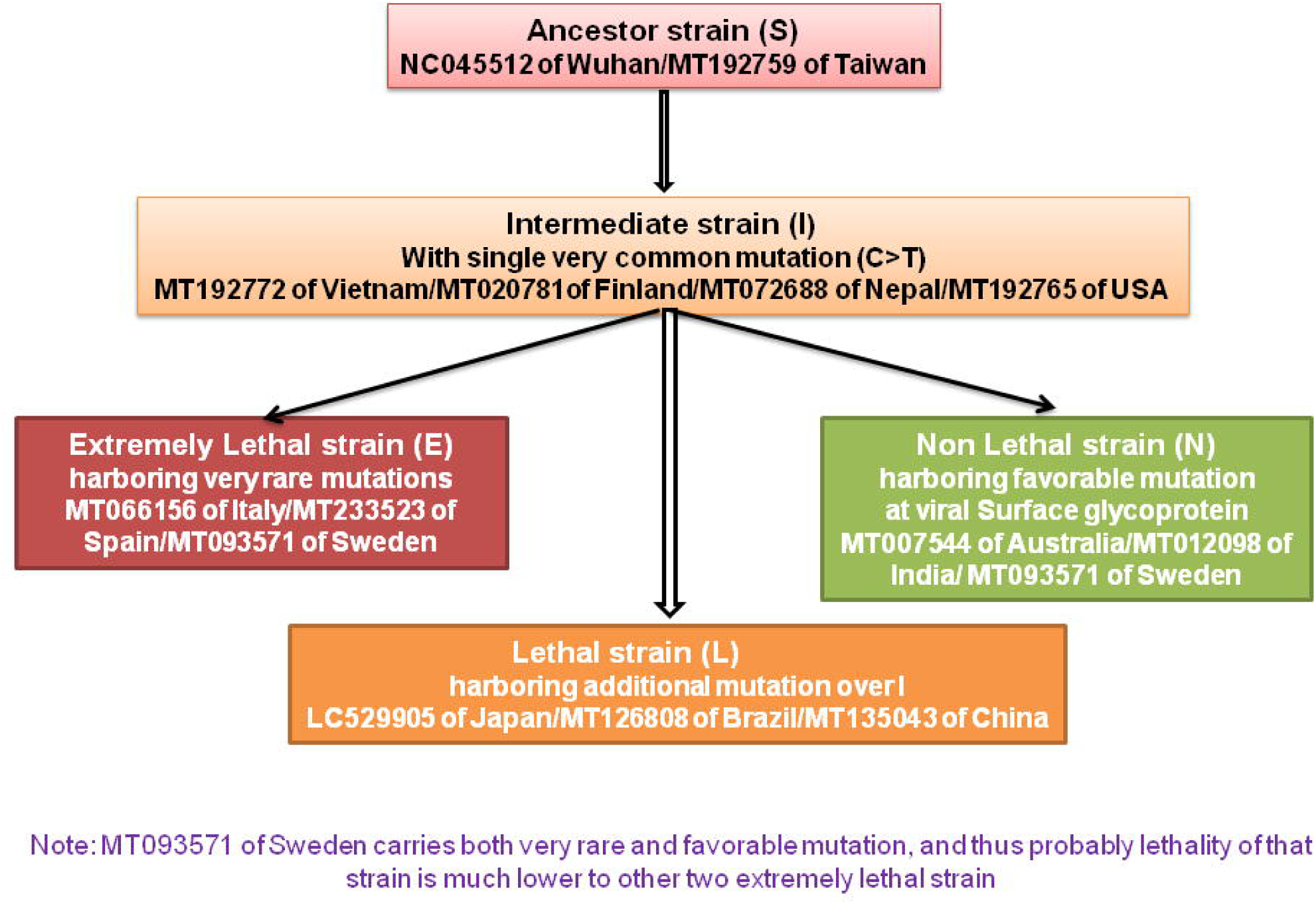

As a RNA virus, it was hypothesized that 2019-nCoV mutate faster than DNA virus.^12^ The genome-wide phylogenetic tree indicated that 2019-nCoV was closest to SARS-like coronavirus, with 96% sequence similarity.^20,21^ When we evaluated the *SARS-CoV* (2003) and *SARS-CoV-2* (2020) from Italy, sequence identity was 79.4%, suggesting new mutations, might be deleterious in later strain.

Our first approach was to categorize the countries depending on number of mutations and type of mutation they had. It was found that similar number of mutations does not correlate well with similar extent of fatality outcome. Example, patients from Australia, Japan and Italy all show 3 mutations in the viral genome, but Australia has least fatality rate, while Italy has maximum. Again, numbers of mutations are 5 in China, 6 in India and 7 in both Sweden and Spain. The observations mentioned above strongly indicate that not the total number of mutation, but the nature of mutations finally guides the overall fatality outcome. In the case of Italy, we found three deadly mutations in 2020 (i.e. 2269 A>T, very rare allele; 11083 G>A, very rare allele and 26144 G>T) with high disease outcome but the wild counterpart of these found in 2003 did not lead to any fatality, suggesting the significance of newly evolved mutations. Although, 2269 A>T mutation does not alter amino acid (A668A), increasing number of evidences suggest that synonymous mutations could have effects on splicing, transcription, that ultimately alter the phenotype, disrupting their silence.^22^ Spain is also carrying a very rare allele transversion T>A, that occurs in orf1ab gene of virus. Orf1ab gene transcribes into a polyprotein and cleaving by protease (3CLpro) and papain-like protease (PLpro) produces several non-structural proteins, which are important for replication as well as virulence for coronavirus. Thus, an alteration in this region might alter the virulence, and associated fatality outcome.

We assume, while some mutations are pathogenic, some will be favorable and will undergo positive selection pressure. Herein, we tried to elucidate the possible interaction between ACE2 and spike glycoprotein. It is well established that for virus entry, spike glycoprotein (S) present on the CoV can be a neutralization antibody and it binds to its receptor followed by membrane fusion. It has been inferred that some favorable changes in viral glycoprotein may limit the increase of fatality. This kind of favorable mutation was found in some strains of COVID-19 from India, Australia and Sweden. Detailed screening depicted that Australian strain carry a rare allele transversion (T>G) which results into a NS mutation (S256R) on surface glycoprotein S1 domain which may affect the binding of hACE2 molecules, thus these strains become non-lethal to human. In this study, only one tri-nucleotide deletion at S1 domain of surface glycoprotein has been found and that is found in Indian strains along with a NS mutation (R417I) in the Receptor Binding Domain (RBD) of S1 subunit that disfavor viral entry by inhibiting hACE2 interaction.

On the other hand, Brazilian strain acquires four common allelic transitions, of which only two non-synonymous mutations affect protein alteration. Surprisingly, Swedish strain carry both extreme lethal (2717 G>A, very rare allele leading to G818S at Orf1ab polyprotein) and also favorable mutations (23952 T>G, rare allele leading to F806C at surface glycoprotein) and thus cumulatively lower the severity of the disease.

In summary, the present study reveals that the fatality rate increases with not only the number of mutations but also depending on its allelic rarity as well as functional alteration of protein. Surface glycoprotein domain is very important for host-carrier interactions and hence the mutations affecting surface glycoprotein can be one of the important mechanisms which alter the viral entry and pathogenesis. Future studies may uncover more genetic information at the molecular level as well as structural levels of the proteins, because without this knowledge it’ll be difficult to identify drug target and prepare vaccine. We hope our work will help in that direction.

## Supporting information

Location of the mutation and the corresponding genetic variation

Pairwise Sequence Alignment between the first submitted Ancestor viral Strain of Wuhan(NC_045512.2) and the viral Strain of Taiwan

Alignment file between a SARS-CoV strain and SARS-CoV-2 strain

## Author Contribution

SB, PB designed the study; SB, SD analyzed the data; SB, PB interpreted the data; SB, SD, SBh searched literatures; SD, SBh, PB wrote the manuscript; SD prepared figures; PB supervised overall study.

## Acknowledgement

The authors acknowledge UGC-DAE CSR for providing fellowship to Shuvam Banerjee.

