## Supplementary material for "Decoding the lethal effect of *SARS-CoV-2* (novel coronavirus) strains from global perspective: molecular pathogenesis and evolutionary divergence": Location of the mutation and the corresponding genetic variation

**Supplementary Material: 1**

**Location of the mutation and the corresponding genetic variation**

```
                                                              2269    2277
MT233523.1      CTCAACCTGTGCTTGTGAAATTGTCGGTGGACAAATTGTCACCTGTGCAAAGGAAATTAA  Spain
MT135043.1      CTCAACCTGTGCTTGTGAAATTGTCGGTGGACAAATTGTCACCTGTGCAAAGGAAATTAA  China
MT012098.1      CTCAACCTGTGCTTGTGAAATTGTCGGTGGACAAATTGTCACCTGTGCAAAGGAAACTAA  India
MT072688.1      CTCAACCTGTGCTTGTGAAATTGTCGGTGGACAAATTGTCACCTGTGCAAAGGAAATTAA  Nepal
MT027064.1      CTCAACCTGTGCTTGTGAAATTGTCGGTGGACAAATTGTCACCTGTGCAAAGGAAATTAA  USA
MT020781.2      CTCAACCTGTGCTTGTGAAATTGTCGGTGGACAAATTGTCACCTGTGCAAAGGAAATTAA  Finland
LC529905.1      CTCAACCTGTGCTTGTGAAATTGTCGGTGGACAAATTGTCACCTGTGCAAAGGAAATTAA  Japan
MT066156.1      CTCAACCTGTGCTTGTGAAATTGTCGGTGGACAAATTGTCACCTGTGCTAAGGAAATTAA  Italy
MT192772.1      CTCAACCTGTGCTTGTGAAATTGTCGGTGGACAAATTGTCACCTGTGCAAAGGAAATTAA  VietNam
MT192759.1      CTCAACCTGTGCTTGTGAAATTGTCGGTGGACAAATTGTCACCTGTGCAAAGGAAATTAA  Taiwan
MT126808.1      CTCAACCTGTGCTTGTGAAATTGTCGGTGGACAAATTGTCACCTGTGCAAAGGAAATTAA  Brazil
MT093571.1      CTCAACCTGTGCTTGTGAAATTGTCGGTGGACAAATTGTCACCTGTGCAAAGGAAATTAA  Sweden
MT007544.1      CTCAACCTGTGCTTGTGAAATTGTCGGTGGACAAATTGTCACCTGTGCAAAGGAAATTAA  Australia
                ************************************************ ******* ***

                               2717
MT233523.1      CTTCACACTCAAAGGCGGTGCACCAACAAAGGTTACTTTTGGTGATGACACTGTGATAGA
MT135043.1      CTTCACACTCAAAGGCGGTGCACCAACAAAGGTTACTTTTGGTGATGACACTGTGATAGA
MT012098.1      CTTCACACTCAAAGGCGGTGCACCAACAAAGGTTACTTTTGGTGATGACACTGTGATAGA
MT072688.1      CTTCACACTCAAAGGCGGTGCACCAACAAAGGTTACTTTTGGTGATGACACTGTGATAGA
MT027064.1      CTTCACACTCAAAGGCGGTGCACCAACAAAGGTTACTTTTGGTGATGACACTGTGATAGA
MT020781.2      CTTCACACTCAAAGGCGGTGCACCAACAAAGGTTACTTTTGGTGATGACACTGTGATAGA
LC529905.1      CTTCACACTCAAAGGCGGTGCACCAACAAAGGTTACTTTTGGTGATGACACTGTGATAGA
MT066156.1      CTTCACACTCAAAGGCGGTGCACCAACAAAGGTTACTTTTGGTGATGACACTGTGATAGA
MT192772.1      CTTCACACTCAAAGGCGGTGCACCAACAAAGGTTACTTTTGGTGATGACACTGTGATAGA
MT192759.1      CTTCACACTCAAAGGCGGTGCACCAACAAAGGTTACTTTTGGTGATGACACTGTGATAGA
MT126808.1      CTTCACACTCAAAGGCGGTGCACCAACAAAGGTTACTTTTGGTGATGACACTGTGATAGA
MT093571.1      CTTCACACTCAAAGGCAGTGCACCAACAAAGGTTACTTTTGGTGATGACACTGTGATAGA
MT007544.1      CTTCACACTCAAAGGCGGTGCACCAACAAAGGTTACTTTTGGTGATGACACTGTGATAGA
                **************** *******************************************


                                    4402
MT233523.1      TTGGAATTTGCGAGAAATGCTTGCACATGCAGAAGAAACACGCAAATTAATGCCTGTCTG
MT135043.1      TTGGAATTTGCGAGAAATGCTCGCACATGCAGAAGAAACACGCAAATTAATGCCTGTCTG
MT012098.1      TTGGAATTTGCGAGAAATGCTTGCACATGCAGAAGAAACACGCAAATTAATGCCTGTCTG
MT072688.1      TTGGAATTTGCGAGAAATGCTTGCACATGCAGAAGAAACACGCAAATTAATGCCTGTCTG
MT027064.1      TTGGAATTTGCGAGAAATGCTTGCACATGCAGAAGAAACACGCAAATTAATGCCTGTCTG
MT020781.2      TTGGAATTTGCGAGAAATGCTTGCACATGCAGAAGAAACACGCAAATTAATGCCTGTCTG
LC529905.1      TTGGAATTTGCGAGAAATGCTTGCACATGCAGAAGAAACACGCAAATTAATGCCTGTCTG
MT066156.1      TTGGAATTTGCGAGAAATGCTTGCACATGCAGAAGAAACACGCAAATTAATGCCTGTCTG
MT192772.1      TTGGAATTTGCGAGAAATGCTTGCACATGCAGAAGAAACACGCAAATTAATGCCTGTCTG
MT192759.1      TTGGAATTTGCGAGAAATGCTTGCACATGCAGAAGAAACACGCAAATTAATGCCTGTCTG
MT126808.1      TTGGAATTTGCGAGAAATGCTTGCACATGCAGAAGAAACACGCAAATTAATGCCTGTCTG
MT093571.1      TTGGAATTTGCGAGAAATGCTTGCACATGCAGAAGAAACACGCAAATTAATGCCTGTCTG
MT007544.1      TTGGAATTTGCGAGAAATGCTTGCACATGCAGAAGAAACACGCAAATTAATGCCTGTCTG
                ********************* **************************************

                                    5062
MT233523.1      ACAGTTTGGTCCAACTTATTTGGATGGAGCTGATGTTACTAAAATAAAACCTCATAATTC
MT135043.1      ACAGTTTGGTCCAACTTATTTTGATGGAGCTGATGTTACTAAAATAAAACCTCATAATTC
MT012098.1      ACAGTTTGGTCCAACTTATTTGGATGGAGCTGATGTTACTAAAATAAAACCTCATAATTC
MT072688.1      ACAGTTTGGTCCAACTTATTTGGATGGAGCTGATGTTACTAAAATAAAACCTCATAATTC
MT027064.1      ACAGTTTGGTCCAACTTATTTGGATGGAGCTGATGTTACTAAAATAAAACCTCATAATTC
MT020781.2      ACAGTTTGGTCCAACTTATTTGGATGGAGCTGATGTTACTAAAATAAAACCTCATAATTC
LC529905.1      ACAGTTTGGTCCAACTTATTTGGATGGAGCTGATGTTACTAAAATAAAACCTCATAATTC
MT066156.1      ACAGTTTGGTCCAACTTATTTGGATGGAGCTGATGTTACTAAAATAAAACCTCATAATTC
MT192772.1      ACAGTTTGGTCCAACTTATTTGGATGGAGCTGATGTTACTAAAATAAAACCTCATAATTC
MT192759.1      ACAGTTTGGTCCAACTTATTTGGATGGAGCTGATGTTACTAAAATAAAACCTCATAATTC
MT126808.1      ACAGTTTGGTCCAACTTATTTGGATGGAGCTGATGTTACTAAAATAAAACCTCATAATTC
MT093571.1      ACAGTTTGGTCCAACTTATTTGGATGGAGCTGATGTTACTAAAATAAAACCTCATAATTC
MT007544.1      ACAGTTTGGTCCAACTTATTTGGATGGAGCTGATGTTACTAAAATAAAACCTCATAATTC
                ********************* **************************************


                                                6695
MT233523.1      TGTCCCTTGGGATACTATAGCTAATTATGCTAAGCCTTTTCTTAACAAAGTTGTTAGTAC
MT135043.1      TGTCCCTTGGGATACTATAGCTAATTATGCTAAGCCTTTTCTTAACAAAGTTGTTAGTAC
MT012098.1      TGTCCCTTGGGATACTATAGCTAATTATGCTAAGTCTTTTCTTAACAAAGTTGTTAGTAC
MT072688.1      TGTCCCTTGGGATACTATAGCTAATTATGCTAAGCCTTTTCTTAACAAAGTTGTTAGTAC
MT027064.1      TGTCCCTTGGGATACTATAGCTAATTATGCTAAGCCTTTTCTTAACAAAGTTGTTAGTAC
MT020781.2      TGTCCCTTGGGATACTATAGCTAATTATGCTAAGCCTTTTCTTAACAAAGTTGTTAGTAC
LC529905.1      TGTCCCTTGGGATACTATAGCTAATTATGCTAAGCCTTTTCTTAACAAAGTTGTTAGTAC
MT066156.1      TGTCCCTTGGGATACTATAGCTAATTATGCTAAGCCTTTTCTTAACAAAGTTGTTAGTAC
MT192772.1      TGTCCCTTGGGATACTATAGCTAATTATGCTAAGCCTTTTCTTAACAAAGTTGTTAGTAC
MT192759.1      TGTCCCTTGGGATACTATAGCTAATTATGCTAAGCCTTTTCTTAACAAAGTTGTTAGTAC
MT126808.1      TGTCCCTTGGGATACTATAGCTAATTATGCTAAGCCTTTTCTTAACAAAGTTGTTAGTAC
MT093571.1      TGTCCCTTGGGATACTATAGCTAATTATGCTAAGCCTTTTCTTAACAAAGTTGTTAGTAC
MT007544.1      TGTCCCTTGGGATACTATAGCTAATTATGCTAAGCCTTTTCTTAACAAAGTTGTTAGTAC
                ********************************** *************************

                                   8782
MT233523.1      TGATTTTGACACATGGTTTAGTCAGCGTGGTGGTAGTTATACTAATGACAAAGCTTGCCC
MT135043.1      TGATTTTGACACATGGTTTAGTCAGCGTGGTGGTAGTTATACTAATGACAAAGCTTGCCC
MT012098.1      TGATTTTGACACATGGTTTAGCCAGCGTGGTGGTAGTTATACTAATGACAAAGCTTGCCC
MT072688.1      TGATTTTGACACATGGTTTAGCCAGCGTGGTGGTAGTTATACTAATGACAAAGCTTGCCC
MT027064.1      TGATTTTGACACATGGTTTAGCCAGCGTGGTGGTAGTTATACTAATGACAAAGCTTGCCC
MT020781.2      TGATTTTGACACATGGTTTAGCCAGCGTGGTGGTAGTTATACTAATGACAAAGCTTGCCC
LC529905.1      TGATTTTGACACATGGTTTAGCCAGCGTGGTGGTAGTTATACTAATGACAAAGCTTGCCC
MT066156.1      TGATTTTGACACATGGTTTAGCCAGCGTGGTGGTAGTTATACTAATGACAAAGCTTGCCC
MT192772.1      TGATTTTGACACATGGTTTAGCCAGCGTGGTGGTAGTTATACTAATGACAAAGCTTGCCC
MT192759.1      TGATTTTGACACATGGTTTAGCCAGCGTGGTGGTAGTTATACTAATGACAAAGCTTGCCC
MT126808.1      TGATTTTGACACATGGTTTAGCCAGCGTGGTGGTAGTTATACTAATGACAAAGCTTGCCC
MT093571.1      TGATTTTGACACATGGTTTAGCCAGCGTGGTGGTAGTTATACTAATGACAAAGCTTGCCC
MT007544.1      TGATTTTGACACATGGTTTAGCCAGCGTGGTGGTAGTTATACTAATGACAAAGCTTGCCC
                ********************* **************************************

                                               9274
MT233523.1      AGAAGCTGGTGTTTGTGTATCTACTAGTGGTAGATGGGTACTTAACAATGATTATTACAG
MT135043.1      AGAAGCTGGTGTTTGTGTATCTACTAGTGGTAGATGGGTACTTAACAATGATTATTACAG
MT012098.1      AGAAGCTGGTGTTTGTGTATCTACTAGTGGTAGATGGGTACTTAACAATGATTATTACAG
MT072688.1      AGAAGCTGGTGTTTGTGTATCTACTAGTGGTAGATGGGTACTTAACAATGATTATTACAG
MT027064.1      AGAAGCTGGTGTTTGTGTATCTACTAGTGGTAGATGGGTACTTAACAATGATTATTACAG
MT020781.2      AGAAGCTGGTGTTTGTGTATCTACTAGTGGTAGATGGGTACTTAACAATGATTATTACAG
LC529905.1      AGAAGCTGGTGTTTGTGTATCTACTAGTGGTAGATGGGTACTTAACAATGATTATTACAG
MT066156.1      AGAAGCTGGTGTTTGTGTATCTACTAGTGGTAGATGGGTACTTAACAATGATTATTACAG
MT192772.1      AGAAGCTGGTGTTTGTGTATCTACTAGTGGTAGATGGGTACTTAACAATGATTATTACAG
MT192759.1      AGAAGCTGGTGTTTGTGTATCTACTAGTGGTAGATGGGTACTTAACAATGATTATTACAG
MT126808.1      AGAAGCTGGTGTTTGTGTATCTACTAGTGGTAGATGGGTACTTAACAATGATTATTACAG
MT093571.1      AGAAGCTGGTGTTTGTGTATCTACTAGTGGTAGGTGGGTACTTAACAATGATTATTACAG
MT007544.1      AGAAGCTGGTGTTTGTGTATCTACTAGTGGTAGATGGGTACTTAACAATGATTATTACAG
                ********************************* **************************

                                                                      9477
MT233523.1      TGTAGCTATCGTAGTAACATGCCTTGCCTACTATTTTATGAGGTTTAGAAGAGCTTATGG
MT135043.1      TGTAGCTATCGTAGTAACATGCCTTGCCTACTATTTTATGAGGTTTAGAAGAGCTTTTGG
MT012098.1      TGTAGCTATCGTAGTAACATGCCTTGCCTACTATTTTATGAGGTTTAGAAGAGCTTTTGG
MT072688.1      TGTAGCTATCGTAGTAACATGCCTTGCCTACTATTTTATGAGGTTTAGAAGAGCTTTTGG
MT027064.1      TGTAGCTATCGTAGTAACATGCCTTGCCTACTATTTTATGAGGTTTAGAAGAGCTTTTGG
MT020781.2      TGTAGCTATCGTAGTAACATGCCTTGCCTACTATTTTATGAGGTTTAGAAGAGCTTTTGG
LC529905.1      TGTAGCTATCGTAGTAACATGCCTTGCCTACTATTTTATGAGGTTTAGAAGAGCTTTTGG
MT066156.1      TGTAGCTATCGTAGTAACATGCCTTGCCTACTATTTTATGAGGTTTAGAAGAGCTTTTGG
MT192772.1      TGTAGCTATCGTAGTAACATGCCTTGCCTACTATTTTATGAGGTTTAGAAGAGCTTTTGG
MT192759.1      TGTAGCTATCGTAGTAACATGCCTTGCCTACTATTTTATGAGGTTTAGAAGAGCTTTTGG
MT126808.1      TGTAGCTATCGTAGTAACATGCCTTGCCTACTATTTTATGAGGTTTAGAAGAGCTTTTGG
MT093571.1      TGTAGCTATCGTAGTAACATGCCTTGCCTACTATTTTATGAGGTTTAGAAGAGCTTTTGG
MT007544.1      TGTAGCTATCGTAGTAACATGCCTTGCCTACTATTTTATGAGGTTTAGAAGAGCTTTTGG
                ******************************************************** ***


                                             10232
MT233523.1      GCTTAACCCTAATTATGAAGATTTACTCATTCGTAAGTCTAATCATAATTTCTTGGTACA
MT135043.1      GCTTAACCCTAATTATGAAGATTTACTCATTCGTAAGTCTAATCATAATTTCTTGGTACA
MT012098.1      GCTTAACCCTAATTATGAAGATTTACTCATTCGTAAGTCTAATCATAATTTCTTGGTACA
MT072688.1      GCTTAACCCTAATTATGAAGATTTACTCATTCGTAAGTCTAATCATAATTTCTTGGTACA
MT027064.1      GCTTAACCCTAATTATGAAGATTTACTCATTCGTAAGTCTAATCATAATTTCTTGGTACA
MT020781.2      GCTTAACCCTAATTATGAAGATTTACTCATTCGTAAGTCTAATCATAATTTCTTGGTACA
LC529905.1      GCTTAACCCTAATTATGAAGATTTACTCATTCGTAAGTCTAATCATAATTTCTTGGTACA
MT066156.1      GCTTAACCCTAATTATGAAGATTTACTCATTCGTAAGTCTAATCATAATTTCTTGGTACA
MT192772.1      GCTTAACCCTAATTATGAAGATTTACTCATTTGTAAGTCTAATCATAATTTCTTGGTACA
MT192759.1      GCTTAACCCTAATTATGAAGATTTACTCATTCGTAAGTCTAATCATAATTTCTTGGTACA
MT126808.1      GCTTAACCCTAATTATGAAGATTTACTCATTCGTAAGTCTAATCATAATTTCTTGGTACA
MT093571.1      GCTTAACCCTAATTATGAAGATTTACTCATTCGTAAGTCTAATCATAATTTCTTGGTACA
MT007544.1      GCTTAACCCTAATTATGAAGATTTACTCATTCGTAAGTCTAATCATAATTTCTTGGTACA
                ******************************* ****************************

                                                         11083
MT233523.1      AGTTTTAGTCCAGAGTACTCAATGGTCTTTGTTCTTTTTTTTGTATGAAAATGCCTTTTT
MT135043.1      AGTTTTAGTCCAGAGTACTCAATGGTCTTTGTTCTTTTTTTTGTATGAAAATGCCTTTTT
MT012098.1      AGTTTTAGTCCAGAGTACTCAATGGTCTTTGTTCTTTTTTTTGTATGAAAATGCCTTTTT
MT072688.1      AGTTTTAGTCCAGAGTACTCAATGGTCTTTGTTCTTTTTTTTGTATGAAAATGCCTTTTT
MT027064.1      AGTTTTAGTCCAGAGTACTCAATGGTCTTTGTTCTTTTTTTTGTATGAAAATGCCTTTTT
MT020781.2      AGTTTTAGTCCAGAGTACTCAATGGTCTTTGTTCTTTTTTTTGTATGAAAATGCCTTTTT
LC529905.1      AGTTTTAGTCCAGAGTACTCAATGGTCTTTGTTCTTTTTTTTGTATGAAAATGCCTTTTT
MT066156.1      AGTTTTAGTCCAGAGTACTCAATGGTCTTTGTTCTTTTTTTTNTATGAAAATGCCTTTTT
MT192772.1      AGTTTTAGTCCAGAGTACTCAATGGTCTTTGTTCTTTTTTTTGTATGAAAATGCCTTTTT
MT192759.1      AGTTTTAGTCCAGAGTACTCAATGGTCTTTGTTCTTTTTTTTGTATGAAAATGCCTTTTT
MT126808.1      AGTTTTAGTCCAGAGTACTCAATGGTCTTTGTTCTTTTTTTTTTATGAAAATGCCTTTTT
MT093571.1      AGTTTTAGTCCAGAGTACTCAATGGTCTTTGTTCTTTTTTTTGTATGAAAATGCCTTTTT
MT007544.1      AGTTTTAGTCCAGAGTACTCAATGGTCTTTGTTCTTTTTTTTGTATGAAAATGCCTTTTT
                ****************************************** *****************

                                      13225/26
MT233523.1      GGAAGCCAATATGGATCAAGAATCCTTTGGTGGTGCATCGTGTTGTCTGTACTGCCGTTG
MT135043.1      GGAAGCCAATATGGATCAAGAATCCTTTGGTGGTGCATCGTGTTGTCTGTACTGCCGTTG
MT012098.1      GGAAGCCAATATGGATCAAGAATCCTTTGGTGGTGCATCGTGTTGTCTGTACTGCCGTTG
MT072688.1      GGAAGCCAATATGGATCAAGAATCCTTTGGTGGTGCATCGTGTTGTCTGTACTGCCGTTG
MT027064.1      GGAAGCCAATATGGATCAAGAATCCTTTGGTGGTGCATCGTGTTGTCTGTACTGCCGTTG
MT020781.2      GGAAGCCAATATGGATCAAGAATCCTTTGGTGGTGCATCGTGTTGTCTGTACTGCCGTTG
LC529905.1      GGAAGCCAATATGGATCAAGAATCCTTTGGTGGTGCATCGTGTTGTCTGTACTGCCGTTG
MT066156.1      GGAAGCCAATATGGATCAAGAATCCTTTGGTGGTGCATCGTGTTGTCTGTACTGCCGTTG
MT192772.1      GGAAGCCAATATGGATCAAGAATCCTTTGGTGGTGCATCGTGTTGTCTGTACTGCCGTTG
MT192759.1      GGAAGCCAATATGGATCAAGAATCCTTTGGTGGTGCATCGTGTTGTCTGTACTGCCGTTG
MT126808.1      GGAAGCCAATATGGATCAAGAATCCTTTGGTGGTGCATCGTGTTGTCTGTACTGCCGTTG
MT093571.1      GGAAGCCAATATGGATCAAGAATCGCTTGGTGGTGCATCGTGTTGTCTGTACTGCCGTTG
MT007544.1      GGAAGCCAATATGGATCAAGAATCCTTTGGTGGTGCATCGTGTTGTCTGTACTGCCGTTG
                ************************  **********************************

                              14657
MT233523.1      CTTACTAACAATGTTGCTTTTCAAACTGTCAAACCCGGTAATTTTAACAAAGACTTCTAT
MT135043.1      CTTACTAACAATGTTGCTTTTCAAACTGTCAAACCCGGTAATTTTAACAAAGACTTCTAT
MT012098.1      CTTACTAACAATGTTGTTTTTCAAACTGTCAAACCCGGTAATTTTAACAAAGACTTCTAT
MT072688.1      CTTACTAACAATGTTGCTTTTCAAACTGTCAAACCCGGTAATTTTAACAAAGACTTCTAT
MT027064.1      CTTACTAACAATGTTGCTTTTCAAACTGTCAAACCCGGTAATTTTAACAAAGACTTCTAT
MT020781.2      CTTACTAACAATGTTGCTTTTCAAACTGTCAAACCCGGTAATTTTAACAAAGACTTCTAT
LC529905.1      CTTACTAACAATGTTGCTTTTCAAACTGTCAAACCCGGTAATTTTAACAAAGACTTCTAT
MT066156.1      CTTACTAACAATGTTGCTTTTCAAACTGTCAAACCCGGTAATTTTAACAAAGACTTCTAT
MT192772.1      CTTACTAACAATGTTGCTTTTCAAACTGTCAAACCCGGTAATTTTAACAAAGACTTCTAT
MT192759.1      CTTACTAACAATGTTGCTTTTCAAACTGTCAAACCCGGTAATTTTAACAAAGACTTCTAT
MT126808.1      CTTACTAACAATGTTGCTTTTCAAACTGTCAAACCCGGTAATTTTAACAAAGACTTCTAT
MT093571.1      CTTACTAACAATGTTGCTTTTCAAACTGTCAAACCCGGTAATTTTAACAAAGACTTCTAT
MT007544.1      CTTACTAACAATGTTGCTTTTCAAACTGTCAAACCCGGTAATTTTAACAAAGACTTCTAT
                **************** *******************************************


                                                                                                    14805
MT233523.1      TTCTTTGCTCAGGATGGTAATGCTGCTATCAGCGATTATGACTATTATCGTTATAATCTA
MT135043.1      TTCTTTGCTCAGGATGGTAATGCTGCTATCAGCGATTATGACTACTATCGTTATAATCTA
MT012098.1      TTCTTTGCTCAGGATGGTAATGCTGCTATCAGCGATTATGACTACTATCGTTATAATCTA
MT072688.1      TTCTTTGCTCAGGATGGTAATGCTGCTATCAGCGATTATGACTACTATCGTTATAATCTA
MT027064.1      TTCTTTGCTCAGGATGGTAATGCTGCTATCAGCGATTATGACTACTATCGTTATAATCTA
MT020781.2      TTCTTTGCTCAGGATGGTAATGCTGCTATCAGCGATTATGACTACTATCGTTATAATCTA
LC529905.1      TTCTTTGCTCAGGATGGTAATGCTGCTATCAGCGATTATGACTACTATCGTTATAATCTA
MT066156.1      TTCTTTGCTCAGGATGGTAATGCTGCTATCAGCGATTATGACTACTATCGTTATAATCTA
MT192772.1      TTCTTTGCTCAGGATGGTAATGCTGCTATCAGCGATTATGACTACTATCGTTATAATCTA
MT192759.1      TTCTTTGCTCAGGATGGTAATGCTGCTATCAGCGATTATGACTACTATCGTTATAATCTA
MT126808.1      TTCTTTGCTCAGGATGGTAATGCTGCTATCAGCGATTATGACTATTATCGTTATAATCTA
MT093571.1      TTCTTTGCTCAGGATGGTAATGCTGCTATCAGCGATTATGACTACTATCGTTATAATCTA
MT007544.1      TTCTTTGCTCAGGATGGTAATGCTGCTATCAGCGATTATGACTACTATCGTTATAATCTA
                ******************************************** ***************

                                     15324
MT233523.1      AAATGTGATAGAGCCATGCCTAACATGCTTAGAATTATGGCCTCACTTGTTCTTGCTCGC
MT135043.1      AAATGTGATAGAGCCATGCCTAACATGCTTAGAATTATGGCCTCACTTGTTCTTGCTCGC
MT012098.1      AAATGTGATAGAGCCATGCCTAACATGCTTAGAATTATGGCCTCACTTGTTCTTGCTCGC
MT072688.1      AAATGTGATAGAGCCATGCCTAACATGCTTAGAATTATGGCCTCACTTGTTCTTGCTCGC
MT027064.1      AAATGTGATAGAGCCATGCCTAACATGCTTAGAATTATGGCCTCACTTGTTCTTGCTCGC
MT020781.2      AAATGTGATAGAGCCATGCCTAACATGCTTAGAATTATGGCCTCACTTGTTCTTGCTCGC
LC529905.1      AAATGTGATAGAGCCATGCCTAATATGCTTAGAATTATGGCCTCACTTGTTCTTGCTCGC
MT066156.1      AAATGTGATAGAGCCATGCCTAACATGCTTAGAATTATGGCCTCACTTGTTCTTGCTCGC
MT192772.1      AAATGTGATAGAGCCATGCCTAACATGCTTAGAATTATGGCCTCACTTGTTCTTGCTCGC
MT192759.1      AAATGTGATAGAGCCATGCCTAACATGCTTAGAATTATGGCCTCACTTGTTCTTGCTCGC
MT126808.1      AAATGTGATAGAGCCATGCCTAACATGCTTAGAATTATGGCCTCACTTGTTCTTGCTCGC
MT093571.1      AAATGTGATAGAGCCATGCCTAACATGCTTAGAATTATGGCCTCACTTGTTCTTGCTCGC
MT007544.1      AAATGTGATAGAGCCATGCCTAACATGCTTAGAATTATGGCCTCACTTGTTCTTGCTCGC
                *********************** ************************************

                                        17247
MT233523.1      AAATGTAGTAGAATTATACCTGCACGTGCTCGTGTAGAGTGTTTTGATAAATTCAAAGTG
MT135043.1      AAATGTAGTAGAATTATACCTGCACGTGCTCGTGTAGAGTGTTTTGATAAATTCAAAGTG
MT012098.1      AAATGTAGTAGAATTATACCTGCACGTGCTCGTGTAGAGTGTTTTGATAAATTCAAAGTG
MT072688.1      AAATGTAGTAGAATTATACCTGCACGTGCTCGTGTAGAGTGTTTTGATAAATTCAAAGTG
MT027064.1      AAATGTAGTAGAATTATACCTGCACGTGCTCGTGTAGAGTGTTTTGATAAATTCAAAGTG
MT020781.2      AAATGTAGTAGAATTATACCTGCACGTGCTCGTGTAGAGTGTTTTGATAAATTCAAAGTG
LC529905.1      AAATGTAGTAGAATTATACCTGCACGTGCTCGTGTAGAGTGTTTTGATAAATTCAAAGTG
MT066156.1      AAATGTAGTAGAATTATACCTGCACGTGCTCGTGTAGAGTGTTTTGATAAATTCAAAGTG
MT192772.1      AAATGTAGTAGAATTATACCTGCACGTGCTCGTGTAGAGTGTTTTGATAAATTCAAAGTG
MT192759.1      AAATGTAGTAGAATTATACCTGCACGTGCTCGTGTAGAGTGTTTTGATAAATTCAAAGTG
MT126808.1      AAATGTAGTAGAATTATACCTGCACGCGCTCGTGTAGAGTGTTTTGATAAATTCAAAGTG
MT093571.1      AAATGTAGTAGAATTATACCTGCACGTGCTCGTGTAGAGTGTTTTGATAAATTCAAAGTG
MT007544.1      AAATGTAGTAGAATTATACCTGCACGTGCTCGTGTAGAGTGTTTTGATAAATTCAAAGTG
                ************************** *********************************

                                              17373/76
MT233523.1      GATATAGTTGTCTTTGATGAAATTTCAATGGCCACAAATTATGATTTGAGTGTTGTCAAT
MT135043.1      GATATAGTTGTCTTTGATGAAATTTCAATGGCCACAAATTATGATTTGAGTGTTGTCAAT
MT012098.1      GATATAGTTGTCTTTGATGAAATTTCAATGGCTACAAATTATGATTTGAGTGTTGTCAAT
MT072688.1      GATATAGTTGTCTTTGATGAAATTTCAATGGCCACAAATTATGATTTGAGTGTTGTCAAT
MT027064.1      GATATAGTTGTCTTTGATGAAATTTCAATGGCCACAAATTATGATTTGAGTGTTGTCAAT
MT020781.2      GATATAGTTGTCTTTGATGAAATTTCAATGGCCACAAATTATGATTTGAGTGTTGTCAAT
LC529905.1      GATATAGTTGTCTTTGATGAAATTTCAATGGCCACAAATTATGATTTGAGTGTTGTCAAT
MT066156.1      GATATAGTTGTCTTTGATGAAATTTCAATGGCCACAAATTATGATTTGAGTGTTGTCAAT
MT192772.1      GATATAGTTGTCTTTGATGAAATTTCAATGGCCACAAATTATGATTTGAGTGTTGTCAAT
MT192759.1      GATATAGTTGTCTTTGATGAAATTTCAATGGCCACAAATTATGATTTGAGTGTTGTCAAT
MT126808.1      GATATAGTTGTCTTTGATGAAATTTCAATGGCCACAAATTATGATTTGAGTGTTGTCAAT
MT093571.1      GATATAGTTGTCTTTGATGAAATTTCAATGGCCACGAATTATGATTTGAGTGTTGTCAAT
MT007544.1      GATATAGTTGTCTTTGATGAAATTTCAATGGCCACAAATTATGATTTGAGTGTTGTCAAT
                ******************************** ** ************************


                                                           19065
MT233523.1      GTTCTTCACGACATTGGTAACCCTAAAGCTATTAAGTGTGTACCTCAAGCTGATGTAGAA
MT135043.1      GTTCTTCACGACATTGGTAACCCTAAAGCTATTAAGTGTGTACCTCAAGCTGATGTAGAA
MT012098.1      GTTCTTCACGACATTGGTAACCCTAAAGCTATTAAGTGTGTACCTCAAGCTGATGTAGAA
MT072688.1      GTTCTTCACGACATTGGTAACCCTAAAGCTATTAAGTGTGTACCTCAAGCTGATGTAGAA
MT027064.1      GTTCTTCACGACATTGGTAACCCTAAAGCTATTAAGTGTGTACCTCAAGCTGATGTAGAA
MT020781.2      GTTCTTCACGACATTGGTAACCCTAAAGCTATTAAGTGTGTACCTCAAGCTGATGTAGAA
LC529905.1      GTTCTTCACGACATTGGTAACCCTAAAGCTATTAAGTGTGTACCTCAAGCTGATGTAGAA
MT066156.1      GTTCTTCACGACATTGGTAACCCTAAAGCTATTAAGTGTGTACCTCAAGCTGATGTAGAA
MT192772.1      GTTCTTCACGACATTGGTAACCCTAAAGCTATTAAGTGTGTACCTCAAGCTGATGTAGAA
MT192759.1      GTTCTTCACGACATTGGTAACCCTAAAGCTATTAAGTGTGTACCTCAAGCTGATGTAGAA
MT126808.1      GTTCTTCACGACATTGGTAACCCTAAAGCTATTAAGTGTGTACCTCAAGCTGATGTAGAA
MT093571.1      GTTCTTCACGACATTGGTAACCCTAAAGCTATTAAGTGTGTACCTCAAGCTGATGTAGAA
MT007544.1      GTTCTTCACGACATTGGTAACCCTAAAGCTATTAAGTGTGTACCCCAAGCTGATGTAGAA
                ******************************************** ***************

                                                            21707
MT233523.1      ACGTGGTGTTTATTACCCTGACAAAGTTTTCAGATCCTCAGTTTTACATTCAACTCAGGA
MT135043.1      ACGTGGTGTTTATTACCCTGACAAAGTTTTCAGATCCTCAGTTTTACATTCAACTCAGGA
MT012098.1      ACGTGGTGTTTATTACCCTGACAAAGTTTTCAGATCCTCAGTTTTACATTCAACTCAGGA
MT072688.1      ACGTGGTGTTTATTACCCTGACAAAGTTTTCAGATCCTCAGTTTTACATTCAACTCAGGA
MT027064.1      ACGTGGTGTTTATTACCCTGACAAAGTTTTCAGATCCTCAGTTTTATATTCAACTCAGGA
MT020781.2      ACGTGGTGTTTATTACCCTGACAAAGTTTTCAGATCCTCAGTTTTATATTCAACTCAGGA
LC529905.1      ACGTGGTGTTTATTACCCTGACAAAGTTTTCAGATCCTCAGTTTTACATTCAACTCAGGA
MT066156.1      ACGTGGTGTTTATTACCCTGACAAAGTTTTCAGATCCTCAGTTTTACATTCAACTCAGGA
MT192772.1      ACGTGGTGTTTATTACCCTGACAAAGTTTTCAGATCCTCAGTTTTACATTCAACTCAGGA
MT192759.1      ACGTGGTGTTTATTACCCTGACAAAGTTTTCAGATCCTCAGTTTTACATTCAACTCAGGA
MT126808.1      ACGTGGTGTTTATTACCCTGACAAAGTTTTCAGATCCTCAGTTTTACATTCAACTCAGGA
MT093571.1      ACGTGGTGTTTATTACCCTGACAAAGTTTTCAGATCCTCAGTTTTACATTCAACTCAGGA
MT007544.1      ACGTGGTGTTTATTACCCTGACAAAGTTTTCAGATCCTCAGTTTTACATTCAACTCAGGA
                ********************************************** *************

                                               21964-66
MT233523.1      TCAATTTTGTAATGATCCATTTTTGGGTGTTTATTACCACAAAAACAACAAAAGTTGGAT
MT135043.1      TCAATTTTGTAATGATCCATTTTTGGGTGTTTATTACCACAAAAACAACAAAAGTTGGAT
MT012098.1      TCAATTTTGTAATGATCCATTTTTGGGTGTTTA---CCACAAAAACAACAAAAGTTGGAT
MT072688.1      TCAATTTTGTAATGATCCATTTTTGGGTGTTTATTACCACAAAAACAACAAAAGTTGGAT
MT027064.1      TCAATTTTGTAATGATCCATTTTTGGGTGTTTATTACCACAAAAACAACAAAAGTTGGAT
MT020781.2      TCAATTTTGTAATGATCCATTTTTGGGTGTTTATTACCACAAAAACAACAAAAGTTGGAT
LC529905.1      TCAATTTTGTAATGATCCATTTTTGGGTGTTTATTACCACAAAAACAACAAAAGTTGGAT
MT066156.1      TCAATTTTGTAATGATCCATTTTTGGGTGTTTATTACCACAAAAACAACAAAAGTTGGAT
MT192772.1      TCAATTTTGTAATGATCCATTTTTGGGTGTTTATTACCACAAAAACAACAAAAGTTGGAT
MT192759.1      TCAATTTTGTAATGATCCATTTTTGGGTGTTTATTACCACAAAAACAACAAAAGTTGGAT
MT126808.1      TCAATTTTGTAATGATCCATTTTTGGGTGTTTATTACCACAAAAACAACAAAAGTTGGAT
MT093571.1      TCAATTTTGTAATGATCCATTTTTGGGTGTTTATTACCACAAAAACAACAAAAGTTGGAT
MT007544.1      TCAATTTTGTAATGATCCATTTTTGGGTGTTTATTACCACAAAAACAACAAAAGTTGGAT
                *********************************   ************************

                                                         22303
MT233523.1      TAACATCACTAGGTTTCAAACTTTACTTGCTTTACATAGAAGTTATTTGACTCCTGGTGA
MT135043.1      TAACATCACTAGGTTTCAAACTTTACTTGCTTTACATAGAAGTTATTTGACTCCTGGTGA
MT012098.1      TAACATCACTAGGTTTCAAACTTTACTTGCTTTACATAGAAGTTATTTGACTCCTGGTGA
MT072688.1      TAACATCACTAGGTTTCAAACTTTACTTGCTTTACATAGAAGTTATTTGACTCCTGGTGA
MT027064.1      TAACATCACTAGGTTTCAAACTTTACTTGCTTTACATAGAAGTTATTTGACTCCTGGTGA
MT020781.2      TAACATCACTAGGTTTCAAACTTTACTTGCTTTACATAGAAGTTATTTGACTCCTGGTGA
LC529905.1      TAACATCACTAGGTTTCAAACTTTACTTGCTTTACATAGAAGTTATTTGACTCCTGGTGA
MT066156.1      TAACATCACTAGGTTTCAAACTTTACTTGCTTTACATAGAAGTTATTTGACTCCTGGTGA
MT192772.1      TAACATCACTAGGTTTCAAACTTTACTTGCTTTACATAGAAGTTATTTGACTCCTGGTGA
MT192759.1      TAACATCACTAGGTTTCAAACTTTACTTGCTTTACATAGAAGTTATTTGACTCCTGGTGA
MT126808.1      TAACATCACTAGGTTTCAAACTTTACTTGCTTTACATAGAAGTTATTTGACTCCTGGTGA
MT093571.1      TAACATCACTAGGTTTCAAACTTTACTTGCTTTACATAGAAGTTATTTGACTCCTGGTGA
MT007544.1      TAACATCACTAGGTTTCAAACTTTACTTGCTTTACATAGAAGGTATTTGACTCCTGGTGA
                ****************************************** *****************


                                                          22785
MT233523.1      TAATGTCTATGCAGATTCATTTGTAATTAGAGGTGATGAAGTCAGACAAATCGCTCCAGG
MT135043.1      TAATGTCTATGCAGATTCATTTGTAATTAGAGGTGATGAAGTCAGACAAATCGCTCCAGG
MT012098.1      TAATGTCTATGCAGATTCATTTGTAATTAGAGGTGATGAAGTCATACAAATCGCTCCAGG
MT072688.1      TAATGTCTATGCAGATTCATTTGTAATTAGAGGTGATGAAGTCAGACAAATCGCTCCAGG
MT027064.1      TAATGTCTATGCAGATTCATTTGTAATTAGAGGTGATGAAGTCAGACAAATCGCTCCAGG
MT020781.2      TAATGTCTATGCAGATTCATTTGTAATTAGAGGTGATGAAGTCAGACAAATCGCTCCAGG
LC529905.1      TAATGTCTATGCAGATTCATTTGTAATTAGAGGTGATGAAGTCAGACAAATCGCTCCAGG
MT066156.1      TAATGTCTATGCAGATTCATTTGTAATTAGAGGTGATGAAGTCAGACAAATCGCTCCAGG
MT192772.1      TAATGTCTATGCAGATTCATTTGTAATTAGAGGTGATGAAGTCAGACAAATCGCTCCAGG
MT192759.1      TAATGTCTATGCAGATTCATTTGTAATTAGAGGTGATGAAGTCAGACAAATCGCTCCAGG
MT126808.1      TAATGTCTATGCAGATTCATTTGTAATTAGAGGTGATGAAGTCAGACAAATCGCTCCAGG
MT093571.1      TAATGTCTATGCAGATTCATTTGTAATTAGAGGTGATGAAGTCAGACAAATCGCTCCAGG
MT007544.1      TAATGTCTATGCAGATTCATTTGTAATTAGAGGTGATGAAGTCAGACAAATCGCTCCAGG
                ******************************************** ***************

                        23952
MT233523.1      AATTAAAGATTTTGGTGGTTTTAATTTTTCACAAATATTACCAGATCCATCAAAACCAAG
MT135043.1      AATTAAAGATTTTGGTGGTTTTAATTTTTCACAAATATTACCAGATCCATCAAAACCAAG
MT012098.1      AATTAAAGATTTTGGTGGTTTTAATTTTTCACAAATATTACCAGATCCATCAAAACCAAG
MT072688.1      AATTAAAGATTTTGGTGGTTTTAATTTTTCACAAATATTACCAGATCCATCAAAACCAAG
MT027064.1      AATTAAAGATTTTGGTGGTTTTAATTTTTCACAAATATTACCAGATCCATCAAAACCAAG
MT020781.2      AATTAAAGATTTTGGTGGTTTTAATTTTTCACAAATATTACCAGATCCATCAAAACCAAG
LC529905.1      AATTAAAGATTTTGGTGGTTTTAATTTTTCACAAATATTACCAGATCCATCAAAACCAAG
MT066156.1      AATTAAAGATTTTGGTGGTTTTAATTTTTCACAAATATTACCAGATCCATCAAAACCAAG
MT192772.1      AATTAAAGATTTTGGTGGTTTTAATTTTTCACAAATATTACCAGATCCATCAAAACCAAG
MT192759.1      AATTAAAGATTTTGGTGGTTTTAATTTTTCACAAATATTACCAGATCCATCAAAACCAAG
MT126808.1      AATTAAAGATTTTGGTGGTTTTAATTTTTCACAAATATTACCAGATCCATCAAAACCAAG
MT093571.1      AATTAAAGATTGTGGTGGTTTTAATTTTTCACAAATATTACCAGATCCATCAAAACCAAG
MT007544.1      AATTAAAGATTTTGGTGGTTTTAATTTTTCACAAATATTACCAGATCCATCAAAACCAAG
                *********** ************************************************

                                               24034
MT233523.1      CAAGAGGTCATTTATTGAAGATCTACTTTTCAACAAAGTGACACTTGCAGATGCTGGCTT
MT135043.1      CAAGAGGTCATTTATTGAAGATCTACTTTTCAACAAAGTGACACTTGCAGATGCTGGCTT
MT012098.1      CAAGAGGTCATTTATTGAAGATCTACTTTTCAACAAAGTGACACTTGCAGATGCTGGCTT
MT072688.1      CAAGAGGTCATTTATTGAAGATCTACTTTTCAATAAAGTGACACTTGCAGATGCTGGCTT
MT027064.1      CAAGAGGTCATTTATTGAAGATCTACTTTTCAACAAAGTGACACTTGCAGATGCTGGCTT
MT020781.2      CAAGAGGTCATTTATTGAAGATCTACTTTTCAACAAAGTGACACTTGCAGATGCTGGCTT
LC529905.1      CAAGAGGTCATTTATTGAAGATCTACTTTTCAACAAAGTGACACTTGCAGATGCTGGCTT
MT066156.1      CAAGAGGTCATTTATTGAAGATCTACTTTTCAACAAAGTGACACTTGCAGATGCTGGCTT
MT192772.1      CAAGAGGTCATTTATTGAAGATCTACTTTTCAACAAAGTGACACTTGCAGATGCTGGCTT
MT192759.1      CAAGAGGTCATTTATTGAAGATCTACTTTTCAACAAAGTGACACTTGCAGATGCTGGCTT
MT126808.1      CAAGAGGTCATTTATTGAAGATCTACTTTTCAACAAAGTGACACTTGCAGATGCTGGCTT
MT093571.1      CAAGAGGTCATTTATTGAAGATCTACTTTTCAACAAAGTGACACTTGCAGATGCTGGCTT
MT007544.1      CAAGAGGTCATTTATTGAAGATCTACTTTTCAACAAAGTGACACTTGCAGATGCTGGCTT
                ********************************* **************************

                       25810
MT233523.1      AACCCATTACTTTATGATGCCAACTATTTTCTTTGCTGGCATACTAATTGTTACGACTAT
MT135043.1      AACCCATTACTTTATGATGCCAACTATTTTCTTTGCTGGCATACTAATTGTTACGACTAT
MT012098.1      AACCCATTACTTTATGATGCCAACTATTTTCTTTGCTGGCATACTAATTGTTACGACTAT
MT072688.1      AACCCATTACTTTATGATGCCAACTATTTTCTTTGCTGGCATACTAATTGTTACGACTAT
MT027064.1      AACCCATTACTTTATGATGCCAACTATTTTCTTTGCTGGCATACTAATTGTTACGACTAT
MT020781.2      AACCCATTACTTTATGATGCCAACTATTTTCTTTGCTGGCATACTAATTGTTACGACTAT
LC529905.1      AACCCATTAGTTTATGATGCCAACTATTTTCTTTGCTGGCATACTAATTGTTACGACTAT
MT066156.1      AACCCATTACTTTATGATGCCAACTATTTTCTTTGCTGGCATACTAATTGTTACGACTAT
MT192772.1      AACCCATTACTTTATGATGCCAACTATTTTCTTTGCTGGCATACTAATTGTTACGACTAT
MT192759.1      AACCCATTACTTTATGATGCCAACTATTTTCTTTGCTGGCATACTAATTGTTACGACTAT
MT126808.1      AACCCATTACTTTATGATGCCAACTATTTTCTTTGCTGGCATACTAATTGTTACGACTAT
MT093571.1      AACCCATTACTTTATGATGCCAACTATTTTCTTTGCTGGCATACTAATTGTTACGACTAT
MT007544.1      AACCCATTACTTTATGATGCCAACTATTTTCTTTGCTGGCATACTAATTGTTACGACTAT
                ********* **************************************************


                                                                      25979
MT233523.1      AGTCCTATTTCTGAACATGACTACCAGATTGGTGGTTATACTGAAAAATGGGAATCTGTA
MT135043.1      AGTCCTATTTCTGAACATGACTACCAGATTGGTGGTTATACTGAAAAATGGGAATCTGGA
MT012098.1      AGTCCTATTTCTGAACATGACTACCAGATTGGTGGTTATACTGAAAAATGGGAATCTGGA
MT072688.1      AGTCCTATTTCTGAACATGACTACCAGATTGGTGGTTATACTGAAAAATGGGAATCTGGA
MT027064.1      AGTCCTATTTCTGAACATGACTACCAGATTGGTGGTTATACTGAAAAATGGGAATCTGGA
MT020781.2      AGTCCTATTTCTGAACATGACTACCAGATTGGTGGTTATACTGAAAAATGGGAATCTGGA
LC529905.1      AGTCCTATTTCTGAACATGACTACCAGATTGGTGGTTATACTGAAAAATGGGAATCTGGA
MT066156.1      AGTCCTATTTCTGAACATGACTACCAGATTGGTGGTTATACTGAAAAATGGGAATCTGGA
MT192772.1      AGTCCTATTTCTGAACATGACTACCAGATTGGTGGTTATACTGAAAAATGGGAATCTGGA
MT192759.1      AGTCCTATTTCTGAACATGACTACCAGATTGGTGGTTATACTGAAAAATGGGAATCTGGA
MT126808.1      AGTCCTATTTCTGAACATGACTACCAGATTGGTGGTTATACTGAAAAATGGGAATCTGGA
MT093571.1      AGTCCTATTTCTGAACATGACTACCAGATTGGTGGTTATACTGAAAAATGGGAATCTGGA
MT007544.1      AGTCCTATTTCTGAACATGACTACCAGATTGGTGGTTATACTGAAAAATGGGAATCTGGA
                ********************************************************** *

                                                         26144
MT233523.1      GTTGATGAGCCTGAAGAACATGTCCAAATTCACACAATCGACGGTTCATCCGGAGTTGTT
MT135043.1      GTTGATGAGCCTGAAGAACATGTCCAAATTCACACAATCGACGGTTCATCCGGAGTTGTT
MT012098.1      GTTGATGAGCCTGAAGAACATGTCCAAATTCACACAATCGACGGTTCATCCGGAGTTGTT
MT072688.1      GTTGATGAGCCTGAAGAACATGTCCAAATTCACACAATCGACGGTTCATCCGGAGTTGTT
MT027064.1      GTTGATGAGCCTGAAGAACATGTCCAAATTCACACAATCGACGGTTCATCCGGAGTTGTT
MT020781.2      GTTGATGAGCCTGAAGAACATGTCCAAATTCACACAATCGACGGTTCATCCGGAGTTGTT
LC529905.1      GTTGATGAGCCTGAAGAACATGTCCAAATTCACACAATCGACGGTTCATCCGGAGTTGTT
MT066156.1      GTTGATGAGCCTGAAGAACATGTCCAAATTCACACAATCGACGTTTCATCCGGAGTTGTT
MT192772.1      GTTGATGAGCCTGAAGAACATGTCCAAATTCACACAATCGACGGTTCATCCGGAGTTGTT
MT192759.1      GTTGATGAGCCTGAAGAACATGTCCAAATTCACACAATCGACGGTTCATCCGGAGTTGTT
MT126808.1      GTTGATGAGCCTGAAGAACATGTCCAAATTCACACAATCGACGTTTCATCCGGAGTTGTT
MT093571.1      GTTGATGAGCCTGAAGAACATGTCCAAATTCACACAATCGACGTTTCATCCGGAGTTGTT
MT007544.1      GTTGATGAGCCTGAAGAACATGTCCAAATTCACACAATCGACGTTTCATCCGGAGTTGTT
                ******************************************* ****************

                 28144
MT233523.1      GTTCACCTTTTACAATTAATTGCCAGGAACCTAAATTGGGTAGTCTTGTAGTGCGTTGTT
MT135043.1      GTTCACCTTTTACAATTAATTGCCAGGAACCTAAATTGGGTAGTCTTGTAGTGCGTTGTT
MT012098.1      GTTTACCTTTTACAATTAATTGCCAGGAACCTAAATTGGGTAGTCTTGTAGTGCGTTGTT
MT072688.1      GTTTACCTTTTACAATTAATTGCCAGGAACCTAAATTGGGTAGTCTTGTAGTGCGTTGTT
MT027064.1      GTTTACCTTTTACAATTAATTGCCAGGAACCTAAATTGGGTAGTCTTGTAGTGCGTTGTT
MT020781.2      GTTTACCTTTTACAATTAATTGCCAGGAACCTAAATTGGGTAGTCTTGTAGTGCGTTGTT
LC529905.1      GTTTACCTTTTACAATTAATTGCCAGGAACCTAAATTGGGTAGTCTTGTAGTGCGTTGTT
MT066156.1      GTTTACCTTTTACAATTAATTGCCAGGAACCTAAATTGGGTAGTCTTGTAGTGCGTTGTT
MT192772.1      GTTTACCTTTTACAATTAATTGCCAGGAACCTAAATTGGGTAGTCTTGTAGTGCGTTGTT
MT192759.1      GTTTACCTTTTACAATTAATTGCCAGGAACCTAAATTGGGTAGTCTTGTAGTGCGTTGTT
MT126808.1      GTTTACCTTTTACAATTAATTGCCAGGAACCTAAATTGGGTAGTCTTGTAGTGCGTTGTT
MT093571.1      GTTTACCTTTTACAATTAATTGCCAGGAACCTAAATTGGGTAGTCTTGTAGTGCGTTGTT
MT007544.1      GTTTACCTTTTACAATTAATTGCCAGGAACCTAAATTGGGTAGTCTTGTAGTGCGTTGTT
                *** ********************************************************

                                                  28657
MT233523.1      GCCAGAAGCTGGACTTCCCTATGGTGCTAACAAAGATGGCATCATATGGGTTGCAACTGA
MT135043.1      GCCAGAAGCTGGACTTCCCTATGGTGCTAACAAAGACGGCATCATATGGGTTGCAACTGA
MT012098.1      GCCAGAAGCTGGACTTCCCTATGGTGCTAACAAAGACGGCATCATATGGGTTGCAACTGA
MT072688.1      GCCAGAAGCTGGACTTCCCTATGGTGCTAACAAAGACGGCATCATATGGGTTGCAACTGA
MT027064.1      GCCAGAAGCTGGACTTCCCTATGGTGCTAACAAAGACGGCATCATATGGGTTGCAACTGA
MT020781.2      GCCAGAAGCTGGACTTCCCTATGGTGCTAACAAAGACGGCATCATATGGGTTGCAACTGA
LC529905.1      GCCAGAAGCTGGACTTCCCTATGGTGCTAACAAAGACGGCATCATATGGGTTGCAACTGA
MT066156.1      GCCAGAAGCTGGACTTCCCTATGGTGCTAACAAAGACGGCATCATATGGGTTGCAACTGA
MT192772.1      GCCAGAAGCTGGACTTCCCTATGGTGCTAACAAAGACGGCATCATATGGGTTGCAACTGA
MT192759.1      GCCAGAAGCTGGACTTCCCTATGGTGCTAACAAAGACGGCATCATATGGGTTGCAACTGA
MT126808.1      GCCAGAAGCTGGACTTCCCTATGGTGCTAACAAAGACGGCATCATATGGGTTGCAACTGA
MT093571.1      GCCAGAAGCTGGACTTCCCTATGGTGCTAACAAAGACGGCATCATATGGGTTGCAACTGA
MT007544.1      GCCAGAAGCTGGACTTCCCTATGGTGCTAACAAAGACGGCATCATATGGGTTGCAACTGA
                ************************************ ***********************


                             28863
MT233523.1      TTTAACTCCAGGCAGCAGTAGGGGAACTTCTCCTGCTAGAATGGCTGGCAATGGCGGTGA
MT135043.1      TTCAACTCCAGGCAGCAGTAGGGGAACTTCTCCTGCTAGAATGGCTGGCAATGGCGGTGA
MT012098.1      TTCAACTCCAGGCAGCAGTAGGGGAACTTCTCCTGCTAGAATGGCTGGCAATGGCGGTGA
MT072688.1      TTCAACTCCAGGCAGCAGTAGGGGAACTTCTCCTGCTAGAATGGCTGGCAATGGCGGTGA
MT027064.1      TTCAACTCCAGGCAGCAGTAGGGGAACTTCTCCTGCTAGAATGGCTGGCAATGGCGGTGA
MT020781.2      TTCAACTCCAGGCAGCAGTAGGGGAACTTCTCCTGCTAGAATGGCTGGCAATGGCGGTGA
LC529905.1      TTCAACTCCAGGCAGCAGTAGGGGAACTTCTCCTGCTAGAATGGCTGGCAATGGCGGTGA
MT066156.1      TTCAACTCCAGGCAGCAGTAGGGGAACTTCTCCTGCTAGAATGGCTGGCAATGGCGGTGA
MT192772.1      TTCAACTCCAGGCAGCAGTAGGGGAACTTCTCCTGCTAGAATGGCTGGCAATGGCGGTGA
MT192759.1      TTCAACTCCAGGCAGCAGTAGGGGAACTTCTCCTGCTAGAATGGCTGGCAATGGCGGTGA
MT126808.1      TTCAACTCCAGGCAGCAGTAGGGGAACTTCTCCTGCTAGAATGGCTGGCAATGGCGGTGA
MT093571.1      TTCAACTCCAGGCAGCAGTAGGGGAACTTCTCCTGCTAGAATGGCTGGCAATGGCGGTGA
MT007544.1      TTCAACTCCAGGCAGCAGTAGGGGAACTTCTCCTGCTAGAATGGCTGGCAATGGCGGTGA
                ** *********************************************************


                                  29301/03
MT233523.1      CATCAAATTGGATGACAAAGATCCAAATTTCAAAGATCAAGTCATTTTGCTGAATAAGCA
MT135043.1      CATCAAATTGGATGACAAAGTTCCAAATTTCAAAGATCAAGTCATTTTGCTGAATAAGCA
MT012098.1      CATCAAATTGGATGACAAAGATCCAAATTTCAAAGATCAAGTCATTTTGCTGAATAAGCA
MT072688.1      CATCAAATTGGATGACAAAGATCCAAATTTCAAAGATCAAGTCATTTTGCTGAATAAGCA
MT027064.1      CATCAAATTGGATGACAAAGATCCAAATTTCAAAGATCAAGTCATTTTGCTGAATAAGCA
MT020781.2      CATCAAATTGGATGACAAAGATCCAAATTTCAAAGATCAAGTCATTTTGCTGAATAAGCA
LC529905.1      CATCAAATTGGATGACAAAGATTCAAATTTCAAAGATCAAGTCATTTTGCTGAATAAGCA
MT066156.1      CATCAAATTGGATGACAAAGATCCAAATTTCAAAGATCAAGTCATTTTGCTGAATAAGCA
MT192772.1      CATCAAATTGGATGACAAAGATCCAAATTTCAAAGATCAAGTCATTTTGCTGAATAAGCA
MT192759.1      CATCAAATTGGATGACAAAGATCCAAATTTCAAAGATCAAGTCATTTTGCTGAATAAGCA
MT126808.1      CATCAAATTGGATGACAAAGATCCAAATTTCAAAGATCAAGTCATTTTGCTGAATAAGCA
MT093571.1      CATCAAATTGGATGACAAAGATCCAAATTTCAAAGATCAAGTCATTTTGCTGAATAAGCA
MT007544.1      CATCAAATTGGATGACAAAGATCCAAATTTCAAAGATCAAGTCATTTTGCTGAATAAGCA
                ******************** * *************************************
```
