## Supplementary material for "Decoding the lethal effect of *SARS-CoV-2* (novel coronavirus) strains from global perspective: molecular pathogenesis and evolutionary divergence": Pairwise Sequence Alignment between the first submitted Ancestor viral Strain of Wuhan(NC_045512.2) and the viral Strain of Taiwan

**Supplementary Material 2**

**Pairwise Sequence Alignment between the first submitted Ancestor viral Strain of Wuhan(Submitted 5^th^ Jan, 2020; NC_045512.2) and the viral Strain of Taiwan(MT192759.1)**

```
# Aligned_sequences: 2
# 1: EMBOSS_001 The Ancestor Strain of Wuhan
# 2: EMBOSS_001 Taiwan Strain that we have got as Ancestor Strain
# Matrix: EDNAFULL
# Gap_penalty: 16
# Extend_penalty: 4
#
# Length: 29870
# Identity:   29843/29870 (99.9%)
# Similarity: 29843/29870 (99.9%)
# Gaps:          27/29870 ( 0.1%)
# Score: 149095
#
#
#=======================================

EMBOSS_001         1 ATTAAAGGTTTATACCTTCCCAGGTAACAAACCAACCAACTTTCGATCTC     50 Wuhan Strain
                                                |||||||||||||||||||||||
EMBOSS_001         1 ---------------------------CAAACCAACCAACTTTCGATCTC     23 Taiwan Strain

EMBOSS_001        51 TTGTAGATCTGTTCTCTAAACGAACTTTAAAATCTGTGTGGCTGTCACTC    100
                     ||||||||||||||||||||||||||||||||||||||||||||||||||
EMBOSS_001        24 TTGTAGATCTGTTCTCTAAACGAACTTTAAAATCTGTGTGGCTGTCACTC     73

EMBOSS_001       101 GGCTGCATGCTTAGTGCACTCACGCAGTATAATTAATAACTAATTACTGT    150
                     ||||||||||||||||||||||||||||||||||||||||||||||||||
EMBOSS_001        74 GGCTGCATGCTTAGTGCACTCACGCAGTATAATTAATAACTAATTACTGT    123

EMBOSS_001       151 CGTTGACAGGACACGAGTAACTCGTCTATCTTCTGCAGGCTGCTTACGGT    200
                     ||||||||||||||||||||||||||||||||||||||||||||||||||
EMBOSS_001       124 CGTTGACAGGACACGAGTAACTCGTCTATCTTCTGCAGGCTGCTTACGGT    173

EMBOSS_001       201 TTCGTCCGTGTTGCAGCCGATCATCAGCACATCTAGGTTTCGTCCGGGTG    250
                     ||||||||||||||||||||||||||||||||||||||||||||||||||
EMBOSS_001       174 TTCGTCCGTGTTGCAGCCGATCATCAGCACATCTAGGTTTCGTCCGGGTG    223

EMBOSS_001       251 TGACCGAAAGGTAAGATGGAGAGCCTTGTCCCTGGTTTCAACGAGAAAAC    300
                     ||||||||||||||||||||||||||||||||||||||||||||||||||
EMBOSS_001       224 TGACCGAAAGGTAAGATGGAGAGCCTTGTCCCTGGTTTCAACGAGAAAAC    273

EMBOSS_001       301 ACACGTCCAACTCAGTTTGCCTGTTTTACAGGTTCGCGACGTGCTCGTAC    350
                     ||||||||||||||||||||||||||||||||||||||||||||||||||
EMBOSS_001       274 ACACGTCCAACTCAGTTTGCCTGTTTTACAGGTTCGCGACGTGCTCGTAC    323

EMBOSS_001       351 GTGGCTTTGGAGACTCCGTGGAGGAGGTCTTATCAGAGGCACGTCAACAT    400
                     ||||||||||||||||||||||||||||||||||||||||||||||||||
EMBOSS_001       324 GTGGCTTTGGAGACTCCGTGGAGGAGGTCTTATCAGAGGCACGTCAACAT    373

EMBOSS_001       401 CTTAAAGATGGCACTTGTGGCTTAGTAGAAGTTGAAAAAGGCGTTTTGCC    450
                     ||||||||||||||||||||||||||||||||||||||||||||||||||
EMBOSS_001       374 CTTAAAGATGGCACTTGTGGCTTAGTAGAAGTTGAAAAAGGCGTTTTGCC    423

EMBOSS_001       451 TCAACTTGAACAGCCCTATGTGTTCATCAAACGTTCGGATGCTCGAACTG    500
                     ||||||||||||||||||||||||||||||||||||||||||||||||||
EMBOSS_001       424 TCAACTTGAACAGCCCTATGTGTTCATCAAACGTTCGGATGCTCGAACTG    473

EMBOSS_001       501 CACCTCATGGTCATGTTATGGTTGAGCTGGTAGCAGAACTCGAAGGCATT    550
                     ||||||||||||||||||||||||||||||||||||||||||||||||||
EMBOSS_001       474 CACCTCATGGTCATGTTATGGTTGAGCTGGTAGCAGAACTCGAAGGCATT    523

EMBOSS_001       551 CAGTACGGTCGTAGTGGTGAGACACTTGGTGTCCTTGTCCCTCATGTGGG    600
                     ||||||||||||||||||||||||||||||||||||||||||||||||||
EMBOSS_001       524 CAGTACGGTCGTAGTGGTGAGACACTTGGTGTCCTTGTCCCTCATGTGGG    573

EMBOSS_001       601 CGAAATACCAGTGGCTTACCGCAAGGTTCTTCTTCGTAAGAACGGTAATA    650
                     ||||||||||||||||||||||||||||||||||||||||||||||||||
EMBOSS_001       574 CGAAATACCAGTGGCTTACCGCAAGGTTCTTCTTCGTAAGAACGGTAATA    623

EMBOSS_001       651 AAGGAGCTGGTGGCCATAGTTACGGCGCCGATCTAAAGTCATTTGACTTA    700
                     ||||||||||||||||||||||||||||||||||||||||||||||||||
EMBOSS_001       624 AAGGAGCTGGTGGCCATAGTTACGGCGCCGATCTAAAGTCATTTGACTTA    673

EMBOSS_001       701 GGCGACGAGCTTGGCACTGATCCTTATGAAGATTTTCAAGAAAACTGGAA    750
                     ||||||||||||||||||||||||||||||||||||||||||||||||||
EMBOSS_001       674 GGCGACGAGCTTGGCACTGATCCTTATGAAGATTTTCAAGAAAACTGGAA    723

EMBOSS_001       751 CACTAAACATAGCAGTGGTGTTACCCGTGAACTCATGCGTGAGCTTAACG    800
                     ||||||||||||||||||||||||||||||||||||||||||||||||||
EMBOSS_001       724 CACTAAACATAGCAGTGGTGTTACCCGTGAACTCATGCGTGAGCTTAACG    773

EMBOSS_001       801 GAGGGGCATACACTCGCTATGTCGATAACAACTTCTGTGGCCCTGATGGC    850
                     ||||||||||||||||||||||||||||||||||||||||||||||||||
EMBOSS_001       774 GAGGGGCATACACTCGCTATGTCGATAACAACTTCTGTGGCCCTGATGGC    823

EMBOSS_001       851 TACCCTCTTGAGTGCATTAAAGACCTTCTAGCACGTGCTGGTAAAGCTTC    900
                     ||||||||||||||||||||||||||||||||||||||||||||||||||
EMBOSS_001       824 TACCCTCTTGAGTGCATTAAAGACCTTCTAGCACGTGCTGGTAAAGCTTC    873

EMBOSS_001       901 ATGCACTTTGTCCGAACAACTGGACTTTATTGACACTAAGAGGGGTGTAT    950
                     ||||||||||||||||||||||||||||||||||||||||||||||||||
EMBOSS_001       874 ATGCACTTTGTCCGAACAACTGGACTTTATTGACACTAAGAGGGGTGTAT    923

EMBOSS_001       951 ACTGCTGCCGTGAACATGAGCATGAAATTGCTTGGTACACGGAACGTTCT   1000
                     ||||||||||||||||||||||||||||||||||||||||||||||||||
EMBOSS_001       924 ACTGCTGCCGTGAACATGAGCATGAAATTGCTTGGTACACGGAACGTTCT    973

EMBOSS_001      1001 GAAAAGAGCTATGAATTGCAGACACCTTTTGAAATTAAATTGGCAAAGAA   1050
                     ||||||||||||||||||||||||||||||||||||||||||||||||||
EMBOSS_001       974 GAAAAGAGCTATGAATTGCAGACACCTTTTGAAATTAAATTGGCAAAGAA   1023

EMBOSS_001      1051 ATTTGACACCTTCAATGGGGAATGTCCAAATTTTGTATTTCCCTTAAATT   1100
                     ||||||||||||||||||||||||||||||||||||||||||||||||||
EMBOSS_001      1024 ATTTGACACCTTCAATGGGGAATGTCCAAATTTTGTATTTCCCTTAAATT   1073

EMBOSS_001      1101 CCATAATCAAGACTATTCAACCAAGGGTTGAAAAGAAAAAGCTTGATGGC   1150
                     ||||||||||||||||||||||||||||||||||||||||||||||||||
EMBOSS_001      1074 CCATAATCAAGACTATTCAACCAAGGGTTGAAAAGAAAAAGCTTGATGGC   1123

EMBOSS_001      1151 TTTATGGGTAGAATTCGATCTGTCTATCCAGTTGCGTCACCAAATGAATG   1200
                     ||||||||||||||||||||||||||||||||||||||||||||||||||
EMBOSS_001      1124 TTTATGGGTAGAATTCGATCTGTCTATCCAGTTGCGTCACCAAATGAATG   1173

EMBOSS_001      1201 CAACCAAATGTGCCTTTCAACTCTCATGAAGTGTGATCATTGTGGTGAAA   1250
                     ||||||||||||||||||||||||||||||||||||||||||||||||||
EMBOSS_001      1174 CAACCAAATGTGCCTTTCAACTCTCATGAAGTGTGATCATTGTGGTGAAA   1223

EMBOSS_001      1251 CTTCATGGCAGACGGGCGATTTTGTTAAAGCCACTTGCGAATTTTGTGGC   1300
                     ||||||||||||||||||||||||||||||||||||||||||||||||||
EMBOSS_001      1224 CTTCATGGCAGACGGGCGATTTTGTTAAAGCCACTTGCGAATTTTGTGGC   1273

EMBOSS_001      1301 ACTGAGAATTTGACTAAAGAAGGTGCCACTACTTGTGGTTACTTACCCCA   1350
                     ||||||||||||||||||||||||||||||||||||||||||||||||||
EMBOSS_001      1274 ACTGAGAATTTGACTAAAGAAGGTGCCACTACTTGTGGTTACTTACCCCA   1323

EMBOSS_001      1351 AAATGCTGTTGTTAAAATTTATTGTCCAGCATGTCACAATTCAGAAGTAG   1400
                     ||||||||||||||||||||||||||||||||||||||||||||||||||
EMBOSS_001      1324 AAATGCTGTTGTTAAAATTTATTGTCCAGCATGTCACAATTCAGAAGTAG   1373

EMBOSS_001      1401 GACCTGAGCATAGTCTTGCCGAATACCATAATGAATCTGGCTTGAAAACC   1450
                     ||||||||||||||||||||||||||||||||||||||||||||||||||
EMBOSS_001      1374 GACCTGAGCATAGTCTTGCCGAATACCATAATGAATCTGGCTTGAAAACC   1423

EMBOSS_001      1451 ATTCTTCGTAAGGGTGGTCGCACTATTGCCTTTGGAGGCTGTGTGTTCTC   1500
                     ||||||||||||||||||||||||||||||||||||||||||||||||||
EMBOSS_001      1424 ATTCTTCGTAAGGGTGGTCGCACTATTGCCTTTGGAGGCTGTGTGTTCTC   1473

EMBOSS_001      1501 TTATGTTGGTTGCCATAACAAGTGTGCCTATTGGGTTCCACGTGCTAGCG   1550
                     ||||||||||||||||||||||||||||||||||||||||||||||||||
EMBOSS_001      1474 TTATGTTGGTTGCCATAACAAGTGTGCCTATTGGGTTCCACGTGCTAGCG   1523

EMBOSS_001      1551 CTAACATAGGTTGTAACCATACAGGTGTTGTTGGAGAAGGTTCCGAAGGT   1600
                     ||||||||||||||||||||||||||||||||||||||||||||||||||
EMBOSS_001      1524 CTAACATAGGTTGTAACCATACAGGTGTTGTTGGAGAAGGTTCCGAAGGT   1573

EMBOSS_001      1601 CTTAATGACAACCTTCTTGAAATACTCCAAAAAGAGAAAGTCAACATCAA   1650
                     ||||||||||||||||||||||||||||||||||||||||||||||||||
EMBOSS_001      1574 CTTAATGACAACCTTCTTGAAATACTCCAAAAAGAGAAAGTCAACATCAA   1623

EMBOSS_001      1651 TATTGTTGGTGACTTTAAACTTAATGAAGAGATCGCCATTATTTTGGCAT   1700
                     ||||||||||||||||||||||||||||||||||||||||||||||||||
EMBOSS_001      1624 TATTGTTGGTGACTTTAAACTTAATGAAGAGATCGCCATTATTTTGGCAT   1673

EMBOSS_001      1701 CTTTTTCTGCTTCCACAAGTGCTTTTGTGGAAACTGTGAAAGGTTTGGAT   1750
                     ||||||||||||||||||||||||||||||||||||||||||||||||||
EMBOSS_001      1674 CTTTTTCTGCTTCCACAAGTGCTTTTGTGGAAACTGTGAAAGGTTTGGAT   1723

EMBOSS_001      1751 TATAAAGCATTCAAACAAATTGTTGAATCCTGTGGTAATTTTAAAGTTAC   1800
                     ||||||||||||||||||||||||||||||||||||||||||||||||||
EMBOSS_001      1724 TATAAAGCATTCAAACAAATTGTTGAATCCTGTGGTAATTTTAAAGTTAC   1773

EMBOSS_001      1801 AAAAGGAAAAGCTAAAAAAGGTGCCTGGAATATTGGTGAACAGAAATCAA   1850
                     ||||||||||||||||||||||||||||||||||||||||||||||||||
EMBOSS_001      1774 AAAAGGAAAAGCTAAAAAAGGTGCCTGGAATATTGGTGAACAGAAATCAA   1823

EMBOSS_001      1851 TACTGAGTCCTCTTTATGCATTTGCATCAGAGGCTGCTCGTGTTGTACGA   1900
                     ||||||||||||||||||||||||||||||||||||||||||||||||||
EMBOSS_001      1824 TACTGAGTCCTCTTTATGCATTTGCATCAGAGGCTGCTCGTGTTGTACGA   1873

EMBOSS_001      1901 TCAATTTTCTCCCGCACTCTTGAAACTGCTCAAAATTCTGTGCGTGTTTT   1950
                     ||||||||||||||||||||||||||||||||||||||||||||||||||
EMBOSS_001      1874 TCAATTTTCTCCCGCACTCTTGAAACTGCTCAAAATTCTGTGCGTGTTTT   1923

EMBOSS_001      1951 ACAGAAGGCCGCTATAACAATACTAGATGGAATTTCACAGTATTCACTGA   2000
                     ||||||||||||||||||||||||||||||||||||||||||||||||||
EMBOSS_001      1924 ACAGAAGGCCGCTATAACAATACTAGATGGAATTTCACAGTATTCACTGA   1973

EMBOSS_001      2001 GACTCATTGATGCTATGATGTTCACATCTGATTTGGCTACTAACAATCTA   2050
                     ||||||||||||||||||||||||||||||||||||||||||||||||||
EMBOSS_001      1974 GACTCATTGATGCTATGATGTTCACATCTGATTTGGCTACTAACAATCTA   2023

EMBOSS_001      2051 GTTGTAATGGCCTACATTACAGGTGGTGTTGTTCAGTTGACTTCGCAGTG   2100
                     ||||||||||||||||||||||||||||||||||||||||||||||||||
EMBOSS_001      2024 GTTGTAATGGCCTACATTACAGGTGGTGTTGTTCAGTTGACTTCGCAGTG   2073

EMBOSS_001      2101 GCTAACTAACATCTTTGGCACTGTTTATGAAAAACTCAAACCCGTCCTTG   2150
                     ||||||||||||||||||||||||||||||||||||||||||||||||||
EMBOSS_001      2074 GCTAACTAACATCTTTGGCACTGTTTATGAAAAACTCAAACCCGTCCTTG   2123

EMBOSS_001      2151 ATTGGCTTGAAGAGAAGTTTAAGGAAGGTGTAGAGTTTCTTAGAGACGGT   2200
                     ||||||||||||||||||||||||||||||||||||||||||||||||||
EMBOSS_001      2124 ATTGGCTTGAAGAGAAGTTTAAGGAAGGTGTAGAGTTTCTTAGAGACGGT   2173

EMBOSS_001      2201 TGGGAAATTGTTAAATTTATCTCAACCTGTGCTTGTGAAATTGTCGGTGG   2250
                     ||||||||||||||||||||||||||||||||||||||||||||||||||
EMBOSS_001      2174 TGGGAAATTGTTAAATTTATCTCAACCTGTGCTTGTGAAATTGTCGGTGG   2223

EMBOSS_001      2251 ACAAATTGTCACCTGTGCAAAGGAAATTAAGGAGAGTGTTCAGACATTCT   2300
                     ||||||||||||||||||||||||||||||||||||||||||||||||||
EMBOSS_001      2224 ACAAATTGTCACCTGTGCAAAGGAAATTAAGGAGAGTGTTCAGACATTCT   2273

EMBOSS_001      2301 TTAAGCTTGTAAATAAATTTTTGGCTTTGTGTGCTGACTCTATCATTATT   2350
                     ||||||||||||||||||||||||||||||||||||||||||||||||||
EMBOSS_001      2274 TTAAGCTTGTAAATAAATTTTTGGCTTTGTGTGCTGACTCTATCATTATT   2323

EMBOSS_001      2351 GGTGGAGCTAAACTTAAAGCCTTGAATTTAGGTGAAACATTTGTCACGCA   2400
                     ||||||||||||||||||||||||||||||||||||||||||||||||||
EMBOSS_001      2324 GGTGGAGCTAAACTTAAAGCCTTGAATTTAGGTGAAACATTTGTCACGCA   2373

EMBOSS_001      2401 CTCAAAGGGATTGTACAGAAAGTGTGTTAAATCCAGAGAAGAAACTGGCC   2450
                     ||||||||||||||||||||||||||||||||||||||||||||||||||
EMBOSS_001      2374 CTCAAAGGGATTGTACAGAAAGTGTGTTAAATCCAGAGAAGAAACTGGCC   2423

EMBOSS_001      2451 TACTCATGCCTCTAAAAGCCCCAAAAGAAATTATCTTCTTAGAGGGAGAA   2500
                     ||||||||||||||||||||||||||||||||||||||||||||||||||
EMBOSS_001      2424 TACTCATGCCTCTAAAAGCCCCAAAAGAAATTATCTTCTTAGAGGGAGAA   2473

EMBOSS_001      2501 ACACTTCCCACAGAAGTGTTAACAGAGGAAGTTGTCTTGAAAACTGGTGA   2550
                     ||||||||||||||||||||||||||||||||||||||||||||||||||
EMBOSS_001      2474 ACACTTCCCACAGAAGTGTTAACAGAGGAAGTTGTCTTGAAAACTGGTGA   2523

EMBOSS_001      2551 TTTACAACCATTAGAACAACCTACTAGTGAAGCTGTTGAAGCTCCATTGG   2600
                     ||||||||||||||||||||||||||||||||||||||||||||||||||
EMBOSS_001      2524 TTTACAACCATTAGAACAACCTACTAGTGAAGCTGTTGAAGCTCCATTGG   2573

EMBOSS_001      2601 TTGGTACACCAGTTTGTATTAACGGGCTTATGTTGCTCGAAATCAAAGAC   2650
                     ||||||||||||||||||||||||||||||||||||||||||||||||||
EMBOSS_001      2574 TTGGTACACCAGTTTGTATTAACGGGCTTATGTTGCTCGAAATCAAAGAC   2623

EMBOSS_001      2651 ACAGAAAAGTACTGTGCCCTTGCACCTAATATGATGGTAACAAACAATAC   2700
                     ||||||||||||||||||||||||||||||||||||||||||||||||||
EMBOSS_001      2624 ACAGAAAAGTACTGTGCCCTTGCACCTAATATGATGGTAACAAACAATAC   2673

EMBOSS_001      2701 CTTCACACTCAAAGGCGGTGCACCAACAAAGGTTACTTTTGGTGATGACA   2750
                     ||||||||||||||||||||||||||||||||||||||||||||||||||
EMBOSS_001      2674 CTTCACACTCAAAGGCGGTGCACCAACAAAGGTTACTTTTGGTGATGACA   2723

EMBOSS_001      2751 CTGTGATAGAAGTGCAAGGTTACAAGAGTGTGAATATCACTTTTGAACTT   2800
                     ||||||||||||||||||||||||||||||||||||||||||||||||||
EMBOSS_001      2724 CTGTGATAGAAGTGCAAGGTTACAAGAGTGTGAATATCACTTTTGAACTT   2773

EMBOSS_001      2801 GATGAAAGGATTGATAAAGTACTTAATGAGAAGTGCTCTGCCTATACAGT   2850
                     ||||||||||||||||||||||||||||||||||||||||||||||||||
EMBOSS_001      2774 GATGAAAGGATTGATAAAGTACTTAATGAGAAGTGCTCTGCCTATACAGT   2823

EMBOSS_001      2851 TGAACTCGGTACAGAAGTAAATGAGTTCGCCTGTGTTGTGGCAGATGCTG   2900
                     ||||||||||||||||||||||||||||||||||||||||||||||||||
EMBOSS_001      2824 TGAACTCGGTACAGAAGTAAATGAGTTCGCCTGTGTTGTGGCAGATGCTG   2873

EMBOSS_001      2901 TCATAAAAACTTTGCAACCAGTATCTGAATTACTTACACCACTGGGCATT   2950
                     ||||||||||||||||||||||||||||||||||||||||||||||||||
EMBOSS_001      2874 TCATAAAAACTTTGCAACCAGTATCTGAATTACTTACACCACTGGGCATT   2923

EMBOSS_001      2951 GATTTAGATGAGTGGAGTATGGCTACATACTACTTATTTGATGAGTCTGG   3000
                     ||||||||||||||||||||||||||||||||||||||||||||||||||
EMBOSS_001      2924 GATTTAGATGAGTGGAGTATGGCTACATACTACTTATTTGATGAGTCTGG   2973

EMBOSS_001      3001 TGAGTTTAAATTGGCTTCACATATGTATTGTTCTTTCTACCCTCCAGATG   3050
                     ||||||||||||||||||||||||||||||||||||||||||||||||||
EMBOSS_001      2974 TGAGTTTAAATTGGCTTCACATATGTATTGTTCTTTCTACCCTCCAGATG   3023

EMBOSS_001      3051 AGGATGAAGAAGAAGGTGATTGTGAAGAAGAAGAGTTTGAGCCATCAACT   3100
                     ||||||||||||||||||||||||||||||||||||||||||||||||||
EMBOSS_001      3024 AGGATGAAGAAGAAGGTGATTGTGAAGAAGAAGAGTTTGAGCCATCAACT   3073

EMBOSS_001      3101 CAATATGAGTATGGTACTGAAGATGATTACCAAGGTAAACCTTTGGAATT   3150
                     ||||||||||||||||||||||||||||||||||||||||||||||||||
EMBOSS_001      3074 CAATATGAGTATGGTACTGAAGATGATTACCAAGGTAAACCTTTGGAATT   3123

EMBOSS_001      3151 TGGTGCCACTTCTGCTGCTCTTCAACCTGAAGAAGAGCAAGAAGAAGATT   3200
                     ||||||||||||||||||||||||||||||||||||||||||||||||||
EMBOSS_001      3124 TGGTGCCACTTCTGCTGCTCTTCAACCTGAAGAAGAGCAAGAAGAAGATT   3173

EMBOSS_001      3201 GGTTAGATGATGATAGTCAACAAACTGTTGGTCAACAAGACGGCAGTGAG   3250
                     ||||||||||||||||||||||||||||||||||||||||||||||||||
EMBOSS_001      3174 GGTTAGATGATGATAGTCAACAAACTGTTGGTCAACAAGACGGCAGTGAG   3223

EMBOSS_001      3251 GACAATCAGACAACTACTATTCAAACAATTGTTGAGGTTCAACCTCAATT   3300
                     ||||||||||||||||||||||||||||||||||||||||||||||||||
EMBOSS_001      3224 GACAATCAGACAACTACTATTCAAACAATTGTTGAGGTTCAACCTCAATT   3273

EMBOSS_001      3301 AGAGATGGAACTTACACCAGTTGTTCAGACTATTGAAGTGAATAGTTTTA   3350
                     ||||||||||||||||||||||||||||||||||||||||||||||||||
EMBOSS_001      3274 AGAGATGGAACTTACACCAGTTGTTCAGACTATTGAAGTGAATAGTTTTA   3323

EMBOSS_001      3351 GTGGTTATTTAAAACTTACTGACAATGTATACATTAAAAATGCAGACATT   3400
                     ||||||||||||||||||||||||||||||||||||||||||||||||||
EMBOSS_001      3324 GTGGTTATTTAAAACTTACTGACAATGTATACATTAAAAATGCAGACATT   3373

EMBOSS_001      3401 GTGGAAGAAGCTAAAAAGGTAAAACCAACAGTGGTTGTTAATGCAGCCAA   3450
                     ||||||||||||||||||||||||||||||||||||||||||||||||||
EMBOSS_001      3374 GTGGAAGAAGCTAAAAAGGTAAAACCAACAGTGGTTGTTAATGCAGCCAA   3423

EMBOSS_001      3451 TGTTTACCTTAAACATGGAGGAGGTGTTGCAGGAGCCTTAAATAAGGCTA   3500
                     ||||||||||||||||||||||||||||||||||||||||||||||||||
EMBOSS_001      3424 TGTTTACCTTAAACATGGAGGAGGTGTTGCAGGAGCCTTAAATAAGGCTA   3473

EMBOSS_001      3501 CTAACAATGCCATGCAAGTTGAATCTGATGATTACATAGCTACTAATGGA   3550
                     ||||||||||||||||||||||||||||||||||||||||||||||||||
EMBOSS_001      3474 CTAACAATGCCATGCAAGTTGAATCTGATGATTACATAGCTACTAATGGA   3523

EMBOSS_001      3551 CCACTTAAAGTGGGTGGTAGTTGTGTTTTAAGCGGACACAATCTTGCTAA   3600
                     ||||||||||||||||||||||||||||||||||||||||||||||||||
EMBOSS_001      3524 CCACTTAAAGTGGGTGGTAGTTGTGTTTTAAGCGGACACAATCTTGCTAA   3573

EMBOSS_001      3601 ACACTGTCTTCATGTTGTCGGCCCAAATGTTAACAAAGGTGAAGACATTC   3650
                     ||||||||||||||||||||||||||||||||||||||||||||||||||
EMBOSS_001      3574 ACACTGTCTTCATGTTGTCGGCCCAAATGTTAACAAAGGTGAAGACATTC   3623

EMBOSS_001      3651 AACTTCTTAAGAGTGCTTATGAAAATTTTAATCAGCACGAAGTTCTACTT   3700
                     ||||||||||||||||||||||||||||||||||||||||||||||||||
EMBOSS_001      3624 AACTTCTTAAGAGTGCTTATGAAAATTTTAATCAGCACGAAGTTCTACTT   3673

EMBOSS_001      3701 GCACCATTATTATCAGCTGGTATTTTTGGTGCTGACCCTATACATTCTTT   3750
                     ||||||||||||||||||||||||||||||||||||||||||||||||||
EMBOSS_001      3674 GCACCATTATTATCAGCTGGTATTTTTGGTGCTGACCCTATACATTCTTT   3723

EMBOSS_001      3751 AAGAGTTTGTGTAGATACTGTTCGCACAAATGTCTACTTAGCTGTCTTTG   3800
                     ||||||||||||||||||||||||||||||||||||||||||||||||||
EMBOSS_001      3724 AAGAGTTTGTGTAGATACTGTTCGCACAAATGTCTACTTAGCTGTCTTTG   3773

EMBOSS_001      3801 ATAAAAATCTCTATGACAAACTTGTTTCAAGCTTTTTGGAAATGAAGAGT   3850
                     ||||||||||||||||||||||||||||||||||||||||||||||||||
EMBOSS_001      3774 ATAAAAATCTCTATGACAAACTTGTTTCAAGCTTTTTGGAAATGAAGAGT   3823

EMBOSS_001      3851 GAAAAGCAAGTTGAACAAAAGATCGCTGAGATTCCTAAAGAGGAAGTTAA   3900
                     ||||||||||||||||||||||||||||||||||||||||||||||||||
EMBOSS_001      3824 GAAAAGCAAGTTGAACAAAAGATCGCTGAGATTCCTAAAGAGGAAGTTAA   3873

EMBOSS_001      3901 GCCATTTATAACTGAAAGTAAACCTTCAGTTGAACAGAGAAAACAAGATG   3950
                     ||||||||||||||||||||||||||||||||||||||||||||||||||
EMBOSS_001      3874 GCCATTTATAACTGAAAGTAAACCTTCAGTTGAACAGAGAAAACAAGATG   3923

EMBOSS_001      3951 ATAAGAAAATCAAAGCTTGTGTTGAAGAAGTTACAACAACTCTGGAAGAA   4000
                     ||||||||||||||||||||||||||||||||||||||||||||||||||
EMBOSS_001      3924 ATAAGAAAATCAAAGCTTGTGTTGAAGAAGTTACAACAACTCTGGAAGAA   3973

EMBOSS_001      4001 ACTAAGTTCCTCACAGAAAACTTGTTACTTTATATTGACATTAATGGCAA   4050
                     ||||||||||||||||||||||||||||||||||||||||||||||||||
EMBOSS_001      3974 ACTAAGTTCCTCACAGAAAACTTGTTACTTTATATTGACATTAATGGCAA   4023

EMBOSS_001      4051 TCTTCATCCAGATTCTGCCACTCTTGTTAGTGACATTGACATCACTTTCT   4100
                     ||||||||||||||||||||||||||||||||||||||||||||||||||
EMBOSS_001      4024 TCTTCATCCAGATTCTGCCACTCTTGTTAGTGACATTGACATCACTTTCT   4073

EMBOSS_001      4101 TAAAGAAAGATGCTCCATATATAGTGGGTGATGTTGTTCAAGAGGGTGTT   4150
                     ||||||||||||||||||||||||||||||||||||||||||||||||||
EMBOSS_001      4074 TAAAGAAAGATGCTCCATATATAGTGGGTGATGTTGTTCAAGAGGGTGTT   4123

EMBOSS_001      4151 TTAACTGCTGTGGTTATACCTACTAAAAAGGCTGGTGGCACTACTGAAAT   4200
                     ||||||||||||||||||||||||||||||||||||||||||||||||||
EMBOSS_001      4124 TTAACTGCTGTGGTTATACCTACTAAAAAGGCTGGTGGCACTACTGAAAT   4173

EMBOSS_001      4201 GCTAGCGAAAGCTTTGAGAAAAGTGCCAACAGACAATTATATAACCACTT   4250
                     ||||||||||||||||||||||||||||||||||||||||||||||||||
EMBOSS_001      4174 GCTAGCGAAAGCTTTGAGAAAAGTGCCAACAGACAATTATATAACCACTT   4223

EMBOSS_001      4251 ACCCGGGTCAGGGTTTAAATGGTTACACTGTAGAGGAGGCAAAGACAGTG   4300
                     ||||||||||||||||||||||||||||||||||||||||||||||||||
EMBOSS_001      4224 ACCCGGGTCAGGGTTTAAATGGTTACACTGTAGAGGAGGCAAAGACAGTG   4273

EMBOSS_001      4301 CTTAAAAAGTGTAAAAGTGCCTTTTACATTCTACCATCTATTATCTCTAA   4350
                     ||||||||||||||||||||||||||||||||||||||||||||||||||
EMBOSS_001      4274 CTTAAAAAGTGTAAAAGTGCCTTTTACATTCTACCATCTATTATCTCTAA   4323

EMBOSS_001      4351 TGAGAAGCAAGAAATTCTTGGAACTGTTTCTTGGAATTTGCGAGAAATGC   4400
                     ||||||||||||||||||||||||||||||||||||||||||||||||||
EMBOSS_001      4324 TGAGAAGCAAGAAATTCTTGGAACTGTTTCTTGGAATTTGCGAGAAATGC   4373

EMBOSS_001      4401 TTGCACATGCAGAAGAAACACGCAAATTAATGCCTGTCTGTGTGGAAACT   4450
                     ||||||||||||||||||||||||||||||||||||||||||||||||||
EMBOSS_001      4374 TTGCACATGCAGAAGAAACACGCAAATTAATGCCTGTCTGTGTGGAAACT   4423

EMBOSS_001      4451 AAAGCCATAGTTTCAACTATACAGCGTAAATATAAGGGTATTAAAATACA   4500
                     ||||||||||||||||||||||||||||||||||||||||||||||||||
EMBOSS_001      4424 AAAGCCATAGTTTCAACTATACAGCGTAAATATAAGGGTATTAAAATACA   4473

EMBOSS_001      4501 AGAGGGTGTGGTTGATTATGGTGCTAGATTTTACTTTTACACCAGTAAAA   4550
                     ||||||||||||||||||||||||||||||||||||||||||||||||||
EMBOSS_001      4474 AGAGGGTGTGGTTGATTATGGTGCTAGATTTTACTTTTACACCAGTAAAA   4523

EMBOSS_001      4551 CAACTGTAGCGTCACTTATCAACACACTTAACGATCTAAATGAAACTCTT   4600
                     ||||||||||||||||||||||||||||||||||||||||||||||||||
EMBOSS_001      4524 CAACTGTAGCGTCACTTATCAACACACTTAACGATCTAAATGAAACTCTT   4573

EMBOSS_001      4601 GTTACAATGCCACTTGGCTATGTAACACATGGCTTAAATTTGGAAGAAGC   4650
                     ||||||||||||||||||||||||||||||||||||||||||||||||||
EMBOSS_001      4574 GTTACAATGCCACTTGGCTATGTAACACATGGCTTAAATTTGGAAGAAGC   4623

EMBOSS_001      4651 TGCTCGGTATATGAGATCTCTCAAAGTGCCAGCTACAGTTTCTGTTTCTT   4700
                     ||||||||||||||||||||||||||||||||||||||||||||||||||
EMBOSS_001      4624 TGCTCGGTATATGAGATCTCTCAAAGTGCCAGCTACAGTTTCTGTTTCTT   4673

EMBOSS_001      4701 CACCTGATGCTGTTACAGCGTATAATGGTTATCTTACTTCTTCTTCTAAA   4750
                     ||||||||||||||||||||||||||||||||||||||||||||||||||
EMBOSS_001      4674 CACCTGATGCTGTTACAGCGTATAATGGTTATCTTACTTCTTCTTCTAAA   4723

EMBOSS_001      4751 ACACCTGAAGAACATTTTATTGAAACCATCTCACTTGCTGGTTCCTATAA   4800
                     ||||||||||||||||||||||||||||||||||||||||||||||||||
EMBOSS_001      4724 ACACCTGAAGAACATTTTATTGAAACCATCTCACTTGCTGGTTCCTATAA   4773

EMBOSS_001      4801 AGATTGGTCCTATTCTGGACAATCTACACAACTAGGTATAGAATTTCTTA   4850
                     ||||||||||||||||||||||||||||||||||||||||||||||||||
EMBOSS_001      4774 AGATTGGTCCTATTCTGGACAATCTACACAACTAGGTATAGAATTTCTTA   4823

EMBOSS_001      4851 AGAGAGGTGATAAAAGTGTATATTACACTAGTAATCCTACCACATTCCAC   4900
                     ||||||||||||||||||||||||||||||||||||||||||||||||||
EMBOSS_001      4824 AGAGAGGTGATAAAAGTGTATATTACACTAGTAATCCTACCACATTCCAC   4873

EMBOSS_001      4901 CTAGATGGTGAAGTTATCACCTTTGACAATCTTAAGACACTTCTTTCTTT   4950
                     ||||||||||||||||||||||||||||||||||||||||||||||||||
EMBOSS_001      4874 CTAGATGGTGAAGTTATCACCTTTGACAATCTTAAGACACTTCTTTCTTT   4923

EMBOSS_001      4951 GAGAGAAGTGAGGACTATTAAGGTGTTTACAACAGTAGACAACATTAACC   5000
                     ||||||||||||||||||||||||||||||||||||||||||||||||||
EMBOSS_001      4924 GAGAGAAGTGAGGACTATTAAGGTGTTTACAACAGTAGACAACATTAACC   4973

EMBOSS_001      5001 TCCACACGCAAGTTGTGGACATGTCAATGACATATGGACAACAGTTTGGT   5050
                     ||||||||||||||||||||||||||||||||||||||||||||||||||
EMBOSS_001      4974 TCCACACGCAAGTTGTGGACATGTCAATGACATATGGACAACAGTTTGGT   5023

EMBOSS_001      5051 CCAACTTATTTGGATGGAGCTGATGTTACTAAAATAAAACCTCATAATTC   5100
                     ||||||||||||||||||||||||||||||||||||||||||||||||||
EMBOSS_001      5024 CCAACTTATTTGGATGGAGCTGATGTTACTAAAATAAAACCTCATAATTC   5073

EMBOSS_001      5101 ACATGAAGGTAAAACATTTTATGTTTTACCTAATGATGACACTCTACGTG   5150
                     ||||||||||||||||||||||||||||||||||||||||||||||||||
EMBOSS_001      5074 ACATGAAGGTAAAACATTTTATGTTTTACCTAATGATGACACTCTACGTG   5123

EMBOSS_001      5151 TTGAGGCTTTTGAGTACTACCACACAACTGATCCTAGTTTTCTGGGTAGG   5200
                     ||||||||||||||||||||||||||||||||||||||||||||||||||
EMBOSS_001      5124 TTGAGGCTTTTGAGTACTACCACACAACTGATCCTAGTTTTCTGGGTAGG   5173

EMBOSS_001      5201 TACATGTCAGCATTAAATCACACTAAAAAGTGGAAATACCCACAAGTTAA   5250
                     ||||||||||||||||||||||||||||||||||||||||||||||||||
EMBOSS_001      5174 TACATGTCAGCATTAAATCACACTAAAAAGTGGAAATACCCACAAGTTAA   5223

EMBOSS_001      5251 TGGTTTAACTTCTATTAAATGGGCAGATAACAACTGTTATCTTGCCACTG   5300
                     ||||||||||||||||||||||||||||||||||||||||||||||||||
EMBOSS_001      5224 TGGTTTAACTTCTATTAAATGGGCAGATAACAACTGTTATCTTGCCACTG   5273

EMBOSS_001      5301 CATTGTTAACACTCCAACAAATAGAGTTGAAGTTTAATCCACCTGCTCTA   5350
                     ||||||||||||||||||||||||||||||||||||||||||||||||||
EMBOSS_001      5274 CATTGTTAACACTCCAACAAATAGAGTTGAAGTTTAATCCACCTGCTCTA   5323

EMBOSS_001      5351 CAAGATGCTTATTACAGAGCAAGGGCTGGTGAAGCTGCTAACTTTTGTGC   5400
                     ||||||||||||||||||||||||||||||||||||||||||||||||||
EMBOSS_001      5324 CAAGATGCTTATTACAGAGCAAGGGCTGGTGAAGCTGCTAACTTTTGTGC   5373

EMBOSS_001      5401 ACTTATCTTAGCCTACTGTAATAAGACAGTAGGTGAGTTAGGTGATGTTA   5450
                     ||||||||||||||||||||||||||||||||||||||||||||||||||
EMBOSS_001      5374 ACTTATCTTAGCCTACTGTAATAAGACAGTAGGTGAGTTAGGTGATGTTA   5423

EMBOSS_001      5451 GAGAAACAATGAGTTACTTGTTTCAACATGCCAATTTAGATTCTTGCAAA   5500
                     ||||||||||||||||||||||||||||||||||||||||||||||||||
EMBOSS_001      5424 GAGAAACAATGAGTTACTTGTTTCAACATGCCAATTTAGATTCTTGCAAA   5473

EMBOSS_001      5501 AGAGTCTTGAACGTGGTGTGTAAAACTTGTGGACAACAGCAGACAACCCT   5550
                     ||||||||||||||||||||||||||||||||||||||||||||||||||
EMBOSS_001      5474 AGAGTCTTGAACGTGGTGTGTAAAACTTGTGGACAACAGCAGACAACCCT   5523

EMBOSS_001      5551 TAAGGGTGTAGAAGCTGTTATGTACATGGGCACACTTTCTTATGAACAAT   5600
                     ||||||||||||||||||||||||||||||||||||||||||||||||||
EMBOSS_001      5524 TAAGGGTGTAGAAGCTGTTATGTACATGGGCACACTTTCTTATGAACAAT   5573

EMBOSS_001      5601 TTAAGAAAGGTGTTCAGATACCTTGTACGTGTGGTAAACAAGCTACAAAA   5650
                     ||||||||||||||||||||||||||||||||||||||||||||||||||
EMBOSS_001      5574 TTAAGAAAGGTGTTCAGATACCTTGTACGTGTGGTAAACAAGCTACAAAA   5623

EMBOSS_001      5651 TATCTAGTACAACAGGAGTCACCTTTTGTTATGATGTCAGCACCACCTGC   5700
                     ||||||||||||||||||||||||||||||||||||||||||||||||||
EMBOSS_001      5624 TATCTAGTACAACAGGAGTCACCTTTTGTTATGATGTCAGCACCACCTGC   5673

EMBOSS_001      5701 TCAGTATGAACTTAAGCATGGTACATTTACTTGTGCTAGTGAGTACACTG   5750
                     ||||||||||||||||||||||||||||||||||||||||||||||||||
EMBOSS_001      5674 TCAGTATGAACTTAAGCATGGTACATTTACTTGTGCTAGTGAGTACACTG   5723

EMBOSS_001      5751 GTAATTACCAGTGTGGTCACTATAAACATATAACTTCTAAAGAAACTTTG   5800
                     ||||||||||||||||||||||||||||||||||||||||||||||||||
EMBOSS_001      5724 GTAATTACCAGTGTGGTCACTATAAACATATAACTTCTAAAGAAACTTTG   5773

EMBOSS_001      5801 TATTGCATAGACGGTGCTTTACTTACAAAGTCCTCAGAATACAAAGGTCC   5850
                     ||||||||||||||||||||||||||||||||||||||||||||||||||
EMBOSS_001      5774 TATTGCATAGACGGTGCTTTACTTACAAAGTCCTCAGAATACAAAGGTCC   5823

EMBOSS_001      5851 TATTACGGATGTTTTCTACAAAGAAAACAGTTACACAACAACCATAAAAC   5900
                     ||||||||||||||||||||||||||||||||||||||||||||||||||
EMBOSS_001      5824 TATTACGGATGTTTTCTACAAAGAAAACAGTTACACAACAACCATAAAAC   5873

EMBOSS_001      5901 CAGTTACTTATAAATTGGATGGTGTTGTTTGTACAGAAATTGACCCTAAG   5950
                     ||||||||||||||||||||||||||||||||||||||||||||||||||
EMBOSS_001      5874 CAGTTACTTATAAATTGGATGGTGTTGTTTGTACAGAAATTGACCCTAAG   5923

EMBOSS_001      5951 TTGGACAATTATTATAAGAAAGACAATTCTTATTTCACAGAGCAACCAAT   6000
                     ||||||||||||||||||||||||||||||||||||||||||||||||||
EMBOSS_001      5924 TTGGACAATTATTATAAGAAAGACAATTCTTATTTCACAGAGCAACCAAT   5973

EMBOSS_001      6001 TGATCTTGTACCAAACCAACCATATCCAAACGCAAGCTTCGATAATTTTA   6050
                     ||||||||||||||||||||||||||||||||||||||||||||||||||
EMBOSS_001      5974 TGATCTTGTACCAAACCAACCATATCCAAACGCAAGCTTCGATAATTTTA   6023

EMBOSS_001      6051 AGTTTGTATGTGATAATATCAAATTTGCTGATGATTTAAACCAGTTAACT   6100
                     ||||||||||||||||||||||||||||||||||||||||||||||||||
EMBOSS_001      6024 AGTTTGTATGTGATAATATCAAATTTGCTGATGATTTAAACCAGTTAACT   6073

EMBOSS_001      6101 GGTTATAAGAAACCTGCTTCAAGAGAGCTTAAAGTTACATTTTTCCCTGA   6150
                     ||||||||||||||||||||||||||||||||||||||||||||||||||
EMBOSS_001      6074 GGTTATAAGAAACCTGCTTCAAGAGAGCTTAAAGTTACATTTTTCCCTGA   6123

EMBOSS_001      6151 CTTAAATGGTGATGTGGTGGCTATTGATTATAAACACTACACACCCTCTT   6200
                     ||||||||||||||||||||||||||||||||||||||||||||||||||
EMBOSS_001      6124 CTTAAATGGTGATGTGGTGGCTATTGATTATAAACACTACACACCCTCTT   6173

EMBOSS_001      6201 TTAAGAAAGGAGCTAAATTGTTACATAAACCTATTGTTTGGCATGTTAAC   6250
                     ||||||||||||||||||||||||||||||||||||||||||||||||||
EMBOSS_001      6174 TTAAGAAAGGAGCTAAATTGTTACATAAACCTATTGTTTGGCATGTTAAC   6223

EMBOSS_001      6251 AATGCAACTAATAAAGCCACGTATAAACCAAATACCTGGTGTATACGTTG   6300
                     ||||||||||||||||||||||||||||||||||||||||||||||||||
EMBOSS_001      6224 AATGCAACTAATAAAGCCACGTATAAACCAAATACCTGGTGTATACGTTG   6273

EMBOSS_001      6301 TCTTTGGAGCACAAAACCAGTTGAAACATCAAATTCGTTTGATGTACTGA   6350
                     ||||||||||||||||||||||||||||||||||||||||||||||||||
EMBOSS_001      6274 TCTTTGGAGCACAAAACCAGTTGAAACATCAAATTCGTTTGATGTACTGA   6323

EMBOSS_001      6351 AGTCAGAGGACGCGCAGGGAATGGATAATCTTGCCTGCGAAGATCTAAAA   6400
                     ||||||||||||||||||||||||||||||||||||||||||||||||||
EMBOSS_001      6324 AGTCAGAGGACGCGCAGGGAATGGATAATCTTGCCTGCGAAGATCTAAAA   6373

EMBOSS_001      6401 CCAGTCTCTGAAGAAGTAGTGGAAAATCCTACCATACAGAAAGACGTTCT   6450
                     ||||||||||||||||||||||||||||||||||||||||||||||||||
EMBOSS_001      6374 CCAGTCTCTGAAGAAGTAGTGGAAAATCCTACCATACAGAAAGACGTTCT   6423

EMBOSS_001      6451 TGAGTGTAATGTGAAAACTACCGAAGTTGTAGGAGACATTATACTTAAAC   6500
                     ||||||||||||||||||||||||||||||||||||||||||||||||||
EMBOSS_001      6424 TGAGTGTAATGTGAAAACTACCGAAGTTGTAGGAGACATTATACTTAAAC   6473

EMBOSS_001      6501 CAGCAAATAATAGTTTAAAAATTACAGAAGAGGTTGGCCACACAGATCTA   6550
                     ||||||||||||||||||||||||||||||||||||||||||||||||||
EMBOSS_001      6474 CAGCAAATAATAGTTTAAAAATTACAGAAGAGGTTGGCCACACAGATCTA   6523

EMBOSS_001      6551 ATGGCTGCTTATGTAGACAATTCTAGTCTTACTATTAAGAAACCTAATGA   6600
                     ||||||||||||||||||||||||||||||||||||||||||||||||||
EMBOSS_001      6524 ATGGCTGCTTATGTAGACAATTCTAGTCTTACTATTAAGAAACCTAATGA   6573

EMBOSS_001      6601 ATTATCTAGAGTATTAGGTTTGAAAACCCTTGCTACTCATGGTTTAGCTG   6650
                     ||||||||||||||||||||||||||||||||||||||||||||||||||
EMBOSS_001      6574 ATTATCTAGAGTATTAGGTTTGAAAACCCTTGCTACTCATGGTTTAGCTG   6623

EMBOSS_001      6651 CTGTTAATAGTGTCCCTTGGGATACTATAGCTAATTATGCTAAGCCTTTT   6700
                     ||||||||||||||||||||||||||||||||||||||||||||||||||
EMBOSS_001      6624 CTGTTAATAGTGTCCCTTGGGATACTATAGCTAATTATGCTAAGCCTTTT   6673

EMBOSS_001      6701 CTTAACAAAGTTGTTAGTACAACTACTAACATAGTTACACGGTGTTTAAA   6750
                     ||||||||||||||||||||||||||||||||||||||||||||||||||
EMBOSS_001      6674 CTTAACAAAGTTGTTAGTACAACTACTAACATAGTTACACGGTGTTTAAA   6723

EMBOSS_001      6751 CCGTGTTTGTACTAATTATATGCCTTATTTCTTTACTTTATTGCTACAAT   6800
                     ||||||||||||||||||||||||||||||||||||||||||||||||||
EMBOSS_001      6724 CCGTGTTTGTACTAATTATATGCCTTATTTCTTTACTTTATTGCTACAAT   6773

EMBOSS_001      6801 TGTGTACTTTTACTAGAAGTACAAATTCTAGAATTAAAGCATCTATGCCG   6850
                     ||||||||||||||||||||||||||||||||||||||||||||||||||
EMBOSS_001      6774 TGTGTACTTTTACTAGAAGTACAAATTCTAGAATTAAAGCATCTATGCCG   6823

EMBOSS_001      6851 ACTACTATAGCAAAGAATACTGTTAAGAGTGTCGGTAAATTTTGTCTAGA   6900
                     ||||||||||||||||||||||||||||||||||||||||||||||||||
EMBOSS_001      6824 ACTACTATAGCAAAGAATACTGTTAAGAGTGTCGGTAAATTTTGTCTAGA   6873

EMBOSS_001      6901 GGCTTCATTTAATTATTTGAAGTCACCTAATTTTTCTAAACTGATAAATA   6950
                     ||||||||||||||||||||||||||||||||||||||||||||||||||
EMBOSS_001      6874 GGCTTCATTTAATTATTTGAAGTCACCTAATTTTTCTAAACTGATAAATA   6923

EMBOSS_001      6951 TTATAATTTGGTTTTTACTATTAAGTGTTTGCCTAGGTTCTTTAATCTAC   7000
                     ||||||||||||||||||||||||||||||||||||||||||||||||||
EMBOSS_001      6924 TTATAATTTGGTTTTTACTATTAAGTGTTTGCCTAGGTTCTTTAATCTAC   6973

EMBOSS_001      7001 TCAACCGCTGCTTTAGGTGTTTTAATGTCTAATTTAGGCATGCCTTCTTA   7050
                     ||||||||||||||||||||||||||||||||||||||||||||||||||
EMBOSS_001      6974 TCAACCGCTGCTTTAGGTGTTTTAATGTCTAATTTAGGCATGCCTTCTTA   7023

EMBOSS_001      7051 CTGTACTGGTTACAGAGAAGGCTATTTGAACTCTACTAATGTCACTATTG   7100
                     ||||||||||||||||||||||||||||||||||||||||||||||||||
EMBOSS_001      7024 CTGTACTGGTTACAGAGAAGGCTATTTGAACTCTACTAATGTCACTATTG   7073

EMBOSS_001      7101 CAACCTACTGTACTGGTTCTATACCTTGTAGTGTTTGTCTTAGTGGTTTA   7150
                     ||||||||||||||||||||||||||||||||||||||||||||||||||
EMBOSS_001      7074 CAACCTACTGTACTGGTTCTATACCTTGTAGTGTTTGTCTTAGTGGTTTA   7123

EMBOSS_001      7151 GATTCTTTAGACACCTATCCTTCTTTAGAAACTATACAAATTACCATTTC   7200
                     ||||||||||||||||||||||||||||||||||||||||||||||||||
EMBOSS_001      7124 GATTCTTTAGACACCTATCCTTCTTTAGAAACTATACAAATTACCATTTC   7173

EMBOSS_001      7201 ATCTTTTAAATGGGATTTAACTGCTTTTGGCTTAGTTGCAGAGTGGTTTT   7250
                     ||||||||||||||||||||||||||||||||||||||||||||||||||
EMBOSS_001      7174 ATCTTTTAAATGGGATTTAACTGCTTTTGGCTTAGTTGCAGAGTGGTTTT   7223

EMBOSS_001      7251 TGGCATATATTCTTTTCACTAGGTTTTTCTATGTACTTGGATTGGCTGCA   7300
                     ||||||||||||||||||||||||||||||||||||||||||||||||||
EMBOSS_001      7224 TGGCATATATTCTTTTCACTAGGTTTTTCTATGTACTTGGATTGGCTGCA   7273

EMBOSS_001      7301 ATCATGCAATTGTTTTTCAGCTATTTTGCAGTACATTTTATTAGTAATTC   7350
                     ||||||||||||||||||||||||||||||||||||||||||||||||||
EMBOSS_001      7274 ATCATGCAATTGTTTTTCAGCTATTTTGCAGTACATTTTATTAGTAATTC   7323

EMBOSS_001      7351 TTGGCTTATGTGGTTAATAATTAATCTTGTACAAATGGCCCCGATTTCAG   7400
                     ||||||||||||||||||||||||||||||||||||||||||||||||||
EMBOSS_001      7324 TTGGCTTATGTGGTTAATAATTAATCTTGTACAAATGGCCCCGATTTCAG   7373

EMBOSS_001      7401 CTATGGTTAGAATGTACATCTTCTTTGCATCATTTTATTATGTATGGAAA   7450
                     ||||||||||||||||||||||||||||||||||||||||||||||||||
EMBOSS_001      7374 CTATGGTTAGAATGTACATCTTCTTTGCATCATTTTATTATGTATGGAAA   7423

EMBOSS_001      7451 AGTTATGTGCATGTTGTAGACGGTTGTAATTCATCAACTTGTATGATGTG   7500
                     ||||||||||||||||||||||||||||||||||||||||||||||||||
EMBOSS_001      7424 AGTTATGTGCATGTTGTAGACGGTTGTAATTCATCAACTTGTATGATGTG   7473

EMBOSS_001      7501 TTACAAACGTAATAGAGCAACAAGAGTCGAATGTACAACTATTGTTAATG   7550
                     ||||||||||||||||||||||||||||||||||||||||||||||||||
EMBOSS_001      7474 TTACAAACGTAATAGAGCAACAAGAGTCGAATGTACAACTATTGTTAATG   7523

EMBOSS_001      7551 GTGTTAGAAGGTCCTTTTATGTCTATGCTAATGGAGGTAAAGGCTTTTGC   7600
                     ||||||||||||||||||||||||||||||||||||||||||||||||||
EMBOSS_001      7524 GTGTTAGAAGGTCCTTTTATGTCTATGCTAATGGAGGTAAAGGCTTTTGC   7573

EMBOSS_001      7601 AAACTACACAATTGGAATTGTGTTAATTGTGATACATTCTGTGCTGGTAG   7650
                     ||||||||||||||||||||||||||||||||||||||||||||||||||
EMBOSS_001      7574 AAACTACACAATTGGAATTGTGTTAATTGTGATACATTCTGTGCTGGTAG   7623

EMBOSS_001      7651 TACATTTATTAGTGATGAAGTTGCGAGAGACTTGTCACTACAGTTTAAAA   7700
                     ||||||||||||||||||||||||||||||||||||||||||||||||||
EMBOSS_001      7624 TACATTTATTAGTGATGAAGTTGCGAGAGACTTGTCACTACAGTTTAAAA   7673

EMBOSS_001      7701 GACCAATAAATCCTACTGACCAGTCTTCTTACATCGTTGATAGTGTTACA   7750
                     ||||||||||||||||||||||||||||||||||||||||||||||||||
EMBOSS_001      7674 GACCAATAAATCCTACTGACCAGTCTTCTTACATCGTTGATAGTGTTACA   7723

EMBOSS_001      7751 GTGAAGAATGGTTCCATCCATCTTTACTTTGATAAAGCTGGTCAAAAGAC   7800
                     ||||||||||||||||||||||||||||||||||||||||||||||||||
EMBOSS_001      7724 GTGAAGAATGGTTCCATCCATCTTTACTTTGATAAAGCTGGTCAAAAGAC   7773

EMBOSS_001      7801 TTATGAAAGACATTCTCTCTCTCATTTTGTTAACTTAGACAACCTGAGAG   7850
                     ||||||||||||||||||||||||||||||||||||||||||||||||||
EMBOSS_001      7774 TTATGAAAGACATTCTCTCTCTCATTTTGTTAACTTAGACAACCTGAGAG   7823

EMBOSS_001      7851 CTAATAACACTAAAGGTTCATTGCCTATTAATGTTATAGTTTTTGATGGT   7900
                     ||||||||||||||||||||||||||||||||||||||||||||||||||
EMBOSS_001      7824 CTAATAACACTAAAGGTTCATTGCCTATTAATGTTATAGTTTTTGATGGT   7873

EMBOSS_001      7901 AAATCAAAATGTGAAGAATCATCTGCAAAATCAGCGTCTGTTTACTACAG   7950
                     ||||||||||||||||||||||||||||||||||||||||||||||||||
EMBOSS_001      7874 AAATCAAAATGTGAAGAATCATCTGCAAAATCAGCGTCTGTTTACTACAG   7923

EMBOSS_001      7951 TCAGCTTATGTGTCAACCTATACTGTTACTAGATCAGGCATTAGTGTCTG   8000
                     ||||||||||||||||||||||||||||||||||||||||||||||||||
EMBOSS_001      7924 TCAGCTTATGTGTCAACCTATACTGTTACTAGATCAGGCATTAGTGTCTG   7973

EMBOSS_001      8001 ATGTTGGTGATAGTGCGGAAGTTGCAGTTAAAATGTTTGATGCTTACGTT   8050
                     ||||||||||||||||||||||||||||||||||||||||||||||||||
EMBOSS_001      7974 ATGTTGGTGATAGTGCGGAAGTTGCAGTTAAAATGTTTGATGCTTACGTT   8023

EMBOSS_001      8051 AATACGTTTTCATCAACTTTTAACGTACCAATGGAAAAACTCAAAACACT   8100
                     ||||||||||||||||||||||||||||||||||||||||||||||||||
EMBOSS_001      8024 AATACGTTTTCATCAACTTTTAACGTACCAATGGAAAAACTCAAAACACT   8073

EMBOSS_001      8101 AGTTGCAACTGCAGAAGCTGAACTTGCAAAGAATGTGTCCTTAGACAATG   8150
                     ||||||||||||||||||||||||||||||||||||||||||||||||||
EMBOSS_001      8074 AGTTGCAACTGCAGAAGCTGAACTTGCAAAGAATGTGTCCTTAGACAATG   8123

EMBOSS_001      8151 TCTTATCTACTTTTATTTCAGCAGCTCGGCAAGGGTTTGTTGATTCAGAT   8200
                     ||||||||||||||||||||||||||||||||||||||||||||||||||
EMBOSS_001      8124 TCTTATCTACTTTTATTTCAGCAGCTCGGCAAGGGTTTGTTGATTCAGAT   8173

EMBOSS_001      8201 GTAGAAACTAAAGATGTTGTTGAATGTCTTAAATTGTCACATCAATCTGA   8250
                     ||||||||||||||||||||||||||||||||||||||||||||||||||
EMBOSS_001      8174 GTAGAAACTAAAGATGTTGTTGAATGTCTTAAATTGTCACATCAATCTGA   8223

EMBOSS_001      8251 CATAGAAGTTACTGGCGATAGTTGTAATAACTATATGCTCACCTATAACA   8300
                     ||||||||||||||||||||||||||||||||||||||||||||||||||
EMBOSS_001      8224 CATAGAAGTTACTGGCGATAGTTGTAATAACTATATGCTCACCTATAACA   8273

EMBOSS_001      8301 AAGTTGAAAACATGACACCCCGTGACCTTGGTGCTTGTATTGACTGTAGT   8350
                     ||||||||||||||||||||||||||||||||||||||||||||||||||
EMBOSS_001      8274 AAGTTGAAAACATGACACCCCGTGACCTTGGTGCTTGTATTGACTGTAGT   8323

EMBOSS_001      8351 GCGCGTCATATTAATGCGCAGGTAGCAAAAAGTCACAACATTGCTTTGAT   8400
                     ||||||||||||||||||||||||||||||||||||||||||||||||||
EMBOSS_001      8324 GCGCGTCATATTAATGCGCAGGTAGCAAAAAGTCACAACATTGCTTTGAT   8373

EMBOSS_001      8401 ATGGAACGTTAAAGATTTCATGTCATTGTCTGAACAACTACGAAAACAAA   8450
                     ||||||||||||||||||||||||||||||||||||||||||||||||||
EMBOSS_001      8374 ATGGAACGTTAAAGATTTCATGTCATTGTCTGAACAACTACGAAAACAAA   8423

EMBOSS_001      8451 TACGTAGTGCTGCTAAAAAGAATAACTTACCTTTTAAGTTGACATGTGCA   8500
                     ||||||||||||||||||||||||||||||||||||||||||||||||||
EMBOSS_001      8424 TACGTAGTGCTGCTAAAAAGAATAACTTACCTTTTAAGTTGACATGTGCA   8473

EMBOSS_001      8501 ACTACTAGACAAGTTGTTAATGTTGTAACAACAAAGATAGCACTTAAGGG   8550
                     ||||||||||||||||||||||||||||||||||||||||||||||||||
EMBOSS_001      8474 ACTACTAGACAAGTTGTTAATGTTGTAACAACAAAGATAGCACTTAAGGG   8523

EMBOSS_001      8551 TGGTAAAATTGTTAATAATTGGTTGAAGCAGTTAATTAAAGTTACACTTG   8600
                     ||||||||||||||||||||||||||||||||||||||||||||||||||
EMBOSS_001      8524 TGGTAAAATTGTTAATAATTGGTTGAAGCAGTTAATTAAAGTTACACTTG   8573

EMBOSS_001      8601 TGTTCCTTTTTGTTGCTGCTATTTTCTATTTAATAACACCTGTTCATGTC   8650
                     ||||||||||||||||||||||||||||||||||||||||||||||||||
EMBOSS_001      8574 TGTTCCTTTTTGTTGCTGCTATTTTCTATTTAATAACACCTGTTCATGTC   8623

EMBOSS_001      8651 ATGTCTAAACATACTGACTTTTCAAGTGAAATCATAGGATACAAGGCTAT   8700
                     ||||||||||||||||||||||||||||||||||||||||||||||||||
EMBOSS_001      8624 ATGTCTAAACATACTGACTTTTCAAGTGAAATCATAGGATACAAGGCTAT   8673

EMBOSS_001      8701 TGATGGTGGTGTCACTCGTGACATAGCATCTACAGATACTTGTTTTGCTA   8750
                     ||||||||||||||||||||||||||||||||||||||||||||||||||
EMBOSS_001      8674 TGATGGTGGTGTCACTCGTGACATAGCATCTACAGATACTTGTTTTGCTA   8723

EMBOSS_001      8751 ACAAACATGCTGATTTTGACACATGGTTTAGCCAGCGTGGTGGTAGTTAT   8800
                     ||||||||||||||||||||||||||||||||||||||||||||||||||
EMBOSS_001      8724 ACAAACATGCTGATTTTGACACATGGTTTAGCCAGCGTGGTGGTAGTTAT   8773

EMBOSS_001      8801 ACTAATGACAAAGCTTGCCCATTGATTGCTGCAGTCATAACAAGAGAAGT   8850
                     ||||||||||||||||||||||||||||||||||||||||||||||||||
EMBOSS_001      8774 ACTAATGACAAAGCTTGCCCATTGATTGCTGCAGTCATAACAAGAGAAGT   8823

EMBOSS_001      8851 GGGTTTTGTCGTGCCTGGTTTGCCTGGCACGATATTACGCACAACTAATG   8900
                     ||||||||||||||||||||||||||||||||||||||||||||||||||
EMBOSS_001      8824 GGGTTTTGTCGTGCCTGGTTTGCCTGGCACGATATTACGCACAACTAATG   8873

EMBOSS_001      8901 GTGACTTTTTGCATTTCTTACCTAGAGTTTTTAGTGCAGTTGGTAACATC   8950
                     ||||||||||||||||||||||||||||||||||||||||||||||||||
EMBOSS_001      8874 GTGACTTTTTGCATTTCTTACCTAGAGTTTTTAGTGCAGTTGGTAACATC   8923

EMBOSS_001      8951 TGTTACACACCATCAAAACTTATAGAGTACACTGACTTTGCAACATCAGC   9000
                     ||||||||||||||||||||||||||||||||||||||||||||||||||
EMBOSS_001      8924 TGTTACACACCATCAAAACTTATAGAGTACACTGACTTTGCAACATCAGC   8973

EMBOSS_001      9001 TTGTGTTTTGGCTGCTGAATGTACAATTTTTAAAGATGCTTCTGGTAAGC   9050
                     ||||||||||||||||||||||||||||||||||||||||||||||||||
EMBOSS_001      8974 TTGTGTTTTGGCTGCTGAATGTACAATTTTTAAAGATGCTTCTGGTAAGC   9023

EMBOSS_001      9051 CAGTACCATATTGTTATGATACCAATGTACTAGAAGGTTCTGTTGCTTAT   9100
                     ||||||||||||||||||||||||||||||||||||||||||||||||||
EMBOSS_001      9024 CAGTACCATATTGTTATGATACCAATGTACTAGAAGGTTCTGTTGCTTAT   9073

EMBOSS_001      9101 GAAAGTTTACGCCCTGACACACGTTATGTGCTCATGGATGGCTCTATTAT   9150
                     ||||||||||||||||||||||||||||||||||||||||||||||||||
EMBOSS_001      9074 GAAAGTTTACGCCCTGACACACGTTATGTGCTCATGGATGGCTCTATTAT   9123

EMBOSS_001      9151 TCAATTTCCTAACACCTACCTTGAAGGTTCTGTTAGAGTGGTAACAACTT   9200
                     ||||||||||||||||||||||||||||||||||||||||||||||||||
EMBOSS_001      9124 TCAATTTCCTAACACCTACCTTGAAGGTTCTGTTAGAGTGGTAACAACTT   9173

EMBOSS_001      9201 TTGATTCTGAGTACTGTAGGCACGGCACTTGTGAAAGATCAGAAGCTGGT   9250
                     ||||||||||||||||||||||||||||||||||||||||||||||||||
EMBOSS_001      9174 TTGATTCTGAGTACTGTAGGCACGGCACTTGTGAAAGATCAGAAGCTGGT   9223

EMBOSS_001      9251 GTTTGTGTATCTACTAGTGGTAGATGGGTACTTAACAATGATTATTACAG   9300
                     ||||||||||||||||||||||||||||||||||||||||||||||||||
EMBOSS_001      9224 GTTTGTGTATCTACTAGTGGTAGATGGGTACTTAACAATGATTATTACAG   9273

EMBOSS_001      9301 ATCTTTACCAGGAGTTTTCTGTGGTGTAGATGCTGTAAATTTACTTACTA   9350
                     ||||||||||||||||||||||||||||||||||||||||||||||||||
EMBOSS_001      9274 ATCTTTACCAGGAGTTTTCTGTGGTGTAGATGCTGTAAATTTACTTACTA   9323

EMBOSS_001      9351 ATATGTTTACACCACTAATTCAACCTATTGGTGCTTTGGACATATCAGCA   9400
                     ||||||||||||||||||||||||||||||||||||||||||||||||||
EMBOSS_001      9324 ATATGTTTACACCACTAATTCAACCTATTGGTGCTTTGGACATATCAGCA   9373

EMBOSS_001      9401 TCTATAGTAGCTGGTGGTATTGTAGCTATCGTAGTAACATGCCTTGCCTA   9450
                     ||||||||||||||||||||||||||||||||||||||||||||||||||
EMBOSS_001      9374 TCTATAGTAGCTGGTGGTATTGTAGCTATCGTAGTAACATGCCTTGCCTA   9423

EMBOSS_001      9451 CTATTTTATGAGGTTTAGAAGAGCTTTTGGTGAATACAGTCATGTAGTTG   9500
                     ||||||||||||||||||||||||||||||||||||||||||||||||||
EMBOSS_001      9424 CTATTTTATGAGGTTTAGAAGAGCTTTTGGTGAATACAGTCATGTAGTTG   9473

EMBOSS_001      9501 CCTTTAATACTTTACTATTCCTTATGTCATTCACTGTACTCTGTTTAACA   9550
                     ||||||||||||||||||||||||||||||||||||||||||||||||||
EMBOSS_001      9474 CCTTTAATACTTTACTATTCCTTATGTCATTCACTGTACTCTGTTTAACA   9523

EMBOSS_001      9551 CCAGTTTACTCATTCTTACCTGGTGTTTATTCTGTTATTTACTTGTACTT   9600
                     ||||||||||||||||||||||||||||||||||||||||||||||||||
EMBOSS_001      9524 CCAGTTTACTCATTCTTACCTGGTGTTTATTCTGTTATTTACTTGTACTT   9573

EMBOSS_001      9601 GACATTTTATCTTACTAATGATGTTTCTTTTTTAGCACATATTCAGTGGA   9650
                     ||||||||||||||||||||||||||||||||||||||||||||||||||
EMBOSS_001      9574 GACATTTTATCTTACTAATGATGTTTCTTTTTTAGCACATATTCAGTGGA   9623

EMBOSS_001      9651 TGGTTATGTTCACACCTTTAGTACCTTTCTGGATAACAATTGCTTATATC   9700
                     ||||||||||||||||||||||||||||||||||||||||||||||||||
EMBOSS_001      9624 TGGTTATGTTCACACCTTTAGTACCTTTCTGGATAACAATTGCTTATATC   9673

EMBOSS_001      9701 ATTTGTATTTCCACAAAGCATTTCTATTGGTTCTTTAGTAATTACCTAAA   9750
                     ||||||||||||||||||||||||||||||||||||||||||||||||||
EMBOSS_001      9674 ATTTGTATTTCCACAAAGCATTTCTATTGGTTCTTTAGTAATTACCTAAA   9723

EMBOSS_001      9751 GAGACGTGTAGTCTTTAATGGTGTTTCCTTTAGTACTTTTGAAGAAGCTG   9800
                     ||||||||||||||||||||||||||||||||||||||||||||||||||
EMBOSS_001      9724 GAGACGTGTAGTCTTTAATGGTGTTTCCTTTAGTACTTTTGAAGAAGCTG   9773

EMBOSS_001      9801 CGCTGTGCACCTTTTTGTTAAATAAAGAAATGTATCTAAAGTTGCGTAGT   9850
                     ||||||||||||||||||||||||||||||||||||||||||||||||||
EMBOSS_001      9774 CGCTGTGCACCTTTTTGTTAAATAAAGAAATGTATCTAAAGTTGCGTAGT   9823

EMBOSS_001      9851 GATGTGCTATTACCTCTTACGCAATATAATAGATACTTAGCTCTTTATAA   9900
                     ||||||||||||||||||||||||||||||||||||||||||||||||||
EMBOSS_001      9824 GATGTGCTATTACCTCTTACGCAATATAATAGATACTTAGCTCTTTATAA   9873

EMBOSS_001      9901 TAAGTACAAGTATTTTAGTGGAGCAATGGATACAACTAGCTACAGAGAAG   9950
                     ||||||||||||||||||||||||||||||||||||||||||||||||||
EMBOSS_001      9874 TAAGTACAAGTATTTTAGTGGAGCAATGGATACAACTAGCTACAGAGAAG   9923

EMBOSS_001      9951 CTGCTTGTTGTCATCTCGCAAAGGCTCTCAATGACTTCAGTAACTCAGGT  10000
                     ||||||||||||||||||||||||||||||||||||||||||||||||||
EMBOSS_001      9924 CTGCTTGTTGTCATCTCGCAAAGGCTCTCAATGACTTCAGTAACTCAGGT   9973

EMBOSS_001     10001 TCTGATGTTCTTTACCAACCACCACAAACCTCTATCACCTCAGCTGTTTT  10050
                     ||||||||||||||||||||||||||||||||||||||||||||||||||
EMBOSS_001      9974 TCTGATGTTCTTTACCAACCACCACAAACCTCTATCACCTCAGCTGTTTT  10023

EMBOSS_001     10051 GCAGAGTGGTTTTAGAAAAATGGCATTCCCATCTGGTAAAGTTGAGGGTT  10100
                     ||||||||||||||||||||||||||||||||||||||||||||||||||
EMBOSS_001     10024 GCAGAGTGGTTTTAGAAAAATGGCATTCCCATCTGGTAAAGTTGAGGGTT  10073

EMBOSS_001     10101 GTATGGTACAAGTAACTTGTGGTACAACTACACTTAACGGTCTTTGGCTT  10150
                     ||||||||||||||||||||||||||||||||||||||||||||||||||
EMBOSS_001     10074 GTATGGTACAAGTAACTTGTGGTACAACTACACTTAACGGTCTTTGGCTT  10123

EMBOSS_001     10151 GATGACGTAGTTTACTGTCCAAGACATGTGATCTGCACCTCTGAAGACAT  10200
                     ||||||||||||||||||||||||||||||||||||||||||||||||||
EMBOSS_001     10124 GATGACGTAGTTTACTGTCCAAGACATGTGATCTGCACCTCTGAAGACAT  10173

EMBOSS_001     10201 GCTTAACCCTAATTATGAAGATTTACTCATTCGTAAGTCTAATCATAATT  10250
                     ||||||||||||||||||||||||||||||||||||||||||||||||||
EMBOSS_001     10174 GCTTAACCCTAATTATGAAGATTTACTCATTCGTAAGTCTAATCATAATT  10223

EMBOSS_001     10251 TCTTGGTACAGGCTGGTAATGTTCAACTCAGGGTTATTGGACATTCTATG  10300
                     ||||||||||||||||||||||||||||||||||||||||||||||||||
EMBOSS_001     10224 TCTTGGTACAGGCTGGTAATGTTCAACTCAGGGTTATTGGACATTCTATG  10273

EMBOSS_001     10301 CAAAATTGTGTACTTAAGCTTAAGGTTGATACAGCCAATCCTAAGACACC  10350
                     ||||||||||||||||||||||||||||||||||||||||||||||||||
EMBOSS_001     10274 CAAAATTGTGTACTTAAGCTTAAGGTTGATACAGCCAATCCTAAGACACC  10323

EMBOSS_001     10351 TAAGTATAAGTTTGTTCGCATTCAACCAGGACAGACTTTTTCAGTGTTAG  10400
                     ||||||||||||||||||||||||||||||||||||||||||||||||||
EMBOSS_001     10324 TAAGTATAAGTTTGTTCGCATTCAACCAGGACAGACTTTTTCAGTGTTAG  10373

EMBOSS_001     10401 CTTGTTACAATGGTTCACCATCTGGTGTTTACCAATGTGCTATGAGGCCC  10450
                     ||||||||||||||||||||||||||||||||||||||||||||||||||
EMBOSS_001     10374 CTTGTTACAATGGTTCACCATCTGGTGTTTACCAATGTGCTATGAGGCCC  10423

EMBOSS_001     10451 AATTTCACTATTAAGGGTTCATTCCTTAATGGTTCATGTGGTAGTGTTGG  10500
                     ||||||||||||||||||||||||||||||||||||||||||||||||||
EMBOSS_001     10424 AATTTCACTATTAAGGGTTCATTCCTTAATGGTTCATGTGGTAGTGTTGG  10473

EMBOSS_001     10501 TTTTAACATAGATTATGACTGTGTCTCTTTTTGTTACATGCACCATATGG  10550
                     ||||||||||||||||||||||||||||||||||||||||||||||||||
EMBOSS_001     10474 TTTTAACATAGATTATGACTGTGTCTCTTTTTGTTACATGCACCATATGG  10523

EMBOSS_001     10551 AATTACCAACTGGAGTTCATGCTGGCACAGACTTAGAAGGTAACTTTTAT  10600
                     ||||||||||||||||||||||||||||||||||||||||||||||||||
EMBOSS_001     10524 AATTACCAACTGGAGTTCATGCTGGCACAGACTTAGAAGGTAACTTTTAT  10573

EMBOSS_001     10601 GGACCTTTTGTTGACAGGCAAACAGCACAAGCAGCTGGTACGGACACAAC  10650
                     ||||||||||||||||||||||||||||||||||||||||||||||||||
EMBOSS_001     10574 GGACCTTTTGTTGACAGGCAAACAGCACAAGCAGCTGGTACGGACACAAC  10623

EMBOSS_001     10651 TATTACAGTTAATGTTTTAGCTTGGTTGTACGCTGCTGTTATAAATGGAG  10700
                     ||||||||||||||||||||||||||||||||||||||||||||||||||
EMBOSS_001     10624 TATTACAGTTAATGTTTTAGCTTGGTTGTACGCTGCTGTTATAAATGGAG  10673

EMBOSS_001     10701 ACAGGTGGTTTCTCAATCGATTTACCACAACTCTTAATGACTTTAACCTT  10750
                     ||||||||||||||||||||||||||||||||||||||||||||||||||
EMBOSS_001     10674 ACAGGTGGTTTCTCAATCGATTTACCACAACTCTTAATGACTTTAACCTT  10723

EMBOSS_001     10751 GTGGCTATGAAGTACAATTATGAACCTCTAACACAAGACCATGTTGACAT  10800
                     ||||||||||||||||||||||||||||||||||||||||||||||||||
EMBOSS_001     10724 GTGGCTATGAAGTACAATTATGAACCTCTAACACAAGACCATGTTGACAT  10773

EMBOSS_001     10801 ACTAGGACCTCTTTCTGCTCAAACTGGAATTGCCGTTTTAGATATGTGTG  10850
                     ||||||||||||||||||||||||||||||||||||||||||||||||||
EMBOSS_001     10774 ACTAGGACCTCTTTCTGCTCAAACTGGAATTGCCGTTTTAGATATGTGTG  10823

EMBOSS_001     10851 CTTCATTAAAAGAATTACTGCAAAATGGTATGAATGGACGTACCATATTG  10900
                     ||||||||||||||||||||||||||||||||||||||||||||||||||
EMBOSS_001     10824 CTTCATTAAAAGAATTACTGCAAAATGGTATGAATGGACGTACCATATTG  10873

EMBOSS_001     10901 GGTAGTGCTTTATTAGAAGATGAATTTACACCTTTTGATGTTGTTAGACA  10950
                     ||||||||||||||||||||||||||||||||||||||||||||||||||
EMBOSS_001     10874 GGTAGTGCTTTATTAGAAGATGAATTTACACCTTTTGATGTTGTTAGACA  10923

EMBOSS_001     10951 ATGCTCAGGTGTTACTTTCCAAAGTGCAGTGAAAAGAACAATCAAGGGTA  11000
                     ||||||||||||||||||||||||||||||||||||||||||||||||||
EMBOSS_001     10924 ATGCTCAGGTGTTACTTTCCAAAGTGCAGTGAAAAGAACAATCAAGGGTA  10973

EMBOSS_001     11001 CACACCACTGGTTGTTACTCACAATTTTGACTTCACTTTTAGTTTTAGTC  11050
                     ||||||||||||||||||||||||||||||||||||||||||||||||||
EMBOSS_001     10974 CACACCACTGGTTGTTACTCACAATTTTGACTTCACTTTTAGTTTTAGTC  11023

EMBOSS_001     11051 CAGAGTACTCAATGGTCTTTGTTCTTTTTTTTGTATGAAAATGCCTTTTT  11100
                     ||||||||||||||||||||||||||||||||||||||||||||||||||
EMBOSS_001     11024 CAGAGTACTCAATGGTCTTTGTTCTTTTTTTTGTATGAAAATGCCTTTTT  11073

EMBOSS_001     11101 ACCTTTTGCTATGGGTATTATTGCTATGTCTGCTTTTGCAATGATGTTTG  11150
                     ||||||||||||||||||||||||||||||||||||||||||||||||||
EMBOSS_001     11074 ACCTTTTGCTATGGGTATTATTGCTATGTCTGCTTTTGCAATGATGTTTG  11123

EMBOSS_001     11151 TCAAACATAAGCATGCATTTCTCTGTTTGTTTTTGTTACCTTCTCTTGCC  11200
                     ||||||||||||||||||||||||||||||||||||||||||||||||||
EMBOSS_001     11124 TCAAACATAAGCATGCATTTCTCTGTTTGTTTTTGTTACCTTCTCTTGCC  11173

EMBOSS_001     11201 ACTGTAGCTTATTTTAATATGGTCTATATGCCTGCTAGTTGGGTGATGCG  11250
                     ||||||||||||||||||||||||||||||||||||||||||||||||||
EMBOSS_001     11174 ACTGTAGCTTATTTTAATATGGTCTATATGCCTGCTAGTTGGGTGATGCG  11223

EMBOSS_001     11251 TATTATGACATGGTTGGATATGGTTGATACTAGTTTGTCTGGTTTTAAGC  11300
                     ||||||||||||||||||||||||||||||||||||||||||||||||||
EMBOSS_001     11224 TATTATGACATGGTTGGATATGGTTGATACTAGTTTGTCTGGTTTTAAGC  11273

EMBOSS_001     11301 TAAAAGACTGTGTTATGTATGCATCAGCTGTAGTGTTACTAATCCTTATG  11350
                     ||||||||||||||||||||||||||||||||||||||||||||||||||
EMBOSS_001     11274 TAAAAGACTGTGTTATGTATGCATCAGCTGTAGTGTTACTAATCCTTATG  11323

EMBOSS_001     11351 ACAGCAAGAACTGTGTATGATGATGGTGCTAGGAGAGTGTGGACACTTAT  11400
                     ||||||||||||||||||||||||||||||||||||||||||||||||||
EMBOSS_001     11324 ACAGCAAGAACTGTGTATGATGATGGTGCTAGGAGAGTGTGGACACTTAT  11373

EMBOSS_001     11401 GAATGTCTTGACACTCGTTTATAAAGTTTATTATGGTAATGCTTTAGATC  11450
                     ||||||||||||||||||||||||||||||||||||||||||||||||||
EMBOSS_001     11374 GAATGTCTTGACACTCGTTTATAAAGTTTATTATGGTAATGCTTTAGATC  11423

EMBOSS_001     11451 AAGCCATTTCCATGTGGGCTCTTATAATCTCTGTTACTTCTAACTACTCA  11500
                     ||||||||||||||||||||||||||||||||||||||||||||||||||
EMBOSS_001     11424 AAGCCATTTCCATGTGGGCTCTTATAATCTCTGTTACTTCTAACTACTCA  11473

EMBOSS_001     11501 GGTGTAGTTACAACTGTCATGTTTTTGGCCAGAGGTATTGTTTTTATGTG  11550
                     ||||||||||||||||||||||||||||||||||||||||||||||||||
EMBOSS_001     11474 GGTGTAGTTACAACTGTCATGTTTTTGGCCAGAGGTATTGTTTTTATGTG  11523

EMBOSS_001     11551 TGTTGAGTATTGCCCTATTTTCTTCATAACTGGTAATACACTTCAGTGTA  11600
                     ||||||||||||||||||||||||||||||||||||||||||||||||||
EMBOSS_001     11524 TGTTGAGTATTGCCCTATTTTCTTCATAACTGGTAATACACTTCAGTGTA  11573

EMBOSS_001     11601 TAATGCTAGTTTATTGTTTCTTAGGCTATTTTTGTACTTGTTACTTTGGC  11650
                     ||||||||||||||||||||||||||||||||||||||||||||||||||
EMBOSS_001     11574 TAATGCTAGTTTATTGTTTCTTAGGCTATTTTTGTACTTGTTACTTTGGC  11623

EMBOSS_001     11651 CTCTTTTGTTTACTCAACCGCTACTTTAGACTGACTCTTGGTGTTTATGA  11700
                     ||||||||||||||||||||||||||||||||||||||||||||||||||
EMBOSS_001     11624 CTCTTTTGTTTACTCAACCGCTACTTTAGACTGACTCTTGGTGTTTATGA  11673

EMBOSS_001     11701 TTACTTAGTTTCTACACAGGAGTTTAGATATATGAATTCACAGGGACTAC  11750
                     ||||||||||||||||||||||||||||||||||||||||||||||||||
EMBOSS_001     11674 TTACTTAGTTTCTACACAGGAGTTTAGATATATGAATTCACAGGGACTAC  11723

EMBOSS_001     11751 TCCCACCCAAGAATAGCATAGATGCCTTCAAACTCAACATTAAATTGTTG  11800
                     ||||||||||||||||||||||||||||||||||||||||||||||||||
EMBOSS_001     11724 TCCCACCCAAGAATAGCATAGATGCCTTCAAACTCAACATTAAATTGTTG  11773

EMBOSS_001     11801 GGTGTTGGTGGCAAACCTTGTATCAAAGTAGCCACTGTACAGTCTAAAAT  11850
                     ||||||||||||||||||||||||||||||||||||||||||||||||||
EMBOSS_001     11774 GGTGTTGGTGGCAAACCTTGTATCAAAGTAGCCACTGTACAGTCTAAAAT  11823

EMBOSS_001     11851 GTCAGATGTAAAGTGCACATCAGTAGTCTTACTCTCAGTTTTGCAACAAC  11900
                     ||||||||||||||||||||||||||||||||||||||||||||||||||
EMBOSS_001     11824 GTCAGATGTAAAGTGCACATCAGTAGTCTTACTCTCAGTTTTGCAACAAC  11873

EMBOSS_001     11901 TCAGAGTAGAATCATCATCTAAATTGTGGGCTCAATGTGTCCAGTTACAC  11950
                     ||||||||||||||||||||||||||||||||||||||||||||||||||
EMBOSS_001     11874 TCAGAGTAGAATCATCATCTAAATTGTGGGCTCAATGTGTCCAGTTACAC  11923

EMBOSS_001     11951 AATGACATTCTCTTAGCTAAAGATACTACTGAAGCCTTTGAAAAAATGGT  12000
                     ||||||||||||||||||||||||||||||||||||||||||||||||||
EMBOSS_001     11924 AATGACATTCTCTTAGCTAAAGATACTACTGAAGCCTTTGAAAAAATGGT  11973

EMBOSS_001     12001 TTCACTACTTTCTGTTTTGCTTTCCATGCAGGGTGCTGTAGACATAAACA  12050
                     ||||||||||||||||||||||||||||||||||||||||||||||||||
EMBOSS_001     11974 TTCACTACTTTCTGTTTTGCTTTCCATGCAGGGTGCTGTAGACATAAACA  12023

EMBOSS_001     12051 AGCTTTGTGAAGAAATGCTGGACAACAGGGCAACCTTACAAGCTATAGCC  12100
                     ||||||||||||||||||||||||||||||||||||||||||||||||||
EMBOSS_001     12024 AGCTTTGTGAAGAAATGCTGGACAACAGGGCAACCTTACAAGCTATAGCC  12073

EMBOSS_001     12101 TCAGAGTTTAGTTCCCTTCCATCATATGCAGCTTTTGCTACTGCTCAAGA  12150
                     ||||||||||||||||||||||||||||||||||||||||||||||||||
EMBOSS_001     12074 TCAGAGTTTAGTTCCCTTCCATCATATGCAGCTTTTGCTACTGCTCAAGA  12123

EMBOSS_001     12151 AGCTTATGAGCAGGCTGTTGCTAATGGTGATTCTGAAGTTGTTCTTAAAA  12200
                     ||||||||||||||||||||||||||||||||||||||||||||||||||
EMBOSS_001     12124 AGCTTATGAGCAGGCTGTTGCTAATGGTGATTCTGAAGTTGTTCTTAAAA  12173

EMBOSS_001     12201 AGTTGAAGAAGTCTTTGAATGTGGCTAAATCTGAATTTGACCGTGATGCA  12250
                     ||||||||||||||||||||||||||||||||||||||||||||||||||
EMBOSS_001     12174 AGTTGAAGAAGTCTTTGAATGTGGCTAAATCTGAATTTGACCGTGATGCA  12223

EMBOSS_001     12251 GCCATGCAACGTAAGTTGGAAAAGATGGCTGATCAAGCTATGACCCAAAT  12300
                     ||||||||||||||||||||||||||||||||||||||||||||||||||
EMBOSS_001     12224 GCCATGCAACGTAAGTTGGAAAAGATGGCTGATCAAGCTATGACCCAAAT  12273

EMBOSS_001     12301 GTATAAACAGGCTAGATCTGAGGACAAGAGGGCAAAAGTTACTAGTGCTA  12350
                     ||||||||||||||||||||||||||||||||||||||||||||||||||
EMBOSS_001     12274 GTATAAACAGGCTAGATCTGAGGACAAGAGGGCAAAAGTTACTAGTGCTA  12323

EMBOSS_001     12351 TGCAGACAATGCTTTTCACTATGCTTAGAAAGTTGGATAATGATGCACTC  12400
                     ||||||||||||||||||||||||||||||||||||||||||||||||||
EMBOSS_001     12324 TGCAGACAATGCTTTTCACTATGCTTAGAAAGTTGGATAATGATGCACTC  12373

EMBOSS_001     12401 AACAACATTATCAACAATGCAAGAGATGGTTGTGTTCCCTTGAACATAAT  12450
                     ||||||||||||||||||||||||||||||||||||||||||||||||||
EMBOSS_001     12374 AACAACATTATCAACAATGCAAGAGATGGTTGTGTTCCCTTGAACATAAT  12423

EMBOSS_001     12451 ACCTCTTACAACAGCAGCCAAACTAATGGTTGTCATACCAGACTATAACA  12500
                     ||||||||||||||||||||||||||||||||||||||||||||||||||
EMBOSS_001     12424 ACCTCTTACAACAGCAGCCAAACTAATGGTTGTCATACCAGACTATAACA  12473

EMBOSS_001     12501 CATATAAAAATACGTGTGATGGTACAACATTTACTTATGCATCAGCATTG  12550
                     ||||||||||||||||||||||||||||||||||||||||||||||||||
EMBOSS_001     12474 CATATAAAAATACGTGTGATGGTACAACATTTACTTATGCATCAGCATTG  12523

EMBOSS_001     12551 TGGGAAATCCAACAGGTTGTAGATGCAGATAGTAAAATTGTTCAACTTAG  12600
                     ||||||||||||||||||||||||||||||||||||||||||||||||||
EMBOSS_001     12524 TGGGAAATCCAACAGGTTGTAGATGCAGATAGTAAAATTGTTCAACTTAG  12573

EMBOSS_001     12601 TGAAATTAGTATGGACAATTCACCTAATTTAGCATGGCCTCTTATTGTAA  12650
                     ||||||||||||||||||||||||||||||||||||||||||||||||||
EMBOSS_001     12574 TGAAATTAGTATGGACAATTCACCTAATTTAGCATGGCCTCTTATTGTAA  12623

EMBOSS_001     12651 CAGCTTTAAGGGCCAATTCTGCTGTCAAATTACAGAATAATGAGCTTAGT  12700
                     ||||||||||||||||||||||||||||||||||||||||||||||||||
EMBOSS_001     12624 CAGCTTTAAGGGCCAATTCTGCTGTCAAATTACAGAATAATGAGCTTAGT  12673

EMBOSS_001     12701 CCTGTTGCACTACGACAGATGTCTTGTGCTGCCGGTACTACACAAACTGC  12750
                     ||||||||||||||||||||||||||||||||||||||||||||||||||
EMBOSS_001     12674 CCTGTTGCACTACGACAGATGTCTTGTGCTGCCGGTACTACACAAACTGC  12723

EMBOSS_001     12751 TTGCACTGATGACAATGCGTTAGCTTACTACAACACAACAAAGGGAGGTA  12800
                     ||||||||||||||||||||||||||||||||||||||||||||||||||
EMBOSS_001     12724 TTGCACTGATGACAATGCGTTAGCTTACTACAACACAACAAAGGGAGGTA  12773

EMBOSS_001     12801 GGTTTGTACTTGCACTGTTATCCGATTTACAGGATTTGAAATGGGCTAGA  12850
                     ||||||||||||||||||||||||||||||||||||||||||||||||||
EMBOSS_001     12774 GGTTTGTACTTGCACTGTTATCCGATTTACAGGATTTGAAATGGGCTAGA  12823

EMBOSS_001     12851 TTCCCTAAGAGTGATGGAACTGGTACTATCTATACAGAACTGGAACCACC  12900
                     ||||||||||||||||||||||||||||||||||||||||||||||||||
EMBOSS_001     12824 TTCCCTAAGAGTGATGGAACTGGTACTATCTATACAGAACTGGAACCACC  12873

EMBOSS_001     12901 TTGTAGGTTTGTTACAGACACACCTAAAGGTCCTAAAGTGAAGTATTTAT  12950
                     ||||||||||||||||||||||||||||||||||||||||||||||||||
EMBOSS_001     12874 TTGTAGGTTTGTTACAGACACACCTAAAGGTCCTAAAGTGAAGTATTTAT  12923

EMBOSS_001     12951 ACTTTATTAAAGGATTAAACAACCTAAATAGAGGTATGGTACTTGGTAGT  13000
                     ||||||||||||||||||||||||||||||||||||||||||||||||||
EMBOSS_001     12924 ACTTTATTAAAGGATTAAACAACCTAAATAGAGGTATGGTACTTGGTAGT  12973

EMBOSS_001     13001 TTAGCTGCCACAGTACGTCTACAAGCTGGTAATGCAACAGAAGTGCCTGC  13050
                     ||||||||||||||||||||||||||||||||||||||||||||||||||
EMBOSS_001     12974 TTAGCTGCCACAGTACGTCTACAAGCTGGTAATGCAACAGAAGTGCCTGC  13023

EMBOSS_001     13051 CAATTCAACTGTATTATCTTTCTGTGCTTTTGCTGTAGATGCTGCTAAAG  13100
                     ||||||||||||||||||||||||||||||||||||||||||||||||||
EMBOSS_001     13024 CAATTCAACTGTATTATCTTTCTGTGCTTTTGCTGTAGATGCTGCTAAAG  13073

EMBOSS_001     13101 CTTACAAAGATTATCTAGCTAGTGGGGGACAACCAATCACTAATTGTGTT  13150
                     ||||||||||||||||||||||||||||||||||||||||||||||||||
EMBOSS_001     13074 CTTACAAAGATTATCTAGCTAGTGGGGGACAACCAATCACTAATTGTGTT  13123

EMBOSS_001     13151 AAGATGTTGTGTACACACACTGGTACTGGTCAGGCAATAACAGTTACACC  13200
                     ||||||||||||||||||||||||||||||||||||||||||||||||||
EMBOSS_001     13124 AAGATGTTGTGTACACACACTGGTACTGGTCAGGCAATAACAGTTACACC  13173

EMBOSS_001     13201 GGAAGCCAATATGGATCAAGAATCCTTTGGTGGTGCATCGTGTTGTCTGT  13250
                     ||||||||||||||||||||||||||||||||||||||||||||||||||
EMBOSS_001     13174 GGAAGCCAATATGGATCAAGAATCCTTTGGTGGTGCATCGTGTTGTCTGT  13223

EMBOSS_001     13251 ACTGCCGTTGCCACATAGATCATCCAAATCCTAAAGGATTTTGTGACTTA  13300
                     ||||||||||||||||||||||||||||||||||||||||||||||||||
EMBOSS_001     13224 ACTGCCGTTGCCACATAGATCATCCAAATCCTAAAGGATTTTGTGACTTA  13273

EMBOSS_001     13301 AAAGGTAAGTATGTACAAATACCTACAACTTGTGCTAATGACCCTGTGGG  13350
                     ||||||||||||||||||||||||||||||||||||||||||||||||||
EMBOSS_001     13274 AAAGGTAAGTATGTACAAATACCTACAACTTGTGCTAATGACCCTGTGGG  13323

EMBOSS_001     13351 TTTTACACTTAAAAACACAGTCTGTACCGTCTGCGGTATGTGGAAAGGTT  13400
                     ||||||||||||||||||||||||||||||||||||||||||||||||||
EMBOSS_001     13324 TTTTACACTTAAAAACACAGTCTGTACCGTCTGCGGTATGTGGAAAGGTT  13373

EMBOSS_001     13401 ATGGCTGTAGTTGTGATCAACTCCGCGAACCCATGCTTCAGTCAGCTGAT  13450
                     ||||||||||||||||||||||||||||||||||||||||||||||||||
EMBOSS_001     13374 ATGGCTGTAGTTGTGATCAACTCCGCGAACCCATGCTTCAGTCAGCTGAT  13423

EMBOSS_001     13451 GCACAATCGTTTTTAAACGGGTTTGCGGTGTAAGTGCAGCCCGTCTTACA  13500
                     ||||||||||||||||||||||||||||||||||||||||||||||||||
EMBOSS_001     13424 GCACAATCGTTTTTAAACGGGTTTGCGGTGTAAGTGCAGCCCGTCTTACA  13473

EMBOSS_001     13501 CCGTGCGGCACAGGCACTAGTACTGATGTCGTATACAGGGCTTTTGACAT  13550
                     ||||||||||||||||||||||||||||||||||||||||||||||||||
EMBOSS_001     13474 CCGTGCGGCACAGGCACTAGTACTGATGTCGTATACAGGGCTTTTGACAT  13523

EMBOSS_001     13551 CTACAATGATAAAGTAGCTGGTTTTGCTAAATTCCTAAAAACTAATTGTT  13600
                     ||||||||||||||||||||||||||||||||||||||||||||||||||
EMBOSS_001     13524 CTACAATGATAAAGTAGCTGGTTTTGCTAAATTCCTAAAAACTAATTGTT  13573

EMBOSS_001     13601 GTCGCTTCCAAGAAAAGGACGAAGATGACAATTTAATTGATTCTTACTTT  13650
                     ||||||||||||||||||||||||||||||||||||||||||||||||||
EMBOSS_001     13574 GTCGCTTCCAAGAAAAGGACGAAGATGACAATTTAATTGATTCTTACTTT  13623

EMBOSS_001     13651 GTAGTTAAGAGACACACTTTCTCTAACTACCAACATGAAGAAACAATTTA  13700
                     ||||||||||||||||||||||||||||||||||||||||||||||||||
EMBOSS_001     13624 GTAGTTAAGAGACACACTTTCTCTAACTACCAACATGAAGAAACAATTTA  13673

EMBOSS_001     13701 TAATTTACTTAAGGATTGTCCAGCTGTTGCTAAACATGACTTCTTTAAGT  13750
                     ||||||||||||||||||||||||||||||||||||||||||||||||||
EMBOSS_001     13674 TAATTTACTTAAGGATTGTCCAGCTGTTGCTAAACATGACTTCTTTAAGT  13723

EMBOSS_001     13751 TTAGAATAGACGGTGACATGGTACCACATATATCACGTCAACGTCTTACT  13800
                     ||||||||||||||||||||||||||||||||||||||||||||||||||
EMBOSS_001     13724 TTAGAATAGACGGTGACATGGTACCACATATATCACGTCAACGTCTTACT  13773

EMBOSS_001     13801 AAATACACAATGGCAGACCTCGTCTATGCTTTAAGGCATTTTGATGAAGG  13850
                     ||||||||||||||||||||||||||||||||||||||||||||||||||
EMBOSS_001     13774 AAATACACAATGGCAGACCTCGTCTATGCTTTAAGGCATTTTGATGAAGG  13823

EMBOSS_001     13851 TAATTGTGACACATTAAAAGAAATACTTGTCACATACAATTGTTGTGATG  13900
                     ||||||||||||||||||||||||||||||||||||||||||||||||||
EMBOSS_001     13824 TAATTGTGACACATTAAAAGAAATACTTGTCACATACAATTGTTGTGATG  13873

EMBOSS_001     13901 ATGATTATTTCAATAAAAAGGACTGGTATGATTTTGTAGAAAACCCAGAT  13950
                     ||||||||||||||||||||||||||||||||||||||||||||||||||
EMBOSS_001     13874 ATGATTATTTCAATAAAAAGGACTGGTATGATTTTGTAGAAAACCCAGAT  13923

EMBOSS_001     13951 ATATTACGCGTATACGCCAACTTAGGTGAACGTGTACGCCAAGCTTTGTT  14000
                     ||||||||||||||||||||||||||||||||||||||||||||||||||
EMBOSS_001     13924 ATATTACGCGTATACGCCAACTTAGGTGAACGTGTACGCCAAGCTTTGTT  13973

EMBOSS_001     14001 AAAAACAGTACAATTCTGTGATGCCATGCGAAATGCTGGTATTGTTGGTG  14050
                     ||||||||||||||||||||||||||||||||||||||||||||||||||
EMBOSS_001     13974 AAAAACAGTACAATTCTGTGATGCCATGCGAAATGCTGGTATTGTTGGTG  14023

EMBOSS_001     14051 TACTGACATTAGATAATCAAGATCTCAATGGTAACTGGTATGATTTCGGT  14100
                     ||||||||||||||||||||||||||||||||||||||||||||||||||
EMBOSS_001     14024 TACTGACATTAGATAATCAAGATCTCAATGGTAACTGGTATGATTTCGGT  14073

EMBOSS_001     14101 GATTTCATACAAACCACGCCAGGTAGTGGAGTTCCTGTTGTAGATTCTTA  14150
                     ||||||||||||||||||||||||||||||||||||||||||||||||||
EMBOSS_001     14074 GATTTCATACAAACCACGCCAGGTAGTGGAGTTCCTGTTGTAGATTCTTA  14123

EMBOSS_001     14151 TTATTCATTGTTAATGCCTATATTAACCTTGACCAGGGCTTTAACTGCAG  14200
                     ||||||||||||||||||||||||||||||||||||||||||||||||||
EMBOSS_001     14124 TTATTCATTGTTAATGCCTATATTAACCTTGACCAGGGCTTTAACTGCAG  14173

EMBOSS_001     14201 AGTCACATGTTGACACTGACTTAACAAAGCCTTACATTAAGTGGGATTTG  14250
                     ||||||||||||||||||||||||||||||||||||||||||||||||||
EMBOSS_001     14174 AGTCACATGTTGACACTGACTTAACAAAGCCTTACATTAAGTGGGATTTG  14223

EMBOSS_001     14251 TTAAAATATGACTTCACGGAAGAGAGGTTAAAACTCTTTGACCGTTATTT  14300
                     ||||||||||||||||||||||||||||||||||||||||||||||||||
EMBOSS_001     14224 TTAAAATATGACTTCACGGAAGAGAGGTTAAAACTCTTTGACCGTTATTT  14273

EMBOSS_001     14301 TAAATATTGGGATCAGACATACCACCCAAATTGTGTTAACTGTTTGGATG  14350
                     ||||||||||||||||||||||||||||||||||||||||||||||||||
EMBOSS_001     14274 TAAATATTGGGATCAGACATACCACCCAAATTGTGTTAACTGTTTGGATG  14323

EMBOSS_001     14351 ACAGATGCATTCTGCATTGTGCAAACTTTAATGTTTTATTCTCTACAGTG  14400
                     ||||||||||||||||||||||||||||||||||||||||||||||||||
EMBOSS_001     14324 ACAGATGCATTCTGCATTGTGCAAACTTTAATGTTTTATTCTCTACAGTG  14373

EMBOSS_001     14401 TTCCCACCTACAAGTTTTGGACCACTAGTGAGAAAAATATTTGTTGATGG  14450
                     ||||||||||||||||||||||||||||||||||||||||||||||||||
EMBOSS_001     14374 TTCCCACCTACAAGTTTTGGACCACTAGTGAGAAAAATATTTGTTGATGG  14423

EMBOSS_001     14451 TGTTCCATTTGTAGTTTCAACTGGATACCACTTCAGAGAGCTAGGTGTTG  14500
                     ||||||||||||||||||||||||||||||||||||||||||||||||||
EMBOSS_001     14424 TGTTCCATTTGTAGTTTCAACTGGATACCACTTCAGAGAGCTAGGTGTTG  14473

EMBOSS_001     14501 TACATAATCAGGATGTAAACTTACATAGCTCTAGACTTAGTTTTAAGGAA  14550
                     ||||||||||||||||||||||||||||||||||||||||||||||||||
EMBOSS_001     14474 TACATAATCAGGATGTAAACTTACATAGCTCTAGACTTAGTTTTAAGGAA  14523

EMBOSS_001     14551 TTACTTGTGTATGCTGCTGACCCTGCTATGCACGCTGCTTCTGGTAATCT  14600
                     ||||||||||||||||||||||||||||||||||||||||||||||||||
EMBOSS_001     14524 TTACTTGTGTATGCTGCTGACCCTGCTATGCACGCTGCTTCTGGTAATCT  14573

EMBOSS_001     14601 ATTACTAGATAAACGCACTACGTGCTTTTCAGTAGCTGCACTTACTAACA  14650
                     ||||||||||||||||||||||||||||||||||||||||||||||||||
EMBOSS_001     14574 ATTACTAGATAAACGCACTACGTGCTTTTCAGTAGCTGCACTTACTAACA  14623

EMBOSS_001     14651 ATGTTGCTTTTCAAACTGTCAAACCCGGTAATTTTAACAAAGACTTCTAT  14700
                     ||||||||||||||||||||||||||||||||||||||||||||||||||
EMBOSS_001     14624 ATGTTGCTTTTCAAACTGTCAAACCCGGTAATTTTAACAAAGACTTCTAT  14673

EMBOSS_001     14701 GACTTTGCTGTGTCTAAGGGTTTCTTTAAGGAAGGAAGTTCTGTTGAATT  14750
                     ||||||||||||||||||||||||||||||||||||||||||||||||||
EMBOSS_001     14674 GACTTTGCTGTGTCTAAGGGTTTCTTTAAGGAAGGAAGTTCTGTTGAATT  14723

EMBOSS_001     14751 AAAACACTTCTTCTTTGCTCAGGATGGTAATGCTGCTATCAGCGATTATG  14800
                     ||||||||||||||||||||||||||||||||||||||||||||||||||
EMBOSS_001     14724 AAAACACTTCTTCTTTGCTCAGGATGGTAATGCTGCTATCAGCGATTATG  14773

EMBOSS_001     14801 ACTACTATCGTTATAATCTACCAACAATGTGTGATATCAGACAACTACTA  14850
                     ||||||||||||||||||||||||||||||||||||||||||||||||||
EMBOSS_001     14774 ACTACTATCGTTATAATCTACCAACAATGTGTGATATCAGACAACTACTA  14823

EMBOSS_001     14851 TTTGTAGTTGAAGTTGTTGATAAGTACTTTGATTGTTACGATGGTGGCTG  14900
                     ||||||||||||||||||||||||||||||||||||||||||||||||||
EMBOSS_001     14824 TTTGTAGTTGAAGTTGTTGATAAGTACTTTGATTGTTACGATGGTGGCTG  14873

EMBOSS_001     14901 TATTAATGCTAACCAAGTCATCGTCAACAACCTAGACAAATCAGCTGGTT  14950
                     ||||||||||||||||||||||||||||||||||||||||||||||||||
EMBOSS_001     14874 TATTAATGCTAACCAAGTCATCGTCAACAACCTAGACAAATCAGCTGGTT  14923

EMBOSS_001     14951 TTCCATTTAATAAATGGGGTAAGGCTAGACTTTATTATGATTCAATGAGT  15000
                     ||||||||||||||||||||||||||||||||||||||||||||||||||
EMBOSS_001     14924 TTCCATTTAATAAATGGGGTAAGGCTAGACTTTATTATGATTCAATGAGT  14973

EMBOSS_001     15001 TATGAGGATCAAGATGCACTTTTCGCATATACAAAACGTAATGTCATCCC  15050
                     ||||||||||||||||||||||||||||||||||||||||||||||||||
EMBOSS_001     14974 TATGAGGATCAAGATGCACTTTTCGCATATACAAAACGTAATGTCATCCC  15023

EMBOSS_001     15051 TACTATAACTCAAATGAATCTTAAGTATGCCATTAGTGCAAAGAATAGAG  15100
                     ||||||||||||||||||||||||||||||||||||||||||||||||||
EMBOSS_001     15024 TACTATAACTCAAATGAATCTTAAGTATGCCATTAGTGCAAAGAATAGAG  15073

EMBOSS_001     15101 CTCGCACCGTAGCTGGTGTCTCTATCTGTAGTACTATGACCAATAGACAG  15150
                     ||||||||||||||||||||||||||||||||||||||||||||||||||
EMBOSS_001     15074 CTCGCACCGTAGCTGGTGTCTCTATCTGTAGTACTATGACCAATAGACAG  15123

EMBOSS_001     15151 TTTCATCAAAAATTATTGAAATCAATAGCCGCCACTAGAGGAGCTACTGT  15200
                     ||||||||||||||||||||||||||||||||||||||||||||||||||
EMBOSS_001     15124 TTTCATCAAAAATTATTGAAATCAATAGCCGCCACTAGAGGAGCTACTGT  15173

EMBOSS_001     15201 AGTAATTGGAACAAGCAAATTCTATGGTGGTTGGCACAACATGTTAAAAA  15250
                     ||||||||||||||||||||||||||||||||||||||||||||||||||
EMBOSS_001     15174 AGTAATTGGAACAAGCAAATTCTATGGTGGTTGGCACAACATGTTAAAAA  15223

EMBOSS_001     15251 CTGTTTATAGTGATGTAGAAAACCCTCACCTTATGGGTTGGGATTATCCT  15300
                     ||||||||||||||||||||||||||||||||||||||||||||||||||
EMBOSS_001     15224 CTGTTTATAGTGATGTAGAAAACCCTCACCTTATGGGTTGGGATTATCCT  15273

EMBOSS_001     15301 AAATGTGATAGAGCCATGCCTAACATGCTTAGAATTATGGCCTCACTTGT  15350
                     ||||||||||||||||||||||||||||||||||||||||||||||||||
EMBOSS_001     15274 AAATGTGATAGAGCCATGCCTAACATGCTTAGAATTATGGCCTCACTTGT  15323

EMBOSS_001     15351 TCTTGCTCGCAAACATACAACGTGTTGTAGCTTGTCACACCGTTTCTATA  15400
                     ||||||||||||||||||||||||||||||||||||||||||||||||||
EMBOSS_001     15324 TCTTGCTCGCAAACATACAACGTGTTGTAGCTTGTCACACCGTTTCTATA  15373

EMBOSS_001     15401 GATTAGCTAATGAGTGTGCTCAAGTATTGAGTGAAATGGTCATGTGTGGC  15450
                     ||||||||||||||||||||||||||||||||||||||||||||||||||
EMBOSS_001     15374 GATTAGCTAATGAGTGTGCTCAAGTATTGAGTGAAATGGTCATGTGTGGC  15423

EMBOSS_001     15451 GGTTCACTATATGTTAAACCAGGTGGAACCTCATCAGGAGATGCCACAAC  15500
                     ||||||||||||||||||||||||||||||||||||||||||||||||||
EMBOSS_001     15424 GGTTCACTATATGTTAAACCAGGTGGAACCTCATCAGGAGATGCCACAAC  15473

EMBOSS_001     15501 TGCTTATGCTAATAGTGTTTTTAACATTTGTCAAGCTGTCACGGCCAATG  15550
                     ||||||||||||||||||||||||||||||||||||||||||||||||||
EMBOSS_001     15474 TGCTTATGCTAATAGTGTTTTTAACATTTGTCAAGCTGTCACGGCCAATG  15523

EMBOSS_001     15551 TTAATGCACTTTTATCTACTGATGGTAACAAAATTGCCGATAAGTATGTC  15600
                     ||||||||||||||||||||||||||||||||||||||||||||||||||
EMBOSS_001     15524 TTAATGCACTTTTATCTACTGATGGTAACAAAATTGCCGATAAGTATGTC  15573

EMBOSS_001     15601 CGCAATTTACAACACAGACTTTATGAGTGTCTCTATAGAAATAGAGATGT  15650
                     ||||||||||||||||||||||||||||||||||||||||||||||||||
EMBOSS_001     15574 CGCAATTTACAACACAGACTTTATGAGTGTCTCTATAGAAATAGAGATGT  15623

EMBOSS_001     15651 TGACACAGACTTTGTGAATGAGTTTTACGCATATTTGCGTAAACATTTCT  15700
                     ||||||||||||||||||||||||||||||||||||||||||||||||||
EMBOSS_001     15624 TGACACAGACTTTGTGAATGAGTTTTACGCATATTTGCGTAAACATTTCT  15673

EMBOSS_001     15701 CAATGATGATACTCTCTGACGATGCTGTTGTGTGTTTCAATAGCACTTAT  15750
                     ||||||||||||||||||||||||||||||||||||||||||||||||||
EMBOSS_001     15674 CAATGATGATACTCTCTGACGATGCTGTTGTGTGTTTCAATAGCACTTAT  15723

EMBOSS_001     15751 GCATCTCAAGGTCTAGTGGCTAGCATAAAGAACTTTAAGTCAGTTCTTTA  15800
                     ||||||||||||||||||||||||||||||||||||||||||||||||||
EMBOSS_001     15724 GCATCTCAAGGTCTAGTGGCTAGCATAAAGAACTTTAAGTCAGTTCTTTA  15773

EMBOSS_001     15801 TTATCAAAACAATGTTTTTATGTCTGAAGCAAAATGTTGGACTGAGACTG  15850
                     ||||||||||||||||||||||||||||||||||||||||||||||||||
EMBOSS_001     15774 TTATCAAAACAATGTTTTTATGTCTGAAGCAAAATGTTGGACTGAGACTG  15823

EMBOSS_001     15851 ACCTTACTAAAGGACCTCATGAATTTTGCTCTCAACATACAATGCTAGTT  15900
                     ||||||||||||||||||||||||||||||||||||||||||||||||||
EMBOSS_001     15824 ACCTTACTAAAGGACCTCATGAATTTTGCTCTCAACATACAATGCTAGTT  15873

EMBOSS_001     15901 AAACAGGGTGATGATTATGTGTACCTTCCTTACCCAGATCCATCAAGAAT  15950
                     ||||||||||||||||||||||||||||||||||||||||||||||||||
EMBOSS_001     15874 AAACAGGGTGATGATTATGTGTACCTTCCTTACCCAGATCCATCAAGAAT  15923

EMBOSS_001     15951 CCTAGGGGCCGGCTGTTTTGTAGATGATATCGTAAAAACAGATGGTACAC  16000
                     ||||||||||||||||||||||||||||||||||||||||||||||||||
EMBOSS_001     15924 CCTAGGGGCCGGCTGTTTTGTAGATGATATCGTAAAAACAGATGGTACAC  15973

EMBOSS_001     16001 TTATGATTGAACGGTTCGTGTCTTTAGCTATAGATGCTTACCCACTTACT  16050
                     ||||||||||||||||||||||||||||||||||||||||||||||||||
EMBOSS_001     15974 TTATGATTGAACGGTTCGTGTCTTTAGCTATAGATGCTTACCCACTTACT  16023

EMBOSS_001     16051 AAACATCCTAATCAGGAGTATGCTGATGTCTTTCATTTGTACTTACAATA  16100
                     ||||||||||||||||||||||||||||||||||||||||||||||||||
EMBOSS_001     16024 AAACATCCTAATCAGGAGTATGCTGATGTCTTTCATTTGTACTTACAATA  16073

EMBOSS_001     16101 CATAAGAAAGCTACATGATGAGTTAACAGGACACATGTTAGACATGTATT  16150
                     ||||||||||||||||||||||||||||||||||||||||||||||||||
EMBOSS_001     16074 CATAAGAAAGCTACATGATGAGTTAACAGGACACATGTTAGACATGTATT  16123

EMBOSS_001     16151 CTGTTATGCTTACTAATGATAACACTTCAAGGTATTGGGAACCTGAGTTT  16200
                     ||||||||||||||||||||||||||||||||||||||||||||||||||
EMBOSS_001     16124 CTGTTATGCTTACTAATGATAACACTTCAAGGTATTGGGAACCTGAGTTT  16173

EMBOSS_001     16201 TATGAGGCTATGTACACACCGCATACAGTCTTACAGGCTGTTGGGGCTTG  16250
                     ||||||||||||||||||||||||||||||||||||||||||||||||||
EMBOSS_001     16174 TATGAGGCTATGTACACACCGCATACAGTCTTACAGGCTGTTGGGGCTTG  16223

EMBOSS_001     16251 TGTTCTTTGCAATTCACAGACTTCATTAAGATGTGGTGCTTGCATACGTA  16300
                     ||||||||||||||||||||||||||||||||||||||||||||||||||
EMBOSS_001     16224 TGTTCTTTGCAATTCACAGACTTCATTAAGATGTGGTGCTTGCATACGTA  16273

EMBOSS_001     16301 GACCATTCTTATGTTGTAAATGCTGTTACGACCATGTCATATCAACATCA  16350
                     ||||||||||||||||||||||||||||||||||||||||||||||||||
EMBOSS_001     16274 GACCATTCTTATGTTGTAAATGCTGTTACGACCATGTCATATCAACATCA  16323

EMBOSS_001     16351 CATAAATTAGTCTTGTCTGTTAATCCGTATGTTTGCAATGCTCCAGGTTG  16400
                     ||||||||||||||||||||||||||||||||||||||||||||||||||
EMBOSS_001     16324 CATAAATTAGTCTTGTCTGTTAATCCGTATGTTTGCAATGCTCCAGGTTG  16373

EMBOSS_001     16401 TGATGTCACAGATGTGACTCAACTTTACTTAGGAGGTATGAGCTATTATT  16450
                     ||||||||||||||||||||||||||||||||||||||||||||||||||
EMBOSS_001     16374 TGATGTCACAGATGTGACTCAACTTTACTTAGGAGGTATGAGCTATTATT  16423

EMBOSS_001     16451 GTAAATCACATAAACCACCCATTAGTTTTCCATTGTGTGCTAATGGACAA  16500
                     ||||||||||||||||||||||||||||||||||||||||||||||||||
EMBOSS_001     16424 GTAAATCACATAAACCACCCATTAGTTTTCCATTGTGTGCTAATGGACAA  16473

EMBOSS_001     16501 GTTTTTGGTTTATATAAAAATACATGTGTTGGTAGCGATAATGTTACTGA  16550
                     ||||||||||||||||||||||||||||||||||||||||||||||||||
EMBOSS_001     16474 GTTTTTGGTTTATATAAAAATACATGTGTTGGTAGCGATAATGTTACTGA  16523

EMBOSS_001     16551 CTTTAATGCAATTGCAACATGTGACTGGACAAATGCTGGTGATTACATTT  16600
                     ||||||||||||||||||||||||||||||||||||||||||||||||||
EMBOSS_001     16524 CTTTAATGCAATTGCAACATGTGACTGGACAAATGCTGGTGATTACATTT  16573

EMBOSS_001     16601 TAGCTAACACCTGTACTGAAAGACTCAAGCTTTTTGCAGCAGAAACGCTC  16650
                     ||||||||||||||||||||||||||||||||||||||||||||||||||
EMBOSS_001     16574 TAGCTAACACCTGTACTGAAAGACTCAAGCTTTTTGCAGCAGAAACGCTC  16623

EMBOSS_001     16651 AAAGCTACTGAGGAGACATTTAAACTGTCTTATGGTATTGCTACTGTACG  16700
                     ||||||||||||||||||||||||||||||||||||||||||||||||||
EMBOSS_001     16624 AAAGCTACTGAGGAGACATTTAAACTGTCTTATGGTATTGCTACTGTACG  16673

EMBOSS_001     16701 TGAAGTGCTGTCTGACAGAGAATTACATCTTTCATGGGAAGTTGGTAAAC  16750
                     ||||||||||||||||||||||||||||||||||||||||||||||||||
EMBOSS_001     16674 TGAAGTGCTGTCTGACAGAGAATTACATCTTTCATGGGAAGTTGGTAAAC  16723

EMBOSS_001     16751 CTAGACCACCACTTAACCGAAATTATGTCTTTACTGGTTATCGTGTAACT  16800
                     ||||||||||||||||||||||||||||||||||||||||||||||||||
EMBOSS_001     16724 CTAGACCACCACTTAACCGAAATTATGTCTTTACTGGTTATCGTGTAACT  16773

EMBOSS_001     16801 AAAAACAGTAAAGTACAAATAGGAGAGTACACCTTTGAAAAAGGTGACTA  16850
                     ||||||||||||||||||||||||||||||||||||||||||||||||||
EMBOSS_001     16774 AAAAACAGTAAAGTACAAATAGGAGAGTACACCTTTGAAAAAGGTGACTA  16823

EMBOSS_001     16851 TGGTGATGCTGTTGTTTACCGAGGTACAACAACTTACAAATTAAATGTTG  16900
                     ||||||||||||||||||||||||||||||||||||||||||||||||||
EMBOSS_001     16824 TGGTGATGCTGTTGTTTACCGAGGTACAACAACTTACAAATTAAATGTTG  16873

EMBOSS_001     16901 GTGATTATTTTGTGCTGACATCACATACAGTAATGCCATTAAGTGCACCT  16950
                     ||||||||||||||||||||||||||||||||||||||||||||||||||
EMBOSS_001     16874 GTGATTATTTTGTGCTGACATCACATACAGTAATGCCATTAAGTGCACCT  16923

EMBOSS_001     16951 ACACTAGTGCCACAAGAGCACTATGTTAGAATTACTGGCTTATACCCAAC  17000
                     ||||||||||||||||||||||||||||||||||||||||||||||||||
EMBOSS_001     16924 ACACTAGTGCCACAAGAGCACTATGTTAGAATTACTGGCTTATACCCAAC  16973

EMBOSS_001     17001 ACTCAATATCTCAGATGAGTTTTCTAGCAATGTTGCAAATTATCAAAAGG  17050
                     ||||||||||||||||||||||||||||||||||||||||||||||||||
EMBOSS_001     16974 ACTCAATATCTCAGATGAGTTTTCTAGCAATGTTGCAAATTATCAAAAGG  17023

EMBOSS_001     17051 TTGGTATGCAAAAGTATTCTACACTCCAGGGACCACCTGGTACTGGTAAG  17100
                     ||||||||||||||||||||||||||||||||||||||||||||||||||
EMBOSS_001     17024 TTGGTATGCAAAAGTATTCTACACTCCAGGGACCACCTGGTACTGGTAAG  17073

EMBOSS_001     17101 AGTCATTTTGCTATTGGCCTAGCTCTCTACTACCCTTCTGCTCGCATAGT  17150
                     ||||||||||||||||||||||||||||||||||||||||||||||||||
EMBOSS_001     17074 AGTCATTTTGCTATTGGCCTAGCTCTCTACTACCCTTCTGCTCGCATAGT  17123

EMBOSS_001     17151 GTATACAGCTTGCTCTCATGCCGCTGTTGATGCACTATGTGAGAAGGCAT  17200
                     ||||||||||||||||||||||||||||||||||||||||||||||||||
EMBOSS_001     17124 GTATACAGCTTGCTCTCATGCCGCTGTTGATGCACTATGTGAGAAGGCAT  17173

EMBOSS_001     17201 TAAAATATTTGCCTATAGATAAATGTAGTAGAATTATACCTGCACGTGCT  17250
                     ||||||||||||||||||||||||||||||||||||||||||||||||||
EMBOSS_001     17174 TAAAATATTTGCCTATAGATAAATGTAGTAGAATTATACCTGCACGTGCT  17223

EMBOSS_001     17251 CGTGTAGAGTGTTTTGATAAATTCAAAGTGAATTCAACATTAGAACAGTA  17300
                     ||||||||||||||||||||||||||||||||||||||||||||||||||
EMBOSS_001     17224 CGTGTAGAGTGTTTTGATAAATTCAAAGTGAATTCAACATTAGAACAGTA  17273

EMBOSS_001     17301 TGTCTTTTGTACTGTAAATGCATTGCCTGAGACGACAGCAGATATAGTTG  17350
                     ||||||||||||||||||||||||||||||||||||||||||||||||||
EMBOSS_001     17274 TGTCTTTTGTACTGTAAATGCATTGCCTGAGACGACAGCAGATATAGTTG  17323

EMBOSS_001     17351 TCTTTGATGAAATTTCAATGGCCACAAATTATGATTTGAGTGTTGTCAAT  17400
                     ||||||||||||||||||||||||||||||||||||||||||||||||||
EMBOSS_001     17324 TCTTTGATGAAATTTCAATGGCCACAAATTATGATTTGAGTGTTGTCAAT  17373

EMBOSS_001     17401 GCCAGATTACGTGCTAAGCACTATGTGTACATTGGCGACCCTGCTCAATT  17450
                     ||||||||||||||||||||||||||||||||||||||||||||||||||
EMBOSS_001     17374 GCCAGATTACGTGCTAAGCACTATGTGTACATTGGCGACCCTGCTCAATT  17423

EMBOSS_001     17451 ACCTGCACCACGCACATTGCTAACTAAGGGCACACTAGAACCAGAATATT  17500
                     ||||||||||||||||||||||||||||||||||||||||||||||||||
EMBOSS_001     17424 ACCTGCACCACGCACATTGCTAACTAAGGGCACACTAGAACCAGAATATT  17473

EMBOSS_001     17501 TCAATTCAGTGTGTAGACTTATGAAAACTATAGGTCCAGACATGTTCCTC  17550
                     ||||||||||||||||||||||||||||||||||||||||||||||||||
EMBOSS_001     17474 TCAATTCAGTGTGTAGACTTATGAAAACTATAGGTCCAGACATGTTCCTC  17523

EMBOSS_001     17551 GGAACTTGTCGGCGTTGTCCTGCTGAAATTGTTGACACTGTGAGTGCTTT  17600
                     ||||||||||||||||||||||||||||||||||||||||||||||||||
EMBOSS_001     17524 GGAACTTGTCGGCGTTGTCCTGCTGAAATTGTTGACACTGTGAGTGCTTT  17573

EMBOSS_001     17601 GGTTTATGATAATAAGCTTAAAGCACATAAAGACAAATCAGCTCAATGCT  17650
                     ||||||||||||||||||||||||||||||||||||||||||||||||||
EMBOSS_001     17574 GGTTTATGATAATAAGCTTAAAGCACATAAAGACAAATCAGCTCAATGCT  17623

EMBOSS_001     17651 TTAAAATGTTTTATAAGGGTGTTATCACGCATGATGTTTCATCTGCAATT  17700
                     ||||||||||||||||||||||||||||||||||||||||||||||||||
EMBOSS_001     17624 TTAAAATGTTTTATAAGGGTGTTATCACGCATGATGTTTCATCTGCAATT  17673

EMBOSS_001     17701 AACAGGCCACAAATAGGCGTGGTAAGAGAATTCCTTACACGTAACCCTGC  17750
                     ||||||||||||||||||||||||||||||||||||||||||||||||||
EMBOSS_001     17674 AACAGGCCACAAATAGGCGTGGTAAGAGAATTCCTTACACGTAACCCTGC  17723

EMBOSS_001     17751 TTGGAGAAAAGCTGTCTTTATTTCACCTTATAATTCACAGAATGCTGTAG  17800
                     ||||||||||||||||||||||||||||||||||||||||||||||||||
EMBOSS_001     17724 TTGGAGAAAAGCTGTCTTTATTTCACCTTATAATTCACAGAATGCTGTAG  17773

EMBOSS_001     17801 CCTCAAAGATTTTGGGACTACCAACTCAAACTGTTGATTCATCACAGGGC  17850
                     ||||||||||||||||||||||||||||||||||||||||||||||||||
EMBOSS_001     17774 CCTCAAAGATTTTGGGACTACCAACTCAAACTGTTGATTCATCACAGGGC  17823

EMBOSS_001     17851 TCAGAATATGACTATGTCATATTCACTCAAACCACTGAAACAGCTCACTC  17900
                     ||||||||||||||||||||||||||||||||||||||||||||||||||
EMBOSS_001     17824 TCAGAATATGACTATGTCATATTCACTCAAACCACTGAAACAGCTCACTC  17873

EMBOSS_001     17901 TTGTAATGTAAACAGATTTAATGTTGCTATTACCAGAGCAAAAGTAGGCA  17950
                     ||||||||||||||||||||||||||||||||||||||||||||||||||
EMBOSS_001     17874 TTGTAATGTAAACAGATTTAATGTTGCTATTACCAGAGCAAAAGTAGGCA  17923

EMBOSS_001     17951 TACTTTGCATAATGTCTGATAGAGACCTTTATGACAAGTTGCAATTTACA  18000
                     ||||||||||||||||||||||||||||||||||||||||||||||||||
EMBOSS_001     17924 TACTTTGCATAATGTCTGATAGAGACCTTTATGACAAGTTGCAATTTACA  17973

EMBOSS_001     18001 AGTCTTGAAATTCCACGTAGGAATGTGGCAACTTTACAAGCTGAAAATGT  18050
                     ||||||||||||||||||||||||||||||||||||||||||||||||||
EMBOSS_001     17974 AGTCTTGAAATTCCACGTAGGAATGTGGCAACTTTACAAGCTGAAAATGT  18023

EMBOSS_001     18051 AACAGGACTCTTTAAAGATTGTAGTAAGGTAATCACTGGGTTACATCCTA  18100
                     ||||||||||||||||||||||||||||||||||||||||||||||||||
EMBOSS_001     18024 AACAGGACTCTTTAAAGATTGTAGTAAGGTAATCACTGGGTTACATCCTA  18073

EMBOSS_001     18101 CACAGGCACCTACACACCTCAGTGTTGACACTAAATTCAAAACTGAAGGT  18150
                     ||||||||||||||||||||||||||||||||||||||||||||||||||
EMBOSS_001     18074 CACAGGCACCTACACACCTCAGTGTTGACACTAAATTCAAAACTGAAGGT  18123

EMBOSS_001     18151 TTATGTGTTGACATACCTGGCATACCTAAGGACATGACCTATAGAAGACT  18200
                     ||||||||||||||||||||||||||||||||||||||||||||||||||
EMBOSS_001     18124 TTATGTGTTGACATACCTGGCATACCTAAGGACATGACCTATAGAAGACT  18173

EMBOSS_001     18201 CATCTCTATGATGGGTTTTAAAATGAATTATCAAGTTAATGGTTACCCTA  18250
                     ||||||||||||||||||||||||||||||||||||||||||||||||||
EMBOSS_001     18174 CATCTCTATGATGGGTTTTAAAATGAATTATCAAGTTAATGGTTACCCTA  18223

EMBOSS_001     18251 ACATGTTTATCACCCGCGAAGAAGCTATAAGACATGTACGTGCATGGATT  18300
                     ||||||||||||||||||||||||||||||||||||||||||||||||||
EMBOSS_001     18224 ACATGTTTATCACCCGCGAAGAAGCTATAAGACATGTACGTGCATGGATT  18273

EMBOSS_001     18301 GGCTTCGATGTCGAGGGGTGTCATGCTACTAGAGAAGCTGTTGGTACCAA  18350
                     ||||||||||||||||||||||||||||||||||||||||||||||||||
EMBOSS_001     18274 GGCTTCGATGTCGAGGGGTGTCATGCTACTAGAGAAGCTGTTGGTACCAA  18323

EMBOSS_001     18351 TTTACCTTTACAGCTAGGTTTTTCTACAGGTGTTAACCTAGTTGCTGTAC  18400
                     ||||||||||||||||||||||||||||||||||||||||||||||||||
EMBOSS_001     18324 TTTACCTTTACAGCTAGGTTTTTCTACAGGTGTTAACCTAGTTGCTGTAC  18373

EMBOSS_001     18401 CTACAGGTTATGTTGATACACCTAATAATACAGATTTTTCCAGAGTTAGT  18450
                     ||||||||||||||||||||||||||||||||||||||||||||||||||
EMBOSS_001     18374 CTACAGGTTATGTTGATACACCTAATAATACAGATTTTTCCAGAGTTAGT  18423

EMBOSS_001     18451 GCTAAACCACCGCCTGGAGATCAATTTAAACACCTCATACCACTTATGTA  18500
                     ||||||||||||||||||||||||||||||||||||||||||||||||||
EMBOSS_001     18424 GCTAAACCACCGCCTGGAGATCAATTTAAACACCTCATACCACTTATGTA  18473

EMBOSS_001     18501 CAAAGGACTTCCTTGGAATGTAGTGCGTATAAAGATTGTACAAATGTTAA  18550
                     ||||||||||||||||||||||||||||||||||||||||||||||||||
EMBOSS_001     18474 CAAAGGACTTCCTTGGAATGTAGTGCGTATAAAGATTGTACAAATGTTAA  18523

EMBOSS_001     18551 GTGACACACTTAAAAATCTCTCTGACAGAGTCGTATTTGTCTTATGGGCA  18600
                     ||||||||||||||||||||||||||||||||||||||||||||||||||
EMBOSS_001     18524 GTGACACACTTAAAAATCTCTCTGACAGAGTCGTATTTGTCTTATGGGCA  18573

EMBOSS_001     18601 CATGGCTTTGAGTTGACATCTATGAAGTATTTTGTGAAAATAGGACCTGA  18650
                     ||||||||||||||||||||||||||||||||||||||||||||||||||
EMBOSS_001     18574 CATGGCTTTGAGTTGACATCTATGAAGTATTTTGTGAAAATAGGACCTGA  18623

EMBOSS_001     18651 GCGCACCTGTTGTCTATGTGATAGACGTGCCACATGCTTTTCCACTGCTT  18700
                     ||||||||||||||||||||||||||||||||||||||||||||||||||
EMBOSS_001     18624 GCGCACCTGTTGTCTATGTGATAGACGTGCCACATGCTTTTCCACTGCTT  18673

EMBOSS_001     18701 CAGACACTTATGCCTGTTGGCATCATTCTATTGGATTTGATTACGTCTAT  18750
                     ||||||||||||||||||||||||||||||||||||||||||||||||||
EMBOSS_001     18674 CAGACACTTATGCCTGTTGGCATCATTCTATTGGATTTGATTACGTCTAT  18723

EMBOSS_001     18751 AATCCGTTTATGATTGATGTTCAACAATGGGGTTTTACAGGTAACCTACA  18800
                     ||||||||||||||||||||||||||||||||||||||||||||||||||
EMBOSS_001     18724 AATCCGTTTATGATTGATGTTCAACAATGGGGTTTTACAGGTAACCTACA  18773

EMBOSS_001     18801 AAGCAACCATGATCTGTATTGTCAAGTCCATGGTAATGCACATGTAGCTA  18850
                     ||||||||||||||||||||||||||||||||||||||||||||||||||
EMBOSS_001     18774 AAGCAACCATGATCTGTATTGTCAAGTCCATGGTAATGCACATGTAGCTA  18823

EMBOSS_001     18851 GTTGTGATGCAATCATGACTAGGTGTCTAGCTGTCCACGAGTGCTTTGTT  18900
                     ||||||||||||||||||||||||||||||||||||||||||||||||||
EMBOSS_001     18824 GTTGTGATGCAATCATGACTAGGTGTCTAGCTGTCCACGAGTGCTTTGTT  18873

EMBOSS_001     18901 AAGCGTGTTGACTGGACTATTGAATATCCTATAATTGGTGATGAACTGAA  18950
                     ||||||||||||||||||||||||||||||||||||||||||||||||||
EMBOSS_001     18874 AAGCGTGTTGACTGGACTATTGAATATCCTATAATTGGTGATGAACTGAA  18923

EMBOSS_001     18951 GATTAATGCGGCTTGTAGAAAGGTTCAACACATGGTTGTTAAAGCTGCAT  19000
                     ||||||||||||||||||||||||||||||||||||||||||||||||||
EMBOSS_001     18924 GATTAATGCGGCTTGTAGAAAGGTTCAACACATGGTTGTTAAAGCTGCAT  18973

EMBOSS_001     19001 TATTAGCAGACAAATTCCCAGTTCTTCACGACATTGGTAACCCTAAAGCT  19050
                     ||||||||||||||||||||||||||||||||||||||||||||||||||
EMBOSS_001     18974 TATTAGCAGACAAATTCCCAGTTCTTCACGACATTGGTAACCCTAAAGCT  19023

EMBOSS_001     19051 ATTAAGTGTGTACCTCAAGCTGATGTAGAATGGAAGTTCTATGATGCACA  19100
                     ||||||||||||||||||||||||||||||||||||||||||||||||||
EMBOSS_001     19024 ATTAAGTGTGTACCTCAAGCTGATGTAGAATGGAAGTTCTATGATGCACA  19073

EMBOSS_001     19101 GCCTTGTAGTGACAAAGCTTATAAAATAGAAGAATTATTCTATTCTTATG  19150
                     ||||||||||||||||||||||||||||||||||||||||||||||||||
EMBOSS_001     19074 GCCTTGTAGTGACAAAGCTTATAAAATAGAAGAATTATTCTATTCTTATG  19123

EMBOSS_001     19151 CCACACATTCTGACAAATTCACAGATGGTGTATGCCTATTTTGGAATTGC  19200
                     ||||||||||||||||||||||||||||||||||||||||||||||||||
EMBOSS_001     19124 CCACACATTCTGACAAATTCACAGATGGTGTATGCCTATTTTGGAATTGC  19173

EMBOSS_001     19201 AATGTCGATAGATATCCTGCTAATTCCATTGTTTGTAGATTTGACACTAG  19250
                     ||||||||||||||||||||||||||||||||||||||||||||||||||
EMBOSS_001     19174 AATGTCGATAGATATCCTGCTAATTCCATTGTTTGTAGATTTGACACTAG  19223

EMBOSS_001     19251 AGTGCTATCTAACCTTAACTTGCCTGGTTGTGATGGTGGCAGTTTGTATG  19300
                     ||||||||||||||||||||||||||||||||||||||||||||||||||
EMBOSS_001     19224 AGTGCTATCTAACCTTAACTTGCCTGGTTGTGATGGTGGCAGTTTGTATG  19273

EMBOSS_001     19301 TAAATAAACATGCATTCCACACACCAGCTTTTGATAAAAGTGCTTTTGTT  19350
                     ||||||||||||||||||||||||||||||||||||||||||||||||||
EMBOSS_001     19274 TAAATAAACATGCATTCCACACACCAGCTTTTGATAAAAGTGCTTTTGTT  19323

EMBOSS_001     19351 AATTTAAAACAATTACCATTTTTCTATTACTCTGACAGTCCATGTGAGTC  19400
                     ||||||||||||||||||||||||||||||||||||||||||||||||||
EMBOSS_001     19324 AATTTAAAACAATTACCATTTTTCTATTACTCTGACAGTCCATGTGAGTC  19373

EMBOSS_001     19401 TCATGGAAAACAAGTAGTGTCAGATATAGATTATGTACCACTAAAGTCTG  19450
                     ||||||||||||||||||||||||||||||||||||||||||||||||||
EMBOSS_001     19374 TCATGGAAAACAAGTAGTGTCAGATATAGATTATGTACCACTAAAGTCTG  19423

EMBOSS_001     19451 CTACGTGTATAACACGTTGCAATTTAGGTGGTGCTGTCTGTAGACATCAT  19500
                     ||||||||||||||||||||||||||||||||||||||||||||||||||
EMBOSS_001     19424 CTACGTGTATAACACGTTGCAATTTAGGTGGTGCTGTCTGTAGACATCAT  19473

EMBOSS_001     19501 GCTAATGAGTACAGATTGTATCTCGATGCTTATAACATGATGATCTCAGC  19550
                     ||||||||||||||||||||||||||||||||||||||||||||||||||
EMBOSS_001     19474 GCTAATGAGTACAGATTGTATCTCGATGCTTATAACATGATGATCTCAGC  19523

EMBOSS_001     19551 TGGCTTTAGCTTGTGGGTTTACAAACAATTTGATACTTATAACCTCTGGA  19600
                     ||||||||||||||||||||||||||||||||||||||||||||||||||
EMBOSS_001     19524 TGGCTTTAGCTTGTGGGTTTACAAACAATTTGATACTTATAACCTCTGGA  19573

EMBOSS_001     19601 ACACTTTTACAAGACTTCAGAGTTTAGAAAATGTGGCTTTTAATGTTGTA  19650
                     ||||||||||||||||||||||||||||||||||||||||||||||||||
EMBOSS_001     19574 ACACTTTTACAAGACTTCAGAGTTTAGAAAATGTGGCTTTTAATGTTGTA  19623

EMBOSS_001     19651 AATAAGGGACACTTTGATGGACAACAGGGTGAAGTACCAGTTTCTATCAT  19700
                     ||||||||||||||||||||||||||||||||||||||||||||||||||
EMBOSS_001     19624 AATAAGGGACACTTTGATGGACAACAGGGTGAAGTACCAGTTTCTATCAT  19673

EMBOSS_001     19701 TAATAACACTGTTTACACAAAAGTTGATGGTGTTGATGTAGAATTGTTTG  19750
                     ||||||||||||||||||||||||||||||||||||||||||||||||||
EMBOSS_001     19674 TAATAACACTGTTTACACAAAAGTTGATGGTGTTGATGTAGAATTGTTTG  19723

EMBOSS_001     19751 AAAATAAAACAACATTACCTGTTAATGTAGCATTTGAGCTTTGGGCTAAG  19800
                     ||||||||||||||||||||||||||||||||||||||||||||||||||
EMBOSS_001     19724 AAAATAAAACAACATTACCTGTTAATGTAGCATTTGAGCTTTGGGCTAAG  19773

EMBOSS_001     19801 CGCAACATTAAACCAGTACCAGAGGTGAAAATACTCAATAATTTGGGTGT  19850
                     ||||||||||||||||||||||||||||||||||||||||||||||||||
EMBOSS_001     19774 CGCAACATTAAACCAGTACCAGAGGTGAAAATACTCAATAATTTGGGTGT  19823

EMBOSS_001     19851 GGACATTGCTGCTAATACTGTGATCTGGGACTACAAAAGAGATGCTCCAG  19900
                     ||||||||||||||||||||||||||||||||||||||||||||||||||
EMBOSS_001     19824 GGACATTGCTGCTAATACTGTGATCTGGGACTACAAAAGAGATGCTCCAG  19873

EMBOSS_001     19901 CACATATATCTACTATTGGTGTTTGTTCTATGACTGACATAGCCAAGAAA  19950
                     ||||||||||||||||||||||||||||||||||||||||||||||||||
EMBOSS_001     19874 CACATATATCTACTATTGGTGTTTGTTCTATGACTGACATAGCCAAGAAA  19923

EMBOSS_001     19951 CCAACTGAAACGATTTGTGCACCACTCACTGTCTTTTTTGATGGTAGAGT  20000
                     ||||||||||||||||||||||||||||||||||||||||||||||||||
EMBOSS_001     19924 CCAACTGAAACGATTTGTGCACCACTCACTGTCTTTTTTGATGGTAGAGT  19973

EMBOSS_001     20001 TGATGGTCAAGTAGACTTATTTAGAAATGCCCGTAATGGTGTTCTTATTA  20050
                     ||||||||||||||||||||||||||||||||||||||||||||||||||
EMBOSS_001     19974 TGATGGTCAAGTAGACTTATTTAGAAATGCCCGTAATGGTGTTCTTATTA  20023

EMBOSS_001     20051 CAGAAGGTAGTGTTAAAGGTTTACAACCATCTGTAGGTCCCAAACAAGCT  20100
                     ||||||||||||||||||||||||||||||||||||||||||||||||||
EMBOSS_001     20024 CAGAAGGTAGTGTTAAAGGTTTACAACCATCTGTAGGTCCCAAACAAGCT  20073

EMBOSS_001     20101 AGTCTTAATGGAGTCACATTAATTGGAGAAGCCGTAAAAACACAGTTCAA  20150
                     ||||||||||||||||||||||||||||||||||||||||||||||||||
EMBOSS_001     20074 AGTCTTAATGGAGTCACATTAATTGGAGAAGCCGTAAAAACACAGTTCAA  20123

EMBOSS_001     20151 TTATTATAAGAAAGTTGATGGTGTTGTCCAACAATTACCTGAAACTTACT  20200
                     ||||||||||||||||||||||||||||||||||||||||||||||||||
EMBOSS_001     20124 TTATTATAAGAAAGTTGATGGTGTTGTCCAACAATTACCTGAAACTTACT  20173

EMBOSS_001     20201 TTACTCAGAGTAGAAATTTACAAGAATTTAAACCCAGGAGTCAAATGGAA  20250
                     ||||||||||||||||||||||||||||||||||||||||||||||||||
EMBOSS_001     20174 TTACTCAGAGTAGAAATTTACAAGAATTTAAACCCAGGAGTCAAATGGAA  20223

EMBOSS_001     20251 ATTGATTTCTTAGAATTAGCTATGGATGAATTCATTGAACGGTATAAATT  20300
                     ||||||||||||||||||||||||||||||||||||||||||||||||||
EMBOSS_001     20224 ATTGATTTCTTAGAATTAGCTATGGATGAATTCATTGAACGGTATAAATT  20273

EMBOSS_001     20301 AGAAGGCTATGCCTTCGAACATATCGTTTATGGAGATTTTAGTCATAGTC  20350
                     ||||||||||||||||||||||||||||||||||||||||||||||||||
EMBOSS_001     20274 AGAAGGCTATGCCTTCGAACATATCGTTTATGGAGATTTTAGTCATAGTC  20323

EMBOSS_001     20351 AGTTAGGTGGTTTACATCTACTGATTGGACTAGCTAAACGTTTTAAGGAA  20400
                     ||||||||||||||||||||||||||||||||||||||||||||||||||
EMBOSS_001     20324 AGTTAGGTGGTTTACATCTACTGATTGGACTAGCTAAACGTTTTAAGGAA  20373

EMBOSS_001     20401 TCACCTTTTGAATTAGAAGATTTTATTCCTATGGACAGTACAGTTAAAAA  20450
                     ||||||||||||||||||||||||||||||||||||||||||||||||||
EMBOSS_001     20374 TCACCTTTTGAATTAGAAGATTTTATTCCTATGGACAGTACAGTTAAAAA  20423

EMBOSS_001     20451 CTATTTCATAACAGATGCGCAAACAGGTTCATCTAAGTGTGTGTGTTCTG  20500
                     ||||||||||||||||||||||||||||||||||||||||||||||||||
EMBOSS_001     20424 CTATTTCATAACAGATGCGCAAACAGGTTCATCTAAGTGTGTGTGTTCTG  20473

EMBOSS_001     20501 TTATTGATTTATTACTTGATGATTTTGTTGAAATAATAAAATCCCAAGAT  20550
                     ||||||||||||||||||||||||||||||||||||||||||||||||||
EMBOSS_001     20474 TTATTGATTTATTACTTGATGATTTTGTTGAAATAATAAAATCCCAAGAT  20523

EMBOSS_001     20551 TTATCTGTAGTTTCTAAGGTTGTCAAAGTGACTATTGACTATACAGAAAT  20600
                     ||||||||||||||||||||||||||||||||||||||||||||||||||
EMBOSS_001     20524 TTATCTGTAGTTTCTAAGGTTGTCAAAGTGACTATTGACTATACAGAAAT  20573

EMBOSS_001     20601 TTCATTTATGCTTTGGTGTAAAGATGGCCATGTAGAAACATTTTACCCAA  20650
                     ||||||||||||||||||||||||||||||||||||||||||||||||||
EMBOSS_001     20574 TTCATTTATGCTTTGGTGTAAAGATGGCCATGTAGAAACATTTTACCCAA  20623

EMBOSS_001     20651 AATTACAATCTAGTCAAGCGTGGCAACCGGGTGTTGCTATGCCTAATCTT  20700
                     ||||||||||||||||||||||||||||||||||||||||||||||||||
EMBOSS_001     20624 AATTACAATCTAGTCAAGCGTGGCAACCGGGTGTTGCTATGCCTAATCTT  20673

EMBOSS_001     20701 TACAAAATGCAAAGAATGCTATTAGAAAAGTGTGACCTTCAAAATTATGG  20750
                     ||||||||||||||||||||||||||||||||||||||||||||||||||
EMBOSS_001     20674 TACAAAATGCAAAGAATGCTATTAGAAAAGTGTGACCTTCAAAATTATGG  20723

EMBOSS_001     20751 TGATAGTGCAACATTACCTAAAGGCATAATGATGAATGTCGCAAAATATA  20800
                     ||||||||||||||||||||||||||||||||||||||||||||||||||
EMBOSS_001     20724 TGATAGTGCAACATTACCTAAAGGCATAATGATGAATGTCGCAAAATATA  20773

EMBOSS_001     20801 CTCAACTGTGTCAATATTTAAACACATTAACATTAGCTGTACCCTATAAT  20850
                     ||||||||||||||||||||||||||||||||||||||||||||||||||
EMBOSS_001     20774 CTCAACTGTGTCAATATTTAAACACATTAACATTAGCTGTACCCTATAAT  20823

EMBOSS_001     20851 ATGAGAGTTATACATTTTGGTGCTGGTTCTGATAAAGGAGTTGCACCAGG  20900
                     ||||||||||||||||||||||||||||||||||||||||||||||||||
EMBOSS_001     20824 ATGAGAGTTATACATTTTGGTGCTGGTTCTGATAAAGGAGTTGCACCAGG  20873

EMBOSS_001     20901 TACAGCTGTTTTAAGACAGTGGTTGCCTACGGGTACGCTGCTTGTCGATT  20950
                     ||||||||||||||||||||||||||||||||||||||||||||||||||
EMBOSS_001     20874 TACAGCTGTTTTAAGACAGTGGTTGCCTACGGGTACGCTGCTTGTCGATT  20923

EMBOSS_001     20951 CAGATCTTAATGACTTTGTCTCTGATGCAGATTCAACTTTGATTGGTGAT  21000
                     ||||||||||||||||||||||||||||||||||||||||||||||||||
EMBOSS_001     20924 CAGATCTTAATGACTTTGTCTCTGATGCAGATTCAACTTTGATTGGTGAT  20973

EMBOSS_001     21001 TGTGCAACTGTACATACAGCTAATAAATGGGATCTCATTATTAGTGATAT  21050
                     ||||||||||||||||||||||||||||||||||||||||||||||||||
EMBOSS_001     20974 TGTGCAACTGTACATACAGCTAATAAATGGGATCTCATTATTAGTGATAT  21023

EMBOSS_001     21051 GTACGACCCTAAGACTAAAAATGTTACAAAAGAAAATGACTCTAAAGAGG  21100
                     ||||||||||||||||||||||||||||||||||||||||||||||||||
EMBOSS_001     21024 GTACGACCCTAAGACTAAAAATGTTACAAAAGAAAATGACTCTAAAGAGG  21073

EMBOSS_001     21101 GTTTTTTCACTTACATTTGTGGGTTTATACAACAAAAGCTAGCTCTTGGA  21150
                     ||||||||||||||||||||||||||||||||||||||||||||||||||
EMBOSS_001     21074 GTTTTTTCACTTACATTTGTGGGTTTATACAACAAAAGCTAGCTCTTGGA  21123

EMBOSS_001     21151 GGTTCCGTGGCTATAAAGATAACAGAACATTCTTGGAATGCTGATCTTTA  21200
                     ||||||||||||||||||||||||||||||||||||||||||||||||||
EMBOSS_001     21124 GGTTCCGTGGCTATAAAGATAACAGAACATTCTTGGAATGCTGATCTTTA  21173

EMBOSS_001     21201 TAAGCTCATGGGACACTTCGCATGGTGGACAGCCTTTGTTACTAATGTGA  21250
                     ||||||||||||||||||||||||||||||||||||||||||||||||||
EMBOSS_001     21174 TAAGCTCATGGGACACTTCGCATGGTGGACAGCCTTTGTTACTAATGTGA  21223

EMBOSS_001     21251 ATGCGTCATCATCTGAAGCATTTTTAATTGGATGTAATTATCTTGGCAAA  21300
                     ||||||||||||||||||||||||||||||||||||||||||||||||||
EMBOSS_001     21224 ATGCGTCATCATCTGAAGCATTTTTAATTGGATGTAATTATCTTGGCAAA  21273

EMBOSS_001     21301 CCACGCGAACAAATAGATGGTTATGTCATGCATGCAAATTACATATTTTG  21350
                     ||||||||||||||||||||||||||||||||||||||||||||||||||
EMBOSS_001     21274 CCACGCGAACAAATAGATGGTTATGTCATGCATGCAAATTACATATTTTG  21323

EMBOSS_001     21351 GAGGAATACAAATCCAATTCAGTTGTCTTCCTATTCTTTATTTGACATGA  21400
                     ||||||||||||||||||||||||||||||||||||||||||||||||||
EMBOSS_001     21324 GAGGAATACAAATCCAATTCAGTTGTCTTCCTATTCTTTATTTGACATGA  21373

EMBOSS_001     21401 GTAAATTTCCCCTTAAATTAAGGGGTACTGCTGTTATGTCTTTAAAAGAA  21450
                     ||||||||||||||||||||||||||||||||||||||||||||||||||
EMBOSS_001     21374 GTAAATTTCCCCTTAAATTAAGGGGTACTGCTGTTATGTCTTTAAAAGAA  21423

EMBOSS_001     21451 GGTCAAATCAATGATATGATTTTATCTCTTCTTAGTAAAGGTAGACTTAT  21500
                     ||||||||||||||||||||||||||||||||||||||||||||||||||
EMBOSS_001     21424 GGTCAAATCAATGATATGATTTTATCTCTTCTTAGTAAAGGTAGACTTAT  21473

EMBOSS_001     21501 AATTAGAGAAAACAACAGAGTTGTTATTTCTAGTGATGTTCTTGTTAACA  21550
                     ||||||||||||||||||||||||||||||||||||||||||||||||||
EMBOSS_001     21474 AATTAGAGAAAACAACAGAGTTGTTATTTCTAGTGATGTTCTTGTTAACA  21523

EMBOSS_001     21551 ACTAAACGAACAATGTTTGTTTTTCTTGTTTTATTGCCACTAGTCTCTAG  21600
                     ||||||||||||||||||||||||||||||||||||||||||||||||||
EMBOSS_001     21524 ACTAAACGAACAATGTTTGTTTTTCTTGTTTTATTGCCACTAGTCTCTAG  21573

EMBOSS_001     21601 TCAGTGTGTTAATCTTACAACCAGAACTCAATTACCCCCTGCATACACTA  21650
                     ||||||||||||||||||||||||||||||||||||||||||||||||||
EMBOSS_001     21574 TCAGTGTGTTAATCTTACAACCAGAACTCAATTACCCCCTGCATACACTA  21623

EMBOSS_001     21651 ATTCTTTCACACGTGGTGTTTATTACCCTGACAAAGTTTTCAGATCCTCA  21700
                     ||||||||||||||||||||||||||||||||||||||||||||||||||
EMBOSS_001     21624 ATTCTTTCACACGTGGTGTTTATTACCCTGACAAAGTTTTCAGATCCTCA  21673

EMBOSS_001     21701 GTTTTACATTCAACTCAGGACTTGTTCTTACCTTTCTTTTCCAATGTTAC  21750
                     ||||||||||||||||||||||||||||||||||||||||||||||||||
EMBOSS_001     21674 GTTTTACATTCAACTCAGGACTTGTTCTTACCTTTCTTTTCCAATGTTAC  21723

EMBOSS_001     21751 TTGGTTCCATGCTATACATGTCTCTGGGACCAATGGTACTAAGAGGTTTG  21800
                     ||||||||||||||||||||||||||||||||||||||||||||||||||
EMBOSS_001     21724 TTGGTTCCATGCTATACATGTCTCTGGGACCAATGGTACTAAGAGGTTTG  21773

EMBOSS_001     21801 ATAACCCTGTCCTACCATTTAATGATGGTGTTTATTTTGCTTCCACTGAG  21850
                     ||||||||||||||||||||||||||||||||||||||||||||||||||
EMBOSS_001     21774 ATAACCCTGTCCTACCATTTAATGATGGTGTTTATTTTGCTTCCACTGAG  21823

EMBOSS_001     21851 AAGTCTAACATAATAAGAGGCTGGATTTTTGGTACTACTTTAGATTCGAA  21900
                     ||||||||||||||||||||||||||||||||||||||||||||||||||
EMBOSS_001     21824 AAGTCTAACATAATAAGAGGCTGGATTTTTGGTACTACTTTAGATTCGAA  21873

EMBOSS_001     21901 GACCCAGTCCCTACTTATTGTTAATAACGCTACTAATGTTGTTATTAAAG  21950
                     ||||||||||||||||||||||||||||||||||||||||||||||||||
EMBOSS_001     21874 GACCCAGTCCCTACTTATTGTTAATAACGCTACTAATGTTGTTATTAAAG  21923

EMBOSS_001     21951 TCTGTGAATTTCAATTTTGTAATGATCCATTTTTGGGTGTTTATTACCAC  22000
                     ||||||||||||||||||||||||||||||||||||||||||||||||||
EMBOSS_001     21924 TCTGTGAATTTCAATTTTGTAATGATCCATTTTTGGGTGTTTATTACCAC  21973

EMBOSS_001     22001 AAAAACAACAAAAGTTGGATGGAAAGTGAGTTCAGAGTTTATTCTAGTGC  22050
                     ||||||||||||||||||||||||||||||||||||||||||||||||||
EMBOSS_001     21974 AAAAACAACAAAAGTTGGATGGAAAGTGAGTTCAGAGTTTATTCTAGTGC  22023

EMBOSS_001     22051 GAATAATTGCACTTTTGAATATGTCTCTCAGCCTTTTCTTATGGACCTTG  22100
                     ||||||||||||||||||||||||||||||||||||||||||||||||||
EMBOSS_001     22024 GAATAATTGCACTTTTGAATATGTCTCTCAGCCTTTTCTTATGGACCTTG  22073

EMBOSS_001     22101 AAGGAAAACAGGGTAATTTCAAAAATCTTAGGGAATTTGTGTTTAAGAAT  22150
                     ||||||||||||||||||||||||||||||||||||||||||||||||||
EMBOSS_001     22074 AAGGAAAACAGGGTAATTTCAAAAATCTTAGGGAATTTGTGTTTAAGAAT  22123

EMBOSS_001     22151 ATTGATGGTTATTTTAAAATATATTCTAAGCACACGCCTATTAATTTAGT  22200
                     ||||||||||||||||||||||||||||||||||||||||||||||||||
EMBOSS_001     22124 ATTGATGGTTATTTTAAAATATATTCTAAGCACACGCCTATTAATTTAGT  22173

EMBOSS_001     22201 GCGTGATCTCCCTCAGGGTTTTTCGGCTTTAGAACCATTGGTAGATTTGC  22250
                     ||||||||||||||||||||||||||||||||||||||||||||||||||
EMBOSS_001     22174 GCGTGATCTCCCTCAGGGTTTTTCGGCTTTAGAACCATTGGTAGATTTGC  22223

EMBOSS_001     22251 CAATAGGTATTAACATCACTAGGTTTCAAACTTTACTTGCTTTACATAGA  22300
                     ||||||||||||||||||||||||||||||||||||||||||||||||||
EMBOSS_001     22224 CAATAGGTATTAACATCACTAGGTTTCAAACTTTACTTGCTTTACATAGA  22273

EMBOSS_001     22301 AGTTATTTGACTCCTGGTGATTCTTCTTCAGGTTGGACAGCTGGTGCTGC  22350
                     ||||||||||||||||||||||||||||||||||||||||||||||||||
EMBOSS_001     22274 AGTTATTTGACTCCTGGTGATTCTTCTTCAGGTTGGACAGCTGGTGCTGC  22323

EMBOSS_001     22351 AGCTTATTATGTGGGTTATCTTCAACCTAGGACTTTTCTATTAAAATATA  22400
                     ||||||||||||||||||||||||||||||||||||||||||||||||||
EMBOSS_001     22324 AGCTTATTATGTGGGTTATCTTCAACCTAGGACTTTTCTATTAAAATATA  22373

EMBOSS_001     22401 ATGAAAATGGAACCATTACAGATGCTGTAGACTGTGCACTTGACCCTCTC  22450
                     ||||||||||||||||||||||||||||||||||||||||||||||||||
EMBOSS_001     22374 ATGAAAATGGAACCATTACAGATGCTGTAGACTGTGCACTTGACCCTCTC  22423

EMBOSS_001     22451 TCAGAAACAAAGTGTACGTTGAAATCCTTCACTGTAGAAAAAGGAATCTA  22500
                     ||||||||||||||||||||||||||||||||||||||||||||||||||
EMBOSS_001     22424 TCAGAAACAAAGTGTACGTTGAAATCCTTCACTGTAGAAAAAGGAATCTA  22473

EMBOSS_001     22501 TCAAACTTCTAACTTTAGAGTCCAACCAACAGAATCTATTGTTAGATTTC  22550
                     ||||||||||||||||||||||||||||||||||||||||||||||||||
EMBOSS_001     22474 TCAAACTTCTAACTTTAGAGTCCAACCAACAGAATCTATTGTTAGATTTC  22523

EMBOSS_001     22551 CTAATATTACAAACTTGTGCCCTTTTGGTGAAGTTTTTAACGCCACCAGA  22600
                     ||||||||||||||||||||||||||||||||||||||||||||||||||
EMBOSS_001     22524 CTAATATTACAAACTTGTGCCCTTTTGGTGAAGTTTTTAACGCCACCAGA  22573

EMBOSS_001     22601 TTTGCATCTGTTTATGCTTGGAACAGGAAGAGAATCAGCAACTGTGTTGC  22650
                     ||||||||||||||||||||||||||||||||||||||||||||||||||
EMBOSS_001     22574 TTTGCATCTGTTTATGCTTGGAACAGGAAGAGAATCAGCAACTGTGTTGC  22623

EMBOSS_001     22651 TGATTATTCTGTCCTATATAATTCCGCATCATTTTCCACTTTTAAGTGTT  22700
                     ||||||||||||||||||||||||||||||||||||||||||||||||||
EMBOSS_001     22624 TGATTATTCTGTCCTATATAATTCCGCATCATTTTCCACTTTTAAGTGTT  22673

EMBOSS_001     22701 ATGGAGTGTCTCCTACTAAATTAAATGATCTCTGCTTTACTAATGTCTAT  22750
                     ||||||||||||||||||||||||||||||||||||||||||||||||||
EMBOSS_001     22674 ATGGAGTGTCTCCTACTAAATTAAATGATCTCTGCTTTACTAATGTCTAT  22723

EMBOSS_001     22751 GCAGATTCATTTGTAATTAGAGGTGATGAAGTCAGACAAATCGCTCCAGG  22800
                     ||||||||||||||||||||||||||||||||||||||||||||||||||
EMBOSS_001     22724 GCAGATTCATTTGTAATTAGAGGTGATGAAGTCAGACAAATCGCTCCAGG  22773

EMBOSS_001     22801 GCAAACTGGAAAGATTGCTGATTATAATTATAAATTACCAGATGATTTTA  22850
                     ||||||||||||||||||||||||||||||||||||||||||||||||||
EMBOSS_001     22774 GCAAACTGGAAAGATTGCTGATTATAATTATAAATTACCAGATGATTTTA  22823

EMBOSS_001     22851 CAGGCTGCGTTATAGCTTGGAATTCTAACAATCTTGATTCTAAGGTTGGT  22900
                     ||||||||||||||||||||||||||||||||||||||||||||||||||
EMBOSS_001     22824 CAGGCTGCGTTATAGCTTGGAATTCTAACAATCTTGATTCTAAGGTTGGT  22873

EMBOSS_001     22901 GGTAATTATAATTACCTGTATAGATTGTTTAGGAAGTCTAATCTCAAACC  22950
                     ||||||||||||||||||||||||||||||||||||||||||||||||||
EMBOSS_001     22874 GGTAATTATAATTACCTGTATAGATTGTTTAGGAAGTCTAATCTCAAACC  22923

EMBOSS_001     22951 TTTTGAGAGAGATATTTCAACTGAAATCTATCAGGCCGGTAGCACACCTT  23000
                     ||||||||||||||||||||||||||||||||||||||||||||||||||
EMBOSS_001     22924 TTTTGAGAGAGATATTTCAACTGAAATCTATCAGGCCGGTAGCACACCTT  22973

EMBOSS_001     23001 GTAATGGTGTTGAAGGTTTTAATTGTTACTTTCCTTTACAATCATATGGT  23050
                     ||||||||||||||||||||||||||||||||||||||||||||||||||
EMBOSS_001     22974 GTAATGGTGTTGAAGGTTTTAATTGTTACTTTCCTTTACAATCATATGGT  23023

EMBOSS_001     23051 TTCCAACCCACTAATGGTGTTGGTTACCAACCATACAGAGTAGTAGTACT  23100
                     ||||||||||||||||||||||||||||||||||||||||||||||||||
EMBOSS_001     23024 TTCCAACCCACTAATGGTGTTGGTTACCAACCATACAGAGTAGTAGTACT  23073

EMBOSS_001     23101 TTCTTTTGAACTTCTACATGCACCAGCAACTGTTTGTGGACCTAAAAAGT  23150
                     ||||||||||||||||||||||||||||||||||||||||||||||||||
EMBOSS_001     23074 TTCTTTTGAACTTCTACATGCACCAGCAACTGTTTGTGGACCTAAAAAGT  23123

EMBOSS_001     23151 CTACTAATTTGGTTAAAAACAAATGTGTCAATTTCAACTTCAATGGTTTA  23200
                     ||||||||||||||||||||||||||||||||||||||||||||||||||
EMBOSS_001     23124 CTACTAATTTGGTTAAAAACAAATGTGTCAATTTCAACTTCAATGGTTTA  23173

EMBOSS_001     23201 ACAGGCACAGGTGTTCTTACTGAGTCTAACAAAAAGTTTCTGCCTTTCCA  23250
                     ||||||||||||||||||||||||||||||||||||||||||||||||||
EMBOSS_001     23174 ACAGGCACAGGTGTTCTTACTGAGTCTAACAAAAAGTTTCTGCCTTTCCA  23223

EMBOSS_001     23251 ACAATTTGGCAGAGACATTGCTGACACTACTGATGCTGTCCGTGATCCAC  23300
                     ||||||||||||||||||||||||||||||||||||||||||||||||||
EMBOSS_001     23224 ACAATTTGGCAGAGACATTGCTGACACTACTGATGCTGTCCGTGATCCAC  23273

EMBOSS_001     23301 AGACACTTGAGATTCTTGACATTACACCATGTTCTTTTGGTGGTGTCAGT  23350
                     ||||||||||||||||||||||||||||||||||||||||||||||||||
EMBOSS_001     23274 AGACACTTGAGATTCTTGACATTACACCATGTTCTTTTGGTGGTGTCAGT  23323

EMBOSS_001     23351 GTTATAACACCAGGAACAAATACTTCTAACCAGGTTGCTGTTCTTTATCA  23400
                     ||||||||||||||||||||||||||||||||||||||||||||||||||
EMBOSS_001     23324 GTTATAACACCAGGAACAAATACTTCTAACCAGGTTGCTGTTCTTTATCA  23373

EMBOSS_001     23401 GGATGTTAACTGCACAGAAGTCCCTGTTGCTATTCATGCAGATCAACTTA  23450
                     ||||||||||||||||||||||||||||||||||||||||||||||||||
EMBOSS_001     23374 GGATGTTAACTGCACAGAAGTCCCTGTTGCTATTCATGCAGATCAACTTA  23423

EMBOSS_001     23451 CTCCTACTTGGCGTGTTTATTCTACAGGTTCTAATGTTTTTCAAACACGT  23500
                     ||||||||||||||||||||||||||||||||||||||||||||||||||
EMBOSS_001     23424 CTCCTACTTGGCGTGTTTATTCTACAGGTTCTAATGTTTTTCAAACACGT  23473

EMBOSS_001     23501 GCAGGCTGTTTAATAGGGGCTGAACATGTCAACAACTCATATGAGTGTGA  23550
                     ||||||||||||||||||||||||||||||||||||||||||||||||||
EMBOSS_001     23474 GCAGGCTGTTTAATAGGGGCTGAACATGTCAACAACTCATATGAGTGTGA  23523

EMBOSS_001     23551 CATACCCATTGGTGCAGGTATATGCGCTAGTTATCAGACTCAGACTAATT  23600
                     ||||||||||||||||||||||||||||||||||||||||||||||||||
EMBOSS_001     23524 CATACCCATTGGTGCAGGTATATGCGCTAGTTATCAGACTCAGACTAATT  23573

EMBOSS_001     23601 CTCCTCGGCGGGCACGTAGTGTAGCTAGTCAATCCATCATTGCCTACACT  23650
                     ||||||||||||||||||||||||||||||||||||||||||||||||||
EMBOSS_001     23574 CTCCTCGGCGGGCACGTAGTGTAGCTAGTCAATCCATCATTGCCTACACT  23623

EMBOSS_001     23651 ATGTCACTTGGTGCAGAAAATTCAGTTGCTTACTCTAATAACTCTATTGC  23700
                     ||||||||||||||||||||||||||||||||||||||||||||||||||
EMBOSS_001     23624 ATGTCACTTGGTGCAGAAAATTCAGTTGCTTACTCTAATAACTCTATTGC  23673

EMBOSS_001     23701 CATACCCACAAATTTTACTATTAGTGTTACCACAGAAATTCTACCAGTGT  23750
                     ||||||||||||||||||||||||||||||||||||||||||||||||||
EMBOSS_001     23674 CATACCCACAAATTTTACTATTAGTGTTACCACAGAAATTCTACCAGTGT  23723

EMBOSS_001     23751 CTATGACCAAGACATCAGTAGATTGTACAATGTACATTTGTGGTGATTCA  23800
                     ||||||||||||||||||||||||||||||||||||||||||||||||||
EMBOSS_001     23724 CTATGACCAAGACATCAGTAGATTGTACAATGTACATTTGTGGTGATTCA  23773

EMBOSS_001     23801 ACTGAATGCAGCAATCTTTTGTTGCAATATGGCAGTTTTTGTACACAATT  23850
                     ||||||||||||||||||||||||||||||||||||||||||||||||||
EMBOSS_001     23774 ACTGAATGCAGCAATCTTTTGTTGCAATATGGCAGTTTTTGTACACAATT  23823

EMBOSS_001     23851 AAACCGTGCTTTAACTGGAATAGCTGTTGAACAAGACAAAAACACCCAAG  23900
                     ||||||||||||||||||||||||||||||||||||||||||||||||||
EMBOSS_001     23824 AAACCGTGCTTTAACTGGAATAGCTGTTGAACAAGACAAAAACACCCAAG  23873

EMBOSS_001     23901 AAGTTTTTGCACAAGTCAAACAAATTTACAAAACACCACCAATTAAAGAT  23950
                     ||||||||||||||||||||||||||||||||||||||||||||||||||
EMBOSS_001     23874 AAGTTTTTGCACAAGTCAAACAAATTTACAAAACACCACCAATTAAAGAT  23923

EMBOSS_001     23951 TTTGGTGGTTTTAATTTTTCACAAATATTACCAGATCCATCAAAACCAAG  24000
                     ||||||||||||||||||||||||||||||||||||||||||||||||||
EMBOSS_001     23924 TTTGGTGGTTTTAATTTTTCACAAATATTACCAGATCCATCAAAACCAAG  23973

EMBOSS_001     24001 CAAGAGGTCATTTATTGAAGATCTACTTTTCAACAAAGTGACACTTGCAG  24050
                     ||||||||||||||||||||||||||||||||||||||||||||||||||
EMBOSS_001     23974 CAAGAGGTCATTTATTGAAGATCTACTTTTCAACAAAGTGACACTTGCAG  24023

EMBOSS_001     24051 ATGCTGGCTTCATCAAACAATATGGTGATTGCCTTGGTGATATTGCTGCT  24100
                     ||||||||||||||||||||||||||||||||||||||||||||||||||
EMBOSS_001     24024 ATGCTGGCTTCATCAAACAATATGGTGATTGCCTTGGTGATATTGCTGCT  24073

EMBOSS_001     24101 AGAGACCTCATTTGTGCACAAAAGTTTAACGGCCTTACTGTTTTGCCACC  24150
                     ||||||||||||||||||||||||||||||||||||||||||||||||||
EMBOSS_001     24074 AGAGACCTCATTTGTGCACAAAAGTTTAACGGCCTTACTGTTTTGCCACC  24123

EMBOSS_001     24151 TTTGCTCACAGATGAAATGATTGCTCAATACACTTCTGCACTGTTAGCGG  24200
                     ||||||||||||||||||||||||||||||||||||||||||||||||||
EMBOSS_001     24124 TTTGCTCACAGATGAAATGATTGCTCAATACACTTCTGCACTGTTAGCGG  24173

EMBOSS_001     24201 GTACAATCACTTCTGGTTGGACCTTTGGTGCAGGTGCTGCATTACAAATA  24250
                     ||||||||||||||||||||||||||||||||||||||||||||||||||
EMBOSS_001     24174 GTACAATCACTTCTGGTTGGACCTTTGGTGCAGGTGCTGCATTACAAATA  24223

EMBOSS_001     24251 CCATTTGCTATGCAAATGGCTTATAGGTTTAATGGTATTGGAGTTACACA  24300
                     ||||||||||||||||||||||||||||||||||||||||||||||||||
EMBOSS_001     24224 CCATTTGCTATGCAAATGGCTTATAGGTTTAATGGTATTGGAGTTACACA  24273

EMBOSS_001     24301 GAATGTTCTCTATGAGAACCAAAAATTGATTGCCAACCAATTTAATAGTG  24350
                     ||||||||||||||||||||||||||||||||||||||||||||||||||
EMBOSS_001     24274 GAATGTTCTCTATGAGAACCAAAAATTGATTGCCAACCAATTTAATAGTG  24323

EMBOSS_001     24351 CTATTGGCAAAATTCAAGACTCACTTTCTTCCACAGCAAGTGCACTTGGA  24400
                     ||||||||||||||||||||||||||||||||||||||||||||||||||
EMBOSS_001     24324 CTATTGGCAAAATTCAAGACTCACTTTCTTCCACAGCAAGTGCACTTGGA  24373

EMBOSS_001     24401 AAACTTCAAGATGTGGTCAACCAAAATGCACAAGCTTTAAACACGCTTGT  24450
                     ||||||||||||||||||||||||||||||||||||||||||||||||||
EMBOSS_001     24374 AAACTTCAAGATGTGGTCAACCAAAATGCACAAGCTTTAAACACGCTTGT  24423

EMBOSS_001     24451 TAAACAACTTAGCTCCAATTTTGGTGCAATTTCAAGTGTTTTAAATGATA  24500
                     ||||||||||||||||||||||||||||||||||||||||||||||||||
EMBOSS_001     24424 TAAACAACTTAGCTCCAATTTTGGTGCAATTTCAAGTGTTTTAAATGATA  24473

EMBOSS_001     24501 TCCTTTCACGTCTTGACAAAGTTGAGGCTGAAGTGCAAATTGATAGGTTG  24550
                     ||||||||||||||||||||||||||||||||||||||||||||||||||
EMBOSS_001     24474 TCCTTTCACGTCTTGACAAAGTTGAGGCTGAAGTGCAAATTGATAGGTTG  24523

EMBOSS_001     24551 ATCACAGGCAGACTTCAAAGTTTGCAGACATATGTGACTCAACAATTAAT  24600
                     ||||||||||||||||||||||||||||||||||||||||||||||||||
EMBOSS_001     24524 ATCACAGGCAGACTTCAAAGTTTGCAGACATATGTGACTCAACAATTAAT  24573

EMBOSS_001     24601 TAGAGCTGCAGAAATCAGAGCTTCTGCTAATCTTGCTGCTACTAAAATGT  24650
                     ||||||||||||||||||||||||||||||||||||||||||||||||||
EMBOSS_001     24574 TAGAGCTGCAGAAATCAGAGCTTCTGCTAATCTTGCTGCTACTAAAATGT  24623

EMBOSS_001     24651 CAGAGTGTGTACTTGGACAATCAAAAAGAGTTGATTTTTGTGGAAAGGGC  24700
                     ||||||||||||||||||||||||||||||||||||||||||||||||||
EMBOSS_001     24624 CAGAGTGTGTACTTGGACAATCAAAAAGAGTTGATTTTTGTGGAAAGGGC  24673

EMBOSS_001     24701 TATCATCTTATGTCCTTCCCTCAGTCAGCACCTCATGGTGTAGTCTTCTT  24750
                     ||||||||||||||||||||||||||||||||||||||||||||||||||
EMBOSS_001     24674 TATCATCTTATGTCCTTCCCTCAGTCAGCACCTCATGGTGTAGTCTTCTT  24723

EMBOSS_001     24751 GCATGTGACTTATGTCCCTGCACAAGAAAAGAACTTCACAACTGCTCCTG  24800
                     ||||||||||||||||||||||||||||||||||||||||||||||||||
EMBOSS_001     24724 GCATGTGACTTATGTCCCTGCACAAGAAAAGAACTTCACAACTGCTCCTG  24773

EMBOSS_001     24801 CCATTTGTCATGATGGAAAAGCACACTTTCCTCGTGAAGGTGTCTTTGTT  24850
                     ||||||||||||||||||||||||||||||||||||||||||||||||||
EMBOSS_001     24774 CCATTTGTCATGATGGAAAAGCACACTTTCCTCGTGAAGGTGTCTTTGTT  24823

EMBOSS_001     24851 TCAAATGGCACACACTGGTTTGTAACACAAAGGAATTTTTATGAACCACA  24900
                     ||||||||||||||||||||||||||||||||||||||||||||||||||
EMBOSS_001     24824 TCAAATGGCACACACTGGTTTGTAACACAAAGGAATTTTTATGAACCACA  24873

EMBOSS_001     24901 AATCATTACTACAGACAACACATTTGTGTCTGGTAACTGTGATGTTGTAA  24950
                     ||||||||||||||||||||||||||||||||||||||||||||||||||
EMBOSS_001     24874 AATCATTACTACAGACAACACATTTGTGTCTGGTAACTGTGATGTTGTAA  24923

EMBOSS_001     24951 TAGGAATTGTCAACAACACAGTTTATGATCCTTTGCAACCTGAATTAGAC  25000
                     ||||||||||||||||||||||||||||||||||||||||||||||||||
EMBOSS_001     24924 TAGGAATTGTCAACAACACAGTTTATGATCCTTTGCAACCTGAATTAGAC  24973

EMBOSS_001     25001 TCATTCAAGGAGGAGTTAGATAAATATTTTAAGAATCATACATCACCAGA  25050
                     ||||||||||||||||||||||||||||||||||||||||||||||||||
EMBOSS_001     24974 TCATTCAAGGAGGAGTTAGATAAATATTTTAAGAATCATACATCACCAGA  25023

EMBOSS_001     25051 TGTTGATTTAGGTGACATCTCTGGCATTAATGCTTCAGTTGTAAACATTC  25100
                     ||||||||||||||||||||||||||||||||||||||||||||||||||
EMBOSS_001     25024 TGTTGATTTAGGTGACATCTCTGGCATTAATGCTTCAGTTGTAAACATTC  25073

EMBOSS_001     25101 AAAAAGAAATTGACCGCCTCAATGAGGTTGCCAAGAATTTAAATGAATCT  25150
                     ||||||||||||||||||||||||||||||||||||||||||||||||||
EMBOSS_001     25074 AAAAAGAAATTGACCGCCTCAATGAGGTTGCCAAGAATTTAAATGAATCT  25123

EMBOSS_001     25151 CTCATCGATCTCCAAGAACTTGGAAAGTATGAGCAGTATATAAAATGGCC  25200
                     ||||||||||||||||||||||||||||||||||||||||||||||||||
EMBOSS_001     25124 CTCATCGATCTCCAAGAACTTGGAAAGTATGAGCAGTATATAAAATGGCC  25173

EMBOSS_001     25201 ATGGTACATTTGGCTAGGTTTTATAGCTGGCTTGATTGCCATAGTAATGG  25250
                     ||||||||||||||||||||||||||||||||||||||||||||||||||
EMBOSS_001     25174 ATGGTACATTTGGCTAGGTTTTATAGCTGGCTTGATTGCCATAGTAATGG  25223

EMBOSS_001     25251 TGACAATTATGCTTTGCTGTATGACCAGTTGCTGTAGTTGTCTCAAGGGC  25300
                     ||||||||||||||||||||||||||||||||||||||||||||||||||
EMBOSS_001     25224 TGACAATTATGCTTTGCTGTATGACCAGTTGCTGTAGTTGTCTCAAGGGC  25273

EMBOSS_001     25301 TGTTGTTCTTGTGGATCCTGCTGCAAATTTGATGAAGACGACTCTGAGCC  25350
                     ||||||||||||||||||||||||||||||||||||||||||||||||||
EMBOSS_001     25274 TGTTGTTCTTGTGGATCCTGCTGCAAATTTGATGAAGACGACTCTGAGCC  25323

EMBOSS_001     25351 AGTGCTCAAAGGAGTCAAATTACATTACACATAAACGAACTTATGGATTT  25400
                     ||||||||||||||||||||||||||||||||||||||||||||||||||
EMBOSS_001     25324 AGTGCTCAAAGGAGTCAAATTACATTACACATAAACGAACTTATGGATTT  25373

EMBOSS_001     25401 GTTTATGAGAATCTTCACAATTGGAACTGTAACTTTGAAGCAAGGTGAAA  25450
                     ||||||||||||||||||||||||||||||||||||||||||||||||||
EMBOSS_001     25374 GTTTATGAGAATCTTCACAATTGGAACTGTAACTTTGAAGCAAGGTGAAA  25423

EMBOSS_001     25451 TCAAGGATGCTACTCCTTCAGATTTTGTTCGCGCTACTGCAACGATACCG  25500
                     ||||||||||||||||||||||||||||||||||||||||||||||||||
EMBOSS_001     25424 TCAAGGATGCTACTCCTTCAGATTTTGTTCGCGCTACTGCAACGATACCG  25473

EMBOSS_001     25501 ATACAAGCCTCACTCCCTTTCGGATGGCTTATTGTTGGCGTTGCACTTCT  25550
                     ||||||||||||||||||||||||||||||||||||||||||||||||||
EMBOSS_001     25474 ATACAAGCCTCACTCCCTTTCGGATGGCTTATTGTTGGCGTTGCACTTCT  25523

EMBOSS_001     25551 TGCTGTTTTTCAGAGCGCTTCCAAAATCATAACCCTCAAAAAGAGATGGC  25600
                     ||||||||||||||||||||||||||||||||||||||||||||||||||
EMBOSS_001     25524 TGCTGTTTTTCAGAGCGCTTCCAAAATCATAACCCTCAAAAAGAGATGGC  25573

EMBOSS_001     25601 AACTAGCACTCTCCAAGGGTGTTCACTTTGTTTGCAACTTGCTGTTGTTG  25650
                     ||||||||||||||||||||||||||||||||||||||||||||||||||
EMBOSS_001     25574 AACTAGCACTCTCCAAGGGTGTTCACTTTGTTTGCAACTTGCTGTTGTTG  25623

EMBOSS_001     25651 TTTGTAACAGTTTACTCACACCTTTTGCTCGTTGCTGCTGGCCTTGAAGC  25700
                     ||||||||||||||||||||||||||||||||||||||||||||||||||
EMBOSS_001     25624 TTTGTAACAGTTTACTCACACCTTTTGCTCGTTGCTGCTGGCCTTGAAGC  25673

EMBOSS_001     25701 CCCTTTTCTCTATCTTTATGCTTTAGTCTACTTCTTGCAGAGTATAAACT  25750
                     ||||||||||||||||||||||||||||||||||||||||||||||||||
EMBOSS_001     25674 CCCTTTTCTCTATCTTTATGCTTTAGTCTACTTCTTGCAGAGTATAAACT  25723

EMBOSS_001     25751 TTGTAAGAATAATAATGAGGCTTTGGCTTTGCTGGAAATGCCGTTCCAAA  25800
                     ||||||||||||||||||||||||||||||||||||||||||||||||||
EMBOSS_001     25724 TTGTAAGAATAATAATGAGGCTTTGGCTTTGCTGGAAATGCCGTTCCAAA  25773

EMBOSS_001     25801 AACCCATTACTTTATGATGCCAACTATTTTCTTTGCTGGCATACTAATTG  25850
                     ||||||||||||||||||||||||||||||||||||||||||||||||||
EMBOSS_001     25774 AACCCATTACTTTATGATGCCAACTATTTTCTTTGCTGGCATACTAATTG  25823

EMBOSS_001     25851 TTACGACTATTGTATACCTTACAATAGTGTAACTTCTTCAATTGTCATTA  25900
                     ||||||||||||||||||||||||||||||||||||||||||||||||||
EMBOSS_001     25824 TTACGACTATTGTATACCTTACAATAGTGTAACTTCTTCAATTGTCATTA  25873

EMBOSS_001     25901 CTTCAGGTGATGGCACAACAAGTCCTATTTCTGAACATGACTACCAGATT  25950
                     ||||||||||||||||||||||||||||||||||||||||||||||||||
EMBOSS_001     25874 CTTCAGGTGATGGCACAACAAGTCCTATTTCTGAACATGACTACCAGATT  25923

EMBOSS_001     25951 GGTGGTTATACTGAAAAATGGGAATCTGGAGTAAAAGACTGTGTTGTATT  26000
                     ||||||||||||||||||||||||||||||||||||||||||||||||||
EMBOSS_001     25924 GGTGGTTATACTGAAAAATGGGAATCTGGAGTAAAAGACTGTGTTGTATT  25973

EMBOSS_001     26001 ACACAGTTACTTCACTTCAGACTATTACCAGCTGTACTCAACTCAATTGA  26050
                     ||||||||||||||||||||||||||||||||||||||||||||||||||
EMBOSS_001     25974 ACACAGTTACTTCACTTCAGACTATTACCAGCTGTACTCAACTCAATTGA  26023

EMBOSS_001     26051 GTACAGACACTGGTGTTGAACATGTTACCTTCTTCATCTACAATAAAATT  26100
                     ||||||||||||||||||||||||||||||||||||||||||||||||||
EMBOSS_001     26024 GTACAGACACTGGTGTTGAACATGTTACCTTCTTCATCTACAATAAAATT  26073

EMBOSS_001     26101 GTTGATGAGCCTGAAGAACATGTCCAAATTCACACAATCGACGGTTCATC  26150
                     ||||||||||||||||||||||||||||||||||||||||||||||||||
EMBOSS_001     26074 GTTGATGAGCCTGAAGAACATGTCCAAATTCACACAATCGACGGTTCATC  26123

EMBOSS_001     26151 CGGAGTTGTTAATCCAGTAATGGAACCAATTTATGATGAACCGACGACGA  26200
                     ||||||||||||||||||||||||||||||||||||||||||||||||||
EMBOSS_001     26124 CGGAGTTGTTAATCCAGTAATGGAACCAATTTATGATGAACCGACGACGA  26173

EMBOSS_001     26201 CTACTAGCGTGCCTTTGTAAGCACAAGCTGATGAGTACGAACTTATGTAC  26250
                     ||||||||||||||||||||||||||||||||||||||||||||||||||
EMBOSS_001     26174 CTACTAGCGTGCCTTTGTAAGCACAAGCTGATGAGTACGAACTTATGTAC  26223

EMBOSS_001     26251 TCATTCGTTTCGGAAGAGACAGGTACGTTAATAGTTAATAGCGTACTTCT  26300
                     ||||||||||||||||||||||||||||||||||||||||||||||||||
EMBOSS_001     26224 TCATTCGTTTCGGAAGAGACAGGTACGTTAATAGTTAATAGCGTACTTCT  26273

EMBOSS_001     26301 TTTTCTTGCTTTCGTGGTATTCTTGCTAGTTACACTAGCCATCCTTACTG  26350
                     ||||||||||||||||||||||||||||||||||||||||||||||||||
EMBOSS_001     26274 TTTTCTTGCTTTCGTGGTATTCTTGCTAGTTACACTAGCCATCCTTACTG  26323

EMBOSS_001     26351 CGCTTCGATTGTGTGCGTACTGCTGCAATATTGTTAACGTGAGTCTTGTA  26400
                     ||||||||||||||||||||||||||||||||||||||||||||||||||
EMBOSS_001     26324 CGCTTCGATTGTGTGCGTACTGCTGCAATATTGTTAACGTGAGTCTTGTA  26373

EMBOSS_001     26401 AAACCTTCTTTTTACGTTTACTCTCGTGTTAAAAATCTGAATTCTTCTAG  26450
                     ||||||||||||||||||||||||||||||||||||||||||||||||||
EMBOSS_001     26374 AAACCTTCTTTTTACGTTTACTCTCGTGTTAAAAATCTGAATTCTTCTAG  26423

EMBOSS_001     26451 AGTTCCTGATCTTCTGGTCTAAACGAACTAAATATTATATTAGTTTTTCT  26500
                     ||||||||||||||||||||||||||||||||||||||||||||||||||
EMBOSS_001     26424 AGTTCCTGATCTTCTGGTCTAAACGAACTAAATATTATATTAGTTTTTCT  26473

EMBOSS_001     26501 GTTTGGAACTTTAATTTTAGCCATGGCAGATTCCAACGGTACTATTACCG  26550
                     ||||||||||||||||||||||||||||||||||||||||||||||||||
EMBOSS_001     26474 GTTTGGAACTTTAATTTTAGCCATGGCAGATTCCAACGGTACTATTACCG  26523

EMBOSS_001     26551 TTGAAGAGCTTAAAAAGCTCCTTGAACAATGGAACCTAGTAATAGGTTTC  26600
                     ||||||||||||||||||||||||||||||||||||||||||||||||||
EMBOSS_001     26524 TTGAAGAGCTTAAAAAGCTCCTTGAACAATGGAACCTAGTAATAGGTTTC  26573

EMBOSS_001     26601 CTATTCCTTACATGGATTTGTCTTCTACAATTTGCCTATGCCAACAGGAA  26650
                     ||||||||||||||||||||||||||||||||||||||||||||||||||
EMBOSS_001     26574 CTATTCCTTACATGGATTTGTCTTCTACAATTTGCCTATGCCAACAGGAA  26623

EMBOSS_001     26651 TAGGTTTTTGTATATAATTAAGTTAATTTTCCTCTGGCTGTTATGGCCAG  26700
                     ||||||||||||||||||||||||||||||||||||||||||||||||||
EMBOSS_001     26624 TAGGTTTTTGTATATAATTAAGTTAATTTTCCTCTGGCTGTTATGGCCAG  26673

EMBOSS_001     26701 TAACTTTAGCTTGTTTTGTGCTTGCTGCTGTTTACAGAATAAATTGGATC  26750
                     ||||||||||||||||||||||||||||||||||||||||||||||||||
EMBOSS_001     26674 TAACTTTAGCTTGTTTTGTGCTTGCTGCTGTTTACAGAATAAATTGGATC  26723

EMBOSS_001     26751 ACCGGTGGAATTGCTATCGCAATGGCTTGTCTTGTAGGCTTGATGTGGCT  26800
                     ||||||||||||||||||||||||||||||||||||||||||||||||||
EMBOSS_001     26724 ACCGGTGGAATTGCTATCGCAATGGCTTGTCTTGTAGGCTTGATGTGGCT  26773

EMBOSS_001     26801 CAGCTACTTCATTGCTTCTTTCAGACTGTTTGCGCGTACGCGTTCCATGT  26850
                     ||||||||||||||||||||||||||||||||||||||||||||||||||
EMBOSS_001     26774 CAGCTACTTCATTGCTTCTTTCAGACTGTTTGCGCGTACGCGTTCCATGT  26823

EMBOSS_001     26851 GGTCATTCAATCCAGAAACTAACATTCTTCTCAACGTGCCACTCCATGGC  26900
                     ||||||||||||||||||||||||||||||||||||||||||||||||||
EMBOSS_001     26824 GGTCATTCAATCCAGAAACTAACATTCTTCTCAACGTGCCACTCCATGGC  26873

EMBOSS_001     26901 ACTATTCTGACCAGACCGCTTCTAGAAAGTGAACTCGTAATCGGAGCTGT  26950
                     ||||||||||||||||||||||||||||||||||||||||||||||||||
EMBOSS_001     26874 ACTATTCTGACCAGACCGCTTCTAGAAAGTGAACTCGTAATCGGAGCTGT  26923

EMBOSS_001     26951 GATCCTTCGTGGACATCTTCGTATTGCTGGACACCATCTAGGACGCTGTG  27000
                     ||||||||||||||||||||||||||||||||||||||||||||||||||
EMBOSS_001     26924 GATCCTTCGTGGACATCTTCGTATTGCTGGACACCATCTAGGACGCTGTG  26973

EMBOSS_001     27001 ACATCAAGGACCTGCCTAAAGAAATCACTGTTGCTACATCACGAACGCTT  27050
                     ||||||||||||||||||||||||||||||||||||||||||||||||||
EMBOSS_001     26974 ACATCAAGGACCTGCCTAAAGAAATCACTGTTGCTACATCACGAACGCTT  27023

EMBOSS_001     27051 TCTTATTACAAATTGGGAGCTTCGCAGCGTGTAGCAGGTGACTCAGGTTT  27100
                     ||||||||||||||||||||||||||||||||||||||||||||||||||
EMBOSS_001     27024 TCTTATTACAAATTGGGAGCTTCGCAGCGTGTAGCAGGTGACTCAGGTTT  27073

EMBOSS_001     27101 TGCTGCATACAGTCGCTACAGGATTGGCAACTATAAATTAAACACAGACC  27150
                     ||||||||||||||||||||||||||||||||||||||||||||||||||
EMBOSS_001     27074 TGCTGCATACAGTCGCTACAGGATTGGCAACTATAAATTAAACACAGACC  27123

EMBOSS_001     27151 ATTCCAGTAGCAGTGACAATATTGCTTTGCTTGTACAGTAAGTGACAACA  27200
                     ||||||||||||||||||||||||||||||||||||||||||||||||||
EMBOSS_001     27124 ATTCCAGTAGCAGTGACAATATTGCTTTGCTTGTACAGTAAGTGACAACA  27173

EMBOSS_001     27201 GATGTTTCATCTCGTTGACTTTCAGGTTACTATAGCAGAGATATTACTAA  27250
                     ||||||||||||||||||||||||||||||||||||||||||||||||||
EMBOSS_001     27174 GATGTTTCATCTCGTTGACTTTCAGGTTACTATAGCAGAGATATTACTAA  27223

EMBOSS_001     27251 TTATTATGAGGACTTTTAAAGTTTCCATTTGGAATCTTGATTACATCATA  27300
                     ||||||||||||||||||||||||||||||||||||||||||||||||||
EMBOSS_001     27224 TTATTATGAGGACTTTTAAAGTTTCCATTTGGAATCTTGATTACATCATA  27273

EMBOSS_001     27301 AACCTCATAATTAAAAATTTATCTAAGTCACTAACTGAGAATAAATATTC  27350
                     ||||||||||||||||||||||||||||||||||||||||||||||||||
EMBOSS_001     27274 AACCTCATAATTAAAAATTTATCTAAGTCACTAACTGAGAATAAATATTC  27323

EMBOSS_001     27351 TCAATTAGATGAAGAGCAACCAATGGAGATTGATTAAACGAACATGAAAA  27400
                     ||||||||||||||||||||||||||||||||||||||||||||||||||
EMBOSS_001     27324 TCAATTAGATGAAGAGCAACCAATGGAGATTGATTAAACGAACATGAAAA  27373

EMBOSS_001     27401 TTATTCTTTTCTTGGCACTGATAACACTCGCTACTTGTGAGCTTTATCAC  27450
                     ||||||||||||||||||||||||||||||||||||||||||||||||||
EMBOSS_001     27374 TTATTCTTTTCTTGGCACTGATAACACTCGCTACTTGTGAGCTTTATCAC  27423

EMBOSS_001     27451 TACCAAGAGTGTGTTAGAGGTACAACAGTACTTTTAAAAGAACCTTGCTC  27500
                     ||||||||||||||||||||||||||||||||||||||||||||||||||
EMBOSS_001     27424 TACCAAGAGTGTGTTAGAGGTACAACAGTACTTTTAAAAGAACCTTGCTC  27473

EMBOSS_001     27501 TTCTGGAACATACGAGGGCAATTCACCATTTCATCCTCTAGCTGATAACA  27550
                     ||||||||||||||||||||||||||||||||||||||||||||||||||
EMBOSS_001     27474 TTCTGGAACATACGAGGGCAATTCACCATTTCATCCTCTAGCTGATAACA  27523

EMBOSS_001     27551 AATTTGCACTGACTTGCTTTAGCACTCAATTTGCTTTTGCTTGTCCTGAC  27600
                     ||||||||||||||||||||||||||||||||||||||||||||||||||
EMBOSS_001     27524 AATTTGCACTGACTTGCTTTAGCACTCAATTTGCTTTTGCTTGTCCTGAC  27573

EMBOSS_001     27601 GGCGTAAAACACGTCTATCAGTTACGTGCCAGATCAGTTTCACCTAAACT  27650
                     ||||||||||||||||||||||||||||||||||||||||||||||||||
EMBOSS_001     27574 GGCGTAAAACACGTCTATCAGTTACGTGCCAGATCAGTTTCACCTAAACT  27623

EMBOSS_001     27651 GTTCATCAGACAAGAGGAAGTTCAAGAACTTTACTCTCCAATTTTTCTTA  27700
                     ||||||||||||||||||||||||||||||||||||||||||||||||||
EMBOSS_001     27624 GTTCATCAGACAAGAGGAAGTTCAAGAACTTTACTCTCCAATTTTTCTTA  27673

EMBOSS_001     27701 TTGTTGCGGCAATAGTGTTTATAACACTTTGCTTCACACTCAAAAGAAAG  27750
                     ||||||||||||||||||||||||||||||||||||||||||||||||||
EMBOSS_001     27674 TTGTTGCGGCAATAGTGTTTATAACACTTTGCTTCACACTCAAAAGAAAG  27723

EMBOSS_001     27751 ACAGAATGATTGAACTTTCATTAATTGACTTCTATTTGTGCTTTTTAGCC  27800
                     ||||||||||||||||||||||||||||||||||||||||||||||||||
EMBOSS_001     27724 ACAGAATGATTGAACTTTCATTAATTGACTTCTATTTGTGCTTTTTAGCC  27773

EMBOSS_001     27801 TTTCTGCTATTCCTTGTTTTAATTATGCTTATTATCTTTTGGTTCTCACT  27850
                     ||||||||||||||||||||||||||||||||||||||||||||||||||
EMBOSS_001     27774 TTTCTGCTATTCCTTGTTTTAATTATGCTTATTATCTTTTGGTTCTCACT  27823

EMBOSS_001     27851 TGAACTGCAAGATCATAATGAAACTTGTCACGCCTAAACGAACATGAAAT  27900
                     ||||||||||||||||||||||||||||||||||||||||||||||||||
EMBOSS_001     27824 TGAACTGCAAGATCATAATGAAACTTGTCACGCCTAAACGAACATGAAAT  27873

EMBOSS_001     27901 TTCTTGTTTTCTTAGGAATCATCACAACTGTAGCTGCATTTCACCAAGAA  27950
                     ||||||||||||||||||||||||||||||||||||||||||||||||||
EMBOSS_001     27874 TTCTTGTTTTCTTAGGAATCATCACAACTGTAGCTGCATTTCACCAAGAA  27923

EMBOSS_001     27951 TGTAGTTTACAGTCATGTACTCAACATCAACCATATGTAGTTGATGACCC  28000
                     ||||||||||||||||||||||||||||||||||||||||||||||||||
EMBOSS_001     27924 TGTAGTTTACAGTCATGTACTCAACATCAACCATATGTAGTTGATGACCC  27973

EMBOSS_001     28001 GTGTCCTATTCACTTCTATTCTAAATGGTATATTAGAGTAGGAGCTAGAA  28050
                     ||||||||||||||||||||||||||||||||||||||||||||||||||
EMBOSS_001     27974 GTGTCCTATTCACTTCTATTCTAAATGGTATATTAGAGTAGGAGCTAGAA  28023

EMBOSS_001     28051 AATCAGCACCTTTAATTGAATTGTGCGTGGATGAGGCTGGTTCTAAATCA  28100
                     ||||||||||||||||||||||||||||||||||||||||||||||||||
EMBOSS_001     28024 AATCAGCACCTTTAATTGAATTGTGCGTGGATGAGGCTGGTTCTAAATCA  28073

EMBOSS_001     28101 CCCATTCAGTACATCGATATCGGTAATTATACAGTTTCCTGTTTACCTTT  28150
                     ||||||||||||||||||||||||||||||||||||||||||||||||||
EMBOSS_001     28074 CCCATTCAGTACATCGATATCGGTAATTATACAGTTTCCTGTTTACCTTT  28123

EMBOSS_001     28151 TACAATTAATTGCCAGGAACCTAAATTGGGTAGTCTTGTAGTGCGTTGTT  28200
                     ||||||||||||||||||||||||||||||||||||||||||||||||||
EMBOSS_001     28124 TACAATTAATTGCCAGGAACCTAAATTGGGTAGTCTTGTAGTGCGTTGTT  28173

EMBOSS_001     28201 CGTTCTATGAAGACTTTTTAGAGTATCATGACGTTCGTGTTGTTTTAGAT  28250
                     ||||||||||||||||||||||||||||||||||||||||||||||||||
EMBOSS_001     28174 CGTTCTATGAAGACTTTTTAGAGTATCATGACGTTCGTGTTGTTTTAGAT  28223

EMBOSS_001     28251 TTCATCTAAACGAACAAACTAAAATGTCTGATAATGGACCCCAAAATCAG  28300
                     ||||||||||||||||||||||||||||||||||||||||||||||||||
EMBOSS_001     28224 TTCATCTAAACGAACAAACTAAAATGTCTGATAATGGACCCCAAAATCAG  28273

EMBOSS_001     28301 CGAAATGCACCCCGCATTACGTTTGGTGGACCCTCAGATTCAACTGGCAG  28350
                     ||||||||||||||||||||||||||||||||||||||||||||||||||
EMBOSS_001     28274 CGAAATGCACCCCGCATTACGTTTGGTGGACCCTCAGATTCAACTGGCAG  28323

EMBOSS_001     28351 TAACCAGAATGGAGAACGCAGTGGGGCGCGATCAAAACAACGTCGGCCCC  28400
                     ||||||||||||||||||||||||||||||||||||||||||||||||||
EMBOSS_001     28324 TAACCAGAATGGAGAACGCAGTGGGGCGCGATCAAAACAACGTCGGCCCC  28373

EMBOSS_001     28401 AAGGTTTACCCAATAATACTGCGTCTTGGTTCACCGCTCTCACTCAACAT  28450
                     ||||||||||||||||||||||||||||||||||||||||||||||||||
EMBOSS_001     28374 AAGGTTTACCCAATAATACTGCGTCTTGGTTCACCGCTCTCACTCAACAT  28423

EMBOSS_001     28451 GGCAAGGAAGACCTTAAATTCCCTCGAGGACAAGGCGTTCCAATTAACAC  28500
                     ||||||||||||||||||||||||||||||||||||||||||||||||||
EMBOSS_001     28424 GGCAAGGAAGACCTTAAATTCCCTCGAGGACAAGGCGTTCCAATTAACAC  28473

EMBOSS_001     28501 CAATAGCAGTCCAGATGACCAAATTGGCTACTACCGAAGAGCTACCAGAC  28550
                     ||||||||||||||||||||||||||||||||||||||||||||||||||
EMBOSS_001     28474 CAATAGCAGTCCAGATGACCAAATTGGCTACTACCGAAGAGCTACCAGAC  28523

EMBOSS_001     28551 GAATTCGTGGTGGTGACGGTAAAATGAAAGATCTCAGTCCAAGATGGTAT  28600
                     ||||||||||||||||||||||||||||||||||||||||||||||||||
EMBOSS_001     28524 GAATTCGTGGTGGTGACGGTAAAATGAAAGATCTCAGTCCAAGATGGTAT  28573

EMBOSS_001     28601 TTCTACTACCTAGGAACTGGGCCAGAAGCTGGACTTCCCTATGGTGCTAA  28650
                     ||||||||||||||||||||||||||||||||||||||||||||||||||
EMBOSS_001     28574 TTCTACTACCTAGGAACTGGGCCAGAAGCTGGACTTCCCTATGGTGCTAA  28623

EMBOSS_001     28651 CAAAGACGGCATCATATGGGTTGCAACTGAGGGAGCCTTGAATACACCAA  28700
                     ||||||||||||||||||||||||||||||||||||||||||||||||||
EMBOSS_001     28624 CAAAGACGGCATCATATGGGTTGCAACTGAGGGAGCCTTGAATACACCAA  28673

EMBOSS_001     28701 AAGATCACATTGGCACCCGCAATCCTGCTAACAATGCTGCAATCGTGCTA  28750
                     ||||||||||||||||||||||||||||||||||||||||||||||||||
EMBOSS_001     28674 AAGATCACATTGGCACCCGCAATCCTGCTAACAATGCTGCAATCGTGCTA  28723

EMBOSS_001     28751 CAACTTCCTCAAGGAACAACATTGCCAAAAGGCTTCTACGCAGAAGGGAG  28800
                     ||||||||||||||||||||||||||||||||||||||||||||||||||
EMBOSS_001     28724 CAACTTCCTCAAGGAACAACATTGCCAAAAGGCTTCTACGCAGAAGGGAG  28773

EMBOSS_001     28801 CAGAGGCGGCAGTCAAGCCTCTTCTCGTTCCTCATCACGTAGTCGCAACA  28850
                     ||||||||||||||||||||||||||||||||||||||||||||||||||
EMBOSS_001     28774 CAGAGGCGGCAGTCAAGCCTCTTCTCGTTCCTCATCACGTAGTCGCAACA  28823

EMBOSS_001     28851 GTTCAAGAAATTCAACTCCAGGCAGCAGTAGGGGAACTTCTCCTGCTAGA  28900
                     ||||||||||||||||||||||||||||||||||||||||||||||||||
EMBOSS_001     28824 GTTCAAGAAATTCAACTCCAGGCAGCAGTAGGGGAACTTCTCCTGCTAGA  28873

EMBOSS_001     28901 ATGGCTGGCAATGGCGGTGATGCTGCTCTTGCTTTGCTGCTGCTTGACAG  28950
                     ||||||||||||||||||||||||||||||||||||||||||||||||||
EMBOSS_001     28874 ATGGCTGGCAATGGCGGTGATGCTGCTCTTGCTTTGCTGCTGCTTGACAG  28923

EMBOSS_001     28951 ATTGAACCAGCTTGAGAGCAAAATGTCTGGTAAAGGCCAACAACAACAAG  29000
                     ||||||||||||||||||||||||||||||||||||||||||||||||||
EMBOSS_001     28924 ATTGAACCAGCTTGAGAGCAAAATGTCTGGTAAAGGCCAACAACAACAAG  28973

EMBOSS_001     29001 GCCAAACTGTCACTAAGAAATCTGCTGCTGAGGCTTCTAAGAAGCCTCGG  29050
                     ||||||||||||||||||||||||||||||||||||||||||||||||||
EMBOSS_001     28974 GCCAAACTGTCACTAAGAAATCTGCTGCTGAGGCTTCTAAGAAGCCTCGG  29023

EMBOSS_001     29051 CAAAAACGTACTGCCACTAAAGCATACAATGTAACACAAGCTTTCGGCAG  29100
                     ||||||||||||||||||||||||||||||||||||||||||||||||||
EMBOSS_001     29024 CAAAAACGTACTGCCACTAAAGCATACAATGTAACACAAGCTTTCGGCAG  29073

EMBOSS_001     29101 ACGTGGTCCAGAACAAACCCAAGGAAATTTTGGGGACCAGGAACTAATCA  29150
                     ||||||||||||||||||||||||||||||||||||||||||||||||||
EMBOSS_001     29074 ACGTGGTCCAGAACAAACCCAAGGAAATTTTGGGGACCAGGAACTAATCA  29123

EMBOSS_001     29151 GACAAGGAACTGATTACAAACATTGGCCGCAAATTGCACAATTTGCCCCC  29200
                     ||||||||||||||||||||||||||||||||||||||||||||||||||
EMBOSS_001     29124 GACAAGGAACTGATTACAAACATTGGCCGCAAATTGCACAATTTGCCCCC  29173

EMBOSS_001     29201 AGCGCTTCAGCGTTCTTCGGAATGTCGCGCATTGGCATGGAAGTCACACC  29250
                     ||||||||||||||||||||||||||||||||||||||||||||||||||
EMBOSS_001     29174 AGCGCTTCAGCGTTCTTCGGAATGTCGCGCATTGGCATGGAAGTCACACC  29223

EMBOSS_001     29251 TTCGGGAACGTGGTTGACCTACACAGGTGCCATCAAATTGGATGACAAAG  29300
                     ||||||||||||||||||||||||||||||||||||||||||||||||||
EMBOSS_001     29224 TTCGGGAACGTGGTTGACCTACACAGGTGCCATCAAATTGGATGACAAAG  29273

EMBOSS_001     29301 ATCCAAATTTCAAAGATCAAGTCATTTTGCTGAATAAGCATATTGACGCA  29350
                     ||||||||||||||||||||||||||||||||||||||||||||||||||
EMBOSS_001     29274 ATCCAAATTTCAAAGATCAAGTCATTTTGCTGAATAAGCATATTGACGCA  29323

EMBOSS_001     29351 TACAAAACATTCCCACCAACAGAGCCTAAAAAGGACAAAAAGAAGAAGGC  29400
                     ||||||||||||||||||||||||||||||||||||||||||||||||||
EMBOSS_001     29324 TACAAAACATTCCCACCAACAGAGCCTAAAAAGGACAAAAAGAAGAAGGC  29373

EMBOSS_001     29401 TGATGAAACTCAAGCCTTACCGCAGAGACAGAAGAAACAGCAAACTGTGA  29450
                     ||||||||||||||||||||||||||||||||||||||||||||||||||
EMBOSS_001     29374 TGATGAAACTCAAGCCTTACCGCAGAGACAGAAGAAACAGCAAACTGTGA  29423

EMBOSS_001     29451 CTCTTCTTCCTGCTGCAGATTTGGATGATTTCTCCAAACAATTGCAACAA  29500
                     ||||||||||||||||||||||||||||||||||||||||||||||||||
EMBOSS_001     29424 CTCTTCTTCCTGCTGCAGATTTGGATGATTTCTCCAAACAATTGCAACAA  29473

EMBOSS_001     29501 TCCATGAGCAGTGCTGACTCAACTCAGGCCTAAACTCATGCAGACCACAC  29550
                     ||||||||||||||||||||||||||||||||||||||||||||||||||
EMBOSS_001     29474 TCCATGAGCAGTGCTGACTCAACTCAGGCCTAAACTCATGCAGACCACAC  29523

EMBOSS_001     29551 AAGGCAGATGGGCTATATAAACGTTTTCGCTTTTCCGTTTACGATATATA  29600
                     ||||||||||||||||||||||||||||||||||||||||||||||||||
EMBOSS_001     29524 AAGGCAGATGGGCTATATAAACGTTTTCGCTTTTCCGTTTACGATATATA  29573

EMBOSS_001     29601 GTCTACTCTTGTGCAGAATGAATTCTCGTAACTACATAGCACAAGTAGAT  29650
                     ||||||||||||||||||||||||||||||||||||||||||||||||||
EMBOSS_001     29574 GTCTACTCTTGTGCAGAATGAATTCTCGTAACTACATAGCACAAGTAGAT  29623

EMBOSS_001     29651 GTAGTTAACTTTAATCTCACATAGCAATCTTTAATCAGTGTGTAACATTA  29700
                     ||||||||||||||||||||||||||||||||||||||||||||||||||
EMBOSS_001     29624 GTAGTTAACTTTAATCTCACATAGCAATCTTTAATCAGTGTGTAACATTA  29673

EMBOSS_001     29701 GGGAGGACTTGAAAGAGCCACCACATTTTCACCGAGGCCACGCGGAGTAC  29750
                     ||||||||||||||||||||||||||||||||||||||||||||||||||
EMBOSS_001     29674 GGGAGGACTTGAAAGAGCCACCACATTTTCACCGAGGCCACGCGGAGTAC  29723

EMBOSS_001     29751 GATCGAGTGTACAGTGAACAATGCTAGGGAGAGCTGCCTATATGGAAGAG  29800
                     ||||||||||||||||||||||||||||||||||||||||||||||||||
EMBOSS_001     29724 GATCGAGTGTACAGTGAACAATGCTAGGGAGAGCTGCCTATATGGAAGAG  29773

EMBOSS_001     29801 CCCTAATGTGTAAAATTAATTTTAGTAGTGCTATCCCCATGTGATTTTAA  29850
                     ||||||||||||||||||||||||||||||||||||||||||||||||||
EMBOSS_001     29774 CCCTAATGTGTAAAATTAATTTTAGTAGTGCTATCCCCATGTGATTTTAA  29823

EMBOSS_001     29851 TAGCTTCTTAGGAGAATGAC  29870
                     ||||||||||||||||||||
EMBOSS_001     29824 TAGCTTCTTAGGAGAATGAC  29843
```
