## Supplementary material for "Decoding the lethal effect of *SARS-CoV-2* (novel coronavirus) strains from global perspective: molecular pathogenesis and evolutionary divergence": Alignment file between a SARS-CoV strain and SARS-CoV-2 strain

**Supplementary Material 3**

**Alignment file between a SARS-CoV strain and SARS-CoV-2 strain**

```
# Aligned_sequences: 2
# 1: AY323977.2 (SARS-CoV Strain of Italy, 2003)
# 2: MT066156.1 (SARS-CoV Strain of Italy, 2020)
# Matrix: EDNAFULL
# Gap_penalty: 16
# Extend_penalty: 4
#
# Length: 29970
# Identity:   23790/29970 (79.4%)
# Similarity: 23791/29970 (79.4%)
# Gaps:         346/29970 (1.2%)
# Score: 93109
#
#
#=======================================

AY323977.2         1 ATATTAGGTTTTTACCTACCCAGGAAA--AGCCAACCAACCT-CGATCTC     47
                     ||...||||||.|||||.||||||.||  |.|||||||||.| |||||||
MT066156.1         1 ATTAAAGGTTTATACCTTCCCAGGTAACAAACCAACCAACTTTCGATCTC     50

AY323977.2        48 TTGTAGATCTGTTCTCTAAACGAACTTTAAAATCTGTGTAGCTGTCGCTC     97
                     |||||||||||||||||||||||||||||||||||||||.||||||.|||
MT066156.1        51 TTGTAGATCTGTTCTCTAAACGAACTTTAAAATCTGTGTGGCTGTCACTC    100

AY323977.2        98 GGCTGCATGCCTAGTGCACCTACGCAGTATAAACAATAATAAATTTTACT    147
                     ||||||||||.||||||||..|||||||||||..|||||..|||  ||||
MT066156.1       101 GGCTGCATGCTTAGTGCACTCACGCAGTATAATTAATAACTAAT--TACT    148

AY323977.2       148 GTCGTTGACAAGAAACGAGTAACTCGTCCCTCTTCTGCAGACTGCTTACG    197
                     ||||||||||.||.||||||||||||||..||||||||||.|||||||||
MT066156.1       149 GTCGTTGACAGGACACGAGTAACTCGTCTATCTTCTGCAGGCTGCTTACG    198

AY323977.2       198 GTTTCGTCCGTGTTGCAGTCGATCATCAGCATACCTAGGTTTCGTCCGGG    247
                     ||||||||||||||||||.||||||||||||.|.||||||||||||||||
MT066156.1       199 GTTTCGTCCGTGTTGCAGCCGATCATCAGCACATCTAGGTTTCGTCCGGG    248

AY323977.2       248 TGTGACCGAAAGGTAAGATGGAGAGCCTTGTTCTTGGTGTCAACGAGAAA    297
                     |||||||||||||||||||||||||||||||.|.||||.|||||||||||
MT066156.1       249 TGTGACCGAAAGGTAAGATGGAGAGCCTTGTCCCTGGTTTCAACGAGAAA    298

AY323977.2       298 ACACACGTCCAACTCAGTTTGCCTGTCCTTCAGGTTAGAGACGTGCTAGT    347
                     ||||||||||||||||||||||||||..|.||||||.|.||||||||.||
MT066156.1       299 ACACACGTCCAACTCAGTTTGCCTGTTTTACAGGTTCGCGACGTGCTCGT    348

AY323977.2       348 GCGTGGCTTCGGGGACTCTGTGGAAGAGGCCCTATCGGAGGCACGTGAAC    397
                     .||||||||.||.|||||.|||||.||||.|.||||.|||||||||.|||
MT066156.1       349 ACGTGGCTTTGGAGACTCCGTGGAGGAGGTCTTATCAGAGGCACGTCAAC    398

AY323977.2       398 ACCTCAAAAATGGCACTTGTGGTCTAGTAGAGCTGGAAAAAGGCGTACTG    447
                     |.||.|||.|||||||||||||..|||||||..|.|||||||||||..||
MT066156.1       399 ATCTTAAAGATGGCACTTGTGGCTTAGTAGAAGTTGAAAAAGGCGTTTTG    448

AY323977.2       448 CCCCAGCTTGAACAGCCCTATGTGTTCATTAAACGTTCTGATGCCTTAA-    496
                     ||.||.|||||||||||||||||||||||.||||||||.|||||...||
MT066156.1       449 CCTCAACTTGAACAGCCCTATGTGTTCATCAAACGTTCGGATGCTCGAAC    498

AY323977.2       497 -GCACCAATCACGGCCACAAGGTCGTTGAGCTGGTTGCAGAAATGGACGG    545
                      |||||  |||.||.||.....|.|||||||||||.||||||.|.||.||
MT066156.1       499 TGCACC--TCATGGTCATGTTATGGTTGAGCTGGTAGCAGAACTCGAAGG    546

AY323977.2       546 CATTCAGTACGGTCGTAGCGGTATAACACTGGGAGTACTCGTGCCACATG    595
                     ||||||||||||||||||.|||...|||||.||.||.||.||.||.||||
MT066156.1       547 CATTCAGTACGGTCGTAGTGGTGAGACACTTGGTGTCCTTGTCCCTCATG    596

AY323977.2       596 TGGGCGAAACCCCAATTGCATACCGCAATGTTCTTCTTCGTAAGAACGGT    645
                     |||||||||..|||.|.||.||||||||.|||||||||||||||||||||
MT066156.1       597 TGGGCGAAATACCAGTGGCTTACCGCAAGGTTCTTCTTCGTAAGAACGGT    646

AY323977.2       646 AATAAGGGAGCCGGTGGTCATAGCTATGGCATCGATCTAAAGTCTTATGA    695
                     |||||.|||||.|||||.|||||.||.|||..||||||||||||.|.|||
MT066156.1       647 AATAAAGGAGCTGGTGGCCATAGTTACGGCGCCGATCTAAAGTCATTTGA    696

AY323977.2       696 CTTAGGTGACGAGCTTGGCACTGATCCCATTGAAGATTATGAACAAAACT    745
                     ||||||.||||||||||||||||||||...||||||||.|.||.||||||
MT066156.1       697 CTTAGGCGACGAGCTTGGCACTGATCCTTATGAAGATTTTCAAGAAAACT    746

AY323977.2       746 GGAACACTAAGCATGGCAGTGGTGCACTCCGTGAACTCACTCGTGAGCTC    795
                     ||||||||||.|||.|||||||||....|||||||||||..||||||||.
MT066156.1       747 GGAACACTAAACATAGCAGTGGTGTTACCCGTGAACTCATGCGTGAGCTT    796

AY323977.2       796 AATGGAGGTGCAGTCACTCGCTATGTCGACAACAATTTCTGTGGCCCAGA    845
                     ||.|||||.|||..|||||||||||||||.|||||.|||||||||||.||
MT066156.1       797 AACGGAGGGGCATACACTCGCTATGTCGATAACAACTTCTGTGGCCCTGA    846

AY323977.2       846 TGGGTACCCTCTTGATTGCATCAAAGATTTTCTCGCACGCGCGGGCAAGT    895
                     |||.|||||||||||.|||||.|||||..||||.|||||.||.||.||..
MT066156.1       847 TGGCTACCCTCTTGAGTGCATTAAAGACCTTCTAGCACGTGCTGGTAAAG    896

AY323977.2       896 CAATGTGCACTCTTTCCGAACAACTTGATTACATCGAGTCGAAGAGAGGT    945
                     |....||||||.|.|||||||||||.||.|..||.||..|.|||||.|||
MT066156.1       897 CTTCATGCACTTTGTCCGAACAACTGGACTTTATTGACACTAAGAGGGGT    946

AY323977.2       946 GTCTACTGCTGCCGTGACCATGAGCATGAAATTGCCTGGTTCACTGAGCG    995
                     ||.||||||||||||||.|||||||||||||||||.||||.|||.||.||
MT066156.1       947 GTATACTGCTGCCGTGAACATGAGCATGAAATTGCTTGGTACACGGAACG    996

AY323977.2       996 CTCTGATAAGAGCTACGAGCACCAGACACCCTTCGAAATTAAGAGTGCCA   1045
                     .|||||.||||||||.||....||||||||.||.||||||||....||.|
MT066156.1       997 TTCTGAAAAGAGCTATGAATTGCAGACACCTTTTGAAATTAAATTGGCAA   1046

AY323977.2      1046 AGAAATTTGACACTTTCAAAGGGGAATGCCCAAAGTTTGTGTTTCCTCTT   1095
                     |||||||||||||.|||||.||||||||.|||||.|||||.|||||..|.
MT066156.1      1047 AGAAATTTGACACCTTCAATGGGGAATGTCCAAATTTTGTATTTCCCTTA   1096

AY323977.2      1096 AACTCAAAAGTCAAAGTCATTCAACCACGTGTTGAAAAGAAAAAGACTGA   1145
                     ||.||.|.|.||||....|||||||||.|.|||||||||||||||..|||
MT066156.1      1097 AATTCCATAATCAAGACTATTCAACCAAGGGTTGAAAAGAAAAAGCTTGA   1146

AY323977.2      1146 GGGTTTCATGGGGCGTATACGCTCTGTGTACCCTGTTGCATCTCCACAGG   1195
                     .||.||.|||||..|.||.||.|||||.||.||.|||||.||.|||.|.|
MT066156.1      1147 TGGCTTTATGGGTAGAATTCGATCTGTCTATCCAGTTGCGTCACCAAATG   1196

AY323977.2      1196 AGTGTAACAATATGCACTTGTCTACCTTGATGAAATGTAATCATTGCGAT   1245
                     |.||.|||.|.|||..|.|.||.||..|.|||||.|||.|||||||.|.|
MT066156.1      1197 AATGCAACCAAATGTGCCTTTCAACTCTCATGAAGTGTGATCATTGTGGT   1246

AY323977.2      1246 GAAGTTTCATGGCAGACGTGCGACTTTCTGAAAGCCACTTGTGAACATTG   1295
                     |||..|||||||||||||.||||.|||.|.|||||||||||.|||..|||
MT066156.1      1247 GAAACTTCATGGCAGACGGGCGATTTTGTTAAAGCCACTTGCGAATTTTG   1296

AY323977.2      1296 TGGCACTGAAAATTTAGTTATTGAAGGACCTACTACATGTGGGTACCTAC   1345
                     |||||||||.|||||...||..|||||..|.|||||.|||||.|||.|||
MT066156.1      1297 TGGCACTGAGAATTTGACTAAAGAAGGTGCCACTACTTGTGGTTACTTAC   1346

AY323977.2      1346 CTACTAATGCTGTAGTGAAAATGCCATGTCCTGCCTGTCAAGACCCAGAG   1395
                     |....||||||||.||.|||||....|||||.||.|||||..|..||||.
MT066156.1      1347 CCCAAAATGCTGTTGTTAAAATTTATTGTCCAGCATGTCACAATTCAGAA   1396

AY323977.2      1396 ATTGGACCTGAGCATAGTGTTGCAGATTATCACAACCACTCAAACATTGA   1445
                     .|.|||||||||||||||.||||.||.||.||.||..|.||...|.|..|
MT066156.1      1397 GTAGGACCTGAGCATAGTCTTGCCGAATACCATAATGAATCTGGCTTGAA   1446

AY323977.2      1446 AACTCGACTCCGCAAGGGAGGTAGGACTAGATGTTTTGGAGGCTGTGTGT   1495
                     |||....||.||.|||||.|||.|.||||.....||||||||||||||||
MT066156.1      1447 AACCATTCTTCGTAAGGGTGGTCGCACTATTGCCTTTGGAGGCTGTGTGT   1496

AY323977.2      1496 TTGCCTATGTTGGCTGCTATAATAAGCGTGCCTACTGGGTTCCTCGTGCT   1545
                     |..|.||||||||.|||.||||.|||.|||||||.||||||||.||||||
MT066156.1      1497 TCTCTTATGTTGGTTGCCATAACAAGTGTGCCTATTGGGTTCCACGTGCT   1546

AY323977.2      1546 AGTGCTGATATTGGCTCAGGCCATACTGGCATTACTGGTGACAATGTGGA   1595
                     ||.|||.|.||.||.|....||||||.||..||..|||.||...|...||
MT066156.1      1547 AGCGCTAACATAGGTTGTAACCATACAGGTGTTGTTGGAGAAGGTTCCGA   1596

AY323977.2      1596 GACCTTGAATGAGGATCTCCTTGAGATACTGAGTCGTGAACGTGTTAACA   1645
                     .....|.|||||..|.||.|||||.|||||.......||....||.||||
MT066156.1      1597 AGGTCTTAATGACAACCTTCTTGAAATACTCCAAAAAGAGAAAGTCAACA   1646

AY323977.2      1646 TTAACATTGTTGGCGATTTTCATTTGAATGAAGAGGTTGCCATCATTTTG   1695
                     |.||.||||||||.||.|||.|..|.|||||||||.|.|||||.||||||
MT066156.1      1647 TCAATATTGTTGGTGACTTTAAACTTAATGAAGAGATCGCCATTATTTTG   1696

AY323977.2      1696 GCATCTTTCTCTGCTTCTACAAGTGCCTTTATTGACACTATAAAGAGTCT   1745
                     ||||||||.||||||||.||||||||.|||.|.||.|||.|.||..||.|
MT066156.1      1697 GCATCTTTTTCTGCTTCCACAAGTGCTTTTGTGGAAACTGTGAAAGGTTT   1746

AY323977.2      1746 TGATTACAAGTCTTTCAAAACCATTGTTGAGTCCTGCGGTAACTATAAAG   1795
                     .|||||.||..|.||||||...||||||||.|||||.|||||.|.|||||
MT066156.1      1747 GGATTATAAAGCATTCAAACAAATTGTTGAATCCTGTGGTAATTTTAAAG   1796

AY323977.2      1796 TTACCAAGGGAAAGCCCGTAAAAGGTGCTTGGAACATTGGACAACAGAGA   1845
                     ||||.||.|||||..|...|||||||||.|||||.|||||..||||||.|
MT066156.1      1797 TTACAAAAGGAAAAGCTAAAAAAGGTGCCTGGAATATTGGTGAACAGAAA   1846

AY323977.2      1846 TCAGTTTTAACACCACTGTGTGGTTTTCCCTCACAGGCTGCTGGTGTTAT   1895
                     |||.|..|.|..||.||.|.||..|||.|.|||.||||||||.|||||.|
MT066156.1      1847 TCAATACTGAGTCCTCTTTATGCATTTGCATCAGAGGCTGCTCGTGTTGT   1896

AY323977.2      1896 CAGATCAATTTTTGCGCGCACACTTGATGCAGCAAACCACTCAATTCCTG   1945
                     ..||||||||||..|.|||||.|||||..|.||..|..|.||..|.|.||
MT066156.1      1897 ACGATCAATTTTCTCCCGCACTCTTGAAACTGCTCAAAATTCTGTGCGTG   1946

AY323977.2      1946 ATTTGCAAAGAGCAGCTGTCACCATACTTGATGGTATTTCTGAACAGTCA   1995
                     .|||.||.|..||.|||.|.||.|||||.|||||.|||||..|..|.|||
MT066156.1      1947 TTTTACAGAAGGCCGCTATAACAATACTAGATGGAATTTCACAGTATTCA   1996

AY323977.2      1996 TTACGTCTTGTCGACGCCATGGTTTATACTTCAGACCTGCTCACCAACAG   2045
                     .|..|.||..|.||.||.|||.|.|..||.||.||..||...||.||||.
MT066156.1      1997 CTGAGACTCATTGATGCTATGATGTTCACATCTGATTTGGCTACTAACAA   2046

AY323977.2      2046 TGTCATTATTATGGCATATGTAACTGGTGGTCTTGTACAACAGACTTCTC   2095
                     |.|..||.|.|||||.||..|.||.||||||.||||.||...||||||.|
MT066156.1      2047 TCTAGTTGTAATGGCCTACATTACAGGTGGTGTTGTTCAGTTGACTTCGC   2096

AY323977.2      2096 AGTGGTTGTCTAATCTTTTGGGCACTACTGTTGAAAAACTCAGGCCTATC   2145
                     |||||.|..||||..|.||.||||||..|..|||||||||||..||..||
MT066156.1      2097 AGTGGCTAACTAACATCTTTGGCACTGTTTATGAAAAACTCAAACCCGTC   2146

AY323977.2      2146 TTTGAATGGATTGAGGCGAAACTTAGTGCAGGAGTTGAATTTCTCAAGGA   2195
                     .||||.|||.||||.|.|||..|||..|.|||.||.||.|||||.|..||
MT066156.1      2147 CTTGATTGGCTTGAAGAGAAGTTTAAGGAAGGTGTAGAGTTTCTTAGAGA   2196

AY323977.2      2196 TGCTTGGGAGATTCTCAAATTTCTCATTACAGGTGTTTTTGACATCGTCA   2245
                     .|.||||||.|||.|.||||||.||...||..|||.||.|||.||.|||.
MT066156.1      2197 CGGTTGGGAAATTGTTAAATTTATCTCAACCTGTGCTTGTGAAATTGTCG   2246

AY323977.2      2246 AGGGTCAAATACAGGTTGCTTCAGATAAC---ATCAAGGATTGTGTAAAA   2292
                     ..||.|||||.   ||..|.|..|.|||.   ||.|||||..||||..|.
MT066156.1      2247 GTGGACAAATT---GTCACCTGTGCTAAGGAAATTAAGGAGAGTGTTCAG   2293

AY323977.2      2293 TGCTTCATTGATGTTGTTAACAAGGCACTCGAAATGTGCATTGATCAAGT   2342
                     ...|||.||.|..||||.||.||.....|.|...||||...|||.....|
MT066156.1      2294 ACATTCTTTAAGCTTGTAAATAAATTTTTGGCTTTGTGTGCTGACTCTAT   2343

AY323977.2      2343 CACTATCGCTGGCGCAAAGTTGCGATCACTCAACTTAGGTGAAGTCTTCA   2392
                     ||.|||.|.|||.||.||..|...|.|..|.||.|||||||||...||..
MT066156.1      2344 CATTATTGGTGGAGCTAAACTTAAAGCCTTGAATTTAGGTGAAACATTTG   2393

AY323977.2      2393 TCGCTCAAAGCAAGGGACTTTACCGTCAGTGTATACGTGGCA-AGGAGCA   2441
                     ||.|.||....||||||.|.|||.|..|||||.|......|| ||.||.|
MT066156.1      2394 TCACGCACTCAAAGGGATTGTACAGAAAGTGTGTTAAATCCAGAGAAGAA   2443

AY323977.2      2442 GCTGCAACTACTCATGCCTCTTAAGGCACCAAAAGAAGTAACCTTTCTTG   2491
                     .|||.. ||||||||||||||.||.||.|||||||||.|.|.|||..|.|
MT066156.1      2444 ACTGGC-CTACTCATGCCTCTAAAAGCCCCAAAAGAAATTATCTTCTTAG   2492

AY323977.2      2492 AAGGTGATTCACATGACACAGTACTTACCTCTGAGGAGGTTGTTCTCAAG   2541
                     |.||.||..|||.|..|||||.|.|.....|.|||||.|||||..|.||.
MT066156.1      2493 AGGGAGAAACACTTCCCACAGAAGTGTTAACAGAGGAAGTTGTCTTGAAA   2542

AY323977.2      2542 AACGGTGAACTCGAAGCACTCGAGACGCCCGTTGATAGCTTCACAAATGG   2591
                     |..|||||..|..||.||.|.|| ||..||  |..|||.....|...||.
MT066156.1      2543 ACTGGTGATTTACAACCATTAGA-ACAACC--TACTAGTGAAGCTGTTGA   2589

AY323977.2      2592 AGCTA---TCGTTGGCACACCAGTCTGTGTAAATGGCCTCATGCTCTTAG   2638
                     ||||.   |.|||||.||||||||.|||.|.||.||.||.|||.|..|.|
MT066156.1      2590 AGCTCCATTGGTTGGTACACCAGTTTGTATTAACGGGCTTATGTTGCTCG   2639

AY323977.2      2639 AGATTAAGGACAAAGAACAATACTGCGCATTGTCTCCTGGTTTACTGGCT   2688
                     |.||.||.||||.||||.|.|||||.||..|..|.|||..|.|..|||..
MT066156.1      2640 AAATCAAAGACACAGAAAAGTACTGTGCCCTTGCACCTAATATGATGGTA   2689

AY323977.2      2689 ACAAACAATGTCTTTCGCTTAAAAGGGGGTGCACCAATTAAAGGTGTAAC   2738
                     |||||||||..|||.....|.|||||.||||||||||. ||||||.  ||
MT066156.1      2690 ACAAACAATACCTTCACACTCAAAGGCGGTGCACCAAC-AAAGGTT--AC   2736

AY323977.2      2739 CTTTGGAGAAGATACTGTTTGGGAAGTTCAAGGTTACAAGAATGTGAGAA   2788
                     .|||||.||.||.|||||....|||||.|||||||||||||.|||||..|
MT066156.1      2737 TTTTGGTGATGACACTGTGATAGAAGTGCAAGGTTACAAGAGTGTGAATA   2786

AY323977.2      2789 TCACATTTGAGCTTGATGAACGTGTTGACAAAGTGCTTAATGAAAAGTGC   2838
                     ||||.|||||.|||||||||.|..||||.|||||.||||||||.||||||
MT066156.1      2787 TCACTTTTGAACTTGATGAAAGGATTGATAAAGTACTTAATGAGAAGTGC   2836

AY323977.2      2839 TCTGTCTACACTGTTGAATCCGGTACCGAAGTTACTGAGTTTGCATGTGT   2888
                     ||||.|||.||.||||||..||||||.|||||.|.||||||.||.|||||
MT066156.1      2837 TCTGCCTATACAGTTGAACTCGGTACAGAAGTAAATGAGTTCGCCTGTGT   2886

AY323977.2      2889 TGTAGCAGAGGCTGTTGTGAAGACTTTACAACCAGTTTCTGATCTCCTTA   2938
                     |||.|||||.|||||..|.||.|||||.||||||||.|||||..|.||||
MT066156.1      2887 TGTGGCAGATGCTGTCATAAAAACTTTGCAACCAGTATCTGAATTACTTA   2936

AY323977.2      2939 CCAACATGGGTATTGATCTTGATGAGTGGAGTGTAGCTACATTCTACTTA   2988
                     |.....||||.||||||.|.||||||||||||.|.|||||||.|||||||
MT066156.1      2937 CACCACTGGGCATTGATTTAGATGAGTGGAGTATGGCTACATACTACTTA   2986

AY323977.2      2989 TTTGATGATGCTGGTGAAGAAAACTTTTCATCACGTATGTATTGTTCCTT   3038
                     ||||||||..|||||||....||.||..|.||||.||||||||||||.||
MT066156.1      2987 TTTGATGAGTCTGGTGAGTTTAAATTGGCTTCACATATGTATTGTTCTTT   3036

AY323977.2      3039 TTACCCTCCAGATGAGGAAGAAGAGGACGATGCAGAGTGTGAGGAAGAAG   3088
                     .|||||||||||||||||.|||||.||.|.|   ||.|||||.|||||||
MT066156.1      3037 CTACCCTCCAGATGAGGATGAAGAAGAAGGT---GATTGTGAAGAAGAAG   3083

AY323977.2      3089 AAATTGATGAAACCTGTGAACATGAGTACGGTACAGAGGATGATTATCAA   3138
                     |..||||...|.|...|.||.|||||||.|||||.||.||||||||.|||
MT066156.1      3084 AGTTTGAGCCATCAACTCAATATGAGTATGGTACTGAAGATGATTACCAA   3133

AY323977.2      3139 GGTCTCCCTCTGGAATTTGGTGCCTCAGCTGAAACAGTTCGAGTTGAGGA   3188
                     |||...|||.||||||||||||||.|..|||...|..|||.|..|||.||
MT066156.1      3134 GGTAAACCTTTGGAATTTGGTGCCACTTCTGCTGCTCTTCAACCTGAAGA   3183

AY323977.2      3189 AGAAGAAGAGGAAGACTGGCTGGATGAT------------ACT-------   3219
                     |||..||||.|||||.|||.|.||||||            |||
MT066156.1      3184 AGAGCAAGAAGAAGATTGGTTAGATGATGATAGTCAACAAACTGTTGGTC   3233

AY323977.2      3220 -----------ACTGAG--CAATCAGAGA-------------------TT   3237
                                |.||||  ||||||||.|                   ||
MT066156.1      3234 AACAAGACGGCAGTGAGGACAATCAGACAACTACTATTCAAACAATTGTT   3283

AY323977.2      3238 GAGCCAGAACCA------------GAACCTACACCTGAAGAACC------   3269
                     |||....||||.            ||||.||||||.|..|..|.
MT066156.1      3284 GAGGTTCAACCTCAATTAGAGATGGAACTTACACCAGTTGTTCAGACTAT   3333

AY323977.2      3270 ---AGTTAATCAGTTTACTGGTTATTTAAAACTTACTGACAATGTTGCCA   3316
                        |||.|||...||||.|||||||||||||||||||||||||||...||
MT066156.1      3334 TGAAGTGAATAGTTTTAGTGGTTATTTAAAACTTACTGACAATGTATACA   3383

AY323977.2      3317 TTAAATGTGTTGACATCGTTAAGGAGGCACAAAGTGCTAATCCTATGGTG   3366
                     |||||..||..|||||.||..|.||.||..|||..|..||.||.|..|||
MT066156.1      3384 TTAAAAATGCAGACATTGTGGAAGAAGCTAAAAAGGTAAAACCAACAGTG   3433

AY323977.2      3367 ATTGTAAATGCTGCTAACATACACCTGAAACATGGTGGTGGTGTAGCAGG   3416
                     .||||.|||||.||.||..|..||||.||||||||.||.|||||.|||||
MT066156.1      3434 GTTGTTAATGCAGCCAATGTTTACCTTAAACATGGAGGAGGTGTTGCAGG   3483

AY323977.2      3417 TGCACTCAACAAGGCAACCAATGGTGCCATGCAAAAGGAGAGTGATGATT   3466
                     .||..|.||.|||||.||.||...||||||||||...||...||||||||
MT066156.1      3484 AGCCTTAAATAAGGCTACTAACAATGCCATGCAAGTTGAATCTGATGATT   3533

AY323977.2      3467 ACATTAAGCTA--AATGGCCCTCTTACAGTAGGAGGGTCTTGTTTGCTTT   3514
                     ||||  |||||  |||||.||.||||.|||.||.||...||||.|..|..
MT066156.1      3534 ACAT--AGCTACTAATGGACCACTTAAAGTGGGTGGTAGTTGTGTTTTAA   3581

AY323977.2      3515 CTGGACATAATCTTGCTAAGAAGTGTCTGCATGTTGTTGGACCTAACCTA   3564
                     ..|||||.|||||||||||..|.|||||.||||||||.||.||.||..|.
MT066156.1      3582 GCGGACACAATCTTGCTAAACACTGTCTTCATGTTGTCGGCCCAAATGTT   3631

AY323977.2      3565 AATGCAGGTGAGGACATCCAGCTTCTTAAGGCAGCATATGAAAATTTCAA   3614
                     ||...||||||.|||||.||.|||||||||...||.|||||||||||.||
MT066156.1      3632 AACAAAGGTGAAGACATTCAACTTCTTAAGAGTGCTTATGAAAATTTTAA   3681

AY323977.2      3615 TTCACAGGACATCTTACTTGCACCATTGTTGTCAGCAGGCATATTTGGTG   3664
                     |...||.||..|..|||||||||||||.||.|||||.||.||.|||||||
MT066156.1      3682 TCAGCACGAAGTTCTACTTGCACCATTATTATCAGCTGGTATTTTTGGTG   3731

AY323977.2      3665 CTAAACCACTTCAGTCTTTACAAGTGTGCGTGCAGACGGTTCGTACACAG   3714
                     ||.|.||..|.||.||||||..|||.||.||..|.||.|||||.|||.|.
MT066156.1      3732 CTGACCCTATACATTCTTTAAGAGTTTGTGTAGATACTGTTCGCACAAAT   3781

AY323977.2      3715 GTTTATATTGCAGTCAATGACAAAGCTCTTTATGAGCAGGTTGTCATGGA   3764
                     ||.||..|.||.|||..|||.|||..|||.|||||..|..||||. |..|
MT066156.1      3782 GTCTACTTAGCTGTCTTTGATAAAAATCTCTATGACAAACTTGTT-TCAA   3830

AY323977.2      3765 TTATCTTGATAACCTGAAGCCTAGAGTGGAAGC--ACCTAAACAA-GAGG   3811
                     ...|.|||..||  |||||..|..|..|.|||.  |.|.|||.|. |..|
MT066156.1      3831 GCTTTTTGGAAA--TGAAGAGTGAAAAGCAAGTTGAACAAAAGATCGCTG   3878

AY323977.2      3812 AGCCACCAAACACAGAAGAT------TCCAAAACTGAGGAGAAATCT---   3852
                     ||...||.||....||||.|      |..|.||||||....|||.||
MT066156.1      3879 AGATTCCTAAAGAGGAAGTTAAGCCATTTATAACTGAAAGTAAACCTTCA   3928

AY323977.2      3853 GTCGTACAGAAGCCTGTCGATGTGAAGCCAAAAATTAAGGCCTGCATTGA   3902
                     ||.|.|||||........||||..|||   |||||.||.||.||..||||
MT066156.1      3929 GTTGAACAGAGAAAACAAGATGATAAG---AAAATCAAAGCTTGTGTTGA   3975

AY323977.2      3903 TGAGGTTACCACAACACTGGAAGAAACTAAGTTTCTTACCAATAAGTTAC   3952
                     .||.|||||.|||||.|||||||||||||||||.||.||..|.||.||..
MT066156.1      3976 AGAAGTTACAACAACTCTGGAAGAAACTAAGTTCCTCACAGAAAACTTGT   4025

AY323977.2      3953 TCTTGTTTGCTGATATCAATGGTAAGCTTTACCATGATTCTCAGAACATG   4002
                     |..|.|.|..|||.||.|||||.||.|||.|.|..||||||...|...|.
MT066156.1      4026 TACTTTATATTGACATTAATGGCAATCTTCATCCAGATTCTGCCACTCTT   4075

AY323977.2      4003 CTTAGAGGTGAAGATATGTCTTTCCTTGAGAAGGATGCACCTTACATGGT   4052
                     .||||.|.....||.||..|||||.|..||||.|||||.||.||.||.||
MT066156.1      4076 GTTAGTGACATTGACATCACTTTCTTAAAGAAAGATGCTCCATATATAGT   4125

AY323977.2      4053 AGGTGATGTTAT-CACTAGTGGTGATATCACTTGTGTTGTAATACCCTCC   4101
                     .|||||||||.| ||..|| ||||.|.|.|||..|||.||.|||||..|.
MT066156.1      4126 GGGTGATGTTGTTCAAGAG-GGTGTTTTAACTGCTGTGGTTATACCTACT   4174

AY323977.2      4102 AAAAAGGCTGGTGGCACTACTGAGATGCTCTCAAGAGCTTTGAAGAAAGT   4151
                     |||||||||||||||||||||||.|||||..|.|.||||||||..|||||
MT066156.1      4175 AAAAAGGCTGGTGGCACTACTGAAATGCTAGCGAAAGCTTTGAGAAAAGT   4224

AY323977.2      4152 GCCAGTTGATGAGTATATAACCACGTACCCTGGACAAGGATGTGCTGGTT   4201
                     ||||...||..|.|||||||||||.|||||.||.||.||.|....|||||
MT066156.1      4225 GCCAACAGACAATTATATAACCACTTACCCGGGTCAGGGTTTAAATGGTT   4274

AY323977.2      4202 ATACACTTGAGGAAGCTAAGACTGCTCTTAAGAAATGCAAATCTGCATTT   4251
                     |.||..|.|||||.||.|||||.|..|||||.||.||.|||..|||.|||
MT066156.1      4275 ACACTGTAGAGGAGGCAAAGACAGTGCTTAAAAAGTGTAAAAGTGCCTTT   4324

AY323977.2      4252 TATGTACTACCTTCAGAAGCACCTAATGCTAAGGAAGAGATTCTAGGAAC   4301
                     ||..|.|||||.||........||||||..|||.||||.|||||.|||||
MT066156.1      4325 TACATTCTACCATCTATTATCTCTAATGAGAAGCAAGAAATTCTTGGAAC   4374

AY323977.2      4302 TGTATCCTGGAATTTGAGAGAAATGCTTGCTCATGCTGAAGAGACAAGAA   4351
                     |||.||.|||||||||.|||||||||||||.|||||.|||||.|||.|.|
MT066156.1      4375 TGTTTCTTGGAATTTGCGAGAAATGCTTGCACATGCAGAAGAAACACGCA   4424

AY323977.2      4352 AATTAATGCCTATATGCATGGATGTTAGAGCCATAATGGCAACCATCCAA   4401
                     |||||||||||.|.||..||||...||.|||||||.|..||||.||.||.
MT066156.1      4425 AATTAATGCCTGTCTGTGTGGAAACTAAAGCCATAGTTTCAACTATACAG   4474

AY323977.2      4402 CGTAAGTATAAAGGAATTAAAATTCAAGAGGGCATCGTTGACTATGGTGT   4451
                     |||||.|||||.||.||||||||.||||||||..|.|||||.|||||||.
MT066156.1      4475 CGTAAATATAAGGGTATTAAAATACAAGAGGGTGTGGTTGATTATGGTGC   4524

AY323977.2      4452 CCGATTCTTCTTTTATACTAGTAAAGAGCCTGTAGCTTCTATTATTACGA   4501
                     ..||||.|.||||||.||.||||||....|||||||.||..||||.|..|
MT066156.1      4525 TAGATTTTACTTTTACACCAGTAAAACAACTGTAGCGTCACTTATCAACA   4574

AY323977.2      4502 AGCTGAACTCTCTAAATGAGCCGCTTGTCACAATGCCAATTGGTTATGTG   4551
                     ..||.|||..|||||||||..|.|||||.|||||||||.||||.|||||.
MT066156.1      4575 CACTTAACGATCTAAATGAAACTCTTGTTACAATGCCACTTGGCTATGTA   4624

AY323977.2      4552 ACACATGGTTTTAATCTTGAAGAGGCTGCGCGCTGTATGCGTTCTCTTAA   4601
                     ||||||||.||.|||.|.|||||.|||||.||.|.||||.|.|||||.||
MT066156.1      4625 ACACATGGCTTAAATTTGGAAGAAGCTGCTCGGTATATGAGATCTCTCAA   4674

AY323977.2      4602 AGCTCCTGCCGTAGTGTCAGTATCATCACCAGATGCTGTTACTACATATA   4651
                     ||..||.||...|||.||.||.||.|||||.|||||||||||..|.||||
MT066156.1      4675 AGTGCCAGCTACAGTTTCTGTTTCTTCACCTGATGCTGTTACAGCGTATA   4724

AY323977.2      4652 ATGGATACCTCACTTCGTCATCAAAGACATCTGAGGAGCACTTTGTAGAA   4701
                     ||||.||.||.|||||.||.||.||.|||.||||.||.||.|||.|.|||
MT066156.1      4725 ATGGTTATCTTACTTCTTCTTCTAAAACACCTGAAGAACATTTTATTGAA   4774

AY323977.2      4702 ACAGTTTCTTTGGCTGGCTCTTACAGAGATTGGTCCTATTCAGGACAGCG   4751
                     ||..|.||..|.|||||.||.||.|.|||||||||||||||.|||||...
MT066156.1      4775 ACCATCTCACTTGCTGGTTCCTATAAAGATTGGTCCTATTCTGGACAATC   4824

AY323977.2      4752 TACAGAGTTAGGTGTTGAATTTCTTAAGCGTGGTGACAAAATTGTGTACC   4801
                     ||||.|..|||||.|.||||||||||||.|.|||||.||||.|||.||..
MT066156.1      4825 TACACAACTAGGTATAGAATTTCTTAAGAGAGGTGATAAAAGTGTATATT   4874

AY323977.2      4802 ACACTCTGGAGAGCCCCGTCGAGTTTC-ATCTTGACGGTGAGGTTCTTTC   4850
                     |||||    ||....||..|.|..||| |.||.||.|||||.|||.|..|
MT066156.1      4875 ACACT----AGTAATCCTACCACATTCCACCTAGATGGTGAAGTTATCAC   4920

AY323977.2      4851 ACTTGACAAACTAAAGAGTCTCTTATCCCTGCGGGAGGTTAAGACTATAA   4900
                     ..|||||||.||.||||..||..|.||..||.|.||.||.|.||||||.|
MT066156.1      4921 CTTTGACAATCTTAAGACACTTCTTTCTTTGAGAGAAGTGAGGACTATTA   4970

AY323977.2      4901 AAGTGTTCACAACTGTGGACAACACTAATCTCCACACACAGCTTGTGGAT   4950
                     |.|||||.|||||.||.|||||||.|||.||||||||.||..|||||||.
MT066156.1      4971 AGGTGTTTACAACAGTAGACAACATTAACCTCCACACGCAAGTTGTGGAC   5020

AY323977.2      4951 ATGTCTATGACATATGGACAGCAGTTTGGTCCAACATACTTGGATGGTGC   5000
                     |||||.||||||||||||||.||||||||||||||.||.||||||||.||
MT066156.1      5021 ATGTCAATGACATATGGACAACAGTTTGGTCCAACTTATTTGGATGGAGC   5070

AY323977.2      5001 TGATGTTACAAAAATTAAACCTCATGTAAATCATGAGGGTAAGACTTTCT   5050
                     |||||||||.|||||.|||||||||......|||||.|||||.||.||.|
MT066156.1      5071 TGATGTTACTAAAATAAAACCTCATAATTCACATGAAGGTAAAACATTTT   5120

AY323977.2      5051 TTGTACTACCTAGTGATGACACACTACGTAGTGAAGCTTTCGAGTACTAC   5100
                     .|||..||||||.|||||||||.||||||..|||.|||||.|||||||||
MT066156.1      5121 ATGTTTTACCTAATGATGACACTCTACGTGTTGAGGCTTTTGAGTACTAC   5170

AY323977.2      5101 CATACTCTTGATGAGAGTTTTCTTGGTAGGTACATGTCTGCTTTAAACCA   5150
                     ||.||...||||...||||||||.||||||||||||||.||.|||||.||
MT066156.1      5171 CACACAACTGATCCTAGTTTTCTGGGTAGGTACATGTCAGCATTAAATCA   5220

AY323977.2      5151 CACAAAGAAATGGAAATTTCCTCAAGTTGGTGGTTTAACTTCAATTAAAT   5200
                     |||.||.||.|||||||..||.||||||..||||||||||||.|||||||
MT066156.1      5221 CACTAAAAAGTGGAAATACCCACAAGTTAATGGTTTAACTTCTATTAAAT   5270

AY323977.2      5201 GGGCTGATAACAATTGTTATTTGTCTAGTGTTTTATTAGCACTTCAACAG   5250
                     ||||.||||||||.||||||.|..|.|.||..||.|||.||||.|||||.
MT066156.1      5271 GGGCAGATAACAACTGTTATCTTGCCACTGCATTGTTAACACTCCAACAA   5320

AY323977.2      5251 CTTGAAGTCAAATTCAATGCACCAGCACTTCAAGAGGCTTATTATAGAGC   5300
                     .|.||..|.||.||.|||.||||.||.||.|||||.||||||||.|||||
MT066156.1      5321 ATAGAGTTGAAGTTTAATCCACCTGCTCTACAAGATGCTTATTACAGAGC   5370

AY323977.2      5301 CCGTGCTGGTGATGCTGCTAACTTTTGTGCACTCATACTCGCTTACAGTA   5350
                     ..|.||||||||.||||||||||||||||||||.||..|.||.|||.|||
MT066156.1      5371 AAGGGCTGGTGAAGCTGCTAACTTTTGTGCACTTATCTTAGCCTACTGTA   5420

AY323977.2      5351 ATAAAACTGTTGGCGAGCTTGGTGATGTCAGAGAAACTATGACCCATCTT   5400
                     ||||.||.||.||.|||.|.||||||||.||||||||.||||...|..|.
MT066156.1      5421 ATAAGACAGTAGGTGAGTTAGGTGATGTTAGAGAAACAATGAGTTACTTG   5470

AY323977.2      5401 CTACAGCATGCTAATTTGGAATCTGCAAAGCGAGTTCTTAATGTGGTGTG   5450
                     .|.||.|||||.|||||.||.|||...||..||||..|.||.||||||||
MT066156.1      5471 TTTCAACATGCCAATTTAGATTCTTGCAAAAGAGTCTTGAACGTGGTGTG   5520

AY323977.2      5451 TAAACATTGTGGTCAGAAAACTACTACCTTAACGGGTGTAGAAGCTGTGA   5500
                     ||||..||||||.||..|....||.|||.|.|.|||||||||||||||.|
MT066156.1      5521 TAAAACTTGTGGACAACAGCAGACAACCCTTAAGGGTGTAGAAGCTGTTA   5570

AY323977.2      5501 TGTATATGGGTACTCTATCTTATGATAATCTTAAGACAGGTGTTTCCATT   5550
                     ||||.|||||.||.||.||||||||..|..||||||.|||||||...||.
MT066156.1      5571 TGTACATGGGCACACTTTCTTATGAACAATTTAAGAAAGGTGTTCAGATA   5620

AY323977.2      5551 CCATGTGTGTGTGGTCGTGATGCTACACAATATCTAGTACAACAAGAGTC   5600
                     ||.|||..|||||||....|.||||||.||||||||||||||||.|||||
MT066156.1      5621 CCTTGTACGTGTGGTAAACAAGCTACAAAATATCTAGTACAACAGGAGTC   5670

AY323977.2      5601 TTCTTTTGTTATGATGTCTGCACCACCTGCTGAGTATAAATTACAGCAAG   5650
                     ..||||||||||||||||.||||||||||||.|||||.||.|..||||.|
MT066156.1      5671 ACCTTTTGTTATGATGTCAGCACCACCTGCTCAGTATGAACTTAAGCATG   5720

AY323977.2      5651 GTACATTCTTATGTGCGAATGAGTACACTGGTAACTATCAGTGTGGTCAT   5700
                     |||||||....|||||.|.|||||||||||||||.||.|||||||||||.
MT066156.1      5721 GTACATTTACTTGTGCTAGTGAGTACACTGGTAATTACCAGTGTGGTCAC   5770

AY323977.2      5701 TACACTCATATAACTGCTAAGGAGACCCTCTATCGTATTGACGGAGCTCA   5750
                     ||.|..|||||||||.||||.||.||..|.|||.|.||.|||||.|||..
MT066156.1      5771 TATAAACATATAACTTCTAAAGAAACTTTGTATTGCATAGACGGTGCTTT   5820

AY323977.2      5751 CCTTACAAAGATGTCAGAGTACAAAGGACCAGTGACTGATGTTTTCTACA   5800
                     .|||||||||...|||||.||||||||.||..|.||.|||||||||||||
MT066156.1      5821 ACTTACAAAGTCCTCAGAATACAAAGGTCCTATTACGGATGTTTTCTACA   5870

AY323977.2      5801 AGGAAACATCTTACACTACAACCATCAAGCCTGTGTCGTATAAACTCGAT   5850
                     |.||||....||||||.||||||||.||.||.||..|.||||||.|.|||
MT066156.1      5871 AAGAAAACAGTTACACAACAACCATAAAACCAGTTACTTATAAATTGGAT   5920

AY323977.2      5851 GGAGTTACTTACACAGAGATTGAACCAAAATTGGATGGGTATTATAAAAA   5900
                     ||.|||..||..|||||.|||||.||.||.|||||....||||||||.||
MT066156.1      5921 GGTGTTGTTTGTACAGAAATTGACCCTAAGTTGGACAATTATTATAAGAA   5970

AY323977.2      5901 GGATAATGCTTACTATACAGAGCAGCCTATAGACCTTGTACCAACTCAAC   5950
                     .||.|||.||||.|..||||||||.||.||.||.||||||||||..||||
MT066156.1      5971 AGACAATTCTTATTTCACAGAGCAACCAATTGATCTTGTACCAAACCAAC   6020

AY323977.2      5951 CATTACCAAATGCGAGTTTTGATAATTTCAAACTCACATGTTCTAACACA   6000
                     |||..|||||.||.||.||.||||||||.||..|...||||..|||.|..
MT066156.1      6021 CATATCCAAACGCAAGCTTCGATAATTTTAAGTTTGTATGTGATAATATC   6070

AY323977.2      6001 AAATTTGCTGATGATTTAAATCAAATGACAGGCTTCACAAAGCCAGCTTC   6050
                     ||||||||||||||||||||.||..|.||.||.|..|..||.||.|||||
MT066156.1      6071 AAATTTGCTGATGATTTAAACCAGTTAACTGGTTATAAGAAACCTGCTTC   6120

AY323977.2      6051 ACGAGAGCTATCTGTCACATTCTTCCCAGACTTGAATGGCGATGTAGTGG   6100
                     |.|||||||....||.|||||.|||||.|||||.|||||.|||||.||||
MT066156.1      6121 AAGAGAGCTTAAAGTTACATTTTTCCCTGACTTAAATGGTGATGTGGTGG   6170

AY323977.2      6101 CTATTGACTATAGACACTATTCAGCGAGTTTCAAGAAAGGTGCTAAATTA   6150
                     |||||||.||||.||||||..||.|...|||.||||||||.||||||||.
MT066156.1      6171 CTATTGATTATAAACACTACACACCCTCTTTTAAGAAAGGAGCTAAATTG   6220

AY323977.2      6151 CTGCATAAGCCAATTGTTTGGCACATTAACCAGGCTACAACCAAGACAAC   6200
                     .|.|||||.||.|||||||||||..|||||.|.||.||.|..||..|.||
MT066156.1      6221 TTACATAAACCTATTGTTTGGCATGTTAACAATGCAACTAATAAAGCCAC   6270

AY323977.2      6201 GTTCAAACCAAACACTTGGTGTTTACGTTGTCTTTGGAGTACAAAGCCAG   6250
                     ||..||||||||.||.||||||.||||||||||||||||.|||||.||||
MT066156.1      6271 GTATAAACCAAATACCTGGTGTATACGTTGTCTTTGGAGCACAAAACCAG   6320

AY323977.2      6251 TAGATACTTCAAATTCATTTGAAGTTCTGGCAGTAGAAGACACACAAGGA   6300
                     |.||.||.||||||||.|||||.||.|||.....|||.|||.|.||.|||
MT066156.1      6321 TTGAAACATCAAATTCGTTTGATGTACTGAAGTCAGAGGACGCGCAGGGA   6370

AY323977.2      6301 ATGGACAATCTTGCTTGTGAAAGTCAACAACCCACCTCTGAAGAAGTAGT   6350
                     |||||.||||||||.||.|||..||.|.||||...|||||||||||||||
MT066156.1      6371 ATGGATAATCTTGCCTGCGAAGATCTAAAACCAGTCTCTGAAGAAGTAGT   6420

AY323977.2      6351 GGAAAATCCTACCATACAGAAGGAAGTCATAGAGTGTGACGTGAAAACTA   6400
                     |||||||||||||||||||||.||.||..|.||||||.|.||||||||||
MT066156.1      6421 GGAAAATCCTACCATACAGAAAGACGTTCTTGAGTGTAATGTGAAAACTA   6470

AY323977.2      6401 CCGAAGTTGTAGGCAATGTCATACTTAAACCATCAGATGAAGGTGTTAAA   6450
                     |||||||||||||..|..|.||||||||||||.||.||.|..||.|.|||
MT066156.1      6471 CCGAAGTTGTAGGAGACATTATACTTAAACCAGCAAATAATAGTTTAAAA   6520

AY323977.2      6451 GTAACACAAGAGTTAGGTCATGAGGATCTTATGGCTGCTTATGTGGAAAA   6500
                     .|.|||.|||||.|.||.||....|||||.||||||||||||||.||.||
MT066156.1      6521 ATTACAGAAGAGGTTGGCCACACAGATCTAATGGCTGCTTATGTAGACAA   6570

AY323977.2      6501 CACAAGCATTACCATTAAGAAACCTAATGAGCTTTCACTAGCCTTAGGTT   6550
                     ..|.||..||||.|||||||||||||||||..|.||...||..|||||||
MT066156.1      6571 TTCTAGTCTTACTATTAAGAAACCTAATGAATTATCTAGAGTATTAGGTT   6620

AY323977.2      6551 TAAAAACAATTGCCACTCATGGTATTGCTGCAATTAATAGTGTTCCTTGG   6600
                     |.|||||..||||.|||||||||.|.|||||..||||||||||.||||||
MT066156.1      6621 TGAAAACCCTTGCTACTCATGGTTTAGCTGCTGTTAATAGTGTCCCTTGG   6670

AY323977.2      6601 AGTAAAATTTTGGCTTATGTCAAACCATT-CTTAGGACAAGCAGCAATTA   6649
                     ..||..||......|||||..||.||.|| ||||..| |||..|..|.||
MT066156.1      6671 GATACTATAGCTAATTATGCTAAGCCTTTTCTTAACA-AAGTTGTTAGTA   6719

AY323977.2      6650 CAACATCAAATTGCGCTAAGAGATTAGCACAACGTGTGTTTAACAATTAT   6699
                     ||||..|.||....|.||...|.|....|.|.|||||.|.||..||||||
MT066156.1      6720 CAACTACTAACATAGTTACACGGTGTTTAAACCGTGTTTGTACTAATTAT   6769

AY323977.2      6700 ATGCCTTATGTGTTTACATTATTGTTCCAATTGTGTACTTTTACTAAAAG   6749
                     |||||||||.|.|||||.||||||.|.|||||||||||||||||||.|||
MT066156.1      6770 ATGCCTTATTTCTTTACTTTATTGCTACAATTGTGTACTTTTACTAGAAG   6819

AY323977.2      6750 TACCAATTCTAGAATTAGAGCTTCACTACCTACAACTATTGCTAAAAATA   6799
                     |||.|||||||||||||.|||.||..|.||.||.|||||.||.||.||||
MT066156.1      6820 TACAAATTCTAGAATTAAAGCATCTATGCCGACTACTATAGCAAAGAATA   6869

AY323977.2      6800 GTGTTAAGAGTGTTGCTAAATTATGTTTGGATGCCGGCATTAATTATGTG   6849
                     .||||||||||||.|.||||||.|||.|.||.||.....||||||||.||
MT066156.1      6870 CTGTTAAGAGTGTCGGTAAATTTTGTCTAGAGGCTTCATTTAATTATTTG   6919

AY323977.2      6850 AAGTCACCCAAATTTTCTAAATTGTTCACAATCGCTATGTGGCTATTGTT   6899
                     ||||||||.||.|||||||||.||.|.|..||....||.|||.|.||..|
MT066156.1      6920 AAGTCACCTAATTTTTCTAAACTGATAAATATTATAATTTGGTTTTTACT   6969

AY323977.2      6900 GTTAAGTATTTGCTTAGGTTCTCTAATCTGTGTAACTGCTGCTTTTGGTG   6949
                     .||||||.|||||.||||||||.||||||....|||.||||||||.||||
MT066156.1      6970 ATTAAGTGTTTGCCTAGGTTCTTTAATCTACTCAACCGCTGCTTTAGGTG   7019

AY323977.2      6950 TACTCTTATCTAATTTTGGTGCTCCTTCTTATTGTAATGGCGTTAGAGAA   6999
                     |..|..|.||||||||.||....||||||||.||||.|||....||||||
MT066156.1      7020 TTTTAATGTCTAATTTAGGCATGCCTTCTTACTGTACTGGTTACAGAGAA   7069

AY323977.2      7000 TTGTATCTTAATTCGTCTAACGTTACTACTATGGATTTCTGTGAAGGTTC   7049
                     ...|||.|.||.||..||||.||.||||.|......|.||||...|||||
MT066156.1      7070 GGCTATTTGAACTCTACTAATGTCACTATTGCAACCTACTGTACTGGTTC   7119

AY323977.2      7050 TTTTCCTTGCAGCATTTGTTTAAGTGGATTAGACTCCCTTGATTCTTATC   7099
                     |.|.|||||.||..|||||.|.|||||.|||||.||..|.||..|.||||
MT066156.1      7120 TATACCTTGTAGTGTTTGTCTTAGTGGTTTAGATTCTTTAGACACCTATC   7169

AY323977.2      7100 CAGCTCTTGAAACCATTCAGGTGACGATTTCATCGTACAAGCTAGACTTG   7149
                     |..||.|.|||||.||.||..|.||.||||||||.|..||....||.||.
MT066156.1      7170 CTTCTTTAGAAACTATACAAATTACCATTTCATCTTTTAAATGGGATTTA   7219

AY323977.2      7150 ACAATTTTAGGTCTGGCCGCTGAGTGGGTTTTGGCATATATGTTGTTCAC   7199
                     ||...|||.||..|.|..||.||||||.|||||||||||||..|.|||||
MT066156.1      7220 ACTGCTTTTGGCTTAGTTGCAGAGTGGTTTTTGGCATATATTCTTTTCAC   7269

AY323977.2      7200 AAAATTCTTTTATTTATTAGGTCTTTCAGCTATAATGCAGGTGTTCTTTG   7249
                     .|..||.||.|||.||.|.||..|..|.||.||.|||||..||||.||..
MT066156.1      7270 TAGGTTTTTCTATGTACTTGGATTGGCTGCAATCATGCAATTGTTTTTCA   7319

AY323977.2      7250 GCTATTTTGCTAGTCATTTCATCAGCAATTCTTGGCTCATGTGGTTTATC   7299
                     ||||||||||....|||||.||.||.|||||||||||.||||||||.||.
MT066156.1      7320 GCTATTTTGCAGTACATTTTATTAGTAATTCTTGGCTTATGTGGTTAATA   7369

AY323977.2      7300 ATTAGTATTGTACAAATGGCACCCGTTTCTGCAATGGTTAGGATGTACAT   7349
                     ||||.|.|||||||||||||.||..||||.||.||||||||.||||||||
MT066156.1      7370 ATTAATCTTGTACAAATGGCCCCGATTTCAGCTATGGTTAGAATGTACAT   7419

AY323977.2      7350 CTTCTTTGCTTCTTTCTACTACATATGGAAGAGCTATGTTCATATCATGG   7399
                     |||||||||.||.||.||.||..|||||||.||.|||||.|||.|..|.|
MT066156.1      7420 CTTCTTTGCATCATTTTATTATGTATGGAAAAGTTATGTGCATGTTGTAG   7469

AY323977.2      7400 ATGGTTGCACCTCTTCGACTTGCATGATGTGCTATAAGCGCAATCGTGCC   7449
                     |.|||||.|..||.||.|||||.||||||||.||.||.||.|||.|.||.
MT066156.1      7470 ACGGTTGTAATTCATCAACTTGTATGATGTGTTACAAACGTAATAGAGCA   7519

AY323977.2      7450 ACACGCGTTGAGTGTACAACTATTGTTAATGGCATGAAGAGATCTTTCTA   7499
                     |||.|.||.||.||||||||||||||||||||..|.|..||.||.||.||
MT066156.1      7520 ACAAGAGTCGAATGTACAACTATTGTTAATGGTGTTAGAAGGTCCTTTTA   7569

AY323977.2      7500 TGTCTATGCAAATGGAGGCCGTGGCTTCTGCAAGACTCACAATTGGAATT   7549
                     |||||||||.||||||||....|||||.|||||....|||||||||||||
MT066156.1      7570 TGTCTATGCTAATGGAGGTAAAGGCTTTTGCAAACTACACAATTGGAATT   7619

AY323977.2      7550 GTCTCAATTGTGACACATTTTGCACTGGTAGTACATTCATTAGTGATGAA   7599
                     ||.|.||||||||.|||||.||..|||||||||||||.||||||||||||
MT066156.1      7620 GTGTTAATTGTGATACATTCTGTGCTGGTAGTACATTTATTAGTGATGAA   7669

AY323977.2      7600 GTTGCTCGTGATTTGTCACTCCAGTTTAAAAGACCAATCAACCCTACTGA   7649
                     |||||..|.||.||||||||.|||||||||||||||||.||.||||||||
MT066156.1      7670 GTTGCGAGAGACTTGTCACTACAGTTTAAAAGACCAATAAATCCTACTGA   7719

AY323977.2      7650 CCAGTCATCGTATATTGTTGATAGTGTTGCTGTGAAAAATGGCGCGCTTC   7699
                     ||||||.||.||.||.||||||||||||.|.|||||.|||||..|..|.|
MT066156.1      7720 CCAGTCTTCTTACATCGTTGATAGTGTTACAGTGAAGAATGGTTCCATCC   7769

AY323977.2      7700 ACCTCTACTTTGACAAGGCTGGTCAAAAGACCTATGAGAGACATCCGCTC   7749
                     |.||.||||||||.||.||||||||||||||.|||||.||||||.|.|||
MT066156.1      7770 ATCTTTACTTTGATAAAGCTGGTCAAAAGACTTATGAAAGACATTCTCTC   7819

AY323977.2      7750 TCCCATTTTGTCAATTTAGACAATTTGAGAGCTAACAACACTAAAGGTTC   7799
                     ||.||||||||.||.||||||||..||||||||||.||||||||||||||
MT066156.1      7820 TCTCATTTTGTTAACTTAGACAACCTGAGAGCTAATAACACTAAAGGTTC   7869

AY323977.2      7800 ACTGCCTATTAATGTCATAGTTTTTGATGGCAAGTCCAAATGCGACGAGT   7849
                     |.|||||||||||||.||||||||||||||.||.||.|||||.||.||.|
MT066156.1      7870 ATTGCCTATTAATGTTATAGTTTTTGATGGTAAATCAAAATGTGAAGAAT   7919

AY323977.2      7850 CTGCTTCTAAGTCTGCTTCTGTGTACTACAGTCAGCTGATGTGCCAACCT   7899
                     |..||.|.||.||.||.|||||.||||||||||||||.|||||.||||||
MT066156.1      7920 CATCTGCAAAATCAGCGTCTGTTTACTACAGTCAGCTTATGTGTCAACCT   7969

AY323977.2      7900 ATTCTGTTGCTTGACCAAGCTCTTGTATCAGACGTTGGAGATAGTACTGA   7949
                     ||.|||||.||.||.||.||..|.||.||.||.|||||.||||||.|.||
MT066156.1      7970 ATACTGTTACTAGATCAGGCATTAGTGTCTGATGTTGGTGATAGTGCGGA   8019

AY323977.2      7950 AGTTTCCGTTAAGATGTTTGATGCTTATGTCGACACCTTTTCAGCAACTT   7999
                     ||||.|.|||||.||||||||||||||.||..|.||.||||||.||||||
MT066156.1      8020 AGTTGCAGTTAAAATGTTTGATGCTTACGTTAATACGTTTTCATCAACTT   8069

AY323977.2      8000 TTAGTGTTCCTATGGAAAAACTTAAGGCACTTGTTGCTACAGCTCACAGC   8049
                     |||..||.||.|||||||||||.||..||||.|||||.||.||..|....
MT066156.1      8070 TTAACGTACCAATGGAAAAACTCAAAACACTAGTTGCAACTGCAGAAGCT   8119

AY323977.2      8050 GAGTTAGCAAAGGGTGTAGCTTTAGATGGTGTCCTTTCTACATTCGTGTC   8099
                     ||..|.||||||..|||..|.|||||...||||.|.|||||.||..|.||
MT066156.1      8120 GAACTTGCAAAGAATGTGTCCTTAGACAATGTCTTATCTACTTTTATTTC   8169

AY323977.2      8100 AGCTGCCCGACAAGGTGTTGTTGATACCGATGTTGACACAAAGGATGTTA   8149
                     |||.||.||.|||||..||||||||.|.|||||.||.||.||.||||||.
MT066156.1      8170 AGCAGCTCGGCAAGGGTTTGTTGATTCAGATGTAGAAACTAAAGATGTTG   8219

AY323977.2      8150 TTGAATGTCTCAAACTTTCACATCACTCTGACTTAGAAGTGACAGGTGAC   8199
                     ||||||||||.|||.|.||||||||.||||||.|||||||.||.||.||.
MT066156.1      8220 TTGAATGTCTTAAATTGTCACATCAATCTGACATAGAAGTTACTGGCGAT   8269

AY323977.2      8200 AGTTGTAACAATTTCATGCTCACCTATAATAAGGTTGAAAACATGACGCC   8249
                     ||||||||.||.|..||||||||||||||.||.||||||||||||||.||
MT066156.1      8270 AGTTGTAATAACTATATGCTCACCTATAACAAAGTTGAAAACATGACACC   8319

AY323977.2      8250 CAGAGATCTTGGCGCATGTATTGACTGTAATGCAAGGCATATCAATGCCC   8299
                     |.|.||.|||||.||.|||||||||||||.|||..|.|||||.|||||.|
MT066156.1      8320 CCGTGACCTTGGTGCTTGTATTGACTGTAGTGCGCGTCATATTAATGCGC   8369

AY323977.2      8300 AAGTAGCAAAAAGTCACAATGTTTCACTCATCTGGAATGTAAAAGACTAC   8349
                     |.|||||||||||||||||..||.|..|.||.|||||.||.|||||.|.|
MT066156.1      8370 AGGTAGCAAAAAGTCACAACATTGCTTTGATATGGAACGTTAAAGATTTC   8419

AY323977.2      8350 ATGTCTTTATCTGAACAGCTGCGTAAACAAATTCGTAGTGCTGCCAAGAA   8399
                     |||||.||.||||||||.||.||.||||||||.|||||||||||.||.||
MT066156.1      8420 ATGTCATTGTCTGAACAACTACGAAAACAAATACGTAGTGCTGCTAAAAA   8469

AY323977.2      8400 GAACAACATACCTTTTAGACTAACTTGTGCTACAACTAGACAGGTTGTCA   8449
                     |||.|||.|||||||||...|.||.|||||.||.||||||||.|||||.|
MT066156.1      8470 GAATAACTTACCTTTTAAGTTGACATGTGCAACTACTAGACAAGTTGTTA   8519

AY323977.2      8450 ATGTCATAACTACTAAAATCTCACTCAAGGGTGGTAAGATTGTTAGTACT   8499
                     ||||..||||.||.||.||..||||.|||||||||||.|||||||.||.|
MT066156.1      8520 ATGTTGTAACAACAAAGATAGCACTTAAGGGTGGTAAAATTGTTAATAAT   8569

AY323977.2      8500 TGTTTTAAACTTATGCTTAAGGCCACATTATTGTGCGTTCTTGCTGCATT   8549
                     ||.||.||.|...|..||||.|..|||.|..|||.|.||.|||.|||...
MT066156.1      8570 TGGTTGAAGCAGTTAATTAAAGTTACACTTGTGTTCCTTTTTGTTGCTGC   8619

AY323977.2      8550 GGTTTGTTATATCGTTATGCCAGTACATACATTGTCAATCCATGATGGTT   8599
                     ..|||..|||.|..|.|..||.||.|||....||||.|..|||..||..|
MT066156.1      8620 TATTTTCTATTTAATAACACCTGTTCATGTCATGTCTAAACATACTGACT   8669

AY323977.2      8600 ACACAAATGAAATCATTGGTTACAAAGCCATTCAGGATGGTGTCACTCGT   8649
                     ...|||.|||||||||.||.|||||.||.|||.|.|.|||||||||||||
MT066156.1      8670 TTTCAAGTGAAATCATAGGATACAAGGCTATTGATGGTGGTGTCACTCGT   8719

AY323977.2      8650 GACATCATTTCTACTGATGATTGTTTTGCAAATAAACATGCTGGTTTTGA   8699
                     |||||....|||||.|||..|||||||||.||.||||||||||.||||||
MT066156.1      8720 GACATAGCATCTACAGATACTTGTTTTGCTAACAAACATGCTGATTTTGA   8769

AY323977.2      8700 CGCATGGTTTAGCCAGCGTGGTGGTTCATACAAAAATGACAAAAGCTGCC   8749
                     |.|||||||||||||||||||||||...||.|..|||||||||...||||
MT066156.1      8770 CACATGGTTTAGCCAGCGTGGTGGTAGTTATACTAATGACAAAGCTTGCC   8819

AY323977.2      8750 CTGTAGTAGCTGCTATCATTACAAGAGAGATTGGTTTCATAGTGCCTGGC   8799
                     |..|..|.|||||..||||.||||||||..|.|||||..|.||||||||.
MT066156.1      8820 CATTGATTGCTGCAGTCATAACAAGAGAAGTGGGTTTTGTCGTGCCTGGT   8869

AY323977.2      8800 TTACCGGGTACTGTGCTGAGAGCAATCAATGGTGACTTCTTGCATTTTCT   8849
                     ||.||.||.||..|..|..|..|||..|||||||||||.||||||||..|
MT066156.1      8870 TTGCCTGGCACGATATTACGCACAACTAATGGTGACTTTTTGCATTTCTT   8919

AY323977.2      8850 ACCTCGTGTTTTTAGTGCTGTTGGCAACATTTGCTACACACCTTCCAAAC   8899
                     ||||.|.|||||||||||.|||||.|||||.||.||||||||.||.||||
MT066156.1      8920 ACCTAGAGTTTTTAGTGCAGTTGGTAACATCTGTTACACACCATCAAAAC   8969

AY323977.2      8900 TCATTGAGTATAGTGATTTTGCTACCTCTGCTTGCGTTCTTGCTGCTGAG   8949
                     |.||.|||||.|.|||.|||||.||.||.|||||.|||.|.||||||||.
MT066156.1      8970 TTATAGAGTACACTGACTTTGCAACATCAGCTTGTGTTTTGGCTGCTGAA   9019

AY323977.2      8950 TGTACAATTTTTAAGGATGCTATGGGCAAACCTGTGCCATATTGTTATGA   8999
                     ||||||||||||||.||||||...||.||.||.||.||||||||||||||
MT066156.1      9020 TGTACAATTTTTAAAGATGCTTCTGGTAAGCCAGTACCATATTGTTATGA   9069

AY323977.2      9000 CACTAATTTGCTAGAGGGTTCTATTTCTTATAGTGAGCTT-CGTCCAGAC   9048
                     .||.|||.|.|||||.||||||.||.||||| |..||.|| ||.||.|||
MT066156.1      9070 TACCAATGTACTAGAAGGTTCTGTTGCTTAT-GAAAGTTTACGCCCTGAC   9118

AY323977.2      9049 ACTCGTTATGTGCTTATGGATGGTTCCATCATACAGTTTCCTAACACTTA   9098
                     ||.|||||||||||.||||||||.||.||.||.||.|||||||||||.||
MT066156.1      9119 ACACGTTATGTGCTCATGGATGGCTCTATTATTCAATTTCCTAACACCTA   9168

AY323977.2      9099 CCTGGAGGGTTCTGTTAGAGTAGTAACAACTTTTGATGCTGAGTACTGTA   9148
                     |||.||.||||||||||||||.|||||||||||||||.||||||||||||
MT066156.1      9169 CCTTGAAGGTTCTGTTAGAGTGGTAACAACTTTTGATTCTGAGTACTGTA   9218

AY323977.2      9149 GACATGGTACATGCGAAAGGTCAGAAGTAGGTATTTGCCTATCTACCAGT   9198
                     |.||.||.||.||.|||||.|||||||..|||.||||..|||||||.|||
MT066156.1      9219 GGCACGGCACTTGTGAAAGATCAGAAGCTGGTGTTTGTGTATCTACTAGT   9268

AY323977.2      9199 GGTAGATGGGTTCTTAATAATGAGCATTACAGAGCTCTATCAGGAGTTTT   9248
                     |||||||||||.|||||.|||||..||||||||.||.||.||||||||||
MT066156.1      9269 GGTAGATGGGTACTTAACAATGATTATTACAGATCTTTACCAGGAGTTTT   9318

AY323977.2      9249 CTGTGGTGTTGATGCGATGAATCTCATAGCTAACATCTTTACTCCTCTTG   9298
                     |||||||||.|||||..|.|||.|..|..||||.||.|||||.||.||..
MT066156.1      9319 CTGTGGTGTAGATGCTGTAAATTTACTTACTAATATGTTTACACCACTAA   9368

AY323977.2      9299 TGCAACCTGTGGGTGCTTTAGATGTGTCTGCTTCAGTAGTGGCTGGTGGT   9348
                     |.||||||.|.||||||||.||..|.||.||.||..||||.|||||||||
MT066156.1      9369 TTCAACCTATTGGTGCTTTGGACATATCAGCATCTATAGTAGCTGGTGGT   9418

AY323977.2      9349 ATTATTGCCATATTGGTGACTTGTGCTGCCTACTACTTTATGAAATTCAG   9398
                     |||.|.||.||..|.||.||.||...|||||||||.|||||||..||.||
MT066156.1      9419 ATTGTAGCTATCGTAGTAACATGCCTTGCCTACTATTTTATGAGGTTTAG   9468

AY323977.2      9399 ACGTGTTTTTGGTGAGTACAACCATGTTGTTGCTGCTAATGCACTTTTGT   9448
                     |.|.|.|||||||||.||||..|||||.|||||...||||.|..|..|.|
MT066156.1      9469 AAGAGCTTTTGGTGAATACAGTCATGTAGTTGCCTTTAATACTTTACTAT   9518

AY323977.2      9449 TTTTGATGTCTTTCACTATACTCTGTCTGGTACCAGCTTACAGCTTTCTG   9498
                     |..|.|||||.||||||.||||||||.|...|||||.||||...||..|.
MT066156.1      9519 TCCTTATGTCATTCACTGTACTCTGTTTAACACCAGTTTACTCATTCTTA   9568

AY323977.2      9499 CCGGGAGTCTACTCAGTCTTTTACTTGTACTTGACATTCTATTTCACCAA   9548
                     ||.||.||.||.||.||..|||||||||||||||||||.|||.|.||.||
MT066156.1      9569 CCTGGTGTTTATTCTGTTATTTACTTGTACTTGACATTTTATCTTACTAA   9618

AY323977.2      9549 TGATGTTTCATTCTTGGCTCACCTTCAATGGTTTGCCATGTTTTCTCCTA   9598
                     |||||||||.||.||.||.||..||||.|||.|.|..|||||..|.|||.
MT066156.1      9619 TGATGTTTCTTTTTTAGCACATATTCAGTGGATGGTTATGTTCACACCTT   9668

AY323977.2      9599 TTGTGCCTTTTTGGATAACAGCAATCTATGTATTCTGTATTTCTCTGAAG   9648
                     |.||.|||||.|||||||||......|||.|..|.||||||||....|||
MT066156.1      9669 TAGTACCTTTCTGGATAACAATTGCTTATATCATTTGTATTTCCACAAAG   9718

AY323977.2      9649 CACTGCCATTGGTTCTTTAACAACTATCTTAGGAAAAGAGTCATGTTTAA   9698
                     ||.|.|.||||||||||||..||.||.||.|.||.|.|.||..|.|||||
MT066156.1      9719 CATTTCTATTGGTTCTTTAGTAATTACCTAAAGAGACGTGTAGTCTTTAA   9768

AY323977.2      9699 TGGAGTTACATTTAGTACCTTCGAGGAGGCTGCTTTGTGTACCTTTTTGC   9748
                     |||.|||.|.||||||||.||.||.||.|||||..||||.|||||||||.
MT066156.1      9769 TGGTGTTTCCTTTAGTACTTTTGAAGAAGCTGCGCTGTGCACCTTTTTGT   9818

AY323977.2      9749 TCAACAAGGAAATGTACCTAAAATTGCGTAGCGAGACACTGTTGCCACTT   9798
                     |.||.||.||||||||.|||||.||||||||.||....||.||.||.|||
MT066156.1      9819 TAAATAAAGAAATGTATCTAAAGTTGCGTAGTGATGTGCTATTACCTCTT   9868

AY323977.2      9799 ACACAGTATAACAGGTATCTTGCTCTATATAACAAGTACAAGTATTTCAG   9848
                     ||.||.|||||.||.||..|.|||||.|||||.||||||||||||||.||
MT066156.1      9869 ACGCAATATAATAGATACTTAGCTCTTTATAATAAGTACAAGTATTTTAG   9918

AY323977.2      9849 TGGAGCCTTAGATACTACCAGCTATCGTGAAGCAGCTTGCTGCCACTTAG   9898
                     ||||||..|.|||||.||.|||||..|.|||||.|||||.||.||..|.|
MT066156.1      9919 TGGAGCAATGGATACAACTAGCTACAGAGAAGCTGCTTGTTGTCATCTCG   9968

AY323977.2      9899 CAAAGGCTCTAAATGACTTTAGCAACTCAGGTGCTGATGTTCTCTACCAA   9948
                     ||||||||||.||||||||.||.|||||||||.||||||||||.||||||
MT066156.1      9969 CAAAGGCTCTCAATGACTTCAGTAACTCAGGTTCTGATGTTCTTTACCAA  10018

AY323977.2      9949 CCACCACAGACATCAATCACTTCTGCTGTTCTGCAGAGTGGTTTTAGGAA   9998
                     ||||||||.||.||.|||||.||.||||||.||||||||||||||||.||
MT066156.1     10019 CCACCACAAACCTCTATCACCTCAGCTGTTTTGCAGAGTGGTTTTAGAAA  10068

AY323977.2      9999 AATGGCATTCCCGTCAGGCAAAGTTGAAGGGTGCATGGTACAAGTAACCT  10048
                     ||||||||||||.||.||.||||||||.||.||.||||||||||||||.|
MT066156.1     10069 AATGGCATTCCCATCTGGTAAAGTTGAGGGTTGTATGGTACAAGTAACTT  10118

AY323977.2     10049 GTGGAACTACAACTCTTAATGGATTGTGGTTGGATGACACAGTATACTGT  10098
                     ||||.||.||.||.|||||.||..|.|||.|.||||||..|||.||||||
MT066156.1     10119 GTGGTACAACTACACTTAACGGTCTTTGGCTTGATGACGTAGTTTACTGT  10168

AY323977.2     10099 CCAAGACATGTCATTTGCACAGCAGAAGACATGCTTAATCCTAACTATGA  10148
                     |||||||||||.||.|||||..|.||||||||||||||.|||||.|||||
MT066156.1     10169 CCAAGACATGTGATCTGCACCTCTGAAGACATGCTTAACCCTAATTATGA  10218

AY323977.2     10149 AGATCTGCTCATTCGCAAATCCAACCATAGCTTTCTTGTTCAGGCTGGCA  10198
                     ||||.|.||||||||.||.||.||.||||..||..|.||.||||||||.|
MT066156.1     10219 AGATTTACTCATTCGTAAGTCTAATCATAATTTCTTGGTACAGGCTGGTA  10268

AY323977.2     10199 ATGTTCAACTTCGTGTTATTGGCCATTCTATGCAAAATTGTCTGCTTAGG  10248
                     ||||||||||..|.||||||||.||||||||||||||||||.|.||||.|
MT066156.1     10269 ATGTTCAACTCAGGGTTATTGGACATTCTATGCAAAATTGTGTACTTAAG  10318

AY323977.2     10249 CTTAAAGTTGATACTTCTAACCCTAAGACACCCAAGTATAAATTTGTCCG  10298
                     |||||.||||||||..|.||.|||||||||||.||||||||.|||||.||
MT066156.1     10319 CTTAAGGTTGATACAGCCAATCCTAAGACACCTAAGTATAAGTTTGTTCG  10368

AY323977.2     10299 TATCCAACCTGGTCAAACATTTTCAGTTCTAGCATGCTACAATGGTTCAC  10348
                     .||.|||||.||.||.||.||||||||..||||.||.|||||||||||||
MT066156.1     10369 CATTCAACCAGGACAGACTTTTTCAGTGTTAGCTTGTTACAATGGTTCAC  10418

AY323977.2     10349 CATCTGGTGTTTATCAGTGTGCCATGAGACCTAATCATACCATTAAAGGT  10398
                     |||||||||||||.||.|||||.|||||.||.|||...||.|||||.|||
MT066156.1     10419 CATCTGGTGTTTACCAATGTGCTATGAGGCCCAATTTCACTATTAAGGGT  10468

AY323977.2     10399 TCTTTCCTTAATGGATCATGTGGTAGTGTTGGTTTTAACATTGATTATGA  10448
                     ||.|||||||||||.||||||||||||||||||||||||||.||||||||
MT066156.1     10469 TCATTCCTTAATGGTTCATGTGGTAGTGTTGGTTTTAACATAGATTATGA  10518

AY323977.2     10449 TTGCGTGTCTTTCTGCTATATGCATCATATGGAGCTTCCAACAGGAGTAC  10498
                     .||.||.|||||.||.||.|||||.||||||||..|.|||||.|||||.|
MT066156.1     10519 CTGTGTCTCTTTTTGTTACATGCACCATATGGAATTACCAACTGGAGTTC  10568

AY323977.2     10499 ACGCTGGTACTGACTTAGAAGGTAAATTCTATGGTCCATTTGTTGACAGA  10548
                     |.|||||.||.||||||||||||||.||.|||||.||.|||||||||||.
MT066156.1     10569 ATGCTGGCACAGACTTAGAAGGTAACTTTTATGGACCTTTTGTTGACAGG  10618

AY323977.2     10549 CAAACTGCACAGGCTGCAGGTACAGACACAACCATAACATTAAATGTTTT  10598
                     |||||.|||||.||.||.|||||.||||||||.||.|||.|.||||||||
MT066156.1     10619 CAAACAGCACAAGCAGCTGGTACGGACACAACTATTACAGTTAATGTTTT  10668

AY323977.2     10599 GGCATGGCTGTATGCTGCTGTTATCAATGGTGATAGGTGGTTTCTTAATA  10648
                     .||.|||.||||.|||||||||||.|||||.||.|||||||||||.|||.
MT066156.1     10669 AGCTTGGTTGTACGCTGCTGTTATAAATGGAGACAGGTGGTTTCTCAATC  10718

AY323977.2     10649 GATTCACCACTACTTTGAATGACTTTAACCTTGTGGCAATGAAGTACAAC  10698
                     ||||.|||||.|||.|.||||||||||||||||||||.|||||||||||.
MT066156.1     10719 GATTTACCACAACTCTTAATGACTTTAACCTTGTGGCTATGAAGTACAAT  10768

AY323977.2     10699 TATGAACCTTTGACACAAGATCATGTTGACATATTGGGACCTCTTTCTGC  10748
                     |||||||||.|.||||||||.||||||||||||.|.||||||||||||||
MT066156.1     10769 TATGAACCTCTAACACAAGACCATGTTGACATACTAGGACCTCTTTCTGC  10818

AY323977.2     10749 TCAAACAGGAATTGCCGTCTTAGATATGTGTGCTGCTTTGAAAGAGCTGC  10798
                     ||||||.|||||||||||.|||||||||||||||.|.||.|||||..|.|
MT066156.1     10819 TCAAACTGGAATTGCCGTTTTAGATATGTGTGCTTCATTAAAAGAATTAC  10868

AY323977.2     10799 TGCAGAATGGTATGAATGGTCGTACTATCCTTGGTAGCACTATTTTAGAA  10848
                     ||||.||||||||||||||.|||||.||..|.|||||..||.|.||||||
MT066156.1     10869 TGCAAAATGGTATGAATGGACGTACCATATTGGGTAGTGCTTTATTAGAA  10918

AY323977.2     10849 GATGAGTTTACACCATTTGATGTTGTTAGACAATGCTCTGGTGTTACCTT  10898
                     |||||.||||||||.|||||||||||||||||||||||.||||||||.||
MT066156.1     10919 GATGAATTTACACCTTTTGATGTTGTTAGACAATGCTCAGGTGTTACTTT  10968

AY323977.2     10899 CCAAGGTAAGTTCAAGAAAATTGTTAAGGGCACTCATCATTGGATGCTTT  10948
                     ||||.||....|.||.|.||...|.|||||.||.||.||.|||.||.|..
MT066156.1     10969 CCAAAGTGCAGTGAAAAGAACAATCAAGGGTACACACCACTGGTTGTTAC  11018

AY323977.2     10949 TAACTTTCTTGACATCACTATTGATTCTTGTTCAAAGTACACAGTGGTCA  10998
                     |.||..|.|||||.|||||.||..||.|.||.||.|||||.||.|||||.
MT066156.1     11019 TCACAATTTTGACTTCACTTTTAGTTTTAGTCCAGAGTACTCAATGGTCT  11068

AY323977.2     10999 CTGTTTTTCTTTGTTTACGAGAATGCTTTCTTGCCATTTACTCTTGGTAT  11048
                     .||||.||.|||.|.||.||.|||||.||.||.||.|||.||.|.|||||
MT066156.1     11069 TTGTTCTTTTTTTTNTATGAAAATGCCTTTTTACCTTTTGCTATGGGTAT  11118

AY323977.2     11049 TATGGCAATTGCTGCATGTGCTATGCTGCTTGTTAAGCATAAGCACGCAT  11098
                     |||.||.||..||||.|.|||.|||.||.||||.||.||||||||.||||
MT066156.1     11119 TATTGCTATGTCTGCTTTTGCAATGATGTTTGTCAAACATAAGCATGCAT  11168

AY323977.2     11099 TCTTGTGCTTGTTTCTGTTACCTTCTCTTGCAACAGTTGCTTACTTTAAT  11148
                     |..|.||.||||||.||||||||||||||||.||.||.|||||.||||||
MT066156.1     11169 TTCTCTGTTTGTTTTTGTTACCTTCTCTTGCCACTGTAGCTTATTTTAAT  11218

AY323977.2     11149 ATGGTCTACATGCCTGCTAGCTGGGTGATGCGTATCATGACATGGCTTGA  11198
                     ||||||||.|||||||||||.||||||||||||||.|||||||||.|.||
MT066156.1     11219 ATGGTCTATATGCCTGCTAGTTGGGTGATGCGTATTATGACATGGTTGGA  11268

AY323977.2     11199 ATTGGCTGACACTAGCTTGTCTGGTTATAGGCTTAAGGATTGTGTTATGT  11248
                     ..|||.|||.|||||.||||||||||.||.|||.||.||.||||||||||
MT066156.1     11269 TATGGTTGATACTAGTTTGTCTGGTTTTAAGCTAAAAGACTGTGTTATGT  11318

AY323977.2     11249 ATGCTTCAGCTTTAGTTTTGCTTATTCTCATGACAGCTCGCACTGTTTAT  11298
                     ||||.||||||.||||.||.||.||.||.||||||||..|.|||||.|||
MT066156.1     11319 ATGCATCAGCTGTAGTGTTACTAATCCTTATGACAGCAAGAACTGTGTAT  11368

AY323977.2     11299 GATGATGCTGCTAGACGTGTTTGGACACTGATGAATGTCATTACACTTGT  11348
                     |||||||.||||||..|.||.||||||||.|||||||||.|.|||||.||
MT066156.1     11369 GATGATGGTGCTAGGAGAGTGTGGACACTTATGAATGTCTTGACACTCGT  11418

AY323977.2     11349 TTACAAAGTCTACTATGGTAATGCTTTAGATCAAGCTATTTCCATGTGGG  11398
                     |||.|||||.||.|||||||||||||||||||||||.|||||||||||||
MT066156.1     11419 TTATAAAGTTTATTATGGTAATGCTTTAGATCAAGCCATTTCCATGTGGG  11468

AY323977.2     11399 CCTTAGTTATTTCTGTAACCTCTAACTATTCTGGTGTCGTTACGACTATC  11448
                     |..|..|.||.|||||.||.||||||||.||.|||||.|||||.|||.||
MT066156.1     11469 CTCTTATAATCTCTGTTACTTCTAACTACTCAGGTGTAGTTACAACTGTC  11518

AY323977.2     11449 ATGTTTTTAGCTAGAGCTATAGTGTTTGTGTGTGTTGAGTATTACCCATT  11498
                     ||||||||.||.||||.|||.||.|||.|||||||||||||||.|||..|
MT066156.1     11519 ATGTTTTTGGCCAGAGGTATTGTTTTTATGTGTGTTGAGTATTGCCCTAT  11568

AY323977.2     11499 GTTATTTATTACTGGCAACACCTTACAGTGTATCATGCTTGTTTATTGTT  11548
                     .||.||.||.|||||.||.||..|.||||||||.|||||.||||||||||
MT066156.1     11569 TTTCTTCATAACTGGTAATACACTTCAGTGTATAATGCTAGTTTATTGTT  11618

AY323977.2     11549 TCTTAGGCTATTGTTGCTGCTGCTACTTTGGCCTTTTCTGTTTACTCAAC  11598
                     ||||||||||||.|||....||.|||||||||||.||.||||||||||||
MT066156.1     11619 TCTTAGGCTATTTTTGTACTTGTTACTTTGGCCTCTTTTGTTTACTCAAC  11668

AY323977.2     11599 CGTTACTTCAGGCTTACTCTTGGTGTTTATGACTACTTGGTCTCTACACA  11648
                     ||.|||||.||.||.|||||||||||||||||.|||||.||.||||||||
MT066156.1     11669 CGCTACTTTAGACTGACTCTTGGTGTTTATGATTACTTAGTTTCTACACA  11718

AY323977.2     11649 AGAATTTAGGTATATGAACTCCCAGGGGCTTTTGCCTCCTAAGAGTAGTA  11698
                     .||.|||||.||||||||.||.|||||.||..|.||.||.||||.|||.|
MT066156.1     11719 GGAGTTTAGATATATGAATTCACAGGGACTACTCCCACCCAAGAATAGCA  11768

AY323977.2     11699 TTGATGCTTTCAAGCTTAACATTAAGTTGTTGGGTATTGGAGGTAAACCA  11748
                     |.|||||.|||||.||.||||||||.|||||||||.||||.||.|||||.
MT066156.1     11769 TAGATGCCTTCAAACTCAACATTAAATTGTTGGGTGTTGGTGGCAAACCT  11818

AY323977.2     11749 TGTATCAAGGTTGCTACTGTACAGTCTAAAATGTCTGACGTAAAGTGCAC  11798
                     ||||||||.||.||.||||||||||||||||||||.||.|||||||||||
MT066156.1     11819 TGTATCAAAGTAGCCACTGTACAGTCTAAAATGTCAGATGTAAAGTGCAC  11868

AY323977.2     11799 ATCTGTGGTACTGCTCTCGGTTCTTCAACAACTTAGAGTAGAGTCATCTT  11848
                     |||.||.||..|.|||||.|||.|.||||||||.||||||||.|||||.|
MT066156.1     11869 ATCAGTAGTCTTACTCTCAGTTTTGCAACAACTCAGAGTAGAATCATCAT  11918

AY323977.2     11849 CTAAATTGTGGGCACAATGTGTACAACTCCACAATGATATTCTTCTTGCA  11898
                     |||||||||||||.||||||||.||..|.||||||||.|||||..|.||.
MT066156.1     11919 CTAAATTGTGGGCTCAATGTGTCCAGTTACACAATGACATTCTCTTAGCT  11968

AY323977.2     11899 AAAGACACAACTGAAGCTTTCGAGAAGATGGTTTCTCTTTTGTCTGTTTT  11948
                     |||||.||.||||||||.||.||.||.||||||||.||..|.||||||||
MT066156.1     11969 AAAGATACTACTGAAGCCTTTGAAAAAATGGTTTCACTACTTTCTGTTTT  12018

AY323977.2     11949 GCTATCCATGCAGGGTGCTGTAGACATTAATAGGTTGTGCGAGGAAATGC  11998
                     |||.|||||||||||||||||||||||.||.|.|.|.||.||.|||||||
MT066156.1     12019 GCTTTCCATGCAGGGTGCTGTAGACATAAACAAGCTTTGTGAAGAAATGC  12068

AY323977.2     11999 TCGATAACCGTGCTACTCTTCAGGCTATTGCTTCAGAATTTAGTTCTTTA  12048
                     |.||.|||.|.||.||..|.||.|||||.||.|||||.||||||||..|.
MT066156.1     12069 TGGACAACAGGGCAACCTTACAAGCTATAGCCTCAGAGTTTAGTTCCCTT  12118

AY323977.2     12049 CCATCATATGCCGCTTATGCCACTGCCCAGGAGGCCTATGAGCAGGCTGT  12098
                     |||||||||||.||||.|||.|||||.||.||.||.||||||||||||||
MT066156.1     12119 CCATCATATGCAGCTTTTGCTACTGCTCAAGAAGCTTATGAGCAGGCTGT  12168

AY323977.2     12099 AGCTAATGGTGATTCTGAAGTCGTTCTCAAAAAGTTAAAGAAATCTTTGA  12148
                     .||||||||||||||||||||.|||||.||||||||.|||||.|||||||
MT066156.1     12169 TGCTAATGGTGATTCTGAAGTTGTTCTTAAAAAGTTGAAGAAGTCTTTGA  12218

AY323977.2     12149 ATGTGGCTAAATCTGAGTTTGACCGTGATGCTGCCATGCAACGCAAGTTG  12198
                     ||||||||||||||||.||||||||||||||.|||||||||||.||||||
MT066156.1     12219 ATGTGGCTAAATCTGAATTTGACCGTGATGCAGCCATGCAACGTAAGTTG  12268

AY323977.2     12199 GAAAAGATGGCAGATCAGGCTATGACCCAAATGTACAAACAGGCAAGATC  12248
                     |||||||||||.|||||.|||||||||||||||||.||||||||.|||||
MT066156.1     12269 GAAAAGATGGCTGATCAAGCTATGACCCAAATGTATAAACAGGCTAGATC  12318

AY323977.2     12249 TGAGGACAAGAGGGCAAAAGTAACTAGTGCTATGCAAACAATGCTCTTCA  12298
                     |||||||||||||||||||||.||||||||||||||.||||||||.||||
MT066156.1     12319 TGAGGACAAGAGGGCAAAAGTTACTAGTGCTATGCAGACAATGCTTTTCA  12368

AY323977.2     12299 CTATGCTTAGGAAGCTTGATAATGATGCACTTAACAACATTATCAACAAT  12348
                     ||||||||||.|||.|.||||||||||||||.||||||||||||||||||
MT066156.1     12369 CTATGCTTAGAAAGTTGGATAATGATGCACTCAACAACATTATCAACAAT  12418

AY323977.2     12349 GCGCGTGATGGTTGTGTTCCACTCAACATCATACCATTGACTACAGCAGC  12398
                     ||..|.||||||||||||||..|.|||||.|||||..|.||.||||||||
MT066156.1     12419 GCAAGAGATGGTTGTGTTCCCTTGAACATAATACCTCTTACAACAGCAGC  12468

AY323977.2     12399 CAAACTCATGGTTGTTGTCCCTGATTATGGTACCTACAAGAACACTTGTG  12448
                     ||||||.||||||||..|.||.||.|||...||.||.||.||.||.||||
MT066156.1     12469 CAAACTAATGGTTGTCATACCAGACTATAACACATATAAAAATACGTGTG  12518

AY323977.2     12449 ATGGTAACACCTTTACATATGCATCTGCACTCTGGGAAATCCAGCAAGTT  12498
                     ||||||..||.|||||.||||||||.|||.|.|||||||||||.||.|||
MT066156.1     12519 ATGGTACAACATTTACTTATGCATCAGCATTGTGGGAAATCCAACAGGTT  12568

AY323977.2     12499 GTTGATGCGGATAGCAAGATTGTTCAACTTAGTGAAATTAACATGGACAA  12548
                     ||.|||||.|||||.||.||||||||||||||||||||||..||||||||
MT066156.1     12569 GTAGATGCAGATAGTAAAATTGTTCAACTTAGTGAAATTAGTATGGACAA  12618

AY323977.2     12549 TTCACCAAATTTGGCTTGGCCTCTTATTGTTACAGCTCTAAGAGCCAACT  12598
                     ||||||.|||||.||.||||||||||||||.||||||.||||.|||||.|
MT066156.1     12619 TTCACCTAATTTAGCATGGCCTCTTATTGTAACAGCTTTAAGGGCCAATT  12668

AY323977.2     12599 CAGCTGTTAAACTACAGAATAATGAACTGAGTCCAGTAGCACTACGACAG  12648
                     |.|||||.|||.|||||||||||||.||.|||||.||.||||||||||||
MT066156.1     12669 CTGCTGTCAAATTACAGAATAATGAGCTTAGTCCTGTTGCACTACGACAG  12718

AY323977.2     12649 ATGTCCTGTGCGGCTGGTACCACACAAACAGCTTGTACTGATGACAATGC  12698
                     |||||.|||||.||.|||||.||||||||.|||||.||||||||||||||
MT066156.1     12719 ATGTCTTGTGCTGCCGGTACTACACAAACTGCTTGCACTGATGACAATGC  12768

AY323977.2     12699 ACTTGCCTACTATAACAATTCGAAGGGAGGTAGGTTTGTGCTGGCATTAC  12748
                     ..|.||.|||||.||||...|.|||||||||||||||||.||.|||.|..
MT066156.1     12769 GTTAGCTTACTACAACACAACAAAGGGAGGTAGGTTTGTACTTGCACTGT  12818

AY323977.2     12749 TATCAGACCACCAAGATCTCAAATGGGCTAGATTCCCTAAGAGTGATGGT  12798
                     ||||.||....||.|||.|.|||||||||||||||||||||||||||||.
MT066156.1     12819 TATCCGATTTACAGGATTTGAAATGGGCTAGATTCCCTAAGAGTGATGGA  12868

AY323977.2     12799 ACAGGTACAATTTACACAGAACTGGAACCACCTTGTAGGTTTGTTACAGA  12848
                     ||.|||||.||.||.|||||||||||||||||||||||||||||||||||
MT066156.1     12869 ACTGGTACTATCTATACAGAACTGGAACCACCTTGTAGGTTTGTTACAGA  12918

AY323977.2     12849 CACACCAAAAGGGCCTAAAGTGAAATACTTGTACTTCATCAAAGGCTTAA  12898
                     ||||||.|||||.|||||||||||.||.||.|||||.||.|||||.||||
MT066156.1     12919 CACACCTAAAGGTCCTAAAGTGAAGTATTTATACTTTATTAAAGGATTAA  12968

AY323977.2     12899 ACAACCTAAATAGAGGTATGGTGCTGGGCAGTTTAGCTGCTACAGTACGT  12948
                     ||||||||||||||||||||||.||.||.|||||||||||.|||||||||
MT066156.1     12969 ACAACCTAAATAGAGGTATGGTACTTGGTAGTTTAGCTGCCACAGTACGT  13018

AY323977.2     12949 CTTCAGGCTGGAAATGCTACAGAAGTACCTGCCAATTCAACTGTGCTTTC  12998
                     ||.||.|||||.|||||.||||||||.|||||||||||||||||..|.||
MT066156.1     13019 CTACAAGCTGGTAATGCAACAGAAGTGCCTGCCAATTCAACTGTATTATC  13068

AY323977.2     12999 CTTCTGTGCTTTTGCAGTAGACCCTGCTAAAGCATATAAGGATTACCTAG  13048
                     .||||||||||||||.|||||..||||||||||.||.||.|||||.||||
MT066156.1     13069 TTTCTGTGCTTTTGCTGTAGATGCTGCTAAAGCTTACAAAGATTATCTAG  13118

AY323977.2     13049 CAAGTGGAGGACAACCAATCACCAACTGTGTGAAGATGTTGTGTACACAC  13098
                     |.|||||.||||||||||||||.||.|||||.||||||||||||||||||
MT066156.1     13119 CTAGTGGGGGACAACCAATCACTAATTGTGTTAAGATGTTGTGTACACAC  13168

AY323977.2     13099 ACTGGTACAGGACAGGCAATTACTGTAACACCAGAAGCTAACATGGACCA  13148
                     ||||||||.||.||||||||.||.||.|||||.|||||.||.|||||.||
MT066156.1     13169 ACTGGTACTGGTCAGGCAATAACAGTTACACCGGAAGCCAATATGGATCA  13218

AY323977.2     13149 AGAGTCCTTTGGTGGTGCTTCATGTTGTCTGTATTGTAGATGCCACATTG  13198
                     |||.||||||||||||||.||.|||||||||||.||..|.||||||||.|
MT066156.1     13219 AGAATCCTTTGGTGGTGCATCGTGTTGTCTGTACTGCCGTTGCCACATAG  13268

AY323977.2     13199 ACCATCCAAATCCTAAAGGATTCTGTGACTTGAAAGGTAAGTACGTCCAA  13248
                     |.||||||||||||||||||||.||||||||.|||||||||||.||.|||
MT066156.1     13269 ATCATCCAAATCCTAAAGGATTTTGTGACTTAAAAGGTAAGTATGTACAA  13318

AY323977.2     13249 ATACCTACCACTTGTGCTAATGACCCAGTGGGTTTTACACTTAGAAACAC  13298
                     ||||||||.|||||||||||||||||.||||||||||||||||.||||||
MT066156.1     13319 ATACCTACAACTTGTGCTAATGACCCTGTGGGTTTTACACTTAAAAACAC  13368

AY323977.2     13299 AGTCTGTACCGTCTGCGGAATGTGGAAAGGTTATGGCTGTAGTTGTGACC  13348
                     ||||||||||||||||||.|||||||||||||||||||||||||||||.|
MT066156.1     13369 AGTCTGTACCGTCTGCGGTATGTGGAAAGGTTATGGCTGTAGTTGTGATC  13418

AY323977.2     13349 AACTCCGCGAACCCTTGATGCAGTCTGCGGATGCATCAACGTTTTTAAAC  13398
                     ||||||||||||||.||.|.|||||.||.||||||..|.|||||||||||
MT066156.1     13419 AACTCCGCGAACCCATGCTTCAGTCAGCTGATGCACAATCGTTTTTAAAC  13468

AY323977.2     13399 GGGTTTGCGGTGTAAGTGCAGCCCGTCTTACACCGTGCGGCACAGGCACT  13448
                     ||||||||||||||||||||||||||||||||||||||||||||||||||
MT066156.1     13469 GGGTTTGCGGTGTAAGTGCAGCCCGTCTTACACCGTGCGGCACAGGCACT  13518

AY323977.2     13449 AGTACTGATGTCGTCTACAGGGCTTTTGATATTTACAACGAAAAAGTTGC  13498
                     ||||||||||||||.||||||||||||||.||.|||||.||.|||||.||
MT066156.1     13519 AGTACTGATGTCGTATACAGGGCTTTTGACATCTACAATGATAAAGTAGC  13568

AY323977.2     13499 TGGTTTTGCAAAGTTCCTAAAAACTAATTGCTGTCGCTTCCAGGAGAAGG  13548
                     |||||||||.||.|||||||||||||||||.|||||||||||.||.||||
MT066156.1     13569 TGGTTTTGCTAAATTCCTAAAAACTAATTGTTGTCGCTTCCAAGAAAAGG  13618

AY323977.2     13549 ATGAGGAAGGCAATTTATTAGACTCTTACTTTGTAGTTAAGAGGCATACT  13598
                     |.||.||.|.|||||||.|.||.||||||||||||||||||||.||.|||
MT066156.1     13619 ACGAAGATGACAATTTAATTGATTCTTACTTTGTAGTTAAGAGACACACT  13668

AY323977.2     13599 ATGTCTAACTACCAACATGAAGAGACTATTTATAACTTGGTTAAAGATTG  13648
                     .|.||||||||||||||||||||.||.||||||||.||..||||.|||||
MT066156.1     13669 TTCTCTAACTACCAACATGAAGAAACAATTTATAATTTACTTAAGGATTG  13718

AY323977.2     13649 TCCAGCGGTTGCTGTCCATGACTTTTTCAAGTTTAGAGTAGATGGTGACA  13698
                     ||||||.||||||...||||||||.||.|||||||||.||||.|||||||
MT066156.1     13719 TCCAGCTGTTGCTAAACATGACTTCTTTAAGTTTAGAATAGACGGTGACA  13768

AY323977.2     13699 TGGTACCACATATATCACGTCAGCGTCTAACTAAATACACAATGGCTGAT  13748
                     ||||||||||||||||||||||.|||||.|||||||||||||||||.||.
MT066156.1     13769 TGGTACCACATATATCACGTCAACGTCTTACTAAATACACAATGGCAGAC  13818

AY323977.2     13749 TTAGTCTATGCTCTACGTCATTTTGATGAGGGTAATTGTGATACATTAAA  13798
                     .|.|||||||||.||.|.|||||||||||.|||||||||||.||||||||
MT066156.1     13819 CTCGTCTATGCTTTAAGGCATTTTGATGAAGGTAATTGTGACACATTAAA  13868

AY323977.2     13799 AGAAATACTCGTCACATACAATTGCTGTGATGATGATTATTTCAATAAGA  13848
                     |||||||||.||||||||||||||.|||||||||||||||||||||||.|
MT066156.1     13869 AGAAATACTTGTCACATACAATTGTTGTGATGATGATTATTTCAATAAAA  13918

AY323977.2     13849 AGGATTGGTATGACTTCGTAGAGAATCCTGACATCTTACGCGTATATGCT  13898
                     ||||.||||||||.||.|||||.||.||.||.||.|||||||||||.||.
MT066156.1     13919 AGGACTGGTATGATTTTGTAGAAAACCCAGATATATTACGCGTATACGCC  13968

AY323977.2     13899 AACTTAGGTGAGCGTGTACGCCAATCATTATTAAAGACTGTACAATTCTG  13948
                     |||||||||||.||||||||||||.|.||.|||||.||.|||||||||||
MT066156.1     13969 AACTTAGGTGAACGTGTACGCCAAGCTTTGTTAAAAACAGTACAATTCTG  14018

AY323977.2     13949 CGATGCTATGCGTGATGCAGGCATTGTAGGCGTACTGACATTAGATAATC  13998
                     .|||||.|||||..||||.||.|||||.||.|||||||||||||||||||
MT066156.1     14019 TGATGCCATGCGAAATGCTGGTATTGTTGGTGTACTGACATTAGATAATC  14068

AY323977.2     13999 AGGATCTTAATGGGAACTGGTACGATTTCGGTGATTTCGTACAAGTAGCA  14048
                     |.|||||.|||||.||||||||.|||||||||||||||.|||||....|.
MT066156.1     14069 AAGATCTCAATGGTAACTGGTATGATTTCGGTGATTTCATACAAACCACG  14118

AY323977.2     14049 CCAGGCTGCGGAGTTCCTATTGTGGATTCATATTACTCATTGCTGATGCC  14098
                     |||||..|.|||||||||.||||.|||||.|||||.||||||.|.|||||
MT066156.1     14119 CCAGGTAGTGGAGTTCCTGTTGTAGATTCTTATTATTCATTGTTAATGCC  14168

AY323977.2     14099 CATCCTCACTTTGACTAGGGCATTGGCTGCTGAGTCCCATATGGATGCTG  14148
                     .||..|.||.|||||.|||||.||..||||.|||||.|||.|.||..|||
MT066156.1     14169 TATATTAACCTTGACCAGGGCTTTAACTGCAGAGTCACATGTTGACACTG  14218

AY323977.2     14149 ATCTCGCAAAACCACTTATTAAGTGGGATTTGCTGAAATATGATTTTACG  14198
                     |..|..||||.||....|||||||||||||||.|.||||||||.||.|||
MT066156.1     14219 ACTTAACAAAGCCTTACATTAAGTGGGATTTGTTAAAATATGACTTCACG  14268

AY323977.2     14199 GAAGAGAGACTTTGTCTCTTCGACCGTTATTTTAAATATTGGGACCAGAC  14248
                     ||||||||..|....|||||.|||||||||||||||||||||||.|||||
MT066156.1     14269 GAAGAGAGGTTAAAACTCTTTGACCGTTATTTTAAATATTGGGATCAGAC  14318

AY323977.2     14249 ATACCATCCCAATTGTATTAACTGTTTGGATGATAGGTGTATCCTTCATT  14298
                     ||||||.||.||||||.||||||||||||||||.||.||.||.||.||||
MT066156.1     14319 ATACCACCCAAATTGTGTTAACTGTTTGGATGACAGATGCATTCTGCATT  14368

AY323977.2     14299 GTGCAAACTTTAATGTGTTATTTTCTACTGTGTTTCCACCTACAAGTTTT  14348
                     ||||||||||||||||.|||||.|||||.|||||.|||||||||||||||
MT066156.1     14369 GTGCAAACTTTAATGTTTTATTCTCTACAGTGTTCCCACCTACAAGTTTT  14418

AY323977.2     14349 GGACCACTAGTAAGAAAAATATTTGTAGATGGTGTTCCTTTTGTTGTTTC  14398
                     |||||||||||.||||||||||||||.|||||||||||.|||||.|||||
MT066156.1     14419 GGACCACTAGTGAGAAAAATATTTGTTGATGGTGTTCCATTTGTAGTTTC  14468

AY323977.2     14399 AACTGGATACCATTTTCGTGAGTTAGGAGTCGTACATAATCAGGATGTAA  14448
                     ||||||||||||.||..|.|||.||||.||.|||||||||||||||||||
MT066156.1     14469 AACTGGATACCACTTCAGAGAGCTAGGTGTTGTACATAATCAGGATGTAA  14518

AY323977.2     14449 ACTTACATAGCTCGCGTCTCAGTTTCAAGGAACTTTTAGTGTATGCTGCT  14498
                     |||||||||||||..|.||.|||||.||||||.|..|.||||||||||||
MT066156.1     14519 ACTTACATAGCTCTAGACTTAGTTTTAAGGAATTACTTGTGTATGCTGCT  14568

AY323977.2     14499 GATCCAGCTATGCATGCAGCTTCTGGCAATTTATTGCTAGATAAACGCAC  14548
                     ||.||.||||||||.||.||||||||.|||.||||.||||||||||||||
MT066156.1     14569 GACCCTGCTATGCACGCTGCTTCTGGTAATCTATTACTAGATAAACGCAC  14618

AY323977.2     14549 TACATGCTTTTCAGTAGCTGCACTAACAAACAATGTTGCTTTTCAAACTG  14598
                     |||.||||||||||||||||||||.||.||||||||||||||||||||||
MT066156.1     14619 TACGTGCTTTTCAGTAGCTGCACTTACTAACAATGTTGCTTTTCAAACTG  14668

AY323977.2     14599 TCAAACCCGGTAATTTTAATAAAGACTTTTATGACTTTGCTGTGTCTAAA  14648
                     |||||||||||||||||||.||||||||.||||||||||||||||||||.
MT066156.1     14669 TCAAACCCGGTAATTTTAACAAAGACTTCTATGACTTTGCTGTGTCTAAG  14718

AY323977.2     14649 GGTTTCTTTAAGGAAGGAAGTTCTGTTGAACTAAAACACTTCTTCTTTGC  14698
                     ||||||||||||||||||||||||||||||.|||||||||||||||||||
MT066156.1     14719 GGTTTCTTTAAGGAAGGAAGTTCTGTTGAATTAAAACACTTCTTCTTTGC  14768

AY323977.2     14699 TCAGGATGGCAACGCTGCTATCAGTGATTATGACTATTATCGTTATAATC  14748
                     |||||||||.||.|||||||||||.|||||||||||.|||||||||||||
MT066156.1     14769 TCAGGATGGTAATGCTGCTATCAGCGATTATGACTACTATCGTTATAATC  14818

AY323977.2     14749 TGCCAACAATGTGTGATATCAGACAACTCCTATTCGTAGTTGAAGTTGTT  14798
                     |.||||||||||||||||||||||||||.|||||.|||||||||||||||
MT066156.1     14819 TACCAACAATGTGTGATATCAGACAACTACTATTTGTAGTTGAAGTTGTT  14868

AY323977.2     14799 GATAAATACTTTGATTGTTACGATGGTGGCTGTATTAATGCCAACCAAGT  14848
                     |||||.|||||||||||||||||||||||||||||||||||.||||||||
MT066156.1     14869 GATAAGTACTTTGATTGTTACGATGGTGGCTGTATTAATGCTAACCAAGT  14918

AY323977.2     14849 AATCGTTAACAATCTGGATAAATCAGCTGGTTTCCCATTTAATAAATGGG  14898
                     .|||||.|||||.||.||.||||||||||||||.||||||||||||||||
MT066156.1     14919 CATCGTCAACAACCTAGACAAATCAGCTGGTTTTCCATTTAATAAATGGG  14968

AY323977.2     14899 GTAAGGCTAGACTTTATTATGACTCAATGAGTTATGAGGATCAAGATGCA  14948
                     ||||||||||||||||||||||.|||||||||||||||||||||||||||
MT066156.1     14969 GTAAGGCTAGACTTTATTATGATTCAATGAGTTATGAGGATCAAGATGCA  15018

AY323977.2     14949 CTTTTCGCGTATACTAAGCGTAATGTCATCCCTACTATAACTCAAATGAA  14998
                     ||||||||.|||||.||.||||||||||||||||||||||||||||||||
MT066156.1     15019 CTTTTCGCATATACAAAACGTAATGTCATCCCTACTATAACTCAAATGAA  15068

AY323977.2     14999 TCTTAAGTATGCCATTAGTGCAAAGAATAGAGCTCGCACCGTAGCTGGTG  15048
                     ||||||||||||||||||||||||||||||||||||||||||||||||||
MT066156.1     15069 TCTTAAGTATGCCATTAGTGCAAAGAATAGAGCTCGCACCGTAGCTGGTG  15118

AY323977.2     15049 TCTCTATCTGTAGTACTATGACAAATAGACAGTTTCATCAGAAATTATTG  15098
                     ||||||||||||||||||||||.|||||||||||||||||.|||||||||
MT066156.1     15119 TCTCTATCTGTAGTACTATGACCAATAGACAGTTTCATCAAAAATTATTG  15168

AY323977.2     15099 AAGTCAATAGCCGCCACTAGAGGAGCTACTGTGGTAATTGGAACAAGCAA  15148
                     ||.|||||||||||||||||||||||||||||.|||||||||||||||||
MT066156.1     15169 AAATCAATAGCCGCCACTAGAGGAGCTACTGTAGTAATTGGAACAAGCAA  15218

AY323977.2     15149 GTTTTACGGTGGCTGGCATAATATGTTAAAAACTGTTTACAGTGATGTAG  15198
                     .||.||.|||||.|||||.||.|||||||||||||||||.||||||||||
MT066156.1     15219 ATTCTATGGTGGTTGGCACAACATGTTAAAAACTGTTTATAGTGATGTAG  15268

AY323977.2     15199 AAACTCCACACCTTATGGGTTGGGATTATCCAAAATGTGACAGAGCCATG  15248
                     |||..||.|||||||||||||||||||||||.||||||||.|||||||||
MT066156.1     15269 AAAACCCTCACCTTATGGGTTGGGATTATCCTAAATGTGATAGAGCCATG  15318

AY323977.2     15249 CCTAACATGCTTAGGATAATGGCCTCTCTTGTTCTTGCTCGCAAACATAA  15298
                     ||||||||||||||.||.||||||||.||||||||||||||||||||||.
MT066156.1     15319 CCTAACATGCTTAGAATTATGGCCTCACTTGTTCTTGCTCGCAAACATAC  15368

AY323977.2     15299 CACTTGCTGTAACTTATCACACCGTTTCTACAGGTTAGCTAACGAGTGTG  15348
                     .||.||.||||.|||.||||||||||||||.||.||||||||.|||||||
MT066156.1     15369 AACGTGTTGTAGCTTGTCACACCGTTTCTATAGATTAGCTAATGAGTGTG  15418

AY323977.2     15349 CGCAAGTATTAAGTGAGATGGTCATGTGTGGCGGCTCACTATATGTTAAA  15398
                     |.||||||||.|||||.|||||||||||||||||.|||||||||||||||
MT066156.1     15419 CTCAAGTATTGAGTGAAATGGTCATGTGTGGCGGTTCACTATATGTTAAA  15468

AY323977.2     15399 CCAGGTGGAACATCATCCGGTGATGCTACAACTGCTTATGCTAATAGTGT  15448
                     |||||||||||.|||||.||.|||||.|||||||||||||||||||||||
MT066156.1     15469 CCAGGTGGAACCTCATCAGGAGATGCCACAACTGCTTATGCTAATAGTGT  15518

AY323977.2     15449 CTTTAACATTTGTCAAGCTGTTACAGCCAATGTAAATGCACTTCTTTCAA  15498
                     .||||||||||||||||||||.||.||||||||.|||||||||.|.||.|
MT066156.1     15519 TTTTAACATTTGTCAAGCTGTCACGGCCAATGTTAATGCACTTTTATCTA  15568

AY323977.2     15499 CTGATGGTAATAAGATAGCTGACAAGTATGTCCGCAATCTACAACACAGG  15548
                     ||||||||||.||.||.||.||.|||||||||||||||.||||||||||.
MT066156.1     15569 CTGATGGTAACAAAATTGCCGATAAGTATGTCCGCAATTTACAACACAGA  15618

AY323977.2     15549 CTCTATGAGTGTCTCTATAGAAATAGGGATGTTGATCATGAATTCGTGGA  15598
                     ||.|||||||||||||||||||||||.||||||||....||.||.|||.|
MT066156.1     15619 CTTTATGAGTGTCTCTATAGAAATAGAGATGTTGACACAGACTTTGTGAA  15668

AY323977.2     15599 TGAGTTTTACGCTTACCTGCGTAAACATTTCTCCATGATGATTCTTTCTG  15648
                     ||||||||||||.||..||||||||||||||||.||||||||.||.||||
MT066156.1     15669 TGAGTTTTACGCATATTTGCGTAAACATTTCTCAATGATGATACTCTCTG  15718

AY323977.2     15649 ATGATGCCGTTGTGTGCTATAACAGTAACTATGCGGCTCAAGGTTTAGTA  15698
                     |.|||||.||||||||.|..||.||.|..|||||..||||||||.||||.
MT066156.1     15719 ACGATGCTGTTGTGTGTTTCAATAGCACTTATGCATCTCAAGGTCTAGTG  15768

AY323977.2     15699 GCTAGCATTAAGAACTTTAAGGCAGTTCTTTATTATCAAAATAATGTGTT  15748
                     ||||||||.||||||||||||.|||||||||||||||||||.|||||.||
MT066156.1     15769 GCTAGCATAAAGAACTTTAAGTCAGTTCTTTATTATCAAAACAATGTTTT  15818

AY323977.2     15749 CATGTCTGAGGCAAAATGTTGGACTGAGACTGACCTTACTAAAGGACCTC  15798
                     .||||||||.||||||||||||||||||||||||||||||||||||||||
MT066156.1     15819 TATGTCTGAAGCAAAATGTTGGACTGAGACTGACCTTACTAAAGGACCTC  15868

AY323977.2     15799 ACGAATTTTGCTCACAGCATACAATGCTAGTTAAACAAGGAGATGATTAC  15848
                     |.|||||||||||.||.||||||||||||||||||||.||.||||||||.
MT066156.1     15869 ATGAATTTTGCTCTCAACATACAATGCTAGTTAAACAGGGTGATGATTAT  15918

AY323977.2     15849 GTGTACCTGCCTTACCCAGATCCATCAAGAATATTAGGCGCAGGCTGTTT  15898
                     ||||||||.|||||||||||||||||||||||..||||.||.||||||||
MT066156.1     15919 GTGTACCTTCCTTACCCAGATCCATCAAGAATCCTAGGGGCCGGCTGTTT  15968

AY323977.2     15899 TGTCGATGATATTGTCAAAACAGATGGTACACTTATGATTGAAAGGTTCG  15948
                     |||.||||||||.||.|||||||||||||||||||||||||||.||||||
MT066156.1     15969 TGTAGATGATATCGTAAAAACAGATGGTACACTTATGATTGAACGGTTCG  16018

AY323977.2     15949 TGTCACTGGCTATTGATGCTTACCCACTTACAAAACATCCTAATCAGGAG  15998
                     ||||..|.|||||.|||||||||||||||||.||||||||||||||||||
MT066156.1     16019 TGTCTTTAGCTATAGATGCTTACCCACTTACTAAACATCCTAATCAGGAG  16068

AY323977.2     15999 TATGCTGATGTCTTTCACTTGTATTTACAATACATTAGAAAGTTACATGA  16048
                     |||||||||||||||||.|||||.|||||||||||.||||||.|||||||
MT066156.1     16069 TATGCTGATGTCTTTCATTTGTACTTACAATACATAAGAAAGCTACATGA  16118

AY323977.2     16049 TGAGCTTACTGGCCACATGTTGGACATGTATTCCGTAATGCTAACTAATG  16098
                     ||||.|.||.||.||||||||.|||||||||||.||.|||||.|||||||
MT066156.1     16119 TGAGTTAACAGGACACATGTTAGACATGTATTCTGTTATGCTTACTAATG  16168

AY323977.2     16099 ATAACACCTCACGGTACTGGGAACCTGAGTTTTATGAGGCTATGTACACA  16148
                     |||||||.|||.||||.|||||||||||||||||||||||||||||||||
MT066156.1     16169 ATAACACTTCAAGGTATTGGGAACCTGAGTTTTATGAGGCTATGTACACA  16218

AY323977.2     16149 CCACATACAGTCTTGCAGGCTGTAGGTGCTTGTGTATTGTGCAATTCACA  16198
                     ||.|||||||||||.||||||||.||.||||||||..|.|||||||||||
MT066156.1     16219 CCGCATACAGTCTTACAGGCTGTTGGGGCTTGTGTTCTTTGCAATTCACA  16268

AY323977.2     16199 GACTTCACTTCGTTGCGGTGCCTGTATTAGGAGACCATTCCTATGTTGCA  16248
                     |||||||.|..|.||.|||||.||.||..|.|||||||||.|||||||.|
MT066156.1     16269 GACTTCATTAAGATGTGGTGCTTGCATACGTAGACCATTCTTATGTTGTA  16318

AY323977.2     16249 AGTGCTGCTATGACCATGTCATTTCAACATCACACAAATTAGTGTTGTCT  16298
                     |.|||||.||.|||||||||||.|||||||||||.||||||||.||||||
MT066156.1     16319 AATGCTGTTACGACCATGTCATATCAACATCACATAAATTAGTCTTGTCT  16368

AY323977.2     16299 GTTAATCCCTATGTTTGCAATGCCCCAGGTTGTGATGTCACTGATGTGAC  16348
                     ||||||||.||||||||||||||.|||||||||||||||||.||||||||
MT066156.1     16369 GTTAATCCGTATGTTTGCAATGCTCCAGGTTGTGATGTCACAGATGTGAC  16418

AY323977.2     16349 ACAACTGTATCTAGGAGGTATGAGCTATTATTGCAAGTCACATAAGCCTC  16398
                     .|||||.||..||||||||||||||||||||||.||.||||||||.||.|
MT066156.1     16419 TCAACTTTACTTAGGAGGTATGAGCTATTATTGTAAATCACATAAACCAC  16468

AY323977.2     16399 CCATTAGTTTTCCATTATGTGCTAATGGTCAGGTTTTTGGTTTATACAAA  16448
                     ||||||||||||||||.|||||||||||.||.||||||||||||||.|||
MT066156.1     16469 CCATTAGTTTTCCATTGTGTGCTAATGGACAAGTTTTTGGTTTATATAAA  16518

AY323977.2     16449 AACACATGTGTAGGCAGTGACAATGTCACTGACTTCAATGCGATAGCAAC  16498
                     ||.||||||||.||.||.||.|||||.||||||||.|||||.||.|||||
MT066156.1     16519 AATACATGTGTTGGTAGCGATAATGTTACTGACTTTAATGCAATTGCAAC  16568

AY323977.2     16499 ATGTGATTGGACTAATGCTGGCGATTACATACTTGCCAACACTTGTACTG  16548
                     ||||||.|||||.||||||||.||||||||..|.||.|||||.|||||||
MT066156.1     16569 ATGTGACTGGACAAATGCTGGTGATTACATTTTAGCTAACACCTGTACTG  16618

AY323977.2     16549 AGAGACTCAAGCTTTTCGCAGCAGAAACGCTCAAAGCCACTGAGGAAACA  16598
                     |.||||||||||||||.||||||||||||||||||||.||||||||.|||
MT066156.1     16619 AAAGACTCAAGCTTTTTGCAGCAGAAACGCTCAAAGCTACTGAGGAGACA  16668

AY323977.2     16599 TTTAAGCTGTCATATGGTATTGCCACTGTACGCGAAGTACTCTCTGACAG  16648
                     |||||.|||||.|||||||||||.||||||||.|||||.||.||||||||
MT066156.1     16669 TTTAAACTGTCTTATGGTATTGCTACTGTACGTGAAGTGCTGTCTGACAG  16718

AY323977.2     16649 AGAATTGCATCTTTCATGGGAGGTTGGAAAACCTAGACCACCATTGAACA  16698
                     ||||||.||||||||||||||.|||||.|||||||||||||||.|.|||.
MT066156.1     16719 AGAATTACATCTTTCATGGGAAGTTGGTAAACCTAGACCACCACTTAACC  16768

AY323977.2     16699 GAAACTATGTCTTTACTGGTTACCGTGTAACTAAAAATAGTAAAGTACAG  16748
                     ||||.|||||||||||||||||.||||||||||||||.|||||||||||.
MT066156.1     16769 GAAATTATGTCTTTACTGGTTATCGTGTAACTAAAAACAGTAAAGTACAA  16818

AY323977.2     16749 ATTGGAGAGTACACCTTTGAAAAAGGTGACTATGGTGATGCTGTTGTGTA  16798
                     ||.||||||||||||||||||||||||||||||||||||||||||||.||
MT066156.1     16819 ATAGGAGAGTACACCTTTGAAAAAGGTGACTATGGTGATGCTGTTGTTTA  16868

AY323977.2     16799 CAGAGGTACTACGACATACAAGTTGAATGTTGGTGATTACTTTGTGTTGA  16848
                     |.|||||||.||.||.|||||.||.||||||||||||||.||||||.|||
MT066156.1     16869 CCGAGGTACAACAACTTACAAATTAAATGTTGGTGATTATTTTGTGCTGA  16918

AY323977.2     16849 CATCTCACACTGTAATGCCACTTAGTGCACCTACTCTAGTGCCACAAGAG  16898
                     ||||.||.||.|||||||||.|.|||||||||||.|||||||||||||||
MT066156.1     16919 CATCACATACAGTAATGCCATTAAGTGCACCTACACTAGTGCCACAAGAG  16968

AY323977.2     16899 CACTATGTGAGAATTACTGGCTTGTACCCAACACTCAACATCTCAGATGA  16948
                     ||||||||.||||||||||||||.||||||||||||||.|||||||||||
MT066156.1     16969 CACTATGTTAGAATTACTGGCTTATACCCAACACTCAATATCTCAGATGA  17018

AY323977.2     16949 GTTTTCTAGCAATGTTGCAAATTATCAAAAGGTCGGCATGCAAAAGTACT  16998
                     |||||||||||||||||||||||||||||||||.||.|||||||||||.|
MT066156.1     17019 GTTTTCTAGCAATGTTGCAAATTATCAAAAGGTTGGTATGCAAAAGTATT  17068

AY323977.2     16999 CTACACTCCAAGGACCACCTGGTACTGGTAAGAGTCATTTTGCCATCGGA  17048
                     ||||||||||.||||||||||||||||||||||||||||||||.||.||.
MT066156.1     17069 CTACACTCCAGGGACCACCTGGTACTGGTAAGAGTCATTTTGCTATTGGC  17118

AY323977.2     17049 CTTGCTCTCTATTACCCATCTGCTCGCATAGTGTATACGGCATGCTCTCA  17098
                     ||.||||||||.|||||.||||||||||||||||||||.||.||||||||
MT066156.1     17119 CTAGCTCTCTACTACCCTTCTGCTCGCATAGTGTATACAGCTTGCTCTCA  17168

AY323977.2     17099 TGCAGCTGTTGATGCCCTATGTGAAAAGGCATTAAAATATTTGCCCATAG  17148
                     |||.|||||||||||.||||||||.||||||||||||||||||||.||||
MT066156.1     17169 TGCCGCTGTTGATGCACTATGTGAGAAGGCATTAAAATATTTGCCTATAG  17218

AY323977.2     17149 ATAAATGTAGTAGAATCATACCTGCGCGTGCGCGCGTAGAGTGTTTTGAT  17198
                     ||||||||||||||||.||||||||.|||||.||.|||||||||||||||
MT066156.1     17219 ATAAATGTAGTAGAATTATACCTGCACGTGCTCGTGTAGAGTGTTTTGAT  17268

AY323977.2     17199 AAATTCAAAGTGAATTCAACACTAGAACAGTATGTTTTCTGCACTGTAAA  17248
                     |||||||||||||||||||||.|||||||||||||.||.||.||||||||
MT066156.1     17269 AAATTCAAAGTGAATTCAACATTAGAACAGTATGTCTTTTGTACTGTAAA  17318

AY323977.2     17249 TGCATTGCCAGAAACAACTGCTGACATTGTAGTCTTTGATGAAATCTCTA  17298
                     |||||||||.||.||.||.||.||.||.||.||||||||||||||.||.|
MT066156.1     17319 TGCATTGCCTGAGACGACAGCAGATATAGTTGTCTTTGATGAAATTTCAA  17368

AY323977.2     17299 TGGCTACTAATTATGACTTGAGTGTTGTCAATGCTAGACTTCGTGCAAAA  17348
                     ||||.||.||||||||.|||||||||||||||||.|||.|.|||||.||.
MT066156.1     17369 TGGCCACAAATTATGATTTGAGTGTTGTCAATGCCAGATTACGTGCTAAG  17418

AY323977.2     17349 CACTACGTCTATATTGGCGATCCTGCTCAATTACCAGCCCCCCGCACATT  17398
                     |||||.||.||.||||||||.||||||||||||||.||.||.||||||||
MT066156.1     17419 CACTATGTGTACATTGGCGACCCTGCTCAATTACCTGCACCACGCACATT  17468

AY323977.2     17399 GCTGACTAAAGGCACACTAGAACCAGAATATTTTAATTCAGTGTGCAGAC  17448
                     |||.|||||.|||||||||||||||||||||||.|||||||||||.||||
MT066156.1     17469 GCTAACTAAGGGCACACTAGAACCAGAATATTTCAATTCAGTGTGTAGAC  17518

AY323977.2     17449 TTATGAAAACAATAGGTCCAGACATGTTCCTTGGAACTTGTCGCCGTTGT  17498
                     ||||||||||.||||||||||||||||||||.|||||||||||.||||||
MT066156.1     17519 TTATGAAAACTATAGGTCCAGACATGTTCCTCGGAACTTGTCGGCGTTGT  17568

AY323977.2     17499 CCTGCTGAAATTGTTGACACTGTGAGTGCTTTAGTTTATGACAATAAGCT  17548
                     ||||||||||||||||||||||||||||||||.||||||||.||||||||
MT066156.1     17569 CCTGCTGAAATTGTTGACACTGTGAGTGCTTTGGTTTATGATAATAAGCT  17618

AY323977.2     17549 AAAAGCACACAAGGATAAGTCAGCTCAATGCTTCAAAATGTTCTACAAAG  17598
                     .||||||||.||.||.||.||||||||||||||.||||||||.||.||.|
MT066156.1     17619 TAAAGCACATAAAGACAAATCAGCTCAATGCTTTAAAATGTTTTATAAGG  17668

AY323977.2     17599 GTGTTATTACACATGATGTTTCATCTGCAATCAACAGACCTCAAATAGGC  17648
                     |||||||.||.||||||||||||||||||||.|||||.||.|||||||||
MT066156.1     17669 GTGTTATCACGCATGATGTTTCATCTGCAATTAACAGGCCACAAATAGGC  17718

AY323977.2     17649 GTTGTAAGAGAATTTCTTACACGCAATCCTGCTTGGAGAAAAGCTGTTTT  17698
                     ||.|||||||||||.||||||||.||.||||||||||||||||||||.||
MT066156.1     17719 GTGGTAAGAGAATTCCTTACACGTAACCCTGCTTGGAGAAAAGCTGTCTT  17768

AY323977.2     17699 TATCTCACCTTATAATTCACAGAACGCTGTAGCTTCAAAAATCTTAGGAT  17748
                     |||.||||||||||||||||||||.||||||||.|||||.||.||.|||.
MT066156.1     17769 TATTTCACCTTATAATTCACAGAATGCTGTAGCCTCAAAGATTTTGGGAC  17818

AY323977.2     17749 TGCCTACGCAGACTGTTGATTCATCACAGGGTTCTGAATATGACTATGTC  17798
                     |.||.||.||.||||||||||||||||||||.||.|||||||||||||||
MT066156.1     17819 TACCAACTCAAACTGTTGATTCATCACAGGGCTCAGAATATGACTATGTC  17868

AY323977.2     17799 ATATTCACACAAACTACTGAAACAGCACACTCTTGTAATGTCAACCGCTT  17848
                     ||||||||.|||||.|||||||||||.||||||||||||||.|||.|.||
MT066156.1     17869 ATATTCACTCAAACCACTGAAACAGCTCACTCTTGTAATGTAAACAGATT  17918

AY323977.2     17849 CAATGTGGCTATCACAAGGGCAAAAATTGGCATTTTGTGCATAATGTCTG  17898
                     .|||||.|||||.||.||.||||||.|.|||||..|.|||||||||||||
MT066156.1     17919 TAATGTTGCTATTACCAGAGCAAAAGTAGGCATACTTTGCATAATGTCTG  17968

AY323977.2     17899 ATAGAGATCTTTATGACAAACTGCAATTTACAAGTCTAGAAATACCACGT  17948
                     |||||||.|||||||||||..||||||||||||||||.|||||.||||||
MT066156.1     17969 ATAGAGACCTTTATGACAAGTTGCAATTTACAAGTCTTGAAATTCCACGT  18018

AY323977.2     17949 CGCAATGTGGCTACATTACAAGCAGAAAATGTAACTGGACTTTTTAAGGA  17998
                     .|.||||||||.||.||||||||.|||||||||||.|||||.|||||.||
MT066156.1     18019 AGGAATGTGGCAACTTTACAAGCTGAAAATGTAACAGGACTCTTTAAAGA  18068

AY323977.2     17999 CTGTAGTAAGATCATTACTGGTCTTCATCCTACACAGGCACCTACACACC  18048
                     .|||||||||.|.||.|||||..|.|||||||||||||||||||||||||
MT066156.1     18069 TTGTAGTAAGGTAATCACTGGGTTACATCCTACACAGGCACCTACACACC  18118

AY323977.2     18049 TCAGCGTTGATATAAAGTTCAAGACTGAAGGATTATGTGTTGACATACCA  18098
                     ||||.|||||.|..||.|||||.||||||||.|||||||||||||||||.
MT066156.1     18119 TCAGTGTTGACACTAAATTCAAAACTGAAGGTTTATGTGTTGACATACCT  18168

AY323977.2     18099 GGCATACCAAAGGACATGACCTACCGTAGACTCATCTCTATGATGGGTTT  18148
                     ||||||||.||||||||||||||..|.|||||||||||||||||||||||
MT066156.1     18169 GGCATACCTAAGGACATGACCTATAGAAGACTCATCTCTATGATGGGTTT  18218

AY323977.2     18149 CAAAATGAATTACCAAGTCAATGGTTACCCTAATATGTTTATCACCCGCG  18198
                     .|||||||||||.|||||.||||||||||||||.||||||||||||||||
MT066156.1     18219 TAAAATGAATTATCAAGTTAATGGTTACCCTAACATGTTTATCACCCGCG  18268

AY323977.2     18199 AAGAAGCTATTCGTCACGTTCGTGCGTGGATTGGCTTTGATGTAGAGGGC  18248
                     ||||||||||..|.||.||.|||||.|||||||||||.|||||.|||||.
MT066156.1     18269 AAGAAGCTATAAGACATGTACGTGCATGGATTGGCTTCGATGTCGAGGGG  18318

AY323977.2     18249 TGTCATGCAACTAGAGATGCTGTGGGTACTAACCTACCTCTCCAGCTAGG  18298
                     ||||||||.||||||||.|||||.|||||.||..|||||.|.||||||||
MT066156.1     18319 TGTCATGCTACTAGAGAAGCTGTTGGTACCAATTTACCTTTACAGCTAGG  18368

AY323977.2     18299 ATTTTCTACAGGTGTTAACTTAGTAGCTGTACCGACTGGTTATGTTGACA  18348
                     .||||||||||||||||||.||||.||||||||.||.|||||||||||.|
MT066156.1     18369 TTTTTCTACAGGTGTTAACCTAGTTGCTGTACCTACAGGTTATGTTGATA  18418

AY323977.2     18349 CTGAAAATAACACAGAATTCACCAGAGTTAATGCAAAACCTCCACCAGGT  18398
                     |....|||||.|||||.||..|||||||||.|||.|||||.||.||.||.
MT066156.1     18419 CACCTAATAATACAGATTTTTCCAGAGTTAGTGCTAAACCACCGCCTGGA  18468

AY323977.2     18399 GACCAGTTTAAACATCTTATACCACTCATGTATAAAGGCTTGCCCTGGAA  18448
                     ||.||.||||||||.||.||||||||.|||||.|||||..|.||.|||||
MT066156.1     18469 GATCAATTTAAACACCTCATACCACTTATGTACAAAGGACTTCCTTGGAA  18518

AY323977.2     18449 TGTAGTGCGTATTAAGATAGTACAAATGCTCAGTGATACACTGAAAGGAT  18498
                     ||||||||||||.|||||.|||||||||.|.|||||.|||||.|||....
MT066156.1     18519 TGTAGTGCGTATAAAGATTGTACAAATGTTAAGTGACACACTTAAAAATC  18568

AY323977.2     18499 TGTCAGACAGAGTCGTGTTCGTCCTTTGGGCGCATGGCTTTGAGCTTACA  18548
                     |.||.|||||||||||.||.|||.|.|||||.||||||||||||.|.|||
MT066156.1     18569 TCTCTGACAGAGTCGTATTTGTCTTATGGGCACATGGCTTTGAGTTGACA  18618

AY323977.2     18549 TCAATGAAGTACTTTGTCAAGATTGGACCTGAAAGAACGTGTTGTCTGTG  18598
                     ||.||||||||.|||||.||.||.||||||||..|.||.||||||||.||
MT066156.1     18619 TCTATGAAGTATTTTGTGAAAATAGGACCTGAGCGCACCTGTTGTCTATG  18668

AY323977.2     18599 TGACAAACGTGCAACTTGCTTTTCTACTTCATCAGATACTTATGCCTGCT  18648
                     |||.|.||||||.||.||||||||.|||.|.|||||.|||||||||||.|
MT066156.1     18669 TGATAGACGTGCCACATGCTTTTCCACTGCTTCAGACACTTATGCCTGTT  18718

AY323977.2     18649 GGAATCATTCTGTGGGTTTTGACTATGTCTATAACCCATTTATGATTGAT  18698
                     ||.||||||||.|.||.|||||.||.||||||||.||.||||||||||||
MT066156.1     18719 GGCATCATTCTATTGGATTTGATTACGTCTATAATCCGTTTATGATTGAT  18768

AY323977.2     18699 GTTCAGCAGTGGGGCTTTACGGGTAACCTTCAGAGTAACCATGACCAACA  18748
                     |||||.||.|||||.|||||.||||||||.||.||.||||||||.|...|
MT066156.1     18769 GTTCAACAATGGGGTTTTACAGGTAACCTACAAAGCAACCATGATCTGTA  18818

AY323977.2     18749 TTGCCAGGTACATGGAAATGCACATGTGGCTAGTTGTGATGCTATCATGA  18798
                     |||.||.||.|||||.|||||||||||.||||||||||||||.|||||||
MT066156.1     18819 TTGTCAAGTCCATGGTAATGCACATGTAGCTAGTTGTGATGCAATCATGA  18868

AY323977.2     18799 CTAGATGTTTAGCAGTCCATGAGTGCTTTGTTAAGCGCGTTGATTGGTCT  18848
                     ||||.|||.||||.|||||.|||||||||||||||||.|||||.|||.||
MT066156.1     18869 CTAGGTGTCTAGCTGTCCACGAGTGCTTTGTTAAGCGTGTTGACTGGACT  18918

AY323977.2     18849 GTTGAATACCCTATTATAGGAGATGAACTGAGGGTTAATTCTGCTTGCAG  18898
                     .|||||||.|||||.||.||.||||||||||.|.|||||.|.|||||.||
MT066156.1     18919 ATTGAATATCCTATAATTGGTGATGAACTGAAGATTAATGCGGCTTGTAG  18968

AY323977.2     18899 AAAAGTACAACACATGGTTGTGAAGTCTGCATTGCTTGCTGATAAGTTTC  18948
                     |||.||.||||||||||||||.||..|||||||..|.||.||.||.||.|
MT066156.1     18969 AAAGGTTCAACACATGGTTGTTAAAGCTGCATTATTAGCAGACAAATTCC  19018

AY323977.2     18949 CAGTTCTTCATGACATTGGAAATCCAAAGGCTATCAAGTGTGTGCCTCAG  18998
                     ||||||||||.||||||||.||.||.||.|||||.||||||||.|||||.
MT066156.1     19019 CAGTTCTTCACGACATTGGTAACCCTAAAGCTATTAAGTGTGTACCTCAA  19068

AY323977.2     18999 GCTGAAGTAGAATGGAAGTTCTACGATGCTCAGCCATGTAGTGACAAAGC  19048
                     |||||.|||||||||||||||||.|||||.|||||.||||||||||||||
MT066156.1     19069 GCTGATGTAGAATGGAAGTTCTATGATGCACAGCCTTGTAGTGACAAAGC  19118

AY323977.2     19049 TTACAAAATAGAGGAACTCTTCTATTCTTATGCTACACATCACGATAAAT  19098
                     |||.||||||||.|||.|.||||||||||||||.||||||...||.||||
MT066156.1     19119 TTATAAAATAGAAGAATTATTCTATTCTTATGCCACACATTCTGACAAAT  19168

AY323977.2     19099 TCACTGATGGTGTTTGTTTGTTTTGGAATTGTAACGTTGATCGTTACCCA  19148
                     ||||.||||||||.||..|.|||||||||||.||.||.|||.|.||.||.
MT066156.1     19169 TCACAGATGGTGTATGCCTATTTTGGAATTGCAATGTCGATAGATATCCT  19218

AY323977.2     19149 GCCAATGCAATTGTGTGTAGGTTTGACACAAGAGTCTTGTCAAACTTGAA  19198
                     ||.|||.|.|||||.|||||.||||||||.|||||..|.||.|||.|.||
MT066156.1     19219 GCTAATTCCATTGTTTGTAGATTTGACACTAGAGTGCTATCTAACCTTAA  19268

AY323977.2     19199 CTTACCAGGCTGTGATGGTGGTAGTTTGTATGTGAATAAGCATGCATTCC  19248
                     |||.||.||.|||||||||||.|||||||||||.|||||.||||||||||
MT066156.1     19269 CTTGCCTGGTTGTGATGGTGGCAGTTTGTATGTAAATAAACATGCATTCC  19318

AY323977.2     19249 ACACTCCAGCTTTCGATAAAAGTGCATTTACTAATTTAAAGCAATTGCCT  19298
                     ||||.||||||||.|||||||||||.|||..|||||||||.|||||.||.
MT066156.1     19319 ACACACCAGCTTTTGATAAAAGTGCTTTTGTTAATTTAAAACAATTACCA  19368

AY323977.2     19299 TTCTTTTACTATTCTGATAGTCCTTGTGAGTCTCATGGCAAACAAGTAGT  19348
                     ||.||.||.||.|||||.|||||.||||||||||||||.|||||||||||
MT066156.1     19369 TTTTTCTATTACTCTGACAGTCCATGTGAGTCTCATGGAAAACAAGTAGT  19418

AY323977.2     19349 GTCGGATATTGATTATGTTCCACTCAAATCTGCTACGTGTATTACACGAT  19398
                     |||.|||||.||||||||.|||||.||.||||||||||||||.|||||.|
MT066156.1     19419 GTCAGATATAGATTATGTACCACTAAAGTCTGCTACGTGTATAACACGTT  19468

AY323977.2     19399 GCAATTTAGGTGGTGCTGTTTGCAGACACCATGCAAATGAGTACCGACAG  19448
                     |||||||||||||||||||.||.|||||.|||||.|||||||||.||..|
MT066156.1     19469 GCAATTTAGGTGGTGCTGTCTGTAGACATCATGCTAATGAGTACAGATTG  19518

AY323977.2     19449 TACTTGGATGCATATAATATGATGATTTCTGCTGGATTTAGCCTATGGAT  19498
                     ||..|.|||||.|||||.||||||||.||.|||||.||||||.|.|||.|
MT066156.1     19519 TATCTCGATGCTTATAACATGATGATCTCAGCTGGCTTTAGCTTGTGGGT  19568

AY323977.2     19499 TTACAAACAATTTGATACTTATAACCTGTGGAATACATTTACCAGGTTAC  19548
                     |||||||||||||||||||||||||||.|||||.||.|||||.||..|.|
MT066156.1     19569 TTACAAACAATTTGATACTTATAACCTCTGGAACACTTTTACAAGACTTC  19618

AY323977.2     19549 AGAGTTTAGAAAATGTGGCTTATAATGTTGTTAATAAAGGACACTTTGAT  19598
                     |||||||||||||||||||||.|||||||||.|||||.||||||||||||
MT066156.1     19619 AGAGTTTAGAAAATGTGGCTTTTAATGTTGTAAATAAGGGACACTTTGAT  19668

AY323977.2     19599 GGACACGCCGGCGAAGCACCTGTTTCCATCATTAATAATGCTGTTTACAC  19648
                     |||||....||.||||.|||.|||||.|||||||||||..||||||||||
MT066156.1     19669 GGACAACAGGGTGAAGTACCAGTTTCTATCATTAATAACACTGTTTACAC  19718

AY323977.2     19649 AAAGGTAGATGGTATTGATGTGGAGATCTTTGAAAATAAGACAACACTTC  19698
                     |||.||.||||||.|||||||.||..|.|||||||||||.||||||.|.|
MT066156.1     19719 AAAAGTTGATGGTGTTGATGTAGAATTGTTTGAAAATAAAACAACATTAC  19768

AY323977.2     19699 CTGTTAATGTTGCATTTGAGCTTTGGGCTAAGCGTAACATTAAACCAGTG  19748
                     ||||||||||.|||||||||||||||||||||||.||||||||||||||.
MT066156.1     19769 CTGTTAATGTAGCATTTGAGCTTTGGGCTAAGCGCAACATTAAACCAGTA  19818

AY323977.2     19749 CCAGAGATTAAGATACTCAATAATTTGGGTGTTGATATCGCTGCTAATAC  19798
                     ||||||.|.||.||||||||||||||||||||.||.||.|||||||||||
MT066156.1     19819 CCAGAGGTGAAAATACTCAATAATTTGGGTGTGGACATTGCTGCTAATAC  19868

AY323977.2     19799 TGTAATCTGGGACTACAAAAGAGAAGCCCCAGCACATGTATCTACAATAG  19848
                     |||.||||||||||||||||||||.||.|||||||||.|||||||.||.|
MT066156.1     19869 TGTGATCTGGGACTACAAAAGAGATGCTCCAGCACATATATCTACTATTG  19918

AY323977.2     19849 GTGTCTGCACAATGACTGACATTGCCAAGAAACCTACTGAGAGTGCTTGT  19898
                     ||||.||..|.|||||||||||.|||||||||||.|||||.|....||||
MT066156.1     19919 GTGTTTGTTCTATGACTGACATAGCCAAGAAACCAACTGAAACGATTTGT  19968

AY323977.2     19899 TCTTCACTTACTGTCTTGTTTGATGGTAGAGTGGAAGGACAGGTAGACCT  19948
                     .|..||||.||||||||.||||||||||||||.||.||.||.||||||.|
MT066156.1     19969 GCACCACTCACTGTCTTTTTTGATGGTAGAGTTGATGGTCAAGTAGACTT  20018

AY323977.2     19949 TTTTAGAAACGCCCGTAATGGTGTTTTAATAACAGAAGGTTCAGTCAAAG  19998
                     .||||||||.|||||||||||||||.|.||.|||||||||...||.||||
MT066156.1     20019 ATTTAGAAATGCCCGTAATGGTGTTCTTATTACAGAAGGTAGTGTTAAAG  20068

AY323977.2     19999 GTCTAACACCTTCAAAGGGACCAGCACAAGCTAGCGTCAATGGAGTCACA  20048
                     ||.||..|||.||....||.||...|||||||||..|.||||||||||||
MT066156.1     20069 GTTTACAACCATCTGTAGGTCCCAAACAAGCTAGTCTTAATGGAGTCACA  20118

AY323977.2     20049 TTAATTGGAGAATCAGTAAAAACACAGTTTAACTACTTTAAGAAAGTAGA  20098
                     ||||||||||||.|.||||||||||||||.||.||.|.|||||||||.||
MT066156.1     20119 TTAATTGGAGAAGCCGTAAAAACACAGTTCAATTATTATAAGAAAGTTGA  20168

AY323977.2     20099 CGGCATTATTCAACAGTTGCCTGAAACCTACTTTACTCAGAGCAGAGACT  20148
                     .||..||.|.|||||.||.||||||||.||||||||||||||.|||.|.|
MT066156.1     20169 TGGTGTTGTCCAACAATTACCTGAAACTTACTTTACTCAGAGTAGAAATT  20218

AY323977.2     20149 TAGAGGATTTTAAGCCCAGATCACAAATGGAAACTGACTTTCTCGAGCTC  20198
                     ||.|.||.|||||.|||||....||||||||||.|||.||..|.||..|.
MT066156.1     20219 TACAAGAATTTAAACCCAGGAGTCAAATGGAAATTGATTTCTTAGAATTA  20268

AY323977.2     20199 GCTATGGATGAATTCATACAGCGATATAAGCTCGAGGGCTATGCCTTCGA  20248
                     |||||||||||||||||..|.||.|||||..|.||.||||||||||||||
MT066156.1     20269 GCTATGGATGAATTCATTGAACGGTATAAATTAGAAGGCTATGCCTTCGA  20318

AY323977.2     20249 ACACATCGTTTATGGAGATTTCAGTCATGGACAACTTGGCGGTCTTCATT  20298
                     |||.|||||||||||||||||.||||||.|.||..|.||.|||.|.|||.
MT066156.1     20319 ACATATCGTTTATGGAGATTTTAGTCATAGTCAGTTAGGTGGTTTACATC  20368

AY323977.2     20299 TAATGATAGGCTTAGCCAAGCGCTCACAAGATTCACCACTTAAATTAGAG  20348
                     ||.||||.||..||||.||.||.|...|.||.|||||..||.|||||||.
MT066156.1     20369 TACTGATTGGACTAGCTAAACGTTTTAAGGAATCACCTTTTGAATTAGAA  20418

AY323977.2     20349 GATTTTATCCCTATGGACAGCACAGTGAAAAATTACTTCATAACAGATGC  20398
                     ||||||||.|||||||||||.|||||.|||||.||.||||||||||||||
MT066156.1     20419 GATTTTATTCCTATGGACAGTACAGTTAAAAACTATTTCATAACAGATGC  20468

AY323977.2     20399 GCAAACAGGTTCATCAAAATGTGTGTGTTCTGTGATTGATCTTTTACTTG  20448
                     |||||||||||||||.||.||||||||||||||.||||||.|.|||||||
MT066156.1     20469 GCAAACAGGTTCATCTAAGTGTGTGTGTTCTGTTATTGATTTATTACTTG  20518

AY323977.2     20449 ATGACTTTGTCGAGATAATAAAGTCACAAGATTTGTCAGTGATTTCAAAA  20498
                     ||||.|||||.||.||||||||.||.||||||||.||.||..||||.||.
MT066156.1     20519 ATGATTTTGTTGAAATAATAAAATCCCAAGATTTATCTGTAGTTTCTAAG  20568

AY323977.2     20499 GTGGTCAAGGTTACAATTGACTATGCTGAAATTTCATTCATGCTTTGGTG  20548
                     ||.|||||.||.||.|||||||||.|.|||||||||||.|||||||||||
MT066156.1     20569 GTTGTCAAAGTGACTATTGACTATACAGAAATTTCATTTATGCTTTGGTG  20618

AY323977.2     20549 TAAGGATGGACATGTTGAAACCTTCTACCCAAAACTACAAGCAAGTCAAG  20598
                     |||.|||||.|||||.|||||.||.|||||||||.|||||.|.|||||||
MT066156.1     20619 TAAAGATGGCCATGTAGAAACATTTTACCCAAAATTACAATCTAGTCAAG  20668

AY323977.2     20599 CGTGGCAACCAGGTGTTGCGATGCCTAACTTGTACAAGATGCAAAGAATG  20648
                     ||||||||||.||||||||.||||||||..|.|||||.||||||||||||
MT066156.1     20669 CGTGGCAACCGGGTGTTGCTATGCCTAATCTTTACAAAATGCAAAGAATG  20718

AY323977.2     20649 CTTCTTGAAAAGTGTGACCTTCAGAATTATGGTGAAAATGCTGTTATACC  20698
                     ||..|.|||||||||||||||||.|||||||||||.|.|||.....||||
MT066156.1     20719 CTATTAGAAAAGTGTGACCTTCAAAATTATGGTGATAGTGCAACATTACC  20768

AY323977.2     20699 AAAAGGAATAATGATGAATGTCGCAAAGTATACTCAACTGTGTCAATACT  20748
                     .|||||.||||||||||||||||||||.||||||||||||||||||||.|
MT066156.1     20769 TAAAGGCATAATGATGAATGTCGCAAAATATACTCAACTGTGTCAATATT  20818

AY323977.2     20749 TAAATACACTTACTTTAGCTGTACCCTACAACATGAGAGTTATTCACTTT  20798
                     ||||.|||.|.||.||||||||||||||.||.|||||||||||.||.|||
MT066156.1     20819 TAAACACATTAACATTAGCTGTACCCTATAATATGAGAGTTATACATTTT  20868

AY323977.2     20799 GGTGCTGGCTCTGATAAAGGAGTTGCACCAGGTACAGCTGTGCTCAGACA  20848
                     ||||||||.||||||||||||||||||||||||||||||||..|.|||||
MT066156.1     20869 GGTGCTGGTTCTGATAAAGGAGTTGCACCAGGTACAGCTGTTTTAAGACA  20918

AY323977.2     20849 ATGGTTGCCAACTGGCACACTACTTGTCGATTCAGATCTTAATGACTTCG  20898
                     .||||||||.||.||.||.||.||||||||||||||||||||||||||.|
MT066156.1     20919 GTGGTTGCCTACGGGTACGCTGCTTGTCGATTCAGATCTTAATGACTTTG  20968

AY323977.2     20899 TCTCCGACGCAGATTCTACTTTAATTGGAGACTGTGCAACAGTACATACG  20948
                     ||||.||.||||||||.|||||.|||||.||.||||||||.||||||||.
MT066156.1     20969 TCTCTGATGCAGATTCAACTTTGATTGGTGATTGTGCAACTGTACATACA  21018

AY323977.2     20949 GCTAATAAATGGGACCTTATTATTAGCGATATGTATGACCCTAGGACCAA  20998
                     ||||||||||||||.||.||||||||.||||||||.|||||||.|||.||
MT066156.1     21019 GCTAATAAATGGGATCTCATTATTAGTGATATGTACGACCCTAAGACTAA  21068

AY323977.2     20999 ACATGTGACAAAAGAGAATGACTCTAAAGAAGGGTTTTTCACTTATCTGT  21048
                     |.||||.||||||||.||||||||||||||.||.|||||||||||..|.|
MT066156.1     21069 AAATGTTACAAAAGAAAATGACTCTAAAGAGGGTTTTTTCACTTACATTT  21118

AY323977.2     21049 GTGGATTTATAAAGCAAAAACTAGCCCTGGGTGGTTCTATAGCTGTAAAG  21098
                     ||||.||||||.|.|||||.|||||.||.||.|||||..|.|||.|||||
MT066156.1     21119 GTGGGTTTATACAACAAAAGCTAGCTCTTGGAGGTTCCGTGGCTATAAAG  21168

AY323977.2     21099 ATAACAGAGCATTCTTGGAATGCTGACCTTTACAAGCTTATGGGCCATTT  21148
                     ||||||||.|||||||||||||||||.|||||.|||||.|||||.||.||
MT066156.1     21169 ATAACAGAACATTCTTGGAATGCTGATCTTTATAAGCTCATGGGACACTT  21218

AY323977.2     21149 CTCATGGTGGACAGCTTTTGTTACAAATGTAAATGCATCATCATCGGAAG  21198
                     |.|||||||||||||.||||||||.|||||.|||||.||||||||.||||
MT066156.1     21219 CGCATGGTGGACAGCCTTTGTTACTAATGTGAATGCGTCATCATCTGAAG  21268

AY323977.2     21199 CATTTTTAATTGGGGCTAACTATCTTGGCAAGCCGAAGGAACAAATTGAT  21248
                     |||||||||||||...|||.|||||||||||.||....||||||||.|||
MT066156.1     21269 CATTTTTAATTGGATGTAATTATCTTGGCAAACCACGCGAACAAATAGAT  21318

AY323977.2     21249 GGCTATACCATGCATGCTAACTACATTTTCTGGAGGAACACAAATCCTAT  21298
                     ||.|||..|||||||||.||.|||||.||.||||||||.||||||||.||
MT066156.1     21319 GGTTATGTCATGCATGCAAATTACATATTTTGGAGGAATACAAATCCAAT  21368

AY323977.2     21299 CCAGTTGTCTTCCTATTCACTCTTTGACATGAGCAAATTTCCTCTTAAAT  21348
                     .|||||||||||||||||..|.|||||||||||.||||||||.|||||||
MT066156.1     21369 TCAGTTGTCTTCCTATTCTTTATTTGACATGAGTAAATTTCCCCTTAAAT  21418

AY323977.2     21349 TAAGAGGAACTGCTGTAATGTCTCTTAAGGAGAATCAAATCAATGATATG  21398
                     ||||.||.||||||||.||||||.|.||.||...||||||||||||||||
MT066156.1     21419 TAAGGGGTACTGCTGTTATGTCTTTAAAAGAAGGTCAAATCAATGATATG  21468

AY323977.2     21399 ATTTATTCTCTTCTGGAAAAAGGTAGGCTTATCATTAGAGAAAACAACAG  21448
                     ||||..||||||||....||||||||.|||||.|||||||||||||||||
MT066156.1     21469 ATTTTATCTCTTCTTAGTAAAGGTAGACTTATAATTAGAGAAAACAACAG  21518

AY323977.2     21449 AGTTGTGGTTTCAAGTGATATTCTTGTTAACAACTAAACGAACA-TGTTT  21497
                     ||||||..||||.||||||.|||||||||||||||||||||||| |||||
MT066156.1     21519 AGTTGTTATTTCTAGTGATGTTCTTGTTAACAACTAAACGAACAATGTTT  21568

AY323977.2     21498 ATTTTCTTATTATTTCTTACTCTCACTAGTGG-TAGTGACCTTGACCGGT  21546
                     .|||  ||.||.||  |||.|..|||||||.. ||||.|         ||
MT066156.1     21569 GTTT--TTCTTGTT--TTATTGCCACTAGTCTCTAGTCA---------GT  21605

AY323977.2     21547 GCACCACTTTTGATGATGTTCAAGCTCCTAATTACACTCAACATAC----  21592
                     |....|.|.|| |..|    |.||..|..||||||.|.|..|||||
MT066156.1     21606 GTGTTAATCTT-ACAA----CCAGAACTCAATTACCCCCTGCATACACTA  21650

AY323977.2     21593 -TTCATCTATGAGGGGGGTTTACTATCCTGATGAAATTTTTAGATCAGAC  21641
                      |||.|..|...|.||.|||||.||.|||||..||.||||.|||||....
MT066156.1     21651 ATTCTTTCACACGTGGTGTTTATTACCCTGACAAAGTTTTCAGATCCTCA  21700

AY323977.2     21642 ACTCTTTATTTAACTCAGGATTTATTTCTTCCATTTTATTCTAATGTTAC  21691
                     ..|.|..|||.|||||||||.||.||..|.||.||.|.|||.||||||||
MT066156.1     21701 GTTTTACATTCAACTCAGGACTTGTTCTTACCTTTCTTTTCCAATGTTAC  21750

AY323977.2     21692 AGGGTTTCATACTAT---------------------TAATCATACGTTTG  21720
                     ..||||.|||.||||                     ||.|.|.|.|||||
MT066156.1     21751 TTGGTTCCATGCTATACATGTCTCTGGGACCAATGGTACTAAGAGGTTTG  21800

AY323977.2     21721 GCAACCCTGTCATACCTTTTAAGGATGGTATTTATTTTGCTGCCACAGAG  21770
                     ..|||||||||.||||.|||||.||||||.|||||||||||.||||.|||
MT066156.1     21801 ATAACCCTGTCCTACCATTTAATGATGGTGTTTATTTTGCTTCCACTGAG  21850

AY323977.2     21771 AAATCAAATGTTGTCCGTGGTTGGGTTTTTGGTTCTACCATGAACAACAA  21820
                     ||.||.||..|..|..|.||.|||.||||||||.||||..|..|....||
MT066156.1     21851 AAGTCTAACATAATAAGAGGCTGGATTTTTGGTACTACTTTAGATTCGAA  21900

AY323977.2     21821 GTCACAGTCGGTGATTATTATTAACAATTCTACTAATGTTGTTATACGAG  21870
                     |.|.|||||..|..|||||.||||.||..||||||||||||||||...||
MT066156.1     21901 GACCCAGTCCCTACTTATTGTTAATAACGCTACTAATGTTGTTATTAAAG  21950

AY323977.2     21871 CATGTAACTTTGAATTGTGTGACAACCCTTTCTTTGCTGTTTCTAAACCC  21920
                     ..|||.|.|||.||||.|||.|..|.||.||.||.|.|||||.|.|.|.|
MT066156.1     21951 TCTGTGAATTTCAATTTTGTAATGATCCATTTTTGGGTGTTTATTACCAC  22000

AY323977.2     21921 A--------------TGGGTACACAGACACATACTATGATATTCGATAAT  21956
                     |              |||.|..|.||..|..|...|...|||||  ||.|
MT066156.1     22001 AAAAACAACAAAAGTTGGATGGAAAGTGAGTTCAGAGTTTATTC--TAGT  22048

AY323977.2     21957 GCATTTAATTGCACTTTCGAGTACATATCTGATGCCTTTTCGCTTG-ATG  22005
                     ||...||||||||||||.||.||..|.|||.| ||||||||...|| |..
MT066156.1     22049 GCGAATAATTGCACTTTTGAATATGTCTCTCA-GCCTTTTCTTATGGACC  22097

AY323977.2     22006 TTTCAGAAAAGTCAGGTAATTTTAAACACTTACGAGAGTTTGTGTTTAAA  22055
                     ||..||.|||....||||||||.|||.|..|..|.||.|||||||||||.
MT066156.1     22098 TTGAAGGAAAACAGGGTAATTTCAAAAATCTTAGGGAATTTGTGTTTAAG  22147

AY323977.2     22056 AATAAAGATGGGTTTCTCTATGTTTATAAGGGCTATCAACCTATAGATGT  22105
                     ||||..|||||.|.|.|..|..|.|||.......|....|||||..||.|
MT066156.1     22148 AATATTGATGGTTATTTTAAAATATATTCTAAGCACACGCCTATTAATTT  22197

AY323977.2     22106 AGTTCGTGATCTACCTTCTGGTTTTAACACTTTGAAACCTATTTTTAAGT  22155
                     |||.||||||||.|||...||||||....||||..||||..|..|..|.|
MT066156.1     22198 AGTGCGTGATCTCCCTCAGGGTTTTTCGGCTTTAGAACCATTGGTAGATT  22247

AY323977.2     22156 TGCCTCTTGGTATTAACATTACAAATTTTAGAGCCATTCTTAC-------  22198
                     ||||..|.|||||||||||.||.|..|||..|.|..|.|||.|
MT066156.1     22248 TGCCAATAGGTATTAACATCACTAGGTTTCAAACTTTACTTGCTTTACAT  22297

AY323977.2     22199 ---AGCCTTTTCAC--CTGCTCAAGACATTTGGGGCACGTCAGCTG----  22239
                        ||...|||.||  |||.|.|......||..||...|.||||||
MT066156.1     22298 AGAAGTTATTTGACTCCTGGTGATTCTTCTTCAGGTTGGACAGCTGGTGC  22347

AY323977.2     22240 --CAGCCTATTTTGTTGGCTATTTAAAGCCAACTACATTTATGCTCAAGT  22287
                       ||||.||||.|||.||.|||.|..|.||.|..||.|||.|..|.||.|
MT066156.1     22348 TGCAGCTTATTATGTGGGTTATCTTCAACCTAGGACTTTTCTATTAAAAT  22397

AY323977.2     22288 ATGATGAAAATGGTACAATCACAGATGCTGTTGATTGTTCTCAAAATCCA  22337
                     ||.||||||||||.||.||.|||||||||||.||.|||.|.|...|.||.
MT066156.1     22398 ATAATGAAAATGGAACCATTACAGATGCTGTAGACTGTGCACTTGACCCT  22447

AY323977.2     22338 CTTGCTGAACTCAAATGCTCTGTTAAGAGCTTTGAGATTGACAAAGGAAT  22387
                     ||..|.|||...||.||..|..|.||...|||.....|.||.||||||||
MT066156.1     22448 CTCTCAGAAACAAAGTGTACGTTGAAATCCTTCACTGTAGAAAAAGGAAT  22497

AY323977.2     22388 TTACCAGACCTCTAATTTCAGGGTTGTTCCCTCAGGAGATGTTGTGAGAT  22437
                     .||.||.||.|||||.||.||.||....||..|||.|..|.||||.||||
MT066156.1     22498 CTATCAAACTTCTAACTTTAGAGTCCAACCAACAGAATCTATTGTTAGAT  22547

AY323977.2     22438 TCCCTAATATTACAAACTTGTGTCCTTTTGGAGAGGTTTTTAATGCTACT  22487
                     |.||||||||||||||||||||.||||||||.||.||||||||.||.||.
MT066156.1     22548 TTCCTAATATTACAAACTTGTGCCCTTTTGGTGAAGTTTTTAACGCCACC  22597

AY323977.2     22488 AAATTCCCTTCTGTCTATGCATGGGAGAGAAAAAAAATTTCTAATTGTGT  22537
                     |.|||..|.|||||.|||||.|||.|.||.||.|.|||....||.|||||
MT066156.1     22598 AGATTTGCATCTGTTTATGCTTGGAACAGGAAGAGAATCAGCAACTGTGT  22647

AY323977.2     22538 TGCTGATTACTCTGTGCTCTACAACTCAACATTTTTTTCAACCTTTAAGT  22587
                     |||||||||.|||||.||.||.||.||..|||..|||||.||.|||||||
MT066156.1     22648 TGCTGATTATTCTGTCCTATATAATTCCGCATCATTTTCCACTTTTAAGT  22697

AY323977.2     22588 GCTATGGCGTTTCTGCCACTAAGTTGAATGATCTTTGCTTCTCCAATGTC  22637
                     |.|||||.||.|||.|.|||||.||.||||||||.|||||..|.||||||
MT066156.1     22698 GTTATGGAGTGTCTCCTACTAAATTAAATGATCTCTGCTTTACTAATGTC  22747

AY323977.2     22638 TATGCAGATTCTTTTGTAGTCAAGGGAGATGATGTAAGACAAATAGCGCC  22687
                     |||||||||||.||||||.|.|..||.|||||.||.||||||||.||.||
MT066156.1     22748 TATGCAGATTCATTTGTAATTAGAGGTGATGAAGTCAGACAAATCGCTCC  22797

AY323977.2     22688 AGGACAAACTGGTGTTATTGCTGATTATAATTATAAATTGCCAGATGATT  22737
                     |||.||||||||....|||||||||||||||||||||||.||||||||||
MT066156.1     22798 AGGGCAAACTGGAAAGATTGCTGATTATAATTATAAATTACCAGATGATT  22847

AY323977.2     22738 TCATGGGTTGTGTCCTTGCTTGGAATACTAGGAACATTGATGCTACTTCA  22787
                     |.|..||.||.||..|.|||||||||.|||..||..|||||.|||.....
MT066156.1     22848 TTACAGGCTGCGTTATAGCTTGGAATTCTAACAATCTTGATTCTAAGGTT  22897

AY323977.2     22788 ACTGGTAATTATAATTATAAATATAGGTATCTTAGACATGGCAAGCTTAG  22837
                     ..|||||||||||||||....|||||.|...||||..|....||.||.|.
MT066156.1     22898 GGTGGTAATTATAATTACCTGTATAGATTGTTTAGGAAGTCTAATCTCAA  22947

AY323977.2     22838 GCCCTTTGAGAGAGACATATCTAATGTGC-CTTTCTCCCCTGATGGCAAA  22886
                     .||.|||||||||||.||.||.|.||... ||.||...||.| |.|||.|
MT066156.1     22948 ACCTTTTGAGAGAGATATTTCAACTGAAATCTATCAGGCCGG-TAGCACA  22996

AY323977.2     22887 CCTTGCACCCCACCTGC---TCTTAATTGTTATTGGCCATTAAATGATTA  22933
                     |||||.|.......||.   |.||||||||||.|..||.|||.|....||
MT066156.1     22997 CCTTGTAATGGTGTTGAAGGTTTTAATTGTTACTTTCCTTTACAATCATA  23046

AY323977.2     22934 TGGTTTTTACACCACTACTGGCATTGGCTACCAACCTTACAGAGTTGTAG  22983
                     ||||||..|..||||||.|||..||||.||||||||.||||||||.||||
MT066156.1     23047 TGGTTTCCAACCCACTAATGGTGTTGGTTACCAACCATACAGAGTAGTAG  23096

AY323977.2     22984 TACTTTCTTTTGAACTTTTAAATGCACCGGCCACGGTTTGTGGACCAAAA  23033
                     |||||||||||||||||.||.|||||||.||.||.|||||||||||.|||
MT066156.1     23097 TACTTTCTTTTGAACTTCTACATGCACCAGCAACTGTTTGTGGACCTAAA  23146

AY323977.2     23034 TTATCCACTGACCTTATTAAGAACCAGTGTGTCAATTTTAATTTTAATGG  23083
                     ...||.|||.|..|..||||.|||.|.|||||||||||.||.||.|||||
MT066156.1     23147 AAGTCTACTAATTTGGTTAAAAACAAATGTGTCAATTTCAACTTCAATGG  23196

AY323977.2     23084 ACTCACTGGTACTGGTGTGTTAACTCCTTCTTCAAAGAGATTTCAACCAT  23133
                     ..|.||.||.||.|||||..|.|||...|||...||.|..||||..||.|
MT066156.1     23197 TTTAACAGGCACAGGTGTTCTTACTGAGTCTAACAAAAAGTTTCTGCCTT  23246

AY323977.2     23134 TTCAACAATTTGGCCGTGATGTTTCTGATTTCACTGATTCCGTTCGAGAT  23183
                     |.||||||||||||.|.||..||.||||....||||||.|.||.||.|||
MT066156.1     23247 TCCAACAATTTGGCAGAGACATTGCTGACACTACTGATGCTGTCCGTGAT  23296

AY323977.2     23184 CCTAAAACATCTGAAATATTAGACATTTCACCTTGCTCTTTTGGGGGTGT  23233
                     ||..|.|||..|||.||..|.||||||.||||.||.||||||||.|||||
MT066156.1     23297 CCACAGACACTTGAGATTCTTGACATTACACCATGTTCTTTTGGTGGTGT  23346

AY323977.2     23234 AAGTGTAATTACACCTGGAACAAATGCTTCATCTGAAGTTGCTGTTCTAT  23283
                     .|||||.||.|||||.|||||||||.||||.....|.|||||||||||.|
MT066156.1     23347 CAGTGTTATAACACCAGGAACAAATACTTCTAACCAGGTTGCTGTTCTTT  23396

AY323977.2     23284 ATCAAGATGTTAACTGCACTGATGTTTCTACAGCAATTCATGCAGATCAA  23333
                     ||||.||||||||||||||.||.||..||...||.|||||||||||||||
MT066156.1     23397 ATCAGGATGTTAACTGCACAGAAGTCCCTGTTGCTATTCATGCAGATCAA  23446

AY323977.2     23334 CTCACACCAGCTTGGCGCATATATTCTACTGGAAACAATGTATTCCAGAC  23383
                     ||.||.||..|||||||..|.||||||||.||....|||||.||.||.||
MT066156.1     23447 CTTACTCCTACTTGGCGTGTTTATTCTACAGGTTCTAATGTTTTTCAAAC  23496

AY323977.2     23384 TCAAGCAGGCTGTCTTATAGGAGCTGAGCATGTCGACACTTCTTATGAGT  23433
                     .|..|||||||||.|.|||||.|||||.||||||.|||..||.|||||||
MT066156.1     23497 ACGTGCAGGCTGTTTAATAGGGGCTGAACATGTCAACAACTCATATGAGT  23546

AY323977.2     23434 GCGACATTCCTATTGGAGCTGGCATTTGTGCTAGTTACCATAC--AGTTT  23481
                     |.|||||.||.|||||.||.||.||.||.||||||||.||.||  ||..|
MT066156.1     23547 GTGACATACCCATTGGTGCAGGTATATGCGCTAGTTATCAGACTCAGACT  23596

AY323977.2     23482 CTTTATT----------ACGTAGTACTAGCCAAA-AATCTATTGTGGCTT  23520
                     ..||.|.          |||||||. ||||.|.. ||||.||..|.||.|
MT066156.1     23597 AATTCTCCTCGGCGGGCACGTAGTG-TAGCTAGTCAATCCATCATTGCCT  23645

AY323977.2     23521 ATACTATGTCTTTAGGTGCTGATAGTTCAATTGCTTACTCTAATAACACC  23570
                     |.||||||||..|.|||||.||.|.||||.|||||||||||||||||.|.
MT066156.1     23646 ACACTATGTCACTTGGTGCAGAAAATTCAGTTGCTTACTCTAATAACTCT  23695

AY323977.2     23571 ATTGCTATACCTACTAACTTTTCAATTAGCATTACTACAGAAGTAATGCC  23620
                     |||||.|||||.||.||.|||.|.|||||..||||.||||||.|..|.||
MT066156.1     23696 ATTGCCATACCCACAAATTTTACTATTAGTGTTACCACAGAAATTCTACC  23745

AY323977.2     23621 TGTTTCTATGGCTAAAACCTCCGTAGATTGTAATATGTACATCTGCGGAG  23670
                     .||.||||||.|.||.||.||.||||||||||..||||||||.||.||.|
MT066156.1     23746 AGTGTCTATGACCAAGACATCAGTAGATTGTACAATGTACATTTGTGGTG  23795

AY323977.2     23671 ATTCTACTGAATGTGCTAATTTGCTTCTCCAATATGGTAGCTTTTGCACA  23720
                     ||||.||||||||....|||.|..|..|.||||||||.||.|||||.|||
MT066156.1     23796 ATTCAACTGAATGCAGCAATCTTTTGTTGCAATATGGCAGTTTTTGTACA  23845

AY323977.2     23721 CAACTAAATCGTGCACTCTCAGGTATTGCTGCTGAACAGGATCGCAACAC  23770
                     |||.||||.|||||..|..|.||.||.||||.||||||.||....|||||
MT066156.1     23846 CAATTAAACCGTGCTTTAACTGGAATAGCTGTTGAACAAGACAAAAACAC  23895

AY323977.2     23771 ACGTGAAGTGTTCGCTCAAGTCAAACAAATGTACAAAACCCCAACTTTGA  23820
                     .|..|||||.||.||.||||||||||||||.||||||||.|||.|..|.|
MT066156.1     23896 CCAAGAAGTTTTTGCACAAGTCAAACAAATTTACAAAACACCACCAATTA  23945

AY323977.2     23821 AATATTTTGGTGGTTTTAATTTTTCACAAATATTACCTGACCCTCTAAAG  23870
                     ||.||||||||||||||||||||||||||||||||||.||.||...|||.
MT066156.1     23946 AAGATTTTGGTGGTTTTAATTTTTCACAAATATTACCAGATCCATCAAAA  23995

AY323977.2     23871 CCAACTAAGAGGTCTTTTATTGAGGACTTGCTCTTTAATAAGGTGACACT  23920
                     ||||..||||||||.||||||||.||..|.||.||.||.||.||||||||
MT066156.1     23996 CCAAGCAAGAGGTCATTTATTGAAGATCTACTTTTCAACAAAGTGACACT  24045

AY323977.2     23921 CGCTGATGCTGGCTTCATGAAGCAATATGGCGAATGCCTAGGTGATATTA  23970
                     .||.||||||||||||||.||.||||||||.||.|||||.|||||||||.
MT066156.1     24046 TGCAGATGCTGGCTTCATCAAACAATATGGTGATTGCCTTGGTGATATTG  24095

AY323977.2     23971 ATGCTAGAGATCTCATTTGTGCGCAGAAGTTCAATGGACTTACAGTGTTG  24020
                     .|||||||||.|||||||||||.||.|||||.||.||.|||||.||.|||
MT066156.1     24096 CTGCTAGAGACCTCATTTGTGCACAAAAGTTTAACGGCCTTACTGTTTTG  24145

AY323977.2     24021 CCACCTCTGCTCACTGATGATATGATTGCTGCCTACACTGCTGCTCTAGT  24070
                     ||||||.|||||||.|||||.|||||||||...||||||.||||.|| ||
MT066156.1     24146 CCACCTTTGCTCACAGATGAAATGATTGCTCAATACACTTCTGCACT-GT  24194

AY323977.2     24071 TAGTGG-TACTGCCACTGCTGGATGGACATTTGGTGCTGGCGCTGCTCTT  24119
                     |||.|| |||...||||.||||.|||||.||||||||.||.|||||..|.
MT066156.1     24195 TAGCGGGTACAATCACTTCTGGTTGGACCTTTGGTGCAGGTGCTGCATTA  24244

AY323977.2     24120 CAAATACCTTTTGCTATGCAAATGGCATATAGGTTCAATGGCATTGGAGT  24169
                     ||||||||.|||||||||||||||||.||||||||.|||||.||||||||
MT066156.1     24245 CAAATACCATTTGCTATGCAAATGGCTTATAGGTTTAATGGTATTGGAGT  24294

AY323977.2     24170 TACCCAAAATGTTCTCTATGAGAACCAAAAACAAATCGCCAACCAATTTA  24219
                     |||.||.||||||||||||||||||||||||...||.|||||||||||||
MT066156.1     24295 TACACAGAATGTTCTCTATGAGAACCAAAAATTGATTGCCAACCAATTTA  24344

AY323977.2     24220 ACAAGGCGATTAGTCAAATTCAAGAATCACTTACAACAACATCAACTGCA  24269
                     |.|..||.|||.|..||||||||||.||||||.|..|.|||.|||.||||
MT066156.1     24345 ATAGTGCTATTGGCAAAATTCAAGACTCACTTTCTTCCACAGCAAGTGCA  24394

AY323977.2     24270 TTGGGCAAGCTGCAAGACGTTGTTAACCAGAATGCTCAAGCATTAAACAC  24319
                     .|.||.||.||.|||||.||.||.|||||.|||||.|||||.||||||||
MT066156.1     24395 CTTGGAAAACTTCAAGATGTGGTCAACCAAAATGCACAAGCTTTAAACAC  24444

AY323977.2     24320 ACTTGTTAAACAACTTAGCTCTAATTTTGGTGCAATTTCAAGTGTGCTAA  24369
                     .||||||||||||||||||||.|||||||||||||||||||||||..|||
MT066156.1     24445 GCTTGTTAAACAACTTAGCTCCAATTTTGGTGCAATTTCAAGTGTTTTAA  24494

AY323977.2     24370 ATGATATCCTTTCGCGACTTGATAAAGTCGAGGCGGAGGTACAAATTGAC  24419
                     |||||||||||||.||.|||||.|||||.|||||.||.||.||||||||.
MT066156.1     24495 ATGATATCCTTTCACGTCTTGACAAAGTTGAGGCTGAAGTGCAAATTGAT  24544

AY323977.2     24420 AGGTTAATTACAGGCAGACTTCAAAGCCTTCAAACCTATGTAACACAACA  24469
                     |||||.||.|||||||||||||||||..|.||.||.|||||.||.|||||
MT066156.1     24545 AGGTTGATCACAGGCAGACTTCAAAGTTTGCAGACATATGTGACTCAACA  24594

AY323977.2     24470 ACTAATCAGGGCTGCTGAAATCAGGGCTTCTGCTAATCTTGCTGCTACTA  24519
                     |.||||.||.|||||.||||||||.|||||||||||||||||||||||||
MT066156.1     24595 ATTAATTAGAGCTGCAGAAATCAGAGCTTCTGCTAATCTTGCTGCTACTA  24644

AY323977.2     24520 AAATGTCTGAGTGTGTTCTTGGACAATCAAAAAGAGTTGACTTTTGTGGA  24569
                     |||||||.||||||||.|||||||||||||||||||||||.|||||||||
MT066156.1     24645 AAATGTCAGAGTGTGTACTTGGACAATCAAAAAGAGTTGATTTTTGTGGA  24694

AY323977.2     24570 AAGGGCTACCACCTTATGTCCTTCCCACAAGCAGCCCCGCATGGTGTTGT  24619
                     ||||||||.||.||||||||||||||.||..||||.||.||||||||.||
MT066156.1     24695 AAGGGCTATCATCTTATGTCCTTCCCTCAGTCAGCACCTCATGGTGTAGT  24744

AY323977.2     24620 CTTCCTACATGTCACGTATGTGCCATCCCAGGAGAGGAACTTCACCACAG  24669
                     ||||.|.|||||.||.|||||.||..|.||.||.|.|||||||||.||.|
MT066156.1     24745 CTTCTTGCATGTGACTTATGTCCCTGCACAAGAAAAGAACTTCACAACTG  24794

AY323977.2     24670 CGCCAGCAATTTGTCATGAAGGCAAAGCATACTTCCCTCGTGAAGGTGTT  24719
                     |.||.||.|||||||||||.||.||||||.||||.||||||||||||||.
MT066156.1     24795 CTCCTGCCATTTGTCATGATGGAAAAGCACACTTTCCTCGTGAAGGTGTC  24844

AY323977.2     24720 TTTGTGTTTAATGGCACTTCTTGGTTTATTACACAGAGGAACTTCTTTTC  24769
                     |||||.|..||||||||....||||||.|.|||||.|||||.||.|.|..
MT066156.1     24845 TTTGTTTCAAATGGCACACACTGGTTTGTAACACAAAGGAATTTTTATGA  24894

AY323977.2     24770 TCCACAAATAATTACTACAGACAATACATTTGTCTCAGGAAATTGTGATG  24819
                     .||||||||.||||||||||||||.||||||||.||.||.||.|||||||
MT066156.1     24895 ACCACAAATCATTACTACAGACAACACATTTGTGTCTGGTAACTGTGATG  24944

AY323977.2     24820 TCGTTATTGGCATCATTAACAACACAGTTTATGATCCTCTGCAACCTGAG  24869
                     |.||.||.||.||..|.|||||||||||||||||||||.||||||||||.
MT066156.1     24945 TTGTAATAGGAATTGTCAACAACACAGTTTATGATCCTTTGCAACCTGAA  24994

AY323977.2     24870 CTTGACTCATTCAAAGAAGAGCTGGACAAGTACTTCAAAAATCATACATC  24919
                     .|.|||||||||||.||.|||.|.||.||.||.||.||.|||||||||||
MT066156.1     24995 TTAGACTCATTCAAGGAGGAGTTAGATAAATATTTTAAGAATCATACATC  25044

AY323977.2     24920 ACCAGATGTTGATCTTGGCGACATTTCAGGCATTAACGCTTCTGTCGTCA  24969
                     |||||||||||||.|.||.|||||.||.||||||||.|||||.||.||.|
MT066156.1     25045 ACCAGATGTTGATTTAGGTGACATCTCTGGCATTAATGCTTCAGTTGTAA  25094

AY323977.2     24970 ACATTCAAAAAGAAATTGACCGCCTCAATGAGGTCGCTAAAAATTTAAAT  25019
                     ||||||||||||||||||||||||||||||||||.||.||.|||||||||
MT066156.1     25095 ACATTCAAAAAGAAATTGACCGCCTCAATGAGGTTGCCAAGAATTTAAAT  25144

AY323977.2     25020 GAATCACTCATTGACCTTCAAGAATTGGGAAAATATGAGCAATATATTAA  25069
                     |||||.|||||.||.||.||||||.|.|||||.||||||||.|||||.||
MT066156.1     25145 GAATCTCTCATCGATCTCCAAGAACTTGGAAAGTATGAGCAGTATATAAA  25194

AY323977.2     25070 ATGGCCTTGGTATGTTTGGCTCGGCTTCATTGCTGGACTAATTGCCATCG  25119
                     ||||||.|||||..|||||||.||.||.||.|||||..|.||||||||.|
MT066156.1     25195 ATGGCCATGGTACATTTGGCTAGGTTTTATAGCTGGCTTGATTGCCATAG  25244

AY323977.2     25120 TCATGGTTACAATCTTGCTTTGTTGCATGACTAGTTGTTGCAGTTGCCTC  25169
                     |.|||||.|||||..|||||||.||.|||||.|||||.||.|||||.|||
MT066156.1     25245 TAATGGTGACAATTATGCTTTGCTGTATGACCAGTTGCTGTAGTTGTCTC  25294

AY323977.2     25170 AAGGGTGCATGCTCTTGTGGTTCTTGCTGCAAGTTTGATGAGGATGACTC  25219
                     |||||....||.||||||||.||.||||||||.||||||||.||.|||||
MT066156.1     25295 AAGGGCTGTTGTTCTTGTGGATCCTGCTGCAAATTTGATGAAGACGACTC  25344

AY323977.2     25220 TGAGCCAGTTCTCAAGGGTGTCAAATTACATTACACATAAACGAACTTAT  25269
                     |||||||||.|||||.||.|||||||||||||||||||||||||||||||
MT066156.1     25345 TGAGCCAGTGCTCAAAGGAGTCAAATTACATTACACATAAACGAACTTAT  25394

AY323977.2     25270 GGATTTGTTTATGAGATTTTTTACTCTTGGATCAATTACTGCACAGCCAG  25319
                     ||||||||||||||||.|.||.||..|||||.|..|.|||....|||.||
MT066156.1     25395 GGATTTGTTTATGAGAATCTTCACAATTGGAACTGTAACTTTGAAGCAAG  25444

AY323977.2     25320 TAAAAATTGACAATGCTTCTCCTGCAAGTACTGTTCATGCTACAGCAACG  25369
                     ...||||..|..|||||.|||||.||..|..|||||..|||||.||||||
MT066156.1     25445 GTGAAATCAAGGATGCTACTCCTTCAGATTTTGTTCGCGCTACTGCAACG  25494

AY323977.2     25370 ATACCGCTACAAGCCTCACTCCCTTTCGGATGGCTTGTTATTGGCGTTGC  25419
                     ||||||.|||||||||||||||||||||||||||||.||.||||||||||
MT066156.1     25495 ATACCGATACAAGCCTCACTCCCTTTCGGATGGCTTATTGTTGGCGTTGC  25544

AY323977.2     25420 ATTTCTTGCTGTTTTTCAGAGCGCTACCAAAATAATTGCGCTCAATAAAA  25469
                     |.|||||||||||||||||||||||.|||||||.||..|.|||||.||.|
MT066156.1     25545 ACTTCTTGCTGTTTTTCAGAGCGCTTCCAAAATCATAACCCTCAAAAAGA  25594

AY323977.2     25470 GATGGCAGCTAGCCCTTTATAAGGGCTTCCAGTTCATTTGCAATTTACTG  25519
                     |||||||.|||||.||.|..|||||..|.||.||..|||||||.||.|||
MT066156.1     25595 GATGGCAACTAGCACTCTCCAAGGGTGTTCACTTTGTTTGCAACTTGCTG  25644

AY323977.2     25520 CTGCTATTTGTTACCATCTATTCACATCTTTTGCTTGTCGCTGCAGGTAT  25569
                     .||.|.|||||.||..|.||.|||||.||||||||.||.|||||.||..|
MT066156.1     25645 TTGTTGTTTGTAACAGTTTACTCACACCTTTTGCTCGTTGCTGCTGGCCT  25694

AY323977.2     25570 GGAGGCGCAATTTTTGTACCTCTATGCCTTGATATATTTTCTACAATGCA  25619
                     .||.||.|..|||.|.||.||.|||||.||..|.||.||..|.||..|.|
MT066156.1     25695 TGAAGCCCCTTTTCTCTATCTTTATGCTTTAGTCTACTTCTTGCAGAGTA  25744

AY323977.2     25620 TCAACGCATGTAGAATTATTATGAGATGTTGGCTTTGTTGGAAGTGCAAA  25669
                     |.|||......|||||.||.|||||...|||||||||.|||||.|||...
MT066156.1     25745 TAAACTTTGTAAGAATAATAATGAGGCTTTGGCTTTGCTGGAAATGCCGT  25794

AY323977.2     25670 TCCAAGAACCCATTACTTTATGATGCCAACTACTTTGTTTGCTGGCACAC  25719
                     |||||.||||||||||||||||||||||||||.|||.||||||||||.||
MT066156.1     25795 TCCAAAAACCCATTACTTTATGATGCCAACTATTTTCTTTGCTGGCATAC  25844

AY323977.2     25720 ACATAACTATGACTACTGTATACCATATAACAGTGTCACAGATACAATTG  25769
                     ..||...||.|||||.||||||||.||.||.|||||.||...|.||||||
MT066156.1     25845 TAATTGTTACGACTATTGTATACCTTACAATAGTGTAACTTCTTCAATTG  25894

AY323977.2     25770 TCGTTACTGAAGGTGACGGCATTTCAACACCAAAACTCAAAGAAGACTAC  25819
                     ||.|||||..||||||.||||...|||..||.|......||.|.||||||
MT066156.1     25895 TCATTACTTCAGGTGATGGCACAACAAGTCCTATTTCTGAACATGACTAC  25944

AY323977.2     25820 CAAATTGGTGGTTATTCTGAGGATAGGCACTCAGGTGTTAAAGACTATGT  25869
                     ||.||||||||||||.||||..|..||.|.||.||.||.|||||||.|||
MT066156.1     25945 CAGATTGGTGGTTATACTGAAAAATGGGAATCTGGAGTAAAAGACTGTGT  25994

AY323977.2     25870 CGTTGTACATGGCTATTTCACCGAAGTTTACTACCAGCTTGAGTCTACAC  25919
                     .||..||||..|.||.|||||...||..||.||||||||..|.||.||.|
MT066156.1     25995 TGTATTACACAGTTACTTCACTTCAGACTATTACCAGCTGTACTCAACTC  26044

AY323977.2     25920 AAATTACTACAGACACTGGTATTGAAAATGCTACATTCTTCATCTTTAAC  25969
                     ||.|.|.|||||||||||||.|||||.|||.|||.||||||||||..||.
MT066156.1     26045 AATTGAGTACAGACACTGGTGTTGAACATGTTACCTTCTTCATCTACAAT  26094

AY323977.2     25970 AAGCTTGTTAAAGACCC---ACCGAATGTGCAAATACACACAATCGACGG  26016
                     ||..|||||.|.||.||   |....||||.|||||.|||||||||||||.
MT066156.1     26095 AAAATTGTTGATGAGCCTGAAGAACATGTCCAAATTCACACAATCGACGT  26144

AY323977.2     26017 CTCTTCAGGAGTTGCTAATCCAGCAATGGATCCAATTTATGATGAGCCGA  26066
                     .||.||.|||||||.||||||||.||||||.||||||||||||||.||||
MT066156.1     26145 TTCATCCGGAGTTGTTAATCCAGTAATGGAACCAATTTATGATGAACCGA  26194

AY323977.2     26067 CGACGACTACTAGCGTGCCTTTGTAAGCACAAGAAAGTGAGTACGAACTT  26116
                     |||||||||||||||||||||||||||||||||....|||||||||||||
MT066156.1     26195 CGACGACTACTAGCGTGCCTTTGTAAGCACAAGCTGATGAGTACGAACTT  26244

AY323977.2     26117 ATGTACTCATTCGTTTCGGAAGAAACAGGTACGTTAATAGTTAATAGCGT  26166
                     |||||||||||||||||||||||.||||||||||||||||||||||||||
MT066156.1     26245 ATGTACTCATTCGTTTCGGAAGAGACAGGTACGTTAATAGTTAATAGCGT  26294

AY323977.2     26167 ACTTCTTTTTCTTGCTTTCGTGGTATTCTTGCTAGTCACACTAGCCATCC  26216
                     ||||||||||||||||||||||||||||||||||||.|||||||||||||
MT066156.1     26295 ACTTCTTTTTCTTGCTTTCGTGGTATTCTTGCTAGTTACACTAGCCATCC  26344

AY323977.2     26217 TTACTGCGCTTCGATTGTGTGCGTACTGCTGCAATATTGTTAACGTGAGT  26266
                     ||||||||||||||||||||||||||||||||||||||||||||||||||
MT066156.1     26345 TTACTGCGCTTCGATTGTGTGCGTACTGCTGCAATATTGTTAACGTGAGT  26394

AY323977.2     26267 TTAGTAAAACCAACGGTTTACGTCTACTCGCGTGTTAAAAATCTGAACTC  26316
                     .|.||||||||..|..|||||||.|||||.|||||||||||||||||.||
MT066156.1     26395 CTTGTAAAACCTTCTTTTTACGTTTACTCTCGTGTTAAAAATCTGAATTC  26444

AY323977.2     26317 TTCTGAAGGAGTTCCTGATCTTCTGGTCTAAACGAACTAACTATTAT--T  26364
                     ||||.   ||||||||||||||||||||||||||||||||.||||||  |
MT066156.1     26445 TTCTA---GAGTTCCTGATCTTCTGGTCTAAACGAACTAAATATTATATT  26491

AY323977.2     26365 A-TTATTCTGTTTGGAACTTTAACATT-GCTTATCATGGCAGA---CAAC  26409
                     | ||.||||||||||||||||||..|| ||    |||||||||   ||||
MT066156.1     26492 AGTTTTTCTGTTTGGAACTTTAATTTTAGC----CATGGCAGATTCCAAC  26537

AY323977.2     26410 GGTACTATTACCGTTGAGGAGCTTAAACAACTCCTGGAACAATGGAACCT  26459
                     |||||||||||||||||.|||||||||.|.|||||.||||||||||||||
MT066156.1     26538 GGTACTATTACCGTTGAAGAGCTTAAAAAGCTCCTTGAACAATGGAACCT  26587

AY323977.2     26460 AGTAATAGGTTTCCTATTCCTAGCCTGGATTATGTTACTACAATTTGCCT  26509
                     |||||||||||||||||||||..|.||||||....|.|||||||||||||
MT066156.1     26588 AGTAATAGGTTTCCTATTCCTTACATGGATTTGTCTTCTACAATTTGCCT  26637

AY323977.2     26510 ATTCTAATCGGAACAGGTTTTTGTACATAATAAAGCTTGTTTTCCTCTGG  26559
                     ||.|.||..||||.|||||||||||.|||||.|||.|..|||||||||||
MT066156.1     26638 ATGCCAACAGGAATAGGTTTTTGTATATAATTAAGTTAATTTTCCTCTGG  26687

AY323977.2     26560 CTCTTGTGGCCAGTAACACTTGCTTGTTTTGTGCTTGCTGCTGTCTACAG  26609
                     ||.||.|||||||||||..|.|||||||||||||||||||||||.|||||
MT066156.1     26688 CTGTTATGGCCAGTAACTTTAGCTTGTTTTGTGCTTGCTGCTGTTTACAG  26737

AY323977.2     26610 AATTAATTGGGTGACTGGCGGGATTGCGATTGCAATGGCTTGTATTGTAG  26659
                     |||.||||||.|.||.||.||.|||||.||.||||||||||||.||||||
MT066156.1     26738 AATAAATTGGATCACCGGTGGAATTGCTATCGCAATGGCTTGTCTTGTAG  26787

AY323977.2     26660 GCTTGATGTGGCTTAGCTACTTCGTTGCTTCCTTCAGGCTGTTTGCTCGT  26709
                     |||||||||||||.|||||||||.|||||||.|||||.||||||||.|||
MT066156.1     26788 GCTTGATGTGGCTCAGCTACTTCATTGCTTCTTTCAGACTGTTTGCGCGT  26837

AY323977.2     26710 ACCCGCTCAATGTGGTCATTCAACCCAGAAACAAACATTCTTCTCAATGT  26759
                     ||.||.||.||||||||||||||.||||||||.||||||||||||||.||
MT066156.1     26838 ACGCGTTCCATGTGGTCATTCAATCCAGAAACTAACATTCTTCTCAACGT  26887

AY323977.2     26760 GCCTCTCCGGGGGACAATTGTGACCAGACCGCTCATGGAAAGTGAACTTG  26809
                     |||.||||..||.||.|||.|||||||||||||..|.|||||||||||.|
MT066156.1     26888 GCCACTCCATGGCACTATTCTGACCAGACCGCTTCTAGAAAGTGAACTCG  26937

AY323977.2     26810 TCATTGGTGCTGTGATCATTCGTGGTCACTTGCGAATGGCCGGACACTCC  26859
                     |.||.||.|||||||||.|||||||.||..|.||.||.||.||||||...
MT066156.1     26938 TAATCGGAGCTGTGATCCTTCGTGGACATCTTCGTATTGCTGGACACCAT  26987

AY323977.2     26860 CTAGGGCGCTGTGACATTAAGGACCTGCCAAAAGAGATCACTGTGGCTAC  26909
                     |||||.|||||||||||.|||||||||||.|||||.||||||||.|||||
MT066156.1     26988 CTAGGACGCTGTGACATCAAGGACCTGCCTAAAGAAATCACTGTTGCTAC  27037

AY323977.2     26910 ATCACGAACGCTTTCTTATTACAAATTAGGAGCGTCGCAGCGTGTAGGCA  26959
                     |||||||||||||||||||||||||||.|||||.|||||||||||||...
MT066156.1     27038 ATCACGAACGCTTTCTTATTACAAATTGGGAGCTTCGCAGCGTGTAGCAG  27087

AY323977.2     26960 CTGATTCAGGTTTTGCTGCATACAACCGCTACCGTATTGGAAACTATAAA  27009
                     .|||.|||||||||||||||||||..||||||.|.|||||.|||||||||
MT066156.1     27088 GTGACTCAGGTTTTGCTGCATACAGTCGCTACAGGATTGGCAACTATAAA  27137

AY323977.2     27010 TTAAATACAGACCACGCCGGTAGCAACGACAATATTGCTTTGCTAGTACA  27059
                     |||||.||||||||..||.||||||..|||||||||||||||||.|||||
MT066156.1     27138 TTAAACACAGACCATTCCAGTAGCAGTGACAATATTGCTTTGCTTGTACA  27187

AY323977.2     27060 GTAAGTGACAACAGATGTTTCATCTTGTTGACTTCCAGGTTACAATAGCA  27109
                     |||||||||||||||||||||||||.||||||||.||||||||.||||||
MT066156.1     27188 GTAAGTGACAACAGATGTTTCATCTCGTTGACTTTCAGGTTACTATAGCA  27237

AY323977.2     27110 GAGATATTGATTATCATTATGAGGACTTTCAGGATTGCTATTTGGAATCT  27159
                     ||||||||..|.||.||||||||||||||.|...||.|.|||||||||||
MT066156.1     27238 GAGATATTACTAATTATTATGAGGACTTTTAAAGTTTCCATTTGGAATCT  27287

AY323977.2     27160 TGACGTTATAATAAGTTCAATAGTGAGACAATTATTTAAGCCTCTAACTA  27209
                     |||....||.||||.....|||.|.|.|.|.||||.||||.|.||||||.
MT066156.1     27288 TGATTACATCATAAACCTCATAATTAAAAATTTATCTAAGTCACTAACTG  27337

AY323977.2     27210 AGAAGAATTATTCGGAGTTAGATGATGAAGAACCTATGGAGTTARATTAT  27259
                     ||||.||.|||||..|.||||||||.||..||||.||||||.|..||||
MT066156.1     27338 AGAATAAATATTCTCAATTAGATGAAGAGCAACCAATGGAGATTGATTA-  27386

AY323977.2     27260 CCATAAAACGAACATGAAAATTATTCTCTTCCTGACATTGATTGTATTTA  27309
                           |||||||||||||||||||||.|||.||.||.||||...|.|..
MT066156.1     27387 ------AACGAACATGAAAATTATTCTTTTCTTGGCACTGATAACACTCG  27430

AY323977.2     27310 CATCTTGCGAGCTATATCACTATCAGGAGTGTGTTAGAGGTACGACTGTA  27359
                     |..||||.|||||.||||||||.||.|||||||||||||||||.||.|||
MT066156.1     27431 CTACTTGTGAGCTTTATCACTACCAAGAGTGTGTTAGAGGTACAACAGTA  27480

AY323977.2     27360 CTACTAAAAGAACCTTGCCCATCAGGAACATACGAGGGCAATTCACCATT  27409
                     ||..||||||||||||||.|.||.||||||||||||||||||||||||||
MT066156.1     27481 CTTTTAAAAGAACCTTGCTCTTCTGGAACATACGAGGGCAATTCACCATT  27530

AY323977.2     27410 TCACCCTCTTGCTGACAATAAATTTGCACTAACTTGCACTAGCACACACT  27459
                     |||.|||||.|||||.||.|||||||||||.||||||..||||||.||.|
MT066156.1     27531 TCATCCTCTAGCTGATAACAAATTTGCACTGACTTGCTTTAGCACTCAAT  27580

AY323977.2     27460 TTGCTTTTGCTTGTGCTGACGGTACTCGACATACCTATCAGCTGCGTGCA  27509
                     ||||||||||||||.|||||||......|||...|||||||.|.|||||.
MT066156.1     27581 TTGCTTTTGCTTGTCCTGACGGCGTAAAACACGTCTATCAGTTACGTGCC  27630

AY323977.2     27510 AGATCAGTTTCACCAAAACTTTTCATCAGACAAGAGGAGGTTCAACAAGA  27559
                     ||||||||||||||.|||||.|||||||||||||||||.|||||   |||
MT066156.1     27631 AGATCAGTTTCACCTAAACTGTTCATCAGACAAGAGGAAGTTCA---AGA  27677

AY323977.2     27560 GCTCTACTCGCCACTTTTTCTCATTGTTGCTGCTCTAGTATTTTTAATAC  27609
                     .||.|||||.|||.|||||||.||||||||.||..||||.|||.|||.||
MT066156.1     27678 ACTTTACTCTCCAATTTTTCTTATTGTTGCGGCAATAGTGTTTATAACAC  27727

AY323977.2     27610 TTTGCTTCACCATTAAGAGAAAGACAGAATGAATGAGCTCACTTTAATTG  27659
                     ||||||||||..|.||.|||||||||||||||.|||.||..|.|||||||
MT066156.1     27728 TTTGCTTCACACTCAAAAGAAAGACAGAATGATTGAACTTTCATTAATTG  27777

AY323977.2     27660 ACTTCTATTTGTGCTTTTTAGCCTTTCTGCTATTCCTTGTTTTAATAATG  27709
                     ||||||||||||||||||||||||||||||||||||||||||||||.|||
MT066156.1     27778 ACTTCTATTTGTGCTTTTTAGCCTTTCTGCTATTCCTTGTTTTAATTATG  27827

AY323977.2     27710 CTTATTATATTTTGGTTTTCACTCGAAATCCAGGATCTAGAAGAACCTTG  27759
                     ||||||||.||||||||.|||||.|||.|.||.||||...|.|||.||||
MT066156.1     27828 CTTATTATCTTTTGGTTCTCACTTGAACTGCAAGATCATAATGAAACTTG  27877

AY323977.2     27760 TACCAAAGTCTAAACGAACATGAAACTTCTCATTGTTTTGACTTGTATT-  27808
                     |..|   |.||||||||||||||||.|||   |||||||.....|.||.
MT066156.1     27878 TCAC---GCCTAAACGAACATGAAATTTC---TTGTTTTCTTAGGAATCA  27921

AY323977.2     27809 TCTCTA-TGCAGTTGCATATGCAC------TGTAGT--ACAGCGCTGTGC  27849
                     ||.|.| ||.||.|||||.| |||      ||||||  ||||...|||.|
MT066156.1     27922 TCACAACTGTAGCTGCATTT-CACCAAGAATGTAGTTTACAGTCATGTAC  27970

AY323977.2     27850 ATCTAATAAACCTCATGTGCTTGAAGATCCTTGT----------------  27883
                     .....||.||||..||||..||||.||.||.|||
MT066156.1     27971 TCAACATCAACCATATGTAGTTGATGACCCGTGTCCTATTCACTTCTATT  28020

AY323977.2     27884 ----AAGGTACA--ACACTAGGGGTAATACTTATAGCACTGCT--TGGCT  27925
                         |.||||.|  |.|.||||.|..|.|.....|||||...|  |.|..
MT066156.1     28021 CTAAATGGTATATTAGAGTAGGAGCTAGAAAATCAGCACCTTTAATTGAA  28070

AY323977.2     27926 TTGTGC-TCTAGGAAA--GGTTTTA-----CCTTTTCA-TAGATGGCACA  27966
                     |||||| |..|.||..  ||||.||     ||..|||| ||.||.|....
MT066156.1     28071 TTGTGCGTGGATGAGGCTGGTTCTAAATCACCCATTCAGTACATCGATAT  28120

AY323977.2     27967 CTATGGTTCAAACATGCACACCTAA--TGTTACTATCAACTGTCAAGATC  28014
                     |..|..||..|...|...|...|.|  |.||||.||.||.||.||.||.|
MT066156.1     28121 CGGTAATTATACAGTTTCCTGTTTACCTTTTACAATTAATTGCCAGGAAC  28170

AY323977.2     28015 CAGCTGGTGGTGCGCTTATAGCTAGGTGTTGGTACCTTCATGAAGGTCAC  28064
                     |.......|||...|||.|||...|.||||.||.|.   ||||||
MT066156.1     28171 CTAAATTGGGTAGTCTTGTAGTGCGTTGTTCGTTCT---ATGAAG-----  28212

AY323977.2     28065 CAAACTGCTGCATTTAGAGACGTACTTGTTGTTTTAAAT-------AAAC  28107
                      |...|...|..|.|...|||||.|.||||||||||.||       ||||
MT066156.1     28213 -ACTTTTTAGAGTATCATGACGTTCGTGTTGTTTTAGATTTCATCTAAAC  28261

AY323977.2     28108 GAACAAATTAAAATGTCTGATAATGGACCCCAATCAAACCAACGTAGTGC  28157
                     |||||||.|||||||||||||||||||||||||   ||.||.||.|.|||
MT066156.1     28262 GAACAAACTAAAATGTCTGATAATGGACCCCAA---AATCAGCGAAATGC  28308

AY323977.2     28158 CCCCCGCATTACATTTGGTGGACCCACAGATTCAACTGACAATAACCAGA  28207
                     .|||||||||||.||||||||||||.||||||||||||.||.||||||||
MT066156.1     28309 ACCCCGCATTACGTTTGGTGGACCCTCAGATTCAACTGGCAGTAACCAGA  28358

AY323977.2     28208 ATGGAGGACGCAATGGGGCAAGGCCAAAACAGCGCCGACCCCAAGGTTTA  28257
                     ||||||.|||||.||||||..|..|||||||.||.||.||||||||||||
MT066156.1     28359 ATGGAGAACGCAGTGGGGCGCGATCAAAACAACGTCGGCCCCAAGGTTTA  28408

AY323977.2     28258 CCCAATAATACTGCGTCTTGGTTCACAGCTCTCACTCAGCATGGCAAGGA  28307
                     ||||||||||||||||||||||||||.|||||||||||.|||||||||||
MT066156.1     28409 CCCAATAATACTGCGTCTTGGTTCACCGCTCTCACTCAACATGGCAAGGA  28458

AY323977.2     28308 GGAACTTAGATTCCCTCGAGGCCAGGGCGTTCCAATCAACACCAATAGTG  28357
                     .||.||||.||||||||||||.||.|||||||||||.|||||||||||..
MT066156.1     28459 AGACCTTAAATTCCCTCGAGGACAAGGCGTTCCAATTAACACCAATAGCA  28508

AY323977.2     28358 GTCCAGATGACCAAATTGGCTACTACCGAAGAGCTACCCGACGAGTTCGT  28407
                     ||||||||||||||||||||||||||||||||||||||.|||||.|||||
MT066156.1     28509 GTCCAGATGACCAAATTGGCTACTACCGAAGAGCTACCAGACGAATTCGT  28558

AY323977.2     28408 GGTGGTGACGGCAAAATGAAAGAGCTCAGCCCCAGATGGTACTTCTATTA  28457
                     |||||||||||.|||||||||||.|||||.||.||||||||.|||||.||
MT066156.1     28559 GGTGGTGACGGTAAAATGAAAGATCTCAGTCCAAGATGGTATTTCTACTA  28608

AY323977.2     28458 CCTAGGAACTGGCCCAGAAGCTTCACTTCCCTACGGCGCTAACAAAGAAG  28507
                     ||||||||||||.|||||||||..|||||||||.||.|||||||||||.|
MT066156.1     28609 CCTAGGAACTGGGCCAGAAGCTGGACTTCCCTATGGTGCTAACAAAGACG  28658

AY323977.2     28508 GCATCGTATGGGTTGCAACTGAGGGAGCCTTGAATACACCCAAAGACCAC  28557
                     |||||.||||||||||||||||||||||||||||||||||.|||||.|||
MT066156.1     28659 GCATCATATGGGTTGCAACTGAGGGAGCCTTGAATACACCAAAAGATCAC  28708

AY323977.2     28558 ATTGGCACCCGCAATCCTAATAACAATGCTGCCACCGTGCTACAACTTCC  28607
                     ||||||||||||||||||..||||||||||||.|.|||||||||||||||
MT066156.1     28709 ATTGGCACCCGCAATCCTGCTAACAATGCTGCAATCGTGCTACAACTTCC  28758

AY323977.2     28608 TCAAGGAACAACATTGCCAAAAGGCTTCTACGCAGAGGGAAGCAGAGGCG  28657
                     ||||||||||||||||||||||||||||||||||||.||.||||||||||
MT066156.1     28759 TCAAGGAACAACATTGCCAAAAGGCTTCTACGCAGAAGGGAGCAGAGGCG  28808

AY323977.2     28658 GCAGTCAAGCCTCTTCTCGCTCCTCATCACGTAGTCGCGGTAATTCAAGA  28707
                     |||||||||||||||||||.||||||||||||||||||...|.|||||||
MT066156.1     28809 GCAGTCAAGCCTCTTCTCGTTCCTCATCACGTAGTCGCAACAGTTCAAGA  28858

AY323977.2     28708 AATTCAACTCCTGGCAGCAGTAGGGGAAATTCTCCTGCTCGAATGGCTAG  28757
                     |||||||||||.||||||||||||||||.||||||||||.||||||||.|
MT066156.1     28859 AATTCAACTCCAGGCAGCAGTAGGGGAACTTCTCCTGCTAGAATGGCTGG  28908

AY323977.2     28758 CGGAGGTGGTGAAACTGCCCTCGCGCTATTGCTGCTAGACAGATTGAACC  28807
                     |...||.|||||..||||.||.||..|..|||||||.|||||||||||||
MT066156.1     28909 CAATGGCGGTGATGCTGCTCTTGCTTTGCTGCTGCTTGACAGATTGAACC  28958

AY323977.2     28808 AGCTTGAGAGCAAAGTTTCTGGTAAAGGCCAACAACAACAAGGCCAAACT  28857
                     ||||||||||||||.|.|||||||||||||||||||||||||||||||||
MT066156.1     28959 AGCTTGAGAGCAAAATGTCTGGTAAAGGCCAACAACAACAAGGCCAAACT  29008

AY323977.2     28858 GTCACTAAGAAATCTGCTGCTGAGGCATCTAAAAAGCCTCGCCAAAAACG  28907
                     ||||||||||||||||||||||||||.|||||.||||||||.||||||||
MT066156.1     29009 GTCACTAAGAAATCTGCTGCTGAGGCTTCTAAGAAGCCTCGGCAAAAACG  29058

AY323977.2     28908 TACTGCCACAAAACAGTACAACGTCACTCAAGCATTTGGGAGACGTGGTC  28957
                     |||||||||.|||...|||||.||.||.|||||.||.||.||||||||||
MT066156.1     29059 TACTGCCACTAAAGCATACAATGTAACACAAGCTTTCGGCAGACGTGGTC  29108

AY323977.2     28958 CAGAACAAACCCAAGGAAATTTCGGGGACCAAGACCTAATCAGACAAGGA  29007
                     ||||||||||||||||||||||.||||||||.||.|||||||||||||||
MT066156.1     29109 CAGAACAAACCCAAGGAAATTTTGGGGACCAGGAACTAATCAGACAAGGA  29158

AY323977.2     29008 ACTGATTACAAACATTGGCCGCAAATTGCACAATTTGCTCCAAGTGCCTC  29057
                     ||||||||||||||||||||||||||||||||||||||.||.||.||.||
MT066156.1     29159 ACTGATTACAAACATTGGCCGCAAATTGCACAATTTGCCCCCAGCGCTTC  29208

AY323977.2     29058 TGCATTCTTTGGAATGTCACGCATTGGCATGGAAGTCACACCTTCGGGAA  29107
                     .||.|||||.||||||||.|||||||||||||||||||||||||||||||
MT066156.1     29209 AGCGTTCTTCGGAATGTCGCGCATTGGCATGGAAGTCACACCTTCGGGAA  29258

AY323977.2     29108 CATGGCTGACTTATCATGGAGCCATTAAATTGGATGACAAAGATCCACAA  29157
                     |.|||.||||.||....||.|||||.|||||||||||||||||||||.|.
MT066156.1     29259 CGTGGTTGACCTACACAGGTGCCATCAAATTGGATGACAAAGATCCAAAT  29308

AY323977.2     29158 TTCAAAGACAACGTCATACTGCTGAACAAGCACATTGACGCATACAAAAC  29207
                     ||||||||..|.|||||..|||||||.|||||.|||||||||||||||||
MT066156.1     29309 TTCAAAGATCAAGTCATTTTGCTGAATAAGCATATTGACGCATACAAAAC  29358

AY323977.2     29208 ATTCCCACCAACAGAGCCTAAAAAGGACAAAAAGAAAAAGACTGATGAAG  29257
                     ||||||||||||||||||||||||||||||||||||.|||.||||||||.
MT066156.1     29359 ATTCCCACCAACAGAGCCTAAAAAGGACAAAAAGAAGAAGGCTGATGAAA  29408

AY323977.2     29258 CTCAGCCTTTGCCGCAGAGACAAAAGAAGCAGCCCACTGTGACTCTTCTT  29307
                     ||||..|.||.|||||||||||.|||||.||||..|||||||||||||||
MT066156.1     29409 CTCAAGCCTTACCGCAGAGACAGAAGAAACAGCAAACTGTGACTCTTCTT  29458

AY323977.2     29308 CCTGCGGCTGACATGGATGATTTCTCCAGACAACTTCAAAATTCCATGAG  29357
                     |||||.||.||..|||||||||||||||.||||.|.|||.|.||||||||
MT066156.1     29459 CCTGCTGCAGATTTGGATGATTTCTCCAAACAATTGCAACAATCCATGAG  29508

AY323977.2     29358 TGGAGCTTCTGCTGATTCAACTCAGGCATAAACACTCATGATGACCACAC  29407
                     ..|      ||||||.|||||||||||.|||||  |||||..||||||||
MT066156.1     29509 CAG------TGCTGACTCAACTCAGGCCTAAAC--TCATGCAGACCACAC  29550

AY323977.2     29408 AAGGCAGATGGGCTATGTAAACGTTTTCGCAATTCCGTTTACGATACATA  29457
                     ||||||||||||||||.|||||||||||||..||||||||||||||.|||
MT066156.1     29551 AAGGCAGATGGGCTATATAAACGTTTTCGCTTTTCCGTTTACGATATATA  29600

AY323977.2     29458 GTCTACTCTTGTGCAGAATGAATTCTCGTAACTAAACAGCACAAGTAGGT  29507
                     ||||||||||||||||||||||||||||||||||.|.|||||||||||.|
MT066156.1     29601 GTCTACTCTTGTGCAGAATGAATTCTCGTAACTACATAGCACAAGTAGAT  29650

AY323977.2     29508 TTAGTTAACTTTAATCTCACATAGCAATCTTTAATCAATGTGTAACATTA  29557
                     .||||||||||||||||||||||||||||||||||||.||||||||||||
MT066156.1     29651 GTAGTTAACTTTAATCTCACATAGCAATCTTTAATCAGTGTGTAACATTA  29700

AY323977.2     29558 GGGAGGACTTGAAAGAGCCACCACATTTTCATCGAGGCCACGCGGAGTAC  29607
                     |||||||||||||||||||||||||||||||.||||||||||||||||||
MT066156.1     29701 GGGAGGACTTGAAAGAGCCACCACATTTTCACCGAGGCCACGCGGAGTAC  29750

AY323977.2     29608 GATCGAGGGTACAGTGAATAATGCTAGGGAGAGCTGCCTATATGGAAGAG  29657
                     |||||||.||||||||||.|||||||||||||||||||||||||||||||
MT066156.1     29751 GATCGAGTGTACAGTGAACAATGCTAGGGAGAGCTGCCTATATGGAAGAG  29800

AY323977.2     29658 CCCTAATGTGTAAAATTAATTTTAGTAGTGCTATCCCCATGTGATTTTAA  29707
                     ||||||||||||||||||||||||||||||||||||||||||||||||||
MT066156.1     29801 CCCTAATGTGTAAAATTAATTTTAGTAGTGCTATCCCCATGTGATTTTAA  29850

AY323977.2     29708 TAGCTTCTTAGGAGAATGAC  29727
                     |||||||||||||||||
MT066156.1     29851 TAGCTTCTTAGGAGAAT---  29867


#---------------------------------------
#---------------------------------------
```
